# SUMOylation of rice DELLA SLR1 modulates transcriptional responses and improves yield under salt stress

**DOI:** 10.1101/2020.03.10.986224

**Authors:** Nuno M. Gonçalves, Telma Fernandes, Cátia Nunes, Margarida T. G. Rosa, Cleverson C. Matiolli, Mafalda A. A. Rodrigues, Pedro M. Barros, M. Margarida Oliveira, Isabel A. Abreu

## Abstract

DELLA proteins modulate GA signalling and are major regulators of plant plasticity to endure stress. DELLAs are mostly regulated at the post-translational level, and their activity relies on the interaction with upstream regulators and transcription factors (TFs). SUMOylation is a post-translational modification (PTM) capable of changing protein interaction and found to influence DELLA activity in Arabidopsis. We determined that SUMOylation of the single rice DELLA SLENDER RICE1 (SLR1) occurs in a lysine residue different from the one previously identified in Arabidopsis REPRESSOR OF GA (RGA). Remarkably, artificially increasing SUMOylated SLR1 (SUMO1SLR1) levels attenuated the penalty of salt stress on plant yield. Gene expression analysis revealed that the overexpression of SUMOylated SLR1 regulates key dioxygenases that modulate active GA levels, namely *GA20ox2* and *GA2ox3*, which could partially explain the sustained productivity upon salt stress imposition. Besides, SLR1 SUMOylation blocked the interaction with the growth regulator YAB4, which may fine-tune *GA20ox2* expression. Mechanistically, we propose that SLR1 SUMOylation disrupts the interaction with members of several transcription factor families to modulate gene expression. We found that SLR1 SUMOylation represents a novel mechanism modulating DELLA activity, which attenuates the impact of stress on plant performance.

**One sentence summary:** Rice plants show increased yield under salt stress when its gibberellin transcriptional regulator DELLA protein is artificially SUMOylated.

## INTRODUCTION

Salinity is one of the most serious threats to food security, affecting 20% of irrigated soils worldwide. Rice (*Oryza sativa* L.) is a crop with high sensitivity to salinity compared to other monocot crops, such as wheat and barley (Munns and Tester, 2008). Plant sessile nature requires rapid physiological responses to cope with stress, which is partly achieved through a complex crosstalk between hormone signalling pathways affecting growth, flowering, and defence against environmental challenges (Vanstraelen and Benková, 2012). Gibberellins (GA) are diterpenoid hormones involved in the regulation of several developmental processes, including seed germination, stem elongation, and flowering (Hirano et al., 2008). GA responses are modulated by DELLA proteins, which act as transcriptional regulators (Hedden and Thomas, 2012). In the presence of GA, DELLAs are targeted to ubiquitin-mediated proteasome degradation by the GA-receptor GA-INSENSITIVE DWARF1 (GID1) through binding to specific DELLA domains (DELLA/TVHYNP), followed by DELLA polyubiquitination mediated by the E3-ligase GA-INSENSITIVE DWARF2/SLEEPY1 (GID2/SLY1). When GA levels decrease, DELLA proteins accumulate and interact with key transcriptional regulators to restrain growth (Hirano et al., 2010; 2008; Ueguchi-Tanaka et al., 2008). For instance, DELLAs can sequester the transcription factors PHYTOCHROME INTERACTING FACTOR3 and 4 (PIF3 and PIF4), in addition to negatively regulate PIF3 protein abundance, mediating the crosstalk of light and GA signalling to modulate cell growth (de Lucas et al., 2008; Li et al., 2016). DELLA transcriptional regulation of GA biosynthetic genes responsible for hormonal homeostasis also potentiates growth arrest. The single rice DELLA SLENDER RICE1 (SLR1) interacts with YAB4 transcription factor to prevent the induction of *GA20ox2*, which encodes a dioxygenase that is essential for the synthesis of bioactive GA (Yang et al., 2016). In Arabidopsis, DELLA interaction with GAI-ASSOCIATED FACTOR1 (GAF1) TF triggers a negative-feedback loop that activates GA biosynthesis and subsequent DELLA degradation (Fukazawa et al., 2017). The increasing number of validated DELLA interactors showcase an overlap with most hormone pathways, impacting development, fertility, and seed number (Davière and Achard, 2015; Van De Velde et al., 2017).

DELLA activity is mostly modulated through post-translational modifications (PTMs), which impact its ability to interact with transcriptional regulators and modulate gene expression (Van De Velde et al., 2017). Besides ubiquitination and subsequent proteasome-mediated degradation, there are other PTMs that influence the activity of the DELLA’s in response to GA signalling. For instance, the phosphorylation by kinase EARLY FLOWERING 1 (EL1) regulates SLR1 protein stability and activity (Dai and Xue, 2010; Itoh et al., 2005). Recently, a mechanism of antagonistic PTMs was proposed to regulate Arabidopsis DELLA’s interaction with specific protein partners. *O*-fucosylation by SPINDLY (SPY) promotes DELLA interaction with essential growth regulators – such as BRASSINOLE-RESISTANT1 (BZR1), PIF3 and PIF4 – leading to growth arrest (Zentella et al., 2017). Conversely, RGA *O*-GlcNAcylation by SECRET AGENT (SEC) blocks these interactions and promotes GA-induced growth resumption (Camut et al., 2017; Zentella et al., 2016). DELLA SUMOylation has also been indicated as a PTM controlling the stability of REPRESSOR OF GA (RGA), one of the Arabidopsis five DELLA proteins. SUMO attachment to RGA hinders its ubiquitin-mediated proteasome degradation through blockage of GID1-RGA interaction in a GA-independent manner (Conti et al., 2014). The stabilization of Arabidopsis DELLA by SUMOylation also positively affected salt stress response, in line with observations that DELLA accumulation leads to increased salinity tolerance (Achard et al., 2006; Conti et al., 2014; Nelis et al., 2015). SUMOylation is a eukaryotic ubiquitination-like process that involves the processing, activation, conjugation, and ligation of mature SMALL UBIQUITIN-LIKE MODIFIER (SUMO) to acceptor lysine’s (K) in the target protein (Miura et al., 2007). Unlike ubiquitination, SUMO-modifications often occur in a canonical consensus motif (ψ-<u>K</u>-X-E/D), which can be used for *in silico* prediction of SUMOylation targets (Xue et al., 2006). SUMOylation regulates protein stability, activity, and interactions with specific protein partners (Cremona et al., 2012). SUMO deconjugation (deSUMOylation) from target proteins is performed by deSUMOylating proteases (DSPs), with some members recently reported to be specific for poly-SUMO chain processing (Eckhoff and Dohmen, 2015; Yates et al., 2016; Augustine et al., 2018). In rice, reduced levels of SUMO conjugates due to DSP activity of OVERTOLERANT TO SALT1/2 (OTS1/OTS2) have been connected to increased salt tolerance (Srivastava et al., 2016a; 2016b). SUMOylation of specific TFs, such as the stress-induced basic LEUCINE ZIPPER 23 (bZIP23), was suggested to be crucial to drought tolerance and fertility (Srivastava et al., 2017).

Few studies have addressed the biological function of SUMOylation of stress-induced targets. Moreover, most of the studies investigating the effects of PTMs on DELLA function were done in Arabidopsis. This dicot model plant encodes five DELLAs in the genome and essentially differs from monocots, which harbour a single DELLA gene. This study demonstrates that SUMOylation of the single rice DELLA SLR1 impacts the transcriptional output in response to salt stress, possibly through modulation of SLR1 interaction with transcription factors regulating transcriptional patterns governing plant growth and stress responses. We suggest that SLR1 SUMOylation affects its interaction with key transcription factors that impacts GA homeostasis (YAB4), ABA signalling (bZIP23), and JA signalling (bHLH089/94). Additionally, the evidence provided suggests that it is possible to uncouple stress response from decreased plant performance by manipulating the levels of SUMOylated SLR1.

## RESULTS

### SLR1 SUMOylation *in vitro* analysis suggests diversification of DELLA regulation by SUMO in plants

SUMOylation of lysine 65 (K65) in Arabidopsis DELLA RGA modulates its stability in a GA-independent manner (Conti et al., 2014). To determine rice SLR1 SUMOylation, we started by performing an *in silico* prediction of SUMOylation motifs using SUMOsp2 software (Xue et al., 2006) in rice DELLA SLR1 and Arabidopsis DELLA RGA protein sequences. We sought for putative conserved SUMOylation motifs in both proteins and found a low probability SUMOylation site which corresponds to Arabidopsis RGA (Figure 1A). The SUMO acceptor lysine (K65) in RGA is part of a canonical SUMOylation motif (LKLE). In contrast, its counterpart in SLR1(K60) is not part of a canonical motif (QKLE), nor are the equivalent ones in another Arabidopsis or monocot DELLAs (Supplemental Figure S1). Nevertheless, both SLR1 and RGA have a high probability SUMOylation site on lysine 2 (K2).

**Figure 1.**
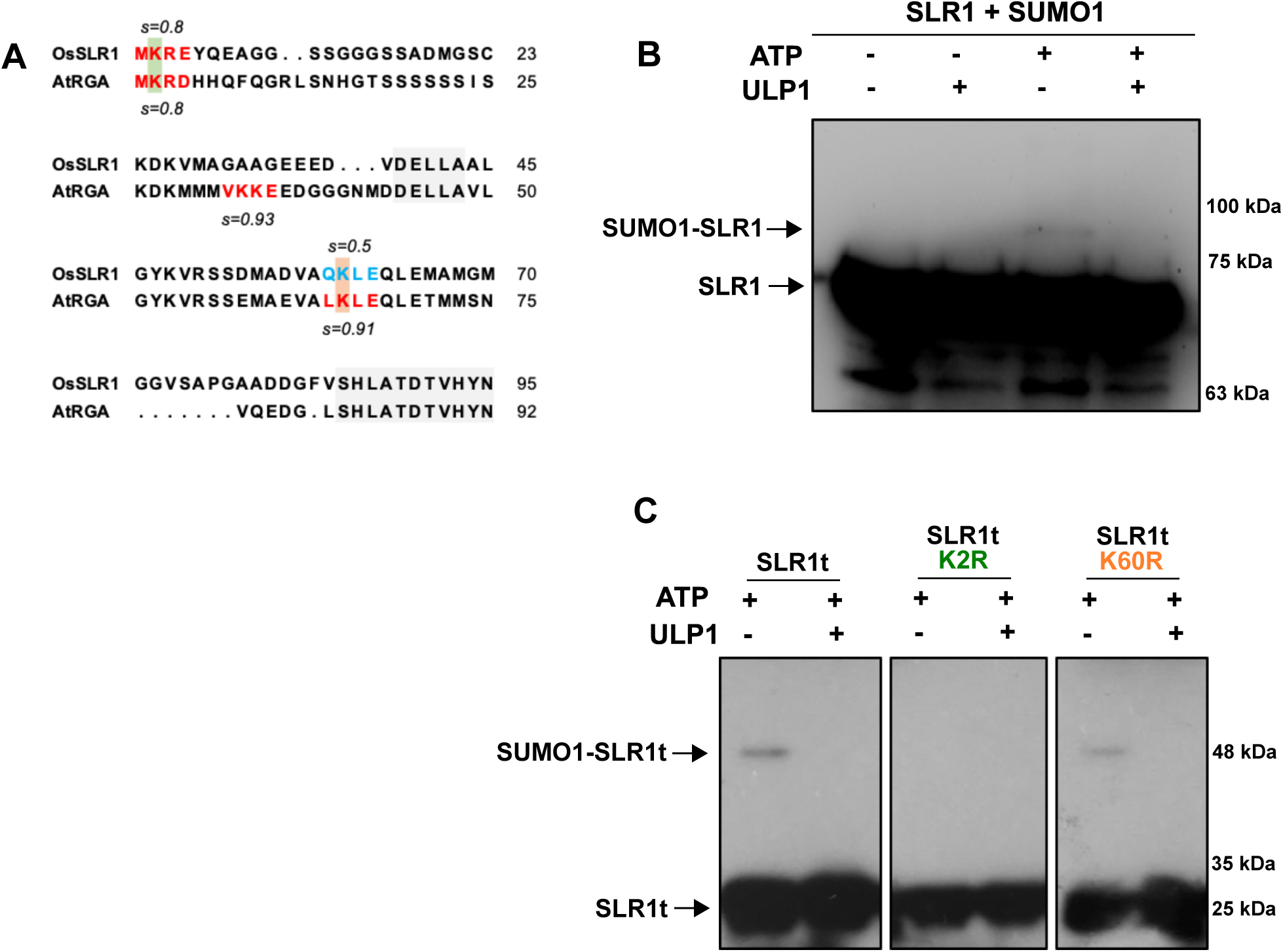
SLR1 is SUMOylated *in vitro* in a novel SUMO-conjugation canonical motif. **(A)** *in silico* prediction of SLR1 canonical motifs (ψ-K-X-E) in the N-terminal region as scored by SUMOsp2 software (Xue et al., 2006), compared to *Arabidopsis thaliana* RGA corresponding region. Motifs showing high SUMOylation probability are marked in red, and low probability motifs are marked in blue. SLR1 K2 and K60 are shown in green and orange, respectively. **(B-C)** *in vitro* SUMOylation assays using rice SUMOylation machinery (OsSCE1, OsSAE1/2, and OsSUMO1) with recombinant SLR1 and SLR1t (SLR1 N-terminal truncated) proteins, treated with ULP1 SUMO protease, as indicated. SLR1 and SLR1t were detected by immunoblotting using custom made anti-SLR1 antibody, where **(B)** corresponds to the full-length SLR1, and **(C)** to the SLR1t coding the leading 190 first SLR1 residues, as well as the mutated isoforms SLR1t K2R and SLR1t K60R.

To verify if the *in silico* prediction of SUMO attachment to SLR1 holds true, we performed an *in vitro* SUMOylation assay targeting both full-length SLR1 and a truncated isoform containing the leading 190 amino acids (SLR1t). Importantly, SLR1t includes the N-terminal domains (DELLA and TVHYNP) responsible for GA-mediated modulation of DELLA stability and SLR1 transactivation properties (Davière and Achard, 2015; Marín-de la Rosa et al., 2015; Hirano et al., 2012), as well as the predicted SUMOylation sites depicted in the Figure 1A. We observed a mobility-shift of both full-length SLR1 (Figure 1B) and truncated SLR1t (Figure 1C; with both SUMO1 and SUMO2 – Supplemental Figure S2A-B), consistent with ATP-dependent SUMO-conjugation in the presence of the rice SUMOylation machinery. The band resulting from *in vitro* SLR1 and SLR1t SUMOylation disappeared upon treatment with recombinant SUMO-specific protease ULP1 that cleaves SUMO from its conjugates (Li and Hochstrasser, 2003), confirming that the observed mobility-shift was caused by attached SUMO (Figure 1B-C). To analyse the contribution of each predicted SUMO acceptor lysine in SLR1, we performed directed mutagenesis changing the lysines K2 and K60 to arginine residues, which cannot be conjugated to SUMO (K2R and K60R). The mutated SLR1t K60R isoform retained the ability to be SUMOylated – which was confirmed by the band disappearance after SUMO proteolytic removal by ULP1 treatment – while K2R did not (Figure 1C). The result shows that K2 is the preferential SUMOylated residue in SLR1, suggesting that SUMO attachment to DELLA may occurs in distinct lysine residues of Arabidopsis RGA (K65) and rice SLR1 (K2). SUMOylation in different sites potentially results in different outcomes regarding protein interactions and activity of SUMOylated SLR1 compared to SUMOylated RGA. However, the *in silico* prediction of SUMOylation sites suggests that RGA K2 could also be SUMOylated (Figure 1A). We need further studies to clarify if K2 SUMOylation occurs in Arabidopsis RGA, or even in DELLA proteins from other species.

### SUMO protease activity analyses provide evidence of *in vivo* SLR1 SUMOylation

To demonstrate the *in vivo* SUMOylation of SLR1, we treated non-denatured rice protein extracts with the ULP1 SUMO protease. The procedure would allow for indirect detection of SUMOylated SLR1 by analysing the increase in non-SUMOylated SLR1 protein amount resulting from ULP1 treatment, in comparison to SUMOylated SLR1. We observed an increase of non-SUMOylated SLR1 upon the addition of ULP1, as detected by the anti-SLR1 antibody in protein extracts (Figure 2A). The result is in line with the previous *in silico* and *in vitro* analyses indicating that SLR1 could be SUMOylated. Our results also suggest that SLR1 mono-SUMOylation may be much less abundant than other forms of SUMOylation since we could not detect an SLR1 band with the expected mobility shift for mono-SUMOylation (Figure 1B-C). To further support the evidence of *in vivo* SLR1 SUMOylation, we tested the levels of SUMOylated SLR1 in knockout mutant lines for genes encoding rice SUMO proteases, namely *els1* and *fug1* (Figure 2B-C). ELS1 and FUG1 are active rice SUMO proteases, as both knockout lines contain increased levels of SUMOylated proteins (Rosa et al., 2018). We treated protein extracts from *els1* and *fug1* homozygous lines with ULP1 and compared their total SUMOylated SLR1 levels with those present in negative segregant lines (NS). The results show that although the overall level of non-SUMOylated SLR1 is not significantly affected in either one of the KO lines when compared to their negative segregant lines (Figure 2B), the levels of SUMOylated SLR1 are increased in *fug1* mutant (Figure 2C). These results demonstrate SUMOylation of SLR1 occurs *in vivo*, as indicated by the processing (*i.e.*, switch from SUMO-SLR1 to SLR1) mediated by FUG1. Interestingly, it also suggests that there is a certain degree of specificity for SLR1 deSUMOylation, as suggested by SUMO cleavage by FUG1 and not ELS1 (Figure 2C). Altogether, the evidence supports that SLR1 is SUMOylated *in planta* and that SUMO specific proteases could regulate the levels of SUMOylated SLR1.

**Figure 2.**
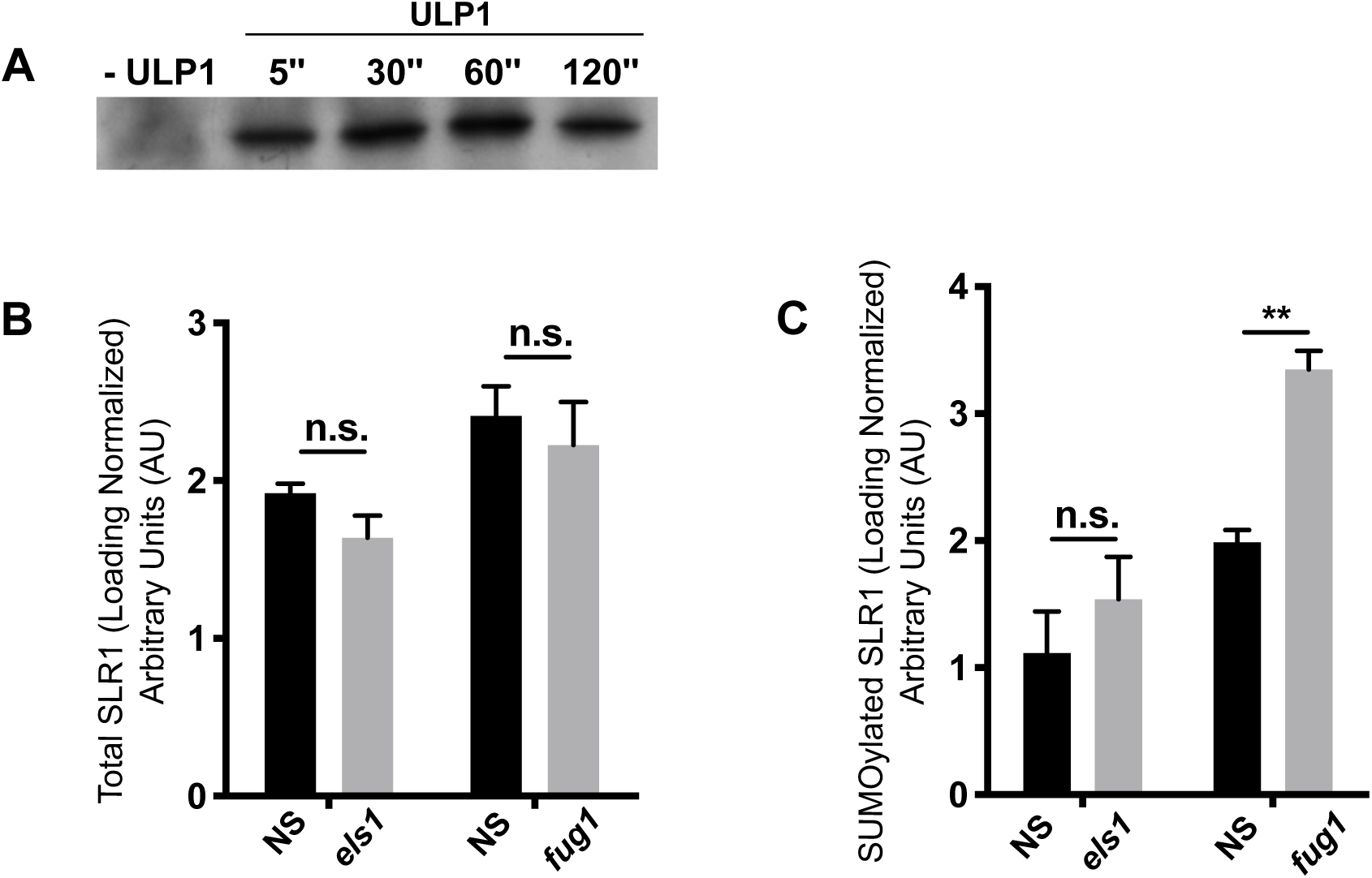
SUMO-specific protease activity reveal SLR1 *in vivo* regulation by SUMO. **(A)** *Oryza sativa* L. cv Nipponbare non-denaturing protein extracts treated with *Saccharomyces cerevisiae* Ulp1 (ULP1) for SLR1 deSUMOylation analysis as detected by immunoblot using anti-SLR1 antibody. **(B)** Loading-normalized relative quantification of total SLR1 protein in *els1* and *fug1* mutants, and the respective negative segregants (NS) after deSUMOylation by ULP1 treatment; and **(C)** total SUMOylated SLR1, as shown by ratio comparison to non-SUMOylated protein after ULP1 treatment. Error bars correspond to standard error (SE; n=4). Statistically significant values are represented as **(p<0.01). The significance of statistics was asserted by t-test with Holm-Sidak method correction for multiple comparisons.

### Overexpression of SUMOylated SLR1 increases salt tolerance

Because DELLA protein accumulation was previously connected with stress tolerance (Achard et al., 2006), we assessed SLR1 expression in response to salt stress. Wild-type seedlings accumulated SLR1 protein in shoots during the first 8 hours of stress (Supplemental Figure S3A). On the other hand, *SLR1* transcript levels decreased, which is consistent with the previously reported DELLA negative feedback-loop regulation on its own transcript abundance (Hedden & Thomas, 2012).

To gain insight on the role of SLR1 SUMOylation in rice development and responses to salt stress, we generated transgenic lines overexpressing SLR1 (SLR1-OX), as well as lines constitutively expressing a mimicry of the SUMOylated form of SRL1 (SUMO1SLR1-OX) originated from the genetic fusion of mature SUMO1 to the K2 residue, which is the SUMOylated lysine of SLR1 (SUMO1_GG_::SLR1_K2_) (Figure 3A). The strategy of using SUMOylated mimetic isoforms originated by gene fusion with the SUMO coding sequence has been used in several biological systems, including plants (Crozet et al., 2015; Ulrich, 2008). We confirmed the overexpression of non-SUMOylated SLR1 isoforms in both SLR1-OX and SUMO1SLR1, with the latter also showing a band with the expected mobility-shift of SLR1 corresponding to mono-SUMOylation (Figure 3B). We also observed an increase in the amount of non-SUMOylated SLR1 protein in SUMO1SLR1-OX compared to WT and NS lines, which could be arising from the processing by endogenous SUMO proteases because the fusion protein retains the SUMO diglycine protease cleavage motif (-GGK-).

**Figure 3.**
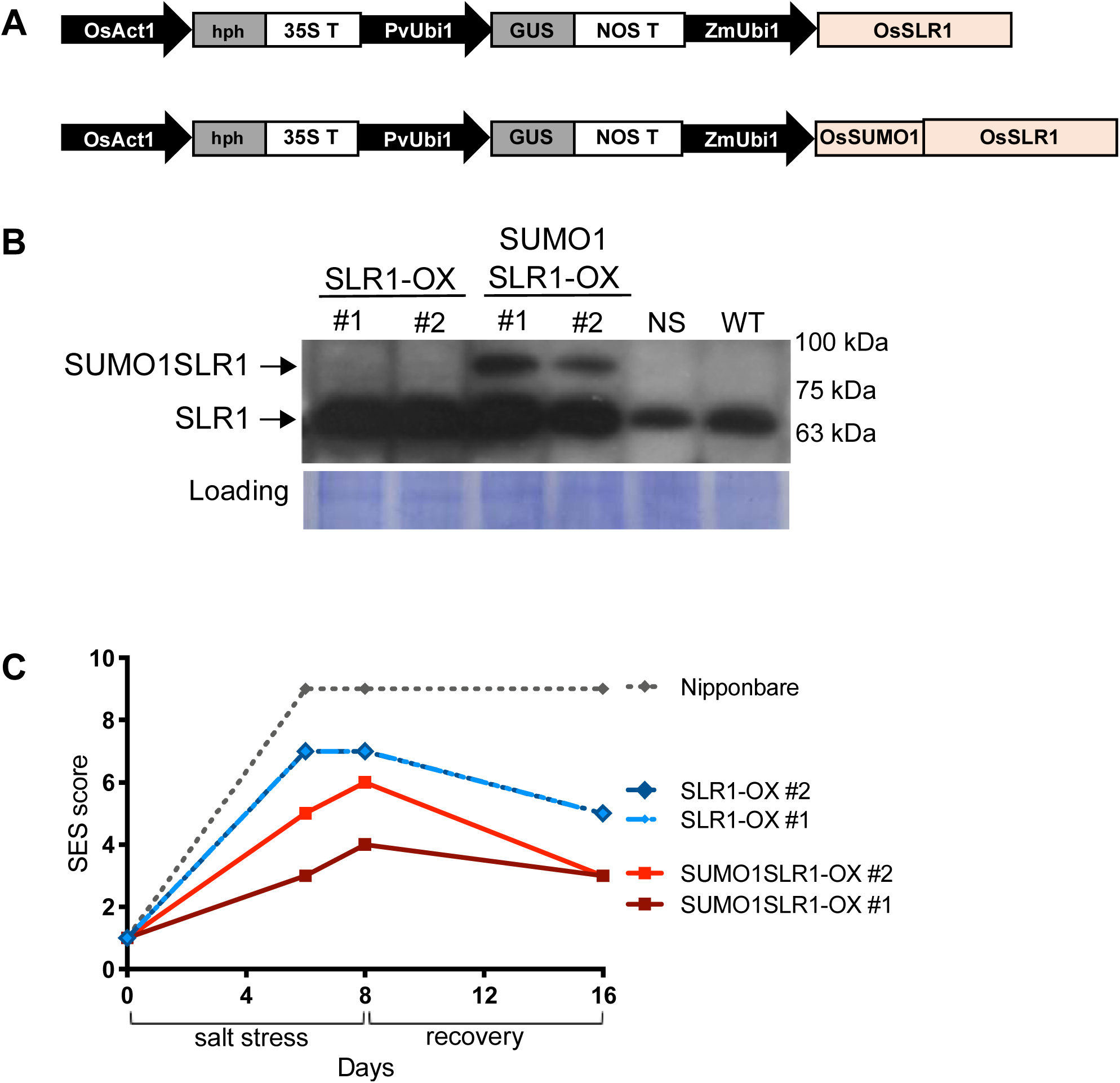
SLR1 SUMOylation confers salt tolerance at the seedling stage. **(A)** Depiction of gene constructs used to generate rice SLR1-OX and SUMO1SLR1-OX transgenic lines. *Zea mays* Ubi1 (ZmUbi1) promoter drives the expression of both OsSLR1 (SLR1-OX) and OsSLR1-OsSUMO1 gene fusion (SUMO1SLR1-OX). **(B)** SLR1 and SUMO1SLR1 protein accumulation in T2 independent transgenic lines (#1 and #2) of SUMO1SLR1-OX and SLR1-OX as detected by immunoblot using custom made anti-SLR1 antibody. **(C)** Stress severity as evaluated by IRRI salinity trial SES score for all lines in 6 and 8 days of stress and after 8 days of recovery period.

Rice is particularly sensitive to salt at both seedling and booting stages (Gregorio et al., 1997). Therefore, we analysed stress marker symptoms of SLR1-OX and SUMO1SLR1-OX lines subjected to high salt concentrations at seedling stage, as well as yield parameters on salt treatments at booting stage. Evaluation of symptoms in seedlings subjected to severe salinity (120mM NaCl) for 8 days showed that both SUMO1SLR1-OX lines performed better under salt stress, displaying decreased symptoms severity according to SES score (Standard Evaluation System for Rice) by IRRI (International Rice Research Institute) (Gregorio et al., 1997), including minimal leaf tip rolling and increased survival (Figure 3C). Importantly, salt tolerance could be correlated with SUMOylated SLR1 levels, as SUMO1SLR1-OX #1 – showing higher SLR1 SUMOylation levels compared to SUMO1SLR1-OX #2 – performed better than the latter (Figure 3B-C). Interestingly, salt-induced SUMOylated SLR1 increased alongside with the non-SUMOylated SLR1 in SUMO1SLR1-OX lines (Supplemental Figure S3B), suggesting that SUMO cleavage processing could be occurring alongside the duration of salt stress. Plants overexpressing SUMO1SLR1 and SLR1-OX also recovered better than the wild type Nipponbare in the 8 following days to salt stress cessation. Thus, mild overexpression of SLR1 can increase rice tolerance to salinity, and this effect is increased by its SUMOylation.

We next tested rice performance under salt stress at the booting stage. Panicle initiation was the cue for mild salt stress imposition (80 mM NaCl), as at this developmental stage pollen viability is most affected by high salinity (Negrão et al., 2011; Sarhadi et al., 2012). Upon measurement of agronomical parameters, we did not detect stress-induced changes in plant height (Figure 4A), tillering or panicle length (Supplemental Figure S4) when compared to the control. However, two critically important parameters for rice yield – namely fertility (filled grains) and productivity (grain weight) – were significantly improved in SUMO1SLR1-OX lines under mild salt stress. Both SLR1-OX and Nipponbare presented around 10% decrease in fertility when comparing to mock conditions, while SUMO1SLR1-OX showed the same fertility in both salt and mock treatments (Figure 4B). This tendency was also observed in grain weight, where SUMO1SLR1-OX presented higher tolerance to mild salt stress regarding productivity compared to Nipponbare and SLR1-OX (Figure 4C). Although the SUMO1SLR1-OX lines had lower productivity under mock conditions compared to Nipponbare, they showed improved performance under salt stress. These results substantiate the significance of SUMOylation of SLR1 in salt stress tolerance acquisition and yield protection observed in two sensitive rice developmental stages.

**Figure 4.**
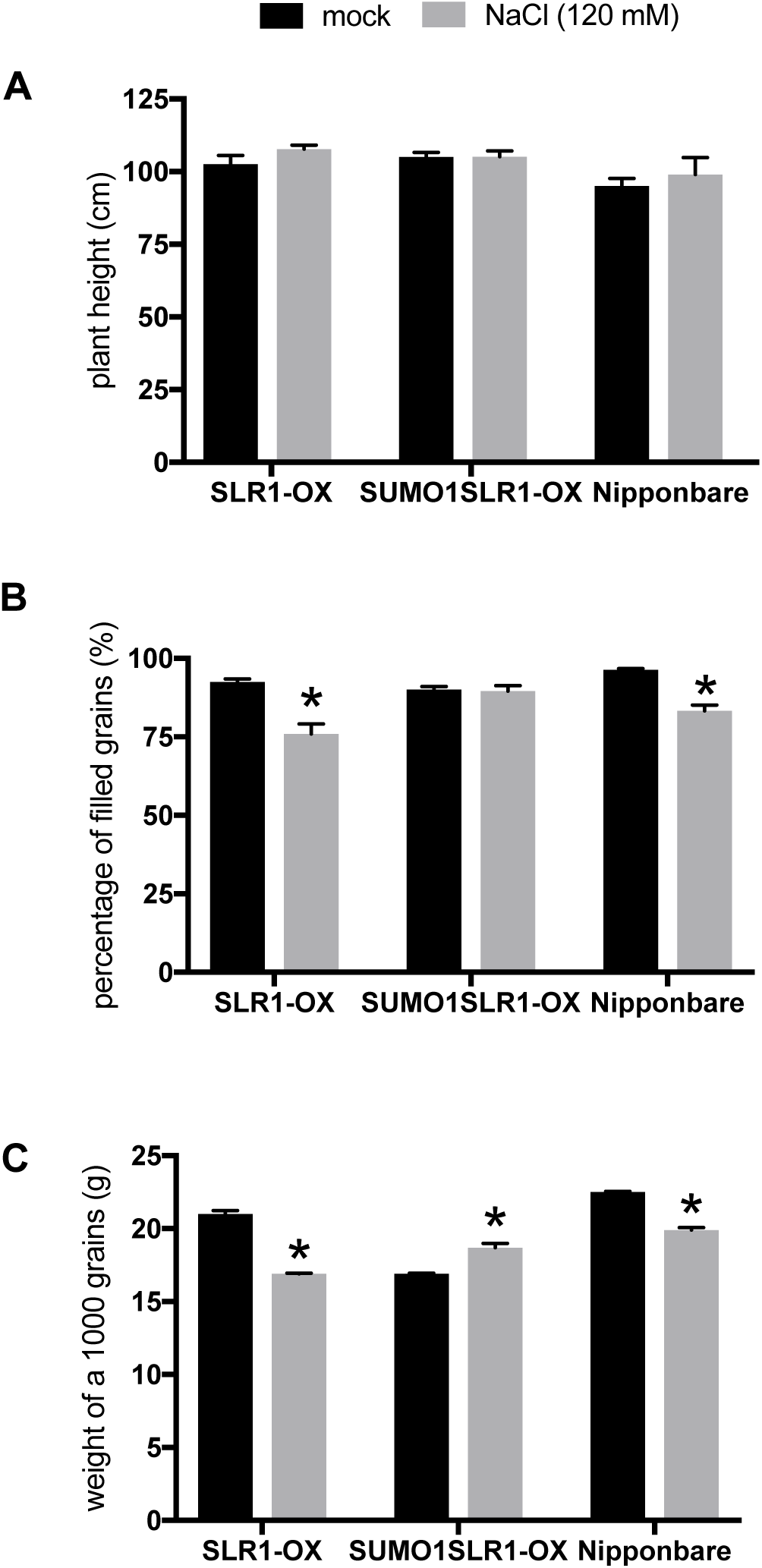
SLR1 SUMOylation reduces the impact of salt stress on yield. Agronomic parameters as measured at the end of rice life cycle: **(A)** plant height (N=6) **(B)** fertility as evaluated by the number of filled grains ratio to total panicle flowers (n>10) and **(C)** productivity as evaluated by the weight of 1000 grains (n>10). Error bars correspond to standard error (SE; 2 independent biological replicates with N=6 each) and statistically significant p-values represented as *(p<0.05). The significance of statistics was asserted by t-test with Holm-Sidak method correction for multiple comparisons.

### SLR1 SUMOylation regulates GA biosynthetic pathway transcripts in response to salt stress

The salt tolerance observed in SUMO1SLR1-OX lines could not be attributed to the increased accumulation of SLR1, as SLR1 protein levels were similar in the SLR1-OX and SUMO1SLR1-OX transgenic lines (Figure 3B). Thus, we inferred that the observed phenotypes in SUMO1SLR1-OX lines could be a consequence of the increased levels of SUMOylated SLR1. Because DELLA proteins regulate transcription, we performed a genome-wide expression analysis to identify differentially regulated genes in SLR1-OX and SUMO1SLR1-OX that could explain the phenotypical differences observed in these lines in response to salt stress. We exposed SLR1-OX, SUMO1SLR1-OX, and Nipponbare genetic background to high salt concentrations (120 mM NaCl) during 8 hours at the seedling stage (2 weeks old), and collected samples for subsequent RNA-seq analysis (Supplemental Table S1). We selected the lines SLR1-OX #2 and SUMO1SLR1-OX #2 for the RNA-seq assay because they showed similar non-SUMOylated SLR1 levels, as we aimed to tackle the effect of SLR1 SUMOylation on gene expression. Both overexpressor lines showed an approximately 1.8-fold increase in SLR1 transcript levels when compared to Nipponbare, which is in line with the levels validated by real-time qPCR for the same samples, *ca.* 1.5-fold increase (Supplemental Figure S5), and with protein levels assessed by immunoblotting (Figure 3B). The induction of salt-induced genes *Rab21* and *P5CS* was used as a transcriptional readout of the salt stress treatment efficiency (Supplemental Figure S5).

Principal Component Analysis (PCA) revealed a clear separation of samples by treatment, this factor being responsible for 47% of the variability in gene expression, while genotype-specific transcriptomic changes explained 21% of the observed variability (Supplemental Figure S6). The constitutive overexpression of SLR1 and SUMO1SLR1 in absence of salt stress resulted in differential regulation of 247 and 207 genes, respectively (Supplemental Tables S1.6 to S1.8). Functional annotation of SLR1-OX DEGs showed enrichment of gene ontology (GO) terms associated with diterpenoid biosynthesis, highlighting the impact of SLR1 on GA metabolism (Supplemental Figure 7A and Supplemental Tables S1.6 to S1.7). Notably, we observed overall downregulation in genes coding for enzymes essential in non-gibberellin diterpenoid synthesis, namely syn-copalyl diphosphate synthase (*CPS4*) and ent-cassa-12,15-diene synthase (*KSL7*) (Supplemental Figures S7A and S7C). These enzymes are part of a diterpenoid pathway branching point that feeds the phytoalexins biosynthetic pathway (Otomo et al., 2004). Concurrently, other genes related to phytoalexin and phytocassane synthesis, such as cytochrome P450 family members *CYP701A8/KO1, CYP99A3, CYP71Z6, CYP76M8*, and *CYP71Z7*, were also downregulated in SLR1-OX.

Phenotypical discrimination of SLR1-OX and SUMO1SLR1-OX was observed under salt stress but not under control conditions (Figure 3C). Interestingly, a closer look into the distribution of SLR1-OX and SUMO1SLR1-OX samples in the PCA, under control and stress conditions highlights a similar tendency (Supplemental Figure S6). Hence, we focused on the analysis on the 236 DEGs specific to SUMO1SLR1-OX exposed to salt stress. After functional annotation of DEGs specific to SUMO1SLR1-OX, we found a significant over-representation of GO terms associated with defence response and gibberellin dioxygenase activity (Supplemental Figures S7B and S7C). We also found a salt stress-dependent induction of *GA20ox2* and *GA2ox3* genes in SUMO1SLR1-OX (Supplemental Table S1.4). *GA20ox2* and *GA2ox3* encode enzymes involved in the regulation of active GA (GA_4_) levels. Therefore, it is reasonable to hypothesize that SLR1 SUMOylation could adjust active GA homeostasis upon exposure to salt stress (Supplemental Figure S7C).

### SUMOylation disrupts SLR1 interaction with stress-associated transcription factors

The increased salt tolerance and sustained productivity of SUMO1SLR1-OX under salt stress likely arise from the presence of increased levels of SUMOylated SLR1 (Figure 3B-C; Figure 4A-B), which seems to be affecting the gene expression profile of plants exposed to stress. The *GA20ox2* transcript was induced around 2-fold in SUMO1SLR1-OX exposed to salt stress, while it showed only a marginal induction in SLR1-OX under the same conditions (Supplemental Table S1.2). One possible mechanism by which SLR1 could affect *GA20ox2* transcription is through the interaction with the transcription factor YAB4, thereby repressing the expression of *GA20ox2* (Yang et al., 2016). We did not detect *YAB4* transcript changes after 8 hours of salt stress exposure, but a search in public transcriptome data revealed that *YAB4* is induced in rice shoots after 3h of salt stress imposition (Genevestigator – Hruz et al., 2008; Supplemental Figure S9A-B), which could lead to increased YAB4 protein accumulation during later stages of plant adaptation to salt stress. In an attempt to dissect the molecular pathways involved in *GA20ox2* upregulation in SUMO1SLR1-OX, we verified the interaction of SLR1 and SUMOylated SLR1 with YAB4 TF using a yeast two-hybrid system and bimolecular fluorescence complementation (BiFC). The aim was to test the effect of SUMOylation on SLR1-YAB4 interaction, which could affect *GA20ox2* transcription. We used a variant of the SUMO1SLR1 fusion protein lacking the SUMO protease recognition domain to avoid possible processing by yeast SUMO proteases (SUMO1(_GG-AA_)::SLR1). We confirmed the previously demonstrated interaction between SLR1 and YAB4 (Yang et al., 2016), and also found that the mimicking SUMOylation in lysine 2 of SLR1 disrupts SLR1-YAB4 interaction in both yeast and *in planta* (Figure 5A-B; negative controls in Supplementary Figure S8). The blocking of the SLR1 interaction with YAB4 by SUMO correlates with the induction of *GA20ox2* in salt-stressed SUMO1SLR1-OX lines.

**Figure 5.**
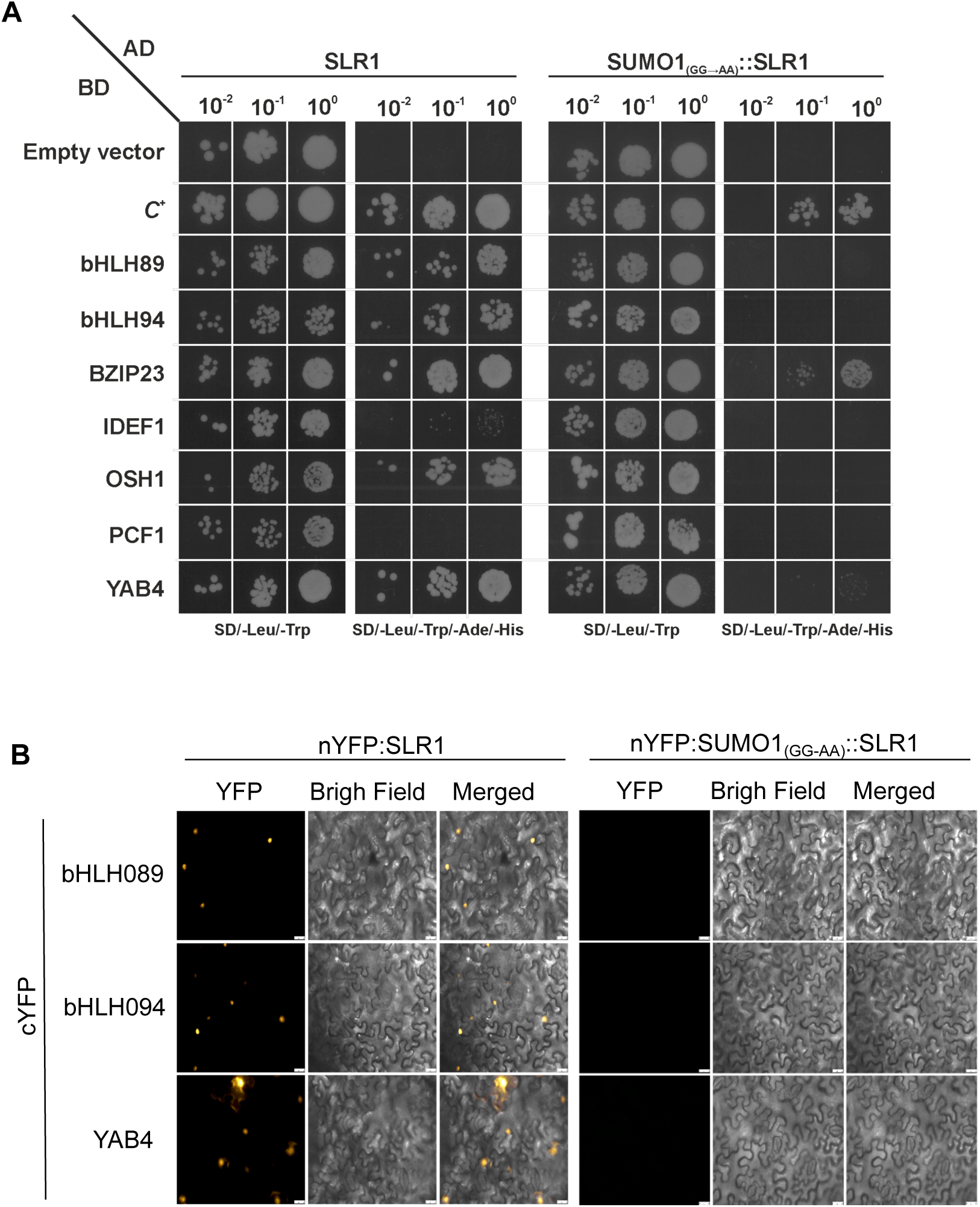
SLR1 SUMOylation affects its interaction with key transcription factors. **(A)** Yeast-two hybrid (Y2H) analysis of both SLR1 and SUMO1SLR1. SUMO1::SLR1 fusion carries two mutations in the SUMO-protease cleavage motif (GG → AA) to avoid its removal by endogenous yeast proteases and guarantee constitutive SUMO attachment to SLR1 during the assay. Positive controls marked as (SD/-Leu/-Trip) and positive interactions marked with (SD/-Leu/-Trip/-Ade/-His). The vectors pAD-WT and pBD-WT (Stratagene) were used as positive control. **(B)** Bimolecular fluorescent complementation (BiFC) assays showing positive interactions between nYFP: SLR1 with cYFP:BHLH089, cYFP:BHLH094, and cYFP:YAB4. The presence of SUMO1 attached to lysine 2 of SLR1 (nYFP:SUMO1_(GG-AA)_::SLR1) disrupts SLR1 interaction with these transcription factors. Negative controls for Y2H and BiFC assays can be found in Supplementary Figure S8.

To further evaluate the molecular aspects of SUMO1SLR1-OX tolerance to salt stress, we explored the notion that SLR1 exerts its function mainly by transcriptional regulation through interaction with transcription factors. We designed a strategy that allowed us to identify SLR1 TF interactors that could explain, at least partially, the differences in gene expression observed in the transgenic lines upon exposure to salt stress. Briefly, we submitted the promoter sequences (1000 bp upstream to the Transcription Start Site – TSS) of 417 selected DEGs (SLR1- and SUMO1SLR1-dependent salt stress responsive genes; Supplemental Tables S1.3 and S1.4) to an *in silico* search for transcription factor binding sites (TFBS) in the PlantPAN online database (Chow et al., 2016). Next, we selected only TFBS and corresponding binding TFs that have been experimentally validated. This allowed us to identify TFs from the TCP, bZIP, bHLH, and B3 families that could be regulating the output gene expression of SLR1-OX and SUMO1SLR1-OX transgenic lines under salt stress (Table 1). To cross-validate the TF identification, we made sure that a substantial overlap between the known target genes of those TFs and the SLR1- and SUMO1SLR1-dependent DEGs were observed (Supplemental Tables S1 and S2).

**Table 1.**
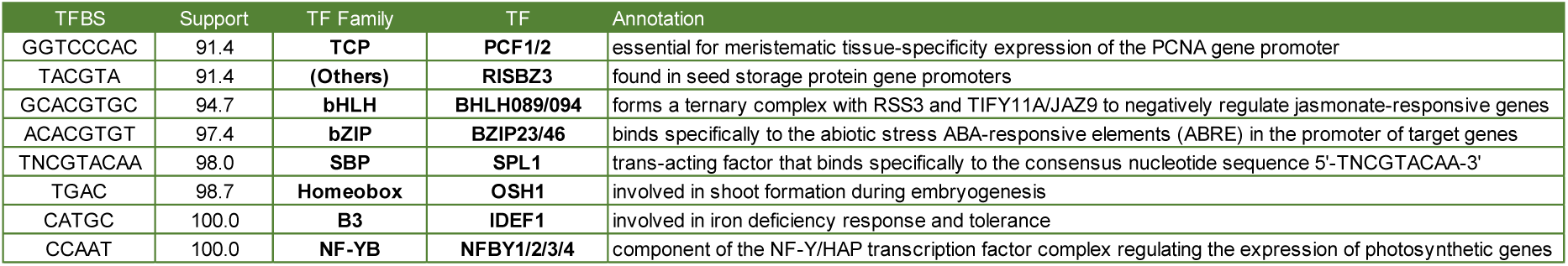
Enriched binding sites and associated transcription factors in SLR1 DEGs promoters. Promoter sequence analysis output from PlantPAN with Support referencing the percentage of transcription factor binding elements (TFBS/Sequence) found in SLR1- and SUMO1SLR1-dependent stress-responsive DEGs. Resulting TF overlap with DEGs detailed in Supplementary Table S4.

The interaction between SLR1 with the selected candidates (Table 1) and its regulation by SUMOylation was assessed through Y2H assays and BiFC analyses (Figure 5A-B). The basic helix-loop-helix transcription factors bHLH089 and bHLH094, as well as the homeobox TF OSH1, showed a strong interaction with SLR1 but not with SUMO1(_GG-AA_)::SLR1. This finding suggests that SUMOylation could modulate the binding of SLR1 to these transcription factors, potentially affecting their transcriptional activities. We also found that bZIP23 can interact with both SUMOylated and non-SUMOylated SLR1 (Figure 5A). The interaction of SLR1 with bHLH089 and bHLH094, as well as its disruption by SLR1 SUMOylation, was validated through BiFC in *Nicotiana benthamiana* leaves, supporting the *in planta* SUMO-dependent regulation of these interactions (Figure 5B). Neither IDEF1 nor PCF1 showed interactions with SLR1 or SUMO1(_GG-AA_)::SLR1.

## DISCUSSION

DELLA proteins are central regulators of plant productivity and stress responses, and the hormonal crosstalk mediated by these proteins is essential for plant plasticity to endure stress. As other hormone modulators, DELLA activity is regulated by post-translational modifications, such as ubiquitination, phosphorylation, *O*-fucosylation, *O*-GlcNAcylation, and SUMOylation (Yano et al., 2019; Zentella et al., 2016; Zentella et al., 2017; Conti et al., 2014; Dai and Xue, 2010; Itoh et al., 2003). Recently, SUMO proteases were linked to salt stress response in rice (Srivastava et al., 2016b; 2017). Although deSUMOylation was proposed as a general mechanism of tolerance acquisition in response to salt, we show that artificially SUMOylated SLR1 could alleviate the salt stress impact on rice productivity, also improving recovery after stress imposition is released.

Following the evidence that SUMO proteases are important for DELLA activity (Campanaro et al., 2016), the knockout mutant of SUMO protease *fug1* showed a higher ratio of overall SUMOylated SLR1. Interestingly, the FUG subclass is phylogenetically close to the SPF subclass, which contains Ulp2-like SUMO proteases (Castro et al., 2018). A Ulp2 from *Saccharomyces cerevisiae* (ScUlp2) was previously reported to be the key protease responsible for poly-SUMO chain processing (Eckhoff and Dohmen, 2015). If FUG1 is indeed related to Ulp2 proteases and it processes SUMO-SLR1, SLR1 deSUMOylation may be occurring in poly-SUMO chain high molecular weight conjugates.

DELLA proteins control several plant processes through interaction with a myriad of transcriptional factors (Van De Velde et al., 2017). By affecting protein-protein interaction, SLR1 SUMOylation has the potential to modulate plant processes by altering the transcriptional outcome of SLR1-TFs interacting proteins. Indeed, transcriptomic analysis of SUMOylated SLR1 overexpressing lines suggested that this isoform plays a role in early transcriptional responses to salt stress, affecting the expression of key genes coding for enzymes controlling the accumulation of active GA, such as *GA20ox2* and *GA2ox3*, involved in activation and inactivation of the active GA4. We were able to determine that SLR1 SUMOylation occurs in lysine 2 (Figure 1A-C), distant from the conserved GA regulatory domains DELLA and TVHYNP. This finding suggests that SLR1 SUMOylation contrasts with Arabidopsis RGA, in which the amino acid residue responsible for SUMO modification is the lysine 65 (Conti et al., 2014), located very close to the GA-regulatory domain TVHYNP. The location of SUMO attachment in SLR1 protein could exert quite different effects on protein-protein interactions compared to its well-described Arabidopsis counterpart RGA. An interesting possibility is that SUMOylation of these residues could have a different outcome regarding the transcriptional activation properties of SLR1 DELLA/TVHYNP motifs (Hirano et al., 2012). Both lysines are within SUMOylation canonical motifs, but while the SUMOylation canonical motif surrounding K2 is conserved in all analysed DELLAs, the one containing K65 is exclusive to RGA. The different SUMOylation sites of SLR1 and RGA might account for alterations in DELLA regulation amongst species that probably determine different interactome networks.

The SUMOylation of SLR1 could also be affecting the interaction with upstream regulatory proteins. For instance, SUMO attachment could affect SLR1 interaction with SPY, which enhances SLR1 activity through *O-*fucosylation and is a major determinant of rice plant architecture (Yano et al., 2019). Crosstalk between SUMOylation and *O-* fucosylation could play a role in GA-promoted growth regulation by decreasing TF-interactions that result in growth repression (Camut et al., 2017). This control of the expression of GA enzymes is demonstrated by our transcriptomic results of SUMO1SLR1-OX lines under salt stress, which shows upregulation of both *GA20ox2* and *GA2ox3*, suggesting that SLR1 SUMOylation could also regulate GA homeostasis. The induction of genes involved in both anabolism and catabolism of bioactive GA, catalized by GA20ox2 and GA2ox3, respectivelly, seems paradoxical. However, each gene could be activated in different tissues under salt stress, determining tissue specific expression and regulating plant plasticity. These differences in spatial expression are lost in whole plant transcriptomic analysis. Further experiments evaluating *GA20ox2* and *GA2ox3* spatial expression under salt stress have to be done to address the issue.

SUMOylation of specific targets was suggested to be crucial for stress tolerance, fertility and yield maintenance in rice. For instance, bZIP23 SUMOylation levels are modulated by OTS1, which affects its stability and the plant’s ability to deal with drought and salt stress (Srivastava et al., 2017). Interestingly, we found that bZIP23 interacts with both SLR1 and SUMO1SLR1. Another novel interaction with both bHLH089 and bHLH094 shows upregulation of JAZ-like repressors like JAZ8, consistent with findings that this TF is regulating genes from the JA pathway as observed in our transcriptomic analysis. Both these interactions may partly explain the JA-induced stress response promoted by SLR1 accumulation (Supplemental Figures S7A and S7B).

It was already shown that YAB4 interacts with SLR1 to repress *GA20ox2* gene expression and restrain development in rice (Yang et al., 2016). We confirmed the interaction between SLR1 and YAB4 by Y2H assays and BiFC (Figure 5B-C). We additionally show that SUMOylation of SLR1 abolished its interaction with YAB4. Accordingly, we found a stress-induced upregulation of *GA20ox2* in SUMO1SLR1-OX (Supplemental Table S1). Moreover, *GA20ox2* was also found to be downregulated by KNOTTED1-like homeobox (KNOX) proteins like OSH1 (Sakamoto et al., 2006). SLR1 SUMOylation also disrupted its interaction with OSH1, suggesting that SLR1 SUMOylation mechanistic action is partly achieved through SLR1 dissociation from transcriptional regulators involved in GA signalling and stress signalling. It could represent a dissociation of DELLA function during salt stress tolerance from its impact on productivity by means of SUMOylation.

The emergence of SUMOylation of SLR1 as an effector of rice tolerance to salt stress, through the growth-maintaining modulation of protein-protein interaction with transcriptional regulators such as YAB4, is evidence of the tight regulation of the GA pathway by rice DELLA during stress. Recent studies have demonstrated that DELLA proteins are targets of different post-translational modifications, which could adjust their interactions with transcriptional regulators in a dynamic fashion, ensuring the best developmental decisions when facing stress. Further studies need to be performed to unveil upstream effectors that perceive stress and the underlying signal transduction pathway that leads to SLR1 SUMOylation. By establishing the trigger for SLR1 SUMOylation and its effect on stress-induced SLR1 protein-protein interactions, new strategies can be developed involving the transient post-translational modification of DELLA, as mutagenesis and selection of growth-favourable SLR1 alleles. This can be used to activate the SUMO-regulated tolerance mechanism when stress is forthcoming and protect crops from yield losses due to abiotic stress.

## METHODS

### SLR1 genomic DNA cloning and vectors construction

The intronless gene SLR1 entire coding sequence was amplified from Nipponbare wild-type genomic DNA, using the primers SLR1-CDS-Fw and SLR1-CDS-Rv (Supplemental Table S4). The fragment was cloned into the pJET 1.2/blunt-end cloning vector (ThermoScientific), and the SLR1 full sequence confirmed by Sanger sequencing. SLR1 CDS was then amplified from pJET-SLR1 using the primers pET-Fw-*Eco*RI and pET-Rv-*Psp*OMI (Supplemental Table S4) to insert the restriction sites for *Eco*RI and *Psp*OMI to clone SLR1 in the pET28a expression vector (Novagen). For the generation of rice transgenic lines overexpressing SLR1, the SLR1 coding sequence was amplified from pJET-SLR1 using OsSLR1-GW-Fw and OsSLR1-GW-Fw primers to insert the attB1 and attB2 recombination sites. The amplified PCR-product was recombined into the pDONR221 (ThermoFisher).The SUMO1SLR1 gene fusion was generated using high-fidelity PCR (Phusion DNA Polymerase; NEB) essentially as described in Atanassov et al. (2009). Primers overlapping both the OsSUMO1 3’-end and OsSLR1 5’-end OL-SLR1+SUMO1-3’-Fw and OL SUMO1+SLR1-5’-Rv, were used in combination with OsSLR1-Gw-Rv and OsSUMO1-GW-Fw, respectively, to generate the fragments to be fused. In the next step, the PCR products were purified and used in the fusion PCR reaction containing the OsSUMO1-GW-Fw and OsSLR1-Gw-Rv primers for the final amplification of the final fusion product (Supplemental Table S4). The gene fusions were confirmed by Sanger sequencing. To obtain rice transgenic plant lines overexpressing SLR1 (SLR1-OX) and SLR1 fused to SUMO1 (SUMO1SLR1-OX), pDONR221 containing the inserts described above was used as entry vector for the pANIC6B destination vector (Mann et al., 2011). The expression of SLR1 and SUMO1SLR1 is controlled by the maize *ZmUbi1* promoter (Figure 3A).

### Site-directed mutagenesis

For the production of truncated recombinant SLR1 protein (SLR1t) containing the leading 190 amino acid residues of SLR1, a stop codon was introduced by site-directed mutagenesis (SDM) in pET28a-SLR1 using *Pfu*Turbo DNA polymerase (Agilent) and the primers (SLR1t-STOP-Fw and SLR1t-STOP-Rv) designed with QuickChangeII online tool (Agilent) (Supplemental Table S4). Unmodified plasmids were digested with methylation-sensitive enzyme *Dpn*I (New England Biolabs). The mutated plasmids were transformed in *E. coli* DH5*α* and the mutagenized site confirmed by sequencing. Mutated pET28a coding the truncated SLR1 isoform (SLR1t) was subjected to further mutagenesis procedures to perform the amino acid substitutions K2R and K60R (primers SLR1t-K2R-Fw, SLR1t-K2R-Rv, SLR1t-K60R-Fw and SLR1t-K60R-Rv, all listed in Supplemental Table S4) following the same protocol.

### Recombinant Protein Production and Purification

pET28a vectors containing the full-length SLR1, N-terminus truncated SLR1 (SLR1t), SLR1t (K2R) and SLR1t (K60R) were transformed into *E. coli* strain BL21 Star (DE3, Sigma-Aldrich) for recombinant protein expression. Cells were grown at 37°C until OD600 = 0.6, and protein production was induced by addition of 200 μM isopropyl-d-1-thiogalactopyranoside (IPTG). After three hours, the cells were harvested by centrifugation in a J2MI centrifuge (Beckman) at 7000 RPM during 20 minutes at 4°C. The cells were resuspended in lysis buffer (phosphate buffer 20 mM pH 7.5, 500 mM NaCl, 10 mM Imidazole) containing 5 μg/mL of DNAse, 50 μg/mL of lysozyme, PMSF 100 mM and MgCl2 25 mM. After freezing at −20°C and thawing, extracts were loaded into a HisTrap chelating column (Amersham Biosciences) and proteins eluted in phosphate buffer containing 500 mM imidazole, following buffer exchange dialysis (phosphate buffer 20 mM, NaCl 150 mM, DTT 1 M, glycerol 2%), for increased protein stability. Recombinant proteins were analysed by Western Blot using custom-made SLR1 specific polyclonal antibody with Mini-Protean II system (Biorad).

### *In vitro* SUMOylation assays and ULP1 treatment

The *in vitro* SUMOylation assays were performed in 20 μL of SUMOylation reaction buffer (Werner et al., 2009) containing 3 μg of SLR1 recombinant protein, 1 μg of E1-activating enzyme (OsSAE1/ OsSAE2), 0.8 μg of E2-conjugating enzyme (OsSCE1a), 0.5 μg of OsSUMO1/2, 0.4 units of creatine phosphokinase, and 10 mM creatine phosphate. Reactions were initiated by adding 1 mM ATP and then incubated at 37°C for 1 h to 3 h. To confirm SUMOylation specificity, 3 μg of purified SUMO protease ULP1 from *Saccharomyces cerevisiae* (Li and Hochstrasser, 2003) was used for deSUMOylation analysis. SUMO conjugates were analysed by Western Blot with custom anti-SLR1 antibody. For ULP1 treatments, whole plants were ground in liquid N2 and 100 mg of ground tissue powder were mixed with non-denaturing buffer (Tris-HCl pH 8.0 50 mM, sodium chloride 150 mM, NP-40 1%, sodium deoxycholate 0.5%, SDS 0.1%, EDTA pH 8.0 1 mM, NEM 50 mM), followed by homogenisation by vortexing, and subsequent centrifugation at 4 °C for 20 minutes to recover the supernatant containing the proteins. The non-denaturing protein extracts were treated with 7 μg of ULP1, incubated at 37 °C with collection points taken at 5, 30, 60, and 120 seconds. Proteins were denatured at 90°C for 10 minutes with Laemmli Buffer with 5 mM DTT and anti-SLR1 immunoblot proceeded as described above.

### SLR1 protein extraction, detection and quantification analysis

Plant shoot samples were ground in liquid N2 and 100 mg of ground tissue used for protein extraction with denaturing TCA/Acetone precipitation, followed by resuspension in lysis buffer (7 M Urea, 2 M Thiourea, 30 mM Tris, 4% (w/v) CHAPS, 4 % (v/v), cOmplete protease inhibitor cocktail EDTA-Free (Sigma-Aldrich), 0.1% (v/v) Nuclease Mix 100x, and NEM 20 mM. Protein quantification was performed with 2-D Quant kit (GE Healthcare) following manufacturer’s instructions. Anti-SLR1 immunoblots were performed in a Mini-Protean II System (Biorad), using custom-made anti-SLR1 polyclonal specific antibody [1:5.000, overnight incubation] and anti-rabbit (Novex) [1:20.000, 90 min incubation] in 5% non-fat dry milk in TBS 1X. Quantification of native SLR1 bands provided by immunoblotting analysis, normalized to protein loading control using stripped membrane stained with Coomassie G-250 (Fluka), was performed with ImageJ software as indicated in the official ImageJ website online tutorial.

### Plant material and Rice transformation

Calluses induced from wild-type *Oryza sativa* (japonica spp.) cv. Nipponbare mature seeds were used for rice transformations according to (Almadanim et al., 2017), and selection for transformants asserted by hygromycin resistance and standard GUS-staining with X-Gluc solution (Roth) of a section of seminal roots. Proteins were extracted, detected and quantified using a modified TCA/Acetone method (Luís et al., 2016). Due to growth arrest phenotype and fertility decrease in homozygous lines of SUMO1SLR1-OX lines, all assays continued with positively-selected heterozygous plants. Moreover, lines #2 for both SUMO1SLR1-OX and SLR1-OX were selected for further analysis at rice booting stage and transcriptomic evaluation according to non-SUMOylated SLR1 levels, in order to avoid DELLA pleiotropic effects (Davière and Achard, 2015). Wild-type Nipponbare was used as a control for SLR1-OX and SUMO1SLR1-OX obtained by transformation as described above. Negative segregant lines of Zhonghua and Hwayoung varieties were used as controls for T-DNA insertion lines knock-out mutants for rice SUMO proteases ELS1 (04Z11JY66) and FUG1 (2A-20225 obtained from RMD (www.rmd.ncpgr.cn), developed by the National Centre of Plant Gene Research) and Postech (www.postech.ac.kr/life/pfg/risd, developed by Gynheung An), respectively. These insertion line seeds were germinated, self-crossed and screened using PCR analysis for homozygosity (Rosa et al., 2018). The T-DNA insertion position was determined by sequencing. Gene expression was analysed by RT-qPCR (Rosa et al., 2018).

### Salinity Stress Assays

For the assays performed at the seedling stage, heterozygous transgenic seeds were germinated and selected through screening for hygromycin resistance and GUS staining. Seedlings were transferred to hydroponics for salinity resistance screening (Almeida et al., 2016) with half-strength Murashige & Skoog Medium and MES buffer 250 mg/L (Duchefa) with pH adjusted to 5.2. Seedlings were grown for 14 days before the salt stress assay. Then, the seedlings were exposed to salt stress, with electric conductivity (EC) set and maintained at 12 dS/m with sodium chloride solution for the extension of the stress treatment. After 8 days, the seedlings were transferred to regular medium for recovery. Stress severity (SES score) – 1 for lower severity and 9 for higher severity – was evaluated as set by IRRI (Gregorio et al., 1997) at 6 and 8 days after stress and 8 days after recovery. Samples for transcriptomic analysis – 4 replicates of a pool of 3 plants – were collected at 8 hours after the initiation of the salt stress, corresponding to 10 hours after the lights went on, for both control and salt treatments, and frozen immediately in liquid nitrogen for grinding and analysis. Samples from 0h, 2h, 4h, and 8h treatments were obtained from shoots and roots collected separately, and frozen immediately in liquid nitrogen. Protein was extracted, detected and quantified as described above. For the booting stage assay, previously germinated and selected heterozygous transgenic seeds were sown in pots with 2-parts soil, 2-parts peat, and 1-part vermiculite. The experiment was set up in the greenhouse from June to September, according to IRRI standard screening procedures (Gregorio et al., 1997). Genotypes were maintained in separate containers, for both stress and control pots, to control for discrepancies in panicle initiation times. Panicle initiation was evaluated by the emergence of the flag-leaf with a distance between 1 and 2 cm to the adjacent leaf and salinity applied individually for each genotype at that point. Electric conductivity (EC) was set and maintained at 8 dS/m with sodium chloride solution for the extension of the life cycle. Upon complete maturation of the seed, agronomical parameters were measured (plant height, tillering, and panicle length) and panicles collected for further characterization of fertility (analysing the number of full matured seeds to total number of flowers ratio) and productivity (weight of 1000 seeds as measured in a Pfeuffer seed counter). The significance of statistics was asserted by t-test with Holm-Sidak method correction for multiple comparisons and a p<0.05 threshold.

### RNA Extraction and expression analysis

RNA was extracted from a total of 100 mg of ground frozen shoot tissue with Direct-zol kit (Zymo Research) according to manufacturer’s protocol with in-column DNAse treatment. The first-strand cDNA was synthesized from 2 μg of total RNA with an anchored-oligo (dT)18 primer using the Transcriptor High Fidelity cDNA Synthesis Kit (Roche) according to the manufacturer’s instructions. Reverse Transcription quantitative PCR (RT-qPCR) analysis was performed using a LightCycler 480 system (Roche) and the SYBR Green I Master mix (Roche). The RT-qPCR reactions were carried out with the primers listed in Supplemental Table S4. PCR running conditions were: one cycle at 95°C for 5 min and 45 cycles of amplification at 95°C for 10 s, 56– 60°C for 10 s and 72°C /10 s. The CT values were calculated using three technical replicates, and gene expression was assessed relative to described internal reference transcript controls, rice *UBIQUITIN CONJUGASE2* (OsUBC2) and *UBIQUITIN10* (OsUBQ10) (Moraes et al., 2015; Pabuayon et al., 2016). For the RNA-seq procedure, the small RNAs in the extracted total RNA samples were cleaned by using the RNeasy spin column from RNeasy Mini Kit (Qiagen) following RNA Cleanup protocol. RNA quality was assessed by Fragment Analyzer (ThermoFisher) with RNA Quality Number over 5. A total of 18 cDNA libraries were prepared, using three biological replicates for each genotype (SLR1-OX, SUMO1SLR1-OX, Nipponbare) and each condition (mock and salt). Enzymatic fragmentation with Pico Nextera kit and Smart-Seq2 multiplexed-library preparation were performed^49^ and single-end sequencing on NextSeq500 (Illumina) with High Output Kit (75 cycles).

### RNA-seq Data Analysis

Raw data FASTQ files were inputted on R/Bioconductor-based user interface analysis software Chipster for RNA-seq. Single-end reads were trimmed with PRINSEQ for 55 bp total with 18 bp trimmed on 5’ end and quality control of 28% set for the 3’ end for adapter removal. Read alignment was performed with TOPHAT2 against the *Oryza sativa* genome (version IRGSP-1.0.36), selecting Sanger-Phred +33 as quality score format. An average of 25.000.000 and 21.000.000 aligned and uniquely mapped reads, respectively, were obtained per library (Supplemental Table S2). For differential expression analysis, BAM files resulting from alignment were submitted to pairwise comparisons with Cufflinks2 tool Cuffdiff for a set 0.05 False Discovery Rate (5% FDR) with sequence-specific bias and multi-mapped read corrections enabled. Differentially expressed genes (DEGs) with a 1.5 fold-change were selected for data visualization and functional annotation using ClueGO plug-in (Bindea et al., 2009) for Cytoscape (Su et al., 2014).

### Promoter Analysis and Interaction Validation by Y2H and BiFC

The 1000 bp promoter region of differential expressed genes DEGs were downloaded from Rice Annotation Project Database (RAP-DB) (Sakai et al., 2013) and submitted to PlantPAN 2.0 (Chow et al., 2016) for *cis*-elements for transcription factor binding site (TFBS) analysis. Gene Group Analysis was used to search for common (90%) *cis*-element binding (TFBS) sites and the transcription factors (TFs) associated with DEG promoters. Rice TFs were examined individually for experimentally verified data concerning the interaction to binding sites and downstream target genes listing (Supplemental Table S2). Resulting target genes were then overlapped with previously verified DEGs from RNA-seq analysis. For the Y2H assays, coding sequences of SLR1 and OsSUMO1(_GG-AA_)::SLR1 were cloned into the pGADT7 vector (Stratagene, USA) while OsbHLH089, OsbHLH094, OsbZIP23, OsIDEF1, OsOSH1, OsPCF1 and OsYAB4 were cloned into the pGBKT7 (Stratagene, USA) vector. These constructs were transformed into *Saccharomyces cerevisiae* strain Y2HGold (Clontech, US) applying the LiAc method. Yeast transformed cells were plated on synthetic defined (SD) medium lacking leucine and tryptophan for plasmid transformation control and in SD lacking leucine (-Leu), tryptophan (-Trip), adenine (-Ade), and histidine (-His) for interaction screening. The interactions were evaluated in three individual colonies transformed with both plasmids. pAD-WT and pBD-WT from the HybriZAP 2.1 kit (Stratagene, USA) were used as a control of positive interaction (C+). pGBKT7 and pGADT7 empty vectors were also co-transformed with pGADT7-SLR1, pGADT7-SUMO1(_GG-AA_)::SLR1 and pGBKT7-bHLH089, pGBKT7-bZIP23, pGBKT7-IDEF1, and pGBKT7-YAB4, respectively, as negative controls. For bimolecular fluorescence complementation (BiFC), OsSLR1, OsSUMO1(_GG-AA_)::SLR1 were cloned in YFN43, and OsbHLH089, OsbHLH094 and OsYAB4 were cloned into YFC43 (Belda-Palazón et al., 2012). *Nicotiana benthamiana* leaves were co-infiltrated with *Agrobacterium tumefaciens* strain EH105 harbouring the appropriate vectors. Fluorescence was visualized using the Leica DM6 B fluorescence microscope (40x) with L5 filter cube (Leica, Germany) 48-72h following infiltration. As negative control, YFN43 (empty vector) was also co-infiltrated with YFC43-bHLH089, YFC43-bHLH094 and YFC43-YAB4. *Agrobacterium* expressing the viral silencing suppressor P19 was included in all infiltrations to increase the expression in *N. benthamiana* leaves. BiFC analyses were performed essentially as described in Belda-Palazón et al., 2012.

### Accession Numbers

Sequence data from this article can be found in the Rice Annotation Project Database (RAP-DB) under the following accession numbers: OsSLR1 (Os03g0707600), OsSUMO1 (Os01g0918300), OsSUMO2 (Os01g0918200), OsSAE1 (Os11g0497000), OsSCE1a (Os03g0123100), OsELS1 (Os01g0355900), OsFUG1 (Os03g0344300), OsUBC2 (Os02g0634800), OsUBQ10 (Os06g0681400), OsP5CS (Os05g0455500), OsRAB21 (Os11g0454300), RGA (At2g01570), OsCPS4 (Os04g0178300), OsKSL7 (Os02g0570400), OsCYP701A8/KO1 (Os06g0569500), OsCYP99A3 (Os04g0178400), OsCYP71Z6 (Os02g0570500), OsCYP76M8 (Os02g0569400), OsCYP71Z7 (Os02g0570700), OsCKX2 (Os01g0197700), OsCKX5 (Os01g0775400), OsLOX2 (Os08g0509100), OsCHT3 (Os06g0726100), OsCHT8 (Os10g054290), OsCHT11 (Os03g0132900), OsGA20ox2 (Os01g0883800), OsGA2ox3 (Os01g0757200), OsPCF1 (Os04g0194600), OsRISBZ3 (Os02g0266800), OsbHLH089 (Os03g0802900), OsbHLH094 (Os07g0193800), OsbZIP23 (Os02g0766700), OsOSH1 (Os03g0727000), OsPCF1 (Os03g0785800), OsIDEF1 (Os11g0175700), OsYAB4 (Os02g0643200).

## Supplementary Materials

Figure S1. Alignment of DELLAs protein sequences of selected monocots and dicots.

Figure S2. SRL1 can be SUMOylated by both OsSUMO1 and OsSUMO2 *in vitro.*

Figure S3. Accumulation dynamics of SRL1 transcript and protein in response to salt stress.

Figure S4. Tillering and panicle length were not affected by mild salt stress imposition.

Figure S5. Analysis of salt treatment efficiency by salt-stress marker gene induction.

Figure S6. Salt treatment had a major contribution to the differences in gene expression among Nipponbare, SLR1-OX and SUMO1SLR1-OX.

Figure S7. Transcriptomics reveal differential stress-induced regulation by SUMOylated SLR1.

Figure S8. Protein-protein interaction (PPI) negative controls for Y2H and BiFC assays.

Figure S9. Genevestigator Results for Gene Expression of YABBY4 (LOC_Os02g4950) in Salinity Assays.

Table S1. RNA seq data showing >1.5 fold Differentially Expressed Genes in Nipponbare, SLR1-OX and SUMOSLR1-OX, in mock and salt stress conditions.

Table S2. Expression changes of selected putative SLR1 TF interactors (A) and the respective downstream target genes (B). Fold-change shown as log2 FC. Comparison of condition A with condition B corresponding to respective selected analysis in Cufflinks (FPKM1 × FPKM2, for condition A and B, respectively). Nipp-Nipponbare; (m)-mock; (s)-stress

Table S3. RNA-seq read count for each of the sequenced libraries and respective legend

Table S4. List of primers used for gene cloning, site directed mutagenesis (SDM) and gene expression quantification (RT-qPCR). Details for the use of specific primers are given in the text.

## Acknowledgments

Portuguese Fundação para a Ciência e a Tecnologia (FCT) for fellowships for NMG (SFRH/BD/85799/2012), TF (PD/BD/135584/2018), MTGR (SFRH/BD/84219/2012), MAAR (SFRH/BD/87420/2012), PMB (SFRH/BPD/86742/2012). FCT Investigator IF/00764/2014 (POPH-QREN) for IAA and PTDC/BIA-FBT/31211/2017 for CM contract. Work was supported by P-KBBE/0001/2012 and also by the FCT research unit GREEN-it “Bioresources4sustainability” (UID/Multi/04551/2013 and UIDB/04551/2020). The funding sources had no involvement in study design, analyses, and interpretation of data, writing, or in the submission decision.

## Notes

### Competing Interest Statement

The authors have declared no competing interest.

### Summary of Updates

- Figure 1B revised: mislabeling (ULP1 treatments) corrected (mistake of minus and plus signals position). Additionally, "SLR1" arrow now indicates the right SLR1 protein band. - Introduction: RGA is O-GlcNAcylated by SECRET AGENT (not SLR1). - mention to "bZIP63" (wrong) instead "bZIP23' (right) was corrected - Discussion: added a short discussion about why GA20ox2 and GA2ox3 are both induced, accordingly with the referee's suggestion.

