## Supplemental Figures S1-S9 for "SUMOylation of rice DELLA SLR1 modulates transcriptional responses and improves yield under salt stress"

```
                 K2                                                DELLA
RGA_ARATH        MKRDHHQFQGRLSNHG----TSSSSSSISKDKMMMVKKEEDGG--GNMDDELLAVLGYKV 54
RGL2_ARATH       MKRGYGETWDPPPKPLPAS-RSG-----EGPSMADKKKADDDNNNSNMDDELLAVLGYKV 54
GAI_ARATH        MKRDHH------------------HHHHQDKKTMMMNEEDDG----NGMDELLAVLGYKV 38
RGL1_ARATH       MKREHNHRESSAGEGG------------SSSMTTVIKEE------AAGVDELLVVLGYKV 42
RGL3_ARATH       MKRSHQ----------ETS-VEE-----EAPSMVEKLENGCGGGGDDNMDEFLAVLGYKV 44
SLR1_ORYSJ       MKREYQEAGGSSGGGSSAD------MG-SCKDKVMAGAAG----EEEDVDELLAALGYKV 49
DWRF8_MAIZE      MKREYQDAGGSG-----GD------MG-SSKDKMMAAAAGAGEQEEEDVDELLAALGYKV 48
GAI_SOLLC        MKRDRDRDREREKRAF--------SNGAVSSGKSKIWEEDEEEKPDAGMDELLAVLGYKV 52
GAI_GOSHI        MKRDHQEI---SGSGS--------NPAESSSIKGKLWEEDPDA--GGMDDELLAVLGYKV 47
SLN1_HORVU       MKREYQDGGGSGGGG--DE------MG-SSRDKMMVSSSEAGE--GEEVDELLAALGYKV 49
GAI1_VITVI       MKREYHHPHHP-------------TCSTSPTGKGKMWDADPQQ--DAGMDELLAVLGYNV 45
RHT1_WHEAT       MKREYQDAGGSGGGG--GG------MG-SSEDKMMVS-AAAGE--GEEVDELLAALGYKV 48
RGA1_BRACM       MKRDLHQFQGPNHGTSIAGSSTSSPAVFGKDKMMMVKEEED--------DELLGVLGYKV 52
GAIP_CUCMA       MKREHHYLHPRPEPPSVATGSNRESY--LNTGKAKLWEEEVQL--DGGMDELLAVLGYKV 56
SLR1_ORYSI       MKREYQEAGGSSGGGSSAD------MG-SCKDKVMAGAAG----EEEDVDELLAALGYKV 49
RGA2_BRACM       MKRDLHQFQGPPDTRF-PNHGTANTGSSSKDKMMMVKEEEDGG--N--MDELLAVLGYKV 55
GAIPB_CUCMA      MKREHHHLHPRPDPPSMAAAPNGDTY--LNTGKAKLWEEDAQL--DGGMDELLAVLGYKV 56

                          K60                            TVHYNP
RGA_ARATH        RSSEMAEVALKLEQLETMMSN---------VQEDGLSHLATDTVHYNPSELYSWLDNMLS 105
RGL2_ARATH       RSSEMAEVAQKLEQLEMVLSN---------DD-VG-STVLNDSVHYNPSDLSNWVESMLS 103
GAI_ARATH        RSSEMADVAQKLEQLEVMMSN---------VQEDDLSQLATETVHYNPAELYTWLDSMLT 89
RGL1_ARATH       RSSDMADVAHKLEQLEMVLG-------------DGISNLSDETVHYNPSDLSGWVESMLS 89
RGL3_ARATH       RSSDMADVAQKLEQLEMVLSN---------DI-ASSSNAFNDTVHYNPSDLSGWAQSMLS 94
SLR1_ORYSJ       RSSDMADVAQKLEQLEMAMGMGGVSAPGA-ADDGFVSHLATDTVHYNPSDLSSWVESMLS 108
DWRF8_MAIZE      RSSDMADVAQKLEQLEMAMGMGGVGGAGATADDGFVSHLATDTVHYNPSDLSSWVESMLS 108
GAI_SOLLC        KSSDMADVAQKLEQLEMAMGT---------TMEDGITHLSTDTVHKNPSDMAGWVQSMLS 103
GAI_GOSHI        RSSDMADVAQKLEMLEKVMGT---------AQEDGISQLG-DTVHFNPSDLSGWVQNLLI 97
SLN1_HORVU       RASDMADVAQKLEQLEMAMGMGG-----PAPDDGFATHLATDTVHYNPTDLSSWVESMLS 104
GAI1_VITVI       KASDMAEVAQKLEQLEEVIVN---------AQEDGLSHLASETVHYNPSDLSNWLGSMLS 96
RHT1_WHEAT       RASDMADVAQKLEQLEMAMGMGGVGA-GAAPDDSFATHLATDTVHYNPTDLSSWVESMLS 107
RGA1_BRACM       RSSEMAEVALKLEQLETMMGN---------AQEDGLAHLATDTVHYNPAELYSWLDNMLT 103
GAIP_CUCMA       KSSDMAEVAQKLEQLEEAMCQ---------VQDTGLSHLAFDTVHYNPSDLSTWVESMLT 107
SLR1_ORYSI       RSSDMADVAQKLEQLEMAMGMGGVSAPGA-ADDGFVSHLATDTVHYNPSDLSSWVESMLS 108
RGA2_BRACM       RSSEMAEVALKLEQLETMMGN---------VQEDGLSNLATDTVHYNPSELYSWLDNMLT 106
GAIPB_CUCMA      KSSDMAEVAQKLEQLEEAMCQ---------VQDTGLSHLAFDTVHYNPSDLSTWLESMIT 107
```

**Supplemental Figure S1. Alignment of DELLAs protein sequences of selected monocots and dicots.** Protein alignment was performed employing CLUSTALO (version 1.2.4). Protein identifiers correspond to UniProt accessions for all reviewed DELLA entries. ARATH (*Arabidopsis thaliana*), ORYSJ (*Oryza sativa* spp. japonica), MAIZE (*Zea mays*), SOLLC (*Solanum lycopersicum*), GOSHI (*Gossypium hirsutum*), HORVU (*Hordeum vulgare*), VITVI (*Vitis vinifera*), WHEAT (*Triticum aestivum*), ORYSI (*Oryza sativa* spp. indica), BRACM (*Brassica rapa*), CUCMA (*Cucurbita maxima*). The motifs marked in the black rectangles delimits the predicted SUMOylation motifs that correspond to putative SUMOylated lysine 2 (K2) and lysine 60 (K60) of *Oryza sativa* SLR1. Domains marked in red highlight the DELLA/TVHYNP N-terminal motifs responsible for DELLA protein stability.

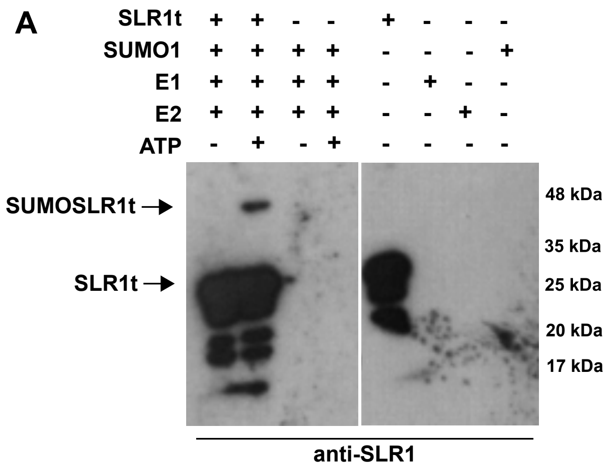


**Supplemental Figure S2. SRL1 can be SUMOylated by both OsSUMO1 and OsSUMO2 *in vitro.*** SUMOylation assays with and without ATP using SLR1t recombinant protein as detected by immunoblot using custom made anti-SLR1. **(A)** N-terminal truncated SLR1t and SUMO1. **(B)** N-terminal truncated SLR1t and SUMO2.

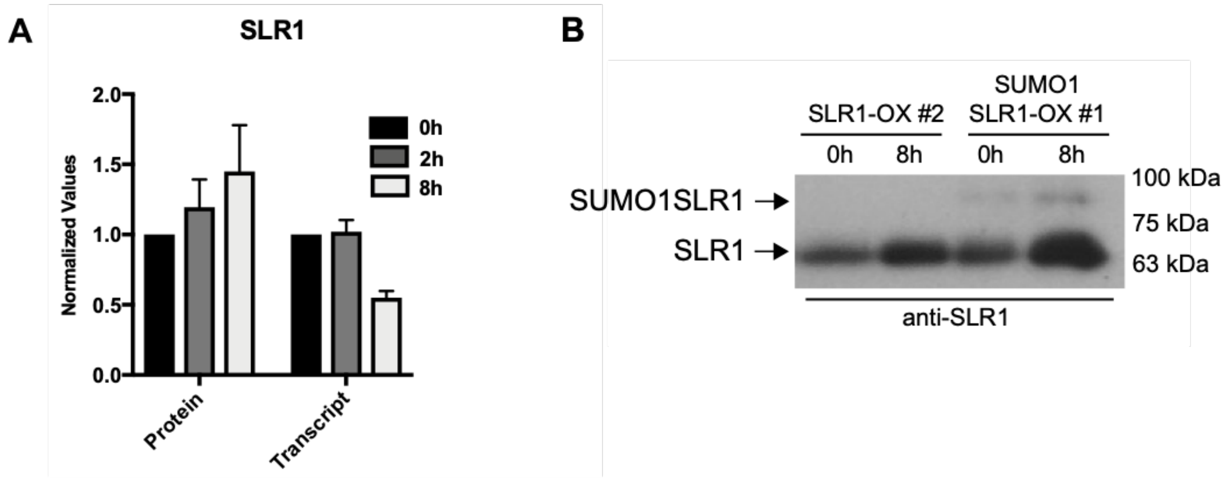


**Supplemental Figure S3. Accumulation dynamics of SRL1 transcript and protein in response to salt stress. (A)** Loading-normalized quantification of total SLR1 protein as detected by immunoblot using custom made anti-SLR1 antibody, and RT-qPCR SLR1 transcript expression analysis after 0, 2 and 8h of 120 mM salt stress imposition in wild-type rice (Nipponbare). The SLR1 mRNA expression was normalized by the average of two reference genes (UBC2 and UBQ10). **(B)** SLR1 and SUMO1SLR1 protein accumulation in response to salinity stress (120 mM for 8h) in T2 transgenic seedlings of SUMO1SLR1-OX and SLR1-OX transgenic lines as detected by anti-SLR1 antibodies.

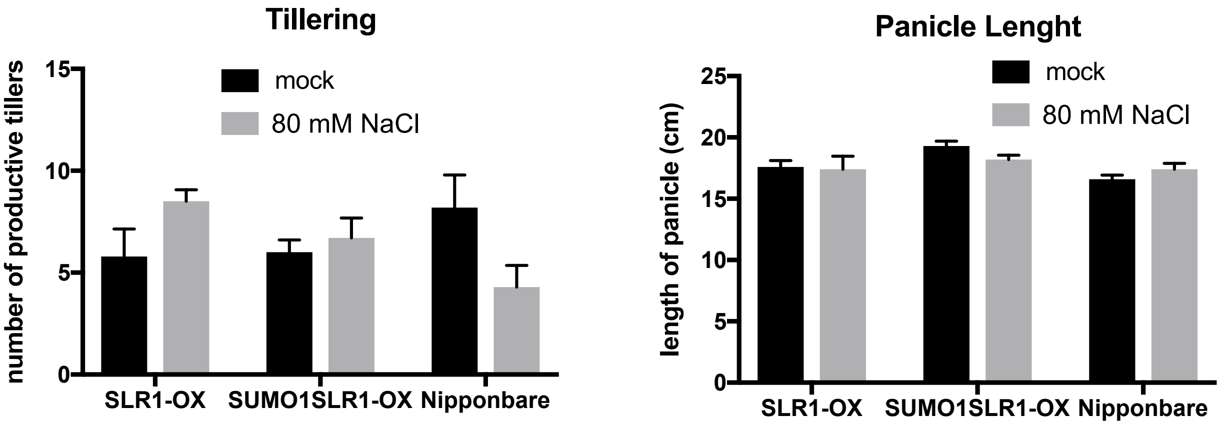


**Supplemental Figure S4. Tillering and panicle length were not affected by mild salt stress imposition.** Agronomic parameters as measured at the end of rice life cycle after 1 month of salt stress (80 mM NaCl) applied at panicle initiation during the booting stage: tillering (n=6); panicle length (n>10). The significance of statistics was asserted by t-test with Holm-Sidak method correction for multiple comparisons and a $p<0.05$ threshold.

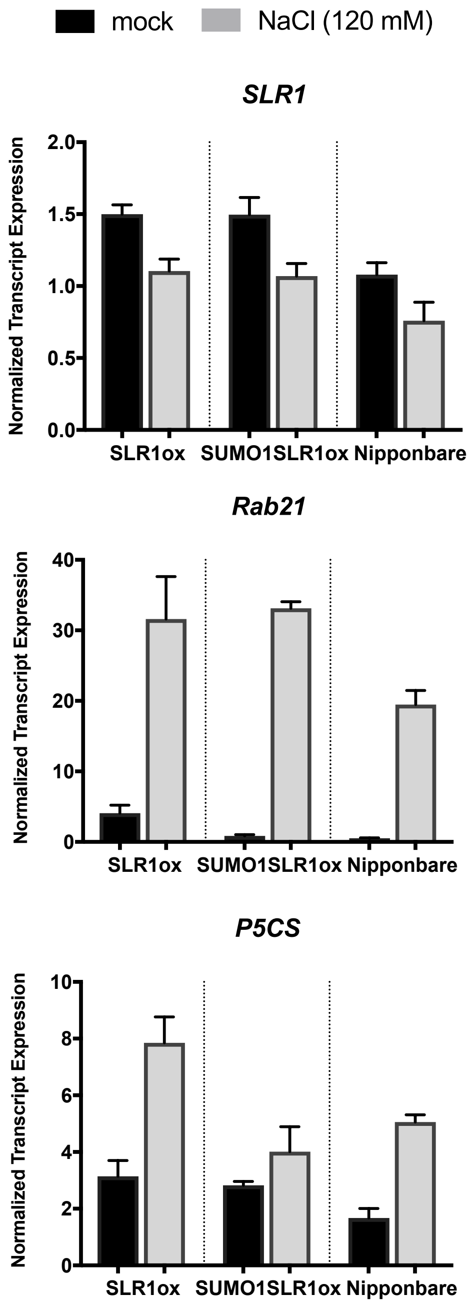


**Supplemental Figure S5. Analysis of salt treatment efficiency by salt-stress marker gene induction.** Changes in gene expression induced by salt stress of SLR1 and the salt-induced genes *Rab21* and *P5CS*, respectively. Transcript amounts were assessed through RT-qPCR analysis in shoots after 8 hours of salt treatment (stress) along with untreated control (mock). Individual gene expression was normalized using the average Ct of two reference genes (UBC2 and UBQ10).

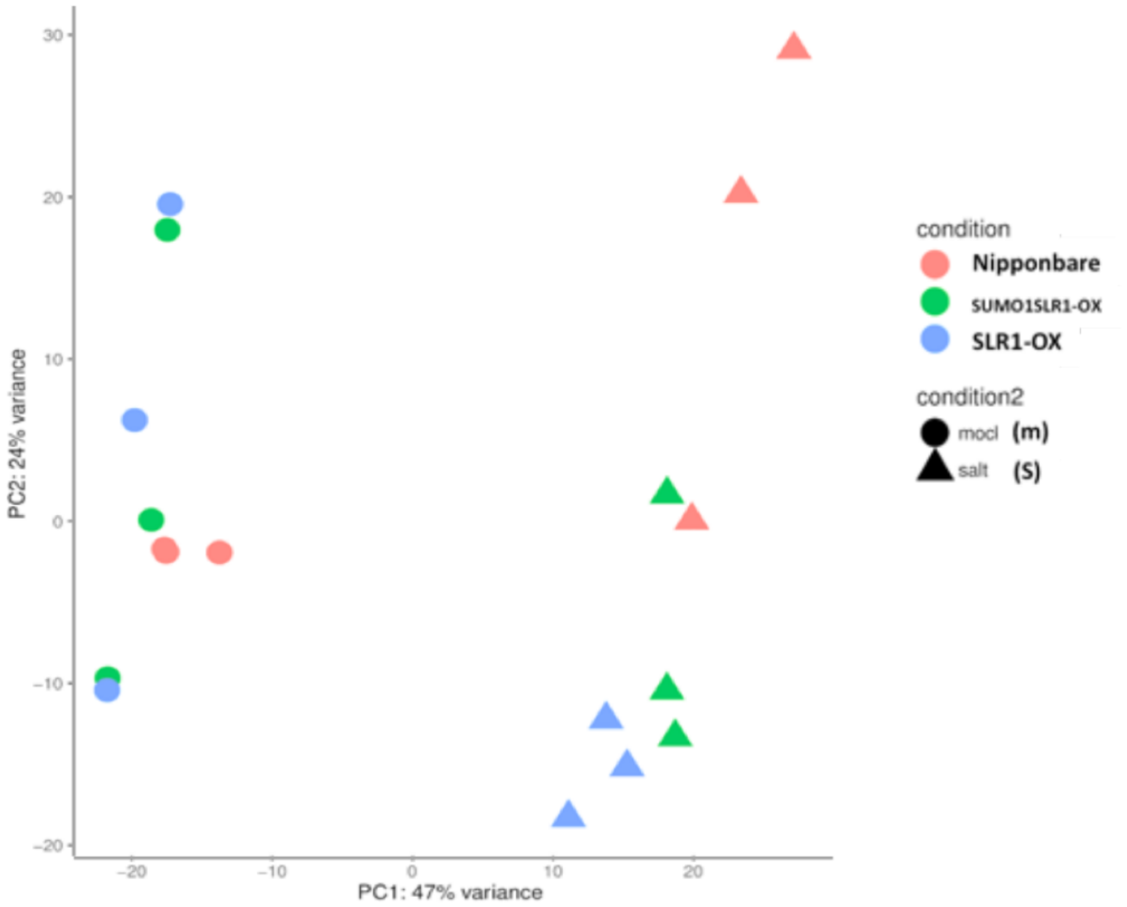


**Supplemental Figure S6. Salt treatment had a major contribution to the differences in gene expression among Nipponbare, SLR1-OX and SUMO1SLR1-OX.** Principal Component Analysis of the three replicates of the libraries subjected to RNA-seq sequencing provided by DESeq2 package. Along PC2 axis, no clear organization was found between samples for control conditions, which may explain the absence of DEGs under an FDR of 5% when comparing the corresponding SLR1-OX and SUMO1SLR1-OX individual mRNA profiles replicates (*i.e.* individual libraries). The mRNA profile replicates of salt-stressed plants were organized according to genotype along PC2, suggesting that an increase in the amount of SUMO1SLR1 in SUMO1SLR1-OX could be driving the gene expression variance between the two OX genotypes at a certain extent.

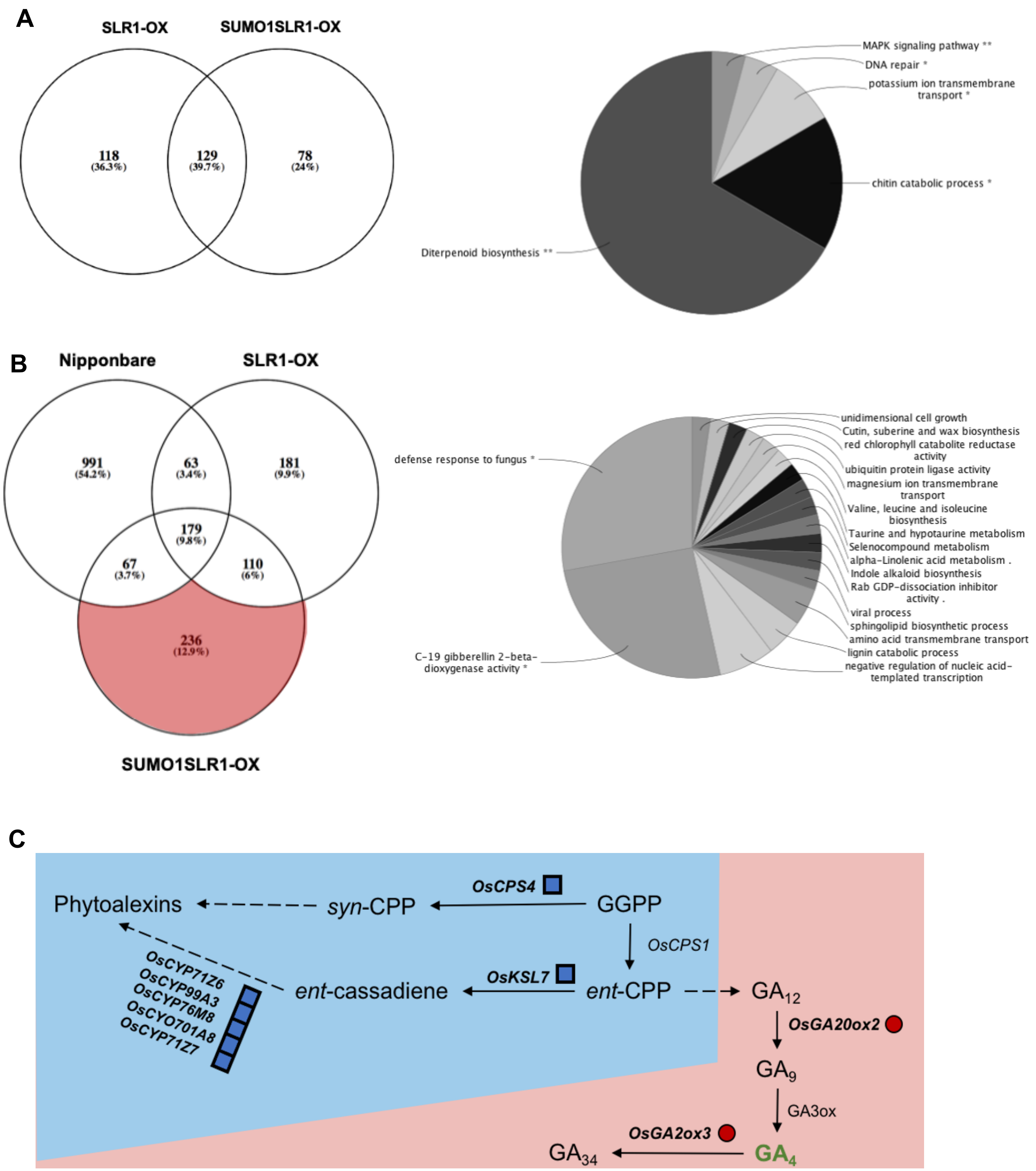


**Supplemental Figure S7. Transcriptomics reveal differential stress-induced regulation by SUMOylated SLR1. (A-B)** Venn diagrams showing the overlaps between differentially expressed genes as detected through RNA-seq analysis of Nipponbare, SLR1-OX and SUMO1SLR1-OX exposed or not to salt stress. Pie charts representing GO term-enrichment for biological process category obtained for selected genes. **(A)** Overlap between SLR1-OX and SUMO1SLR1-OX differentially expressed genes compared to wild-type in control conditions. SLR1-dependent genes (247 genes) were selected for the GO functional enrichment analysis represented in the right. **(B)** Differentially expressed genes found for SLR1-OX, SUMO1SLR1-OX and wild-type

(Nipponbare) lines under salt stress as compared with the respective genotypes under control conditions. SUMOylated SLR1-exclusive genes (236 genes) were selected for GO functional enrichment analysis represented in the right. Statistical testing for functional enrichment analysis was performed using a two-sided hypergeometric test with Bonferroni step down correction (*p value<0.05. **p value<0.01). **(C)** Highlights of diterpenoid and GA biosynthetic pathway with downregulated genes from SLR1-induced variation in blue squares and upregulated genes from SUMOylated SLR1-exclusive variation in red circles. Bioactive gibberellin (GA4) is represented in green font color.

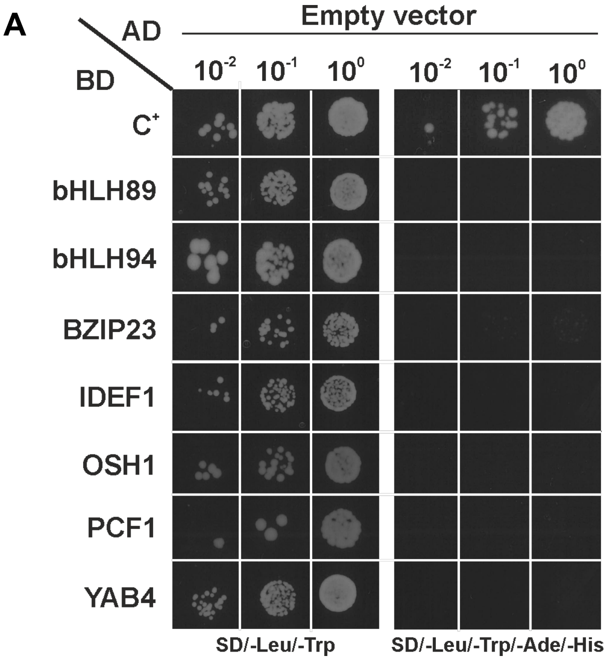


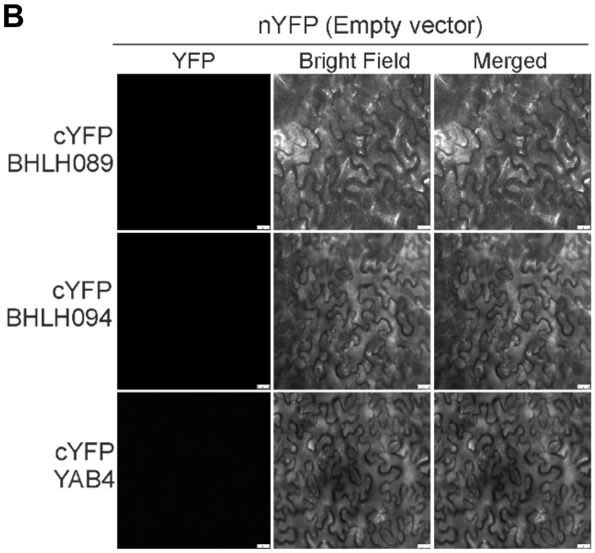


**Supplemental Figure S8. Protein-protein interaction (PPI) negative controls for Y2H and BiFC assays.** AD – Activation Domain (Y2H) and nYFP alone (empty vectors) do not interact with the transcription factors selected for this work **(A)** Y2H assay with positive controls marked as (SD/-Leu/-Trip) and positive interactions marked with (SD/-Leu/-Trip/-Ade/-His) with C+ as an protein-protein interaction control. The vectors pAD-WT and pBD-WT (Stratagene) were used as positive control. **(B)** Bi-molecular fluorescent complementation (BiFC) assay testing YFN43 (nYFP) empty vector for interaction with cYFP (YFC43) fused to bHLH089, bHLH094 and YAB4.

**A**

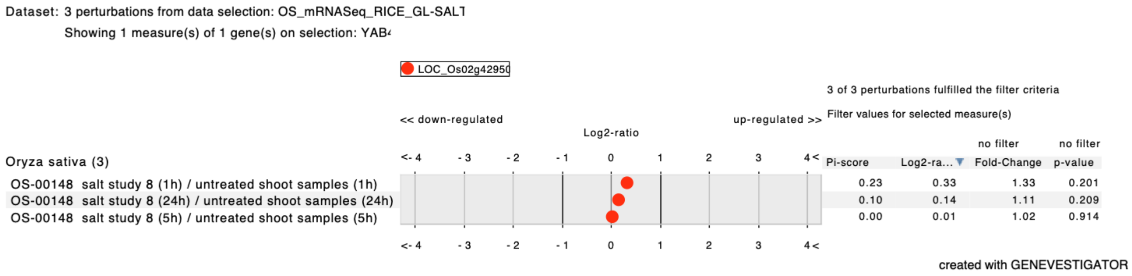


**B**

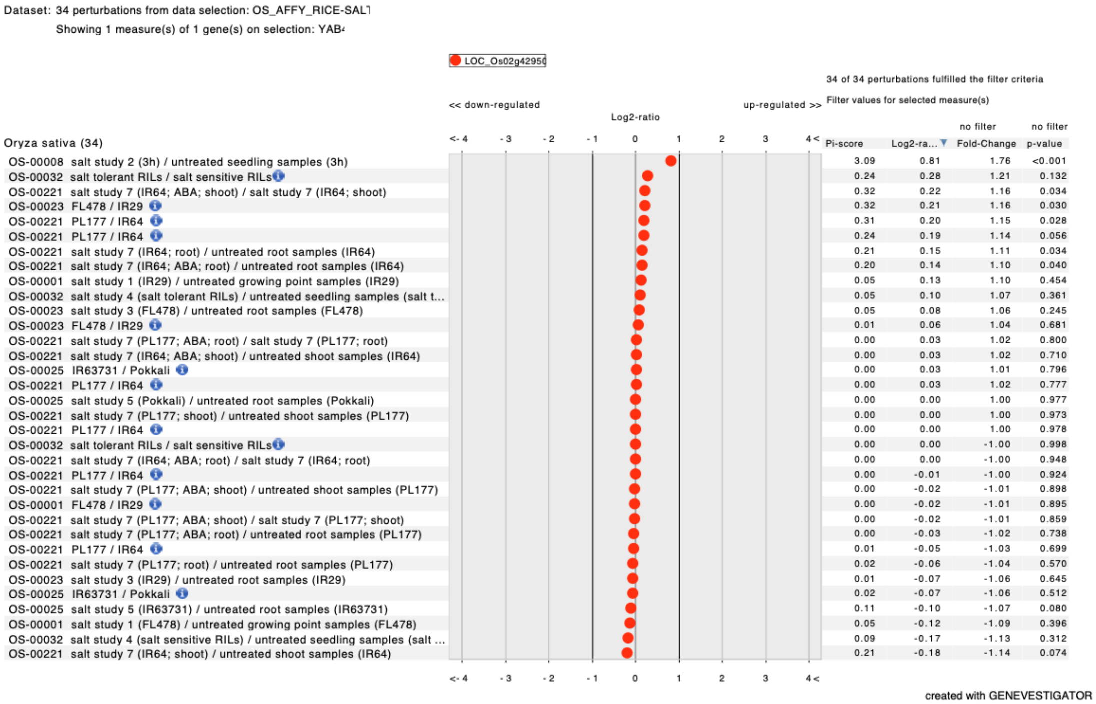


**Supplemental Figure S9.** Genevestigator Results for Gene Expression of YABBY4 (LOC_Os02g4950) in Salinity Assays. Available from **(A)** RNA-seq data and **(B)** Affymetrix Micro Array data.
