## Supplemental Tables S1-S4 for "SUMOylation of rice DELLA SLR1 modulates transcriptional responses and improves yield under salt stress"

**Supplemental Table S1.** Misregulated genes in SLR1-OX and SUMO1-SLR1-OX identified using messenger RNA sequencing (mRNA-seq).

| **Legend:** | |
|---|---|
| **N** | Nipponbare (WT) |
| **SS** | SUMO1SLR1-OX |
| **S** | SLR1-OX |
| **m** | Condition: mock |
| **s** | Condition: salt |
| **1,2,3** | Replicates |
| **FC** | Fold-change |

| 1. Overall stress-responsive | | | | |
|---|---|---|---|---|
| Identifier | Description | FC.Nipp | FC.SSox | FC.Sox |
| OS07G0443500 | Molecular chaperone, heat shock protein, Hsp40, DnaJ domain containing protein. (Os07t0443500-00) | -5.36986 | -3.93429 | -4.19645 |
| OS10G0552600 | Similar to Tfm5 protein. (Os10t0552600-01) | -3.53724 | -2.49074 | -2.59067 |
| OS06G0480500 | Non-protein coding transcript. (Os06t0480500-01) | -3.51976 | -3.64977 | -2.90619 |
| OS10G0320100 | Similar to Flavonoid 3'-monooxygenase (EC 1.14.13.21) (Flavonoid 3'-hydroxylase) (Cytochrome P450 75B2). (Os10t0320100-01) | -3.47092 | -2.88305 | -2.53566 |
| OS01G0663051 | Similar to SANT/MYB protein. (Os01t0663051-00) | -3.20819 | -2.48593 | -2.70729 |
| OS05G0503650 | Non-protein coding transcript. (Os05t0503650-01) | -2.92754 | -3.18161 | -3.30983 |
| OS05G0500600 | GRAS transcription factor domain containing protein. (Os05t0500600-01) | -2.76276 | -2.17042 | -2.92463 |
| OS04G0141400 | Concanavalin A-like lectin/glucanase domain containing protein. (Os04t0141400-01) | -2.68346 | -3.11831 | -2.80984 |
| OS10G0552700 | Similar to Tumor-related protein (Fragment). (Os10t0552700-01) | -2.57808 | -2.62194 | -1.99708 |
| OS07G0142200 | Conserved hypothetical protein. (Os07t0142200-01) | -2.394 | -2.38621 | -2.64425 |
| OS09G0351700 | Protein kinase, catalytic domain domain containing protein. (Os09t0351700-00) | -2.36211 | -1.50426 | -1.82898 |
| OS10G0508900 | Conserved hypothetical protein. (Os10t0508900-01) | -2.33888 | -2.15541 | -1.89753 |
| OS01G0735300 | UDP-glucuronosyl/UDP-glucosyltransferase family protein. (Os01t0735300-01) | -2.31592 | -1.96211 | -2.19383 |
| OS06G0210000 | Protein of unknown function DUF6 domain containing protein. (Os06t0210000-00) | -2.30431 | -3.07956 | -2.53266 |
| OS01G0878700 | Amino acid transporter, transmembrane domain containing protein. (Os01t0878700-02);Amino acid transporter, transmembrane domain containing protein. (Os01t0878700-03) | -2.17062 | -1.68359 | -1.926 |
| OS12G0163700 | Similar to Actin 7 (Actin 2). (Os12t0163700-01) | -2.08911 | -3.75557 | -2.85685 |
| OS09G0246300 | Hypothetical protein. (Os09t0246300-01);Conserved hypothetical protein. (Os09t0246300-02) | -2.08577 | -1.63377 | -1.76023 |
| OS07G0115300 | Similar to Peroxidase2 precursor (EC 1.11.1.7). (Os07t0115300-01) | -1.99848 | -2.14453 | -1.58869 |
| OS07G0142300 | Similar to Mucin-2. (Os07t0142300-01) | -1.98842 | -2.07945 | -2.38192 |
| OS11G0153500 | Protein of unknown function DUF3049 domain containing protein. (Os11t0153500-01) | -1.92427 | -2.1726 | -2.06058 |
| OS03G0664400 | Similar to Protease inhibitor/seed storage/LTP family protein. (Os03t0664400-01);Expressed protein (With alternative splicing). (Os03t0664400-02) | -1.91388 | -1.96971 | -1.87473 |
| OS01G0253400 | Protein of unknown function DUF1218 family protein. (Os01t0253400-01) | -1.89019 | -1.52632 | -1.94361 |
| OS04G0445300 | Pectinesterase inhibitor domain containing protein. (Os04t0445300-01) | -1.87098 | -1.72123 | -2.66737 |
| OS01G0778400 | Hypothetical protein. (Os01t0778400-00) | -1.84291 | -2.56491 | -2.70899 |
| OS07G0693800 | Omega-3 fatty acid desaturase, Acclimation to low-temperature stress, Defense response (Os07t0693800-01);Similar to Plastid omega-3 fatty acid desaturase (Fragment). (Os07t069380 | -1.8027 | -1.77337 | -1.59882 |
| OS02G0783000 | Similar to Pectin methylesterase 5 (Fragment). (Os02t0783000-01) | -1.77681 | -1.81415 | -1.59247 |
| OS10G0552800 | Similar to Tfm5 protein. (Os10t0552800-01) | -1.73844 | -2.47018 | -2.86848 |
| OS05G0499300 | Similar to Peroxidase (EC 1.11.1.7). (Os05t0499300-01) | -1.73588 | -2.18937 | -2.13456 |
| OS08G0355400 | Protein of unknown function DUF247, plant family protein. (Os08t0355400-01);Hypothetical conserved gene. (Os08t0355400-02) | -1.71715 | -2.03224 | -1.98785 |
| OS02G0240300 | Similar to Class III peroxidase GvPx2b (Fragment). (Os02t0240300-01) | -1.70446 | -1.97688 | -2.15965 |
| OS03G0195300 | Similar to Low affinity sulphate transporter 3. (Os03t0195300-01) | -1.62668 | -2.49844 | -1.60253 |
| OS03G0836400 | Harpin-induced 1 domain containing protein. (Os03t0836400-01) | -1.58215 | -2.35521 | -1.80641 |
| OS08G0460000 | Similar to Germin-like protein 1 precursor. (Os08t0460000-01) | -1.57195 | -1.69679 | -1.68612 |
| OS01G0635200 | Homeodomain-like containing protein. (Os01t0635200-01) | -1.56403 | -2.53114 | -2.17593 |
| OS01G0935800 | Similar to BRASSINOSTEROID INSENSITIVE 1-associated receptor kinase 1. (Os01t0935800-01) | -1.54599 | -2.41963 | -3.05532 |
| OS03G0314500 | WD40-like Beta Propeller domain containing protein. (Os03t0314500-01);Hypothetical conserved gene. (Os03t0314500-02) | -1.54185 | -1.50192 | -2.7771 |
| OS10G0150000 | Hypothetical conserved gene. (Os10t0150000-01) | -1.53018 | -2.74271 | -2.77495 |
| OS01G0316600 | Similar to pre-mRNA-splicing factor SF2. (Os01t0316600-01) | 1.51207 | 2.21902 | 1.8996 |
| OS04G0667300 | Hypothetical protein. (Os04t0667300-01) | 1.51761 | 1.96832 | 1.83836 |
| OS09G0418000 | Similar to CBL-interacting protein kinase 16. (Os09t0418000-01);Serine/threonine protein kinase domain containing protein. (Os09t0418000-02);Similar to CBL-interacting protein kinase | 1.52399 | 2.82494 | 2.61663 |
| OS01G0800500 | Purple acid phosphatase, N-terminal domain containing protein. (Os01t0800500-01) | 1.52735 | 1.58572 | 1.72934 |
| OS07G0687900 | WSI76 protein induced by water stress. (Os07t0687900-01) | 1.54065 | 1.76749 | 2.2985 |
| OS03G0386000 | Similar to Salt-responsive WD40 protein 5. (Os03t0386000-01);WD40 subfamily protein, Salt stress (Os03t0386000-02) | 1.54299 | 1.95322 | 1.62363 |
| OS06G0651200 | Conserved hypothetical protein. (Os06t0651200-01) | 1.5505 | 2.80257 | 2.93353 |
| OS01G0849000 | Hypothetical protein. (Os01t0849000-02) | 1.57147 | 2.22081 | 2.13613 |
| OS06G0223700 | Protein of unknown function DUF581 family protein. (Os06t0223700-01) | 1.57454 | 1.70931 | 1.7787 |
| OS05G0138300 | Similar to ICT protein (Fragment). (Os05t0138300-01);Similar to ICT protein (Fragment). (Os05t0138300-03) | 1.58421 | 1.70132 | 1.87582 |
| OS01G0219300 | Conserved hypothetical protein. (Os01t0219300-01) | 1.59159 | 2.87633 | 2.0236 |
| OS06G0683100 | NAD(P)-binding domain containing protein. (Os06t0683100-01) | 1.59741 | 2.77596 | 2.15229 |
| OS01G0846300 | Similar to Protein phosphatase 2C. (Os01t0846300-01);Similar to protein phosphatase 2C. (Os01t0846300-02) | 1.61 | 2.2134 | 2.09606 |
| OS08G0205800 | Hypothetical conserved gene. (Os08t0205800-01) | 1.62369 | 2.2707 | 1.7161 |
| OS04G0201800 | Similar to H0512B01.7 protein. (Os04t0201800-00) | 1.64281 | 2.05165 | 2.41701 |
| OS08G0110600 | Protein of unknown function DUF1442 domain containing protein. (Os08t0110600-00) | 1.64824 | 3.30536 | 2.59091 |
| OS07G0676900 | Similar to Peroxidase (EC 1.11.1.7). (Os07t0676900-01) | 1.68615 | 1.64713 | 1.79229 |
| OS11G0115400 | Lipid transfer protein LPT IV. (Os11t0115400-00) | 1.7132 | 1.79118 | 1.74936 |
| OS03G0746900 | Protein of unknown function DUF1677, plant family protein. (Os03t0746900-01) | 1.7148 | 2.4421 | 2.07248 |
| OS04G0631600 | Basic helix-loop-helix dimerisation region bHLH domain containing protein. (Os04t0631600-01) | 1.75307 | 1.58677 | 1.61772 |
| OS03G0268600 | Similar to Protein phosphatase type 2C. (Os03t0268600-01) | 1.75623 | 2.43004 | 3.24772 |

| | | | | |
|---|---|---|---|---|
| OS05G0595200 | Allergen V5/Tpx-1 related family protein. (Os05t0595200-01);Similar to allergen V5/Tpx-1-related family protein. (Os05t0595200-02) | 1.76929 | 1.58845 | 1.6962 |
| OS01G0192300 | Similar to I-box binding factor (Fragment). (Os01t0192300-01);Hypothetical conserved gene. (Os01t0192300-02) | 1.77149 | 1.93722 | 1.84662 |
| OS04G0122000 | Leucine-rich repeat, N-terminal domain containing protein. (Os04t0122000-01);Leucine-rich repeat, N-terminal domain containing protein. (Os04t0122000-02);Leucine-rich repeat doma | 1.7765 | 2.51149 | 1.97181 |
| OS05G0595100 | Uridine-diphospho-(UDP)-glucose 4-epimerase, Cell wall carbohydrate partitioning during nitrogen (N) limitation (Os05t0595100-01) | 1.77795 | 1.73236 | 1.81028 |
| OS02G0651900 | Hypothetical protein. (Os02t0651900-01) | 1.78276 | 1.70159 | 1.67783 |
| OS07G0586100 | Similar to Esterase precursor (EC 3.1.1.-) (Early nodule-specific protein homolog) (Latex allergen Hev b 13). (Os07t0586100-01) | 1.80115 | 2.30776 | 2.2882 |
| OS04G0244800 | Heavy metal transport/detoxification protein domain containing protein. (Os04t0244800-01) | 1.82455 | 1.86015 | 2.20276 |
| OS10G0450800 | Hypothetical protein. (Os10t0450800-02) | 1.84337 | 2.17696 | 2.04974 |
| OS11G0634200 | Conserved hypothetical protein. (Os11t0634200-01) | 1.85311 | 1.7717 | 1.93947 |
| OS02G0649300 | HD-ZIP I protein, Transcription activator, Stress response, Panicle development (Os02t0649300-01) | 1.854766 | 2.525355 | 1.94931081 |
| OS07G0290200 | Plant lipid transfer protein/seed storage/trypsin-alpha amylase inhibitor domain containing protein. (Os07t0290200-01) | 1.89163 | 1.7594 | 1.73121 |
| OS03G0189400 | Similar to Alcohol dehydrogenase ADH. (Os03t0189400-01) | 1.90577 | 2.26104 | 1.85278 |
| OS01G0726700 | Conserved hypothetical protein. (Os01t0726700-01) | 1.92288 | 2.00097 | 2.67005 |
| OS02G0592833 | Protein of unknown function DUF581 family protein. (Os02t0592833-00) | 1.92486 | 1.99851 | 1.94012 |
| OS08G0485400 | Similar to Oxidoreductase. (Os08t0485400-01);Similar to 2-nitropropane dioxygenase-like protein. (Os08t0485400-02) | 1.93506 | 2.18224 | 2.07669 |
| OS06G0258000 | Similar to Typical P-type R2R3 Myb protein (Fragment). (Os06t0258000-01) | 1.9434 | 1.78045 | 1.87558 |
| OS01G0658900 | OSBZ8. (Os01t0658900-01) | 1.95529 | 3.90876 | 3.66577 |
| OS07G0633200 | Similar to SC35-like splicing factor SCL30a, 30a kD. (Os07t0633200-01);Similar to SC35-like splicing factor SCL30a, 30a kD. (Os07t0633200-02) | 1.95695 | 1.66152 | 1.78613 |
| OS05G0496000 | Conserved hypothetical protein. (Os05t0496000-01) | 1.96681 | 1.66311 | 1.5756 |
| OS04G0266900 | Transketolase C-terminal-like domain containing protein. (Os04t0266900-01) | 2.01376 | 1.9834 | 1.58026 |
| OS04G0494100 | Similar to Chitinase. (Os04t0494100-02) | 2.03299 | 1.64959 | 1.60878 |
| OS01G0511100 | UspA domain containing protein. (Os01t0511100-01) | 2.03841 | 1.921 | 2.1547 |
| OS11G0444900 | Octicosapeptide/Phox/Bem1p domain containing protein. (Os11t0444900-01) | 2.05003 | 1.79635 | 1.65004 |
| OS05G0195101 | Non-protein coding transcript. (Os05t0195101-01) | 2.05209 | 3.44986 | 2.35531 |
| OS03G0340500 | Similar to Sucrose synthase (EC 2.4.1.13). (Os03t0340500-01);Similar to sucrose synthase2. (Os03t0340500-03) | 2.1385 | 1.98021 | 2.04411 |
| OS11G0634350 | Non-protein coding transcript. (Os11t0634350-01) | 2.15093 | 1.762906 | 1.98872831 |
| OS06G0567900 | Similar to Ascorbate oxidase (Fragment). (Os06t0567900-01) | 2.15147 | 2.98079 | 2.4531 |
| OS05G0537400 | Similar to Protein phosphatase 2C. (Os05t0537400-00) | 2.16035 | 2.70446 | 1.63991 |
| OS03G0180900 | Tify domain containing protein. (Os03t0180900-01) | 2.1989 | 1.67315 | 1.64946 |
| OS02G0139000 | Transcription factor, Regulation of Pi signaling and homeostasis, Tolerance to low-Pi stress (Os02t0139000-01) | 2.21372 | 2.23438 | 1.94579 |
| OS02G0622500 | Conserved hypothetical protein. (Os02t0622500-01) | 2.23995 | 2.0742 | 1.6128 |
| OS09G0537700 | S-like ribonuclease, Salinity tolerance, Abiotic stress response, Regulation of photomorphogenesis (Os09t0537700-02) | 2.2403 | 1.77198 | 2.06698 |
| OS02G0106100 | Similar to Fructosyltransferase. (Os02t0106100-01) | 2.26409 | 4.5072 | 3.88203 |
| OS01G0699400 | Serine/threonine protein kinase domain containing protein. (Os01t0699400-01) | 2.26497 | 2.4571 | 3.34352 |
| OS07G0120650 | Protein of unknown function DUF538 family protein. (Os07t0120650-01) | 2.26981 | 3.50647 | 2.06538 |
| OS07G0169600 | 2OG-Fe(II) oxygenase domain containing protein. (Os07t0169600-01) | 2.28203 | 2.80023 | 2.4285 |
| OS04G0607500 | Ion transporter , Na+ transport (Os04t0607500-01);Similar to Cation transporter HKT4. (Os04t0607500-02) | 2.28838 | 2.18542 | 1.81045 |
| OS01G0615100 | Similar to Substilin /chymotrypsin-like inhibitor (Proteinase inhibitor). (Os01t0615100-01) | 2.32488 | 3.72562 | 2.39173 |
| OS11G0427800 | Similar to Lipid transfer protein LPT III. (Os11t0427800-01) | 2.37725 | 2.18998 | 3.1494 |
| OS02G0824500 | Similar to Remorin. (Os02t0824500-01) | 2.4047 | 1.8867 | 3.72028 |
| OS04G0513400 | Similar to Beta-glucosidase. (Os04t0513400-01);Similar to Beta-glucosidase. (Os04t0513400-02) | 2.43141 | 1.72243 | 1.63716 |
| OS11G0582300 | Similar to root hair defective 3 GTP-binding (RHD3) family protein. (Os11t0582300-01);Similar to GTP-binding protein-like; root hair defective 3 protein-like. (Os11t0582300-02) | 2.43282 | 3.22854 | 2.14325 |
| OS01G0123900 | Similar to Bowman-Birk type proteinase inhibitor (EBI). (Os01t0123900-01) | 2.44394 | 2.15654 | 1.80359 |
| OS06G0725000 | Similar to Senescence-associated protein DIN1. (Os06t0725000-01) | 2.45348 | 1.65214 | 1.53044 |
| OS01G0794400 | Thioredoxin domain 2 containing protein. (Os01t0794400-01) | 2.46677 | 3.15432 | 3.56238 |
| OS06G0166500 | Similar to Auxin-responsive protein IAA20 (Indoleacetic acid-induced protein 20). (Os06t0166500-01) | 2.47437 | 3.92491 | 3.50473 |
| OS09G0464066 | Hypothetical protein. (Os09t0464066-00) | 2.50165 | 2.62376 | 1.65394 |
| OS06G0668200 | Similar to Phosphoglycerate kinase, cytosolic (EC 2.7.2.3). (Os06t0668200-01) | 2.55317 | 2.18176 | 2.69325 |
| OS03G0694000 | Similar to Oxalate oxidase 1 (EC 1.2.3.4) (Germin). (Os03t0694000-01) | 2.56046 | 3.45026 | 3.07855 |
| OS03G0607200 | Gibberellin regulated protein family protein. (Os03t0607200-01) | 2.57173 | 2.42164 | 1.93051 |
| OS03G0130300 | Similar to Cp-thionin. (Os03t0130300-01) | 2.57534 | 2.6926 | 2.96181 |
| OS07G0539300 | Glycoside hydrolase, family 17 protein. (Os07t0539300-01);Glycoside hydrolase, family 17 protein. (Os07t0539300-02) | 2.57677 | 2.8696 | 2.36732 |
| OS02G0258800 | Conserved hypothetical protein. (Os02t0258800-01) | 2.60841 | 2.79029 | 2.83127 |
| OS01G0227700 | Cytochrome P450 family protein. (Os01t0227700-01) | 2.61979 | 2.69858 | 2.49784 |
| OS01G0644200 | Similar to Little protein 1. (Os01t0644200-01) | 2.62459 | 3.376249 | 2.61756179 |
| OS10G0492900 | Similar to alpha-galactosidase. (Os10t0492900-01) | 2.62818 | 4.42741 | 4.23101 |
| OS12G0147200 | Conserved hypothetical protein. (Os12t0147200-01) | 2.66291 | 3.47605 | 3.69843 |
| OS04G0188433 | Similar to OSIGBa0102N07.2 protein. (Os04t0188433-01) | 2.6675 | 2.58387 | 1.97085 |
| OS05G0247100 | Similar to Glycosyl hydrolases family 18. (Os05t0247100-01);Chitinase III protein, Xylanase inhibitor, Regulation of abiotic stress response, Resistance to herbivores (Os05t0247100-02) | 2.72764 | 2.24861 | 1.81986 |
| OS01G0644000 | Twin-arginine translocation pathway signal domain containing protein. (Os01t0644000-01);Twin-arginine translocation pathway signal domain containing protein. (Os01t0644000-02) | 2.75389 | 2.67596 | 2.51607 |
| OS12G0448900 | Fatty acid alpha-dioxygenase family, Enzyme that oxygenates fatty acids into 2R-hydroperoxides (Os12t0448900-01) | 2.78865 | 2.26938 | 2.83558 |

| | | | | |
|---|---|---|---|---|
| OS05G0130100 | Hypothetical conserved gene. (Os05t0130100-00) | 2.79439 | 3.90573 | 3.63614 |
| OS01G0901600 | Similar to 4-coumarate--CoA ligase-like 6. (Os01t0901600-01) | 2.79811 | 2.92502 | 2.40178 |
| OS09G0272600 | Conserved hypothetical protein. (Os09t0272600-01) | 2.8115 | 3.23559 | 3.27245 |
| OS08G0112300 | Transferase domain containing protein. (Os08t0112300-01) | 2.82146 | 1.81444 | 1.78908 |
| OS05G0355400 | UspA domain containing protein. (Os05t0355400-01) | 2.82251 | 3.02835 | 2.61633 |
| **OS09G0379600** | **Similar to Homeobox-leucine zipper protein HOX25. (Os09t0379600-00)** | **2.83022** | **2.11391** | **2.52548** |
| **OS04G0541700** | **Similar to Homeobox-leucine zipper protein HOX22. (Os04t0541700-01);Homeodomain-leucine zipper (HD-Zip) protein, Transcription factor, ABA-mediated drought and salt toleranc** | **2.83321** | **2.06379** | **2.18024** |
| OS01G0155800 | Conserved hypothetical protein. (Os01t0155800-01) | 2.85171 | 4.44838 | 7.38163 |
| OS02G0276200 | Similar to Isochorismatase family protein rutB. (Os02t0276200-01) | 2.86881 | 1.6889 | 2.10723 |
| OS04G0493400 | Similar to Chitinase. (Os04t0493400-01) | 2.88454 | 2.02672 | 1.64085 |
| **OS11G0454000** | **Dehydrin RAB 16C. (Os11t0454000-01)** | **2.88999** | **3.80864** | **3.84233** |
| OS06G0698300 | Protein phosphatase 2C family protein. (Os06t0698300-01) | 2.92499 | 2.99899 | 2.51195 |
| OS03G0153900 | Aromatic-ring hydroxylase family protein. (Os03t0153900-01) | 2.92717 | 2.25596 | 2.29423 |
| OS10G0205700 | Pollen Ole e 1 allergen/extensin domain containing protein. (Os10t0205700-01) | 2.94269 | 3.77575 | 3.87048 |
| OS09G0538000 | Ribonuclease T2 family protein. (Os09t0538000-01);RNase S-like protein. (Os09t0538000-04);Non-protein coding transcript. (Os09t0538000-05) | 2.9784 | 2.09363 | 1.86672 |
| **OS11G0454200** | **Dehydrin RAB 16B. (Os11t0454200-01)** | **3.024644** | **4.87375** | **5.90849** |
| OS11G0592200 | Similar to Chitin-binding allergen Bra r 2 (Fragments). (Os11t0592200-01) | 3.06538 | 2.16429 | 1.79308 |
| OS03G0723400 | Similar to UFG2. (Os03t0723400-01) | 3.126443 | 4.33348 | 4.45218 |
| OS01G0124200 | Similar to Bowman Birk trypsin inhibitor. (Os01t0124200-02) | 3.20549 | 3.0923 | 2.41331 |
| OS01G0124700 | Hypothetical protein. (Os01t0124700-01) | 3.23792 | 2.87076 | 2.55543 |
| OS01G0214500 | Conserved hypothetical protein. (Os01t0214500-01) | 3.23982 | 2.56382 | 2.83289 |
| OS12G0428000 | Similar to senescence-associated protein DIN1. (Os12t0428000-01) | 3.25797 | 3.144 | 2.96057 |
| OS01G0832600 | Similar to Leucoanthocyanidin dioxygenase-like protein. (Os01t0832600-01) | 3.31159 | 3.03269 | 3.45363 |
| OS11G0592000 | Similar to Barwin. (Os11t0592000-01) | 3.33697 | 2.24978 | 1.58957 |
| OS01G0127600 | Similar to Bowman-Birk type proteinase inhibitor D-II precursor (IV). (Os01t0127600-01) | 3.36675 | 2.4095 | 2.02494 |
| OS05G0373900 | Similar to Eukaryotic peptide chain release factor subunit 1 (eRF1) (Eukaryotic release factor 1) (TB3-1) (Cl1 protein). (Os05t0373900-01) | 3.3912 | 4.21046 | 3.88457 |
| OS03G0729000 | Similar to Peptidase, M50 family. (Os03t0729000-01) | 3.4243 | 4.2651 | 4.42111 |
| **OS05G0381400** | **AWPM-19-like protein, Stress tolerance through ABA-dependent pathway (Os05t0381400-01)** | **3.444442** | **4.549466** | **4.69309661** |
| OS08G0395700 | Conserved hypothetical protein. (Os08t0395700-01) | 3.47867 | 2.98476 | 2.67344 |
| OS12G0559200 | Lipoxygenase (EC 1.13.11.12). (Os12t0559200-01);Similar to Lipoxygenase. (Os12t0559200-02) | 3.48461 | 3.18445 | 1.87785 |
| OS01G0124650 | Hypothetical conserved gene. (Os01t0124650-02) | 3.4919 | 3.0018 | 2.61229 |
| OS02G0783625 | Similar to Lysine ketoglutarate reductase/saccharopine dehydrogenase. (Os02t0783625-00) | 3.54243 | 3.40462 | 3.0115 |
| OS03G0826800 | Conserved hypothetical protein. (Os03t0826800-01) | 3.67262 | 2.73665 | 2.64492 |
| OS02G0513100 | Similar to MtN3 protein precursor. (Os02t0513100-01) | 3.70172 | 4.90645 | 3.72283 |
| OS12G0548401 | Similar to Proteinase inhibitor. (Os12t0548401-01) | 3.82512 | 3.43253 | 3.59409 |
| OS03G0718800 | Similar to Physical impedance induced protein. (Os03t0718800-01) | 3.84512 | 4.26467 | 4.20775 |
| OS12G0458100 | Transferase family protein. (Os12t0458100-01) | 3.90497 | 3.62784 | 1.81144 |
| OS03G0820500 | Similar to WCOR719. (Os03t0820500-01) | 3.96175 | 4.37808 | 4.30803 |
| **OS12G0247700** | **Similar to Jasmonate-induced protein. (Os12t0247700-01)** | **3.98897** | **4.74645** | **3.34788** |
| OS06G0498800 | Similar to MOTHER of FT and TF1 protein. (Os06t0498800-01) | 4.07408 | 5.1979 | 5.26524 |
| OS01G0124000 | Similar to Bowman Birk trypsin inhibitor. (Os01t0124000-01) | 4.2157 | 4.27422 | 3.70393 |
| OS09G0425200 | Conserved hypothetical protein. (Os09t0425200-00) | 4.3026 | 5.8658 | 5.75486 |
| OS06G0681200 | Cupredoxin domain containing protein. (Os06t0681200-01) | 4.30274 | 4.99788 | 6.85822 |
| OS03G0286900 | RCI2 (rare cold-inducible 2) family protein, Drought resistance (Os03t0286900-01) | 4.31536 | 3.52987 | 3.5026 |
| OS12G0437800 | Similar to MPI. (Os12t0437800-01) | 4.46273 | 3.86167 | 2.47825 |
| **OS09G0325700** | **Protein phosphatase 2C, Abiotic stress response, Early panicle development (Os09t0325700-01)** | **4.4687** | **3.20528** | **4.44344** |
| OS01G0124100 | Proteinase inhibitor I12, Bowman-Birk family protein. (Os01t0124100-01) | 4.47612 | 4.35108 | 2.8181 |
| OS05G0550600 | Similar to Nonspecific lipid-transfer protein 2 (nsLTP2) (7 kDa lipid transfer protein). (Os05t0550600-02) | 4.97718 | 5.85045 | 5.42305 |
| OS08G0508000 | Cytochrome P450 family protein. (Os08t0508000-01) | 5.06685 | 2.46419 | 2.09344 |
| OS07G0529000 | Similar to Isocitrate lyase (Fragment). (Os07t0529000-01) | 5.52987 | 3.43393 | 2.70139 |
| **OS01G0348900** | **SalT gene product (Salt-induced protein). (Os01t0348900-01)** | **5.69996** | **3.97548** | **4.18298** |
| **OS04G0511200** | **EFA27 for EF hand, abscisic acid, 27kD. (Os04t0511200-01)** | **5.74667** | **4.99467** | **4.33382** |
| OS01G0124401 | Similar to Bowman-Birk type bran trypsin inhibitor. (Os01t0124401-01) | 5.98477 | 5.86743 | 5.30277 |
| OS05G0468800 | Phosphatidylethanolamine-binding protein PEBP domain containing protein. (Os05t0468800-01) | 6.10237 | 5.24158 | 4.34189 |
| OS11G0211800 | Similar to Specific abundant protein-like protein 1. (Os11t0211800-01) | 6.6291 | 5.52693 | 4.82037 |
| **OS12G0123700** | **NAC domain containing transcription factor, Abiotic and biotic stress response (Os12t0123700-01)** | **6.75049** | **3.88525** | **3.59979** |
| **OS11G0454300** | **Similar to Water-stress inducible protein RAB21. (Os11t0454300-01)** | **6.76968** | **6.27587** | **7.65379** |
| OS07G0147550 | Similar to Photosystem II 10 kDa polypeptide, chloroplast. (Os07t0147550-00) | 7.57202 | 5.41752 | 4.07309 |
| OS09G0344500 | Similar to O-methyltransferase ZRP4 (EC 2.1.1.-) (OMT). (Os09t0344500-01) | 8.42084 | 4.74334 | 5.38753 |
| OS04G0344100 | Similar to OSIGBa0106G08.3 protein. (Os04t0344100-01) | 8.73221 | 4.90382 | 3.77444 |

#### 2. Stress-responsive in OXs

| Identifier | Description | FC.SSmSSs | FC.SmSs | FC.NmNs | FC.NmSSm | FC.NmSm |
|---|---|---|---|---|---|---|
| OS04G0525100 | UDP-glucuronosyl/UDP-glucosyltransferase family protein. (Os04t0525100-01) | -1.93898 | -2.79212 | - | - | - |
| OS04G0364800 | Barwin-related endoglucanase domain containing protein. (Os04t0364800-00) | -2.19329 | -2.62716 | -1.26063 | - | - |
| OS05G0463100 | Hypothetical gene. (Os05t0463100-01) | -2.47781 | -2.46887 | -1.30485 | - | - |
| OS01G0691400 | Similar to seed maturation protein. (Os01t0691400-00) | -2.80782 | -2.31508 | - | - | - |
| OS07G0142500 | Conserved hypothetical protein. (Os07t0142500-00) | -2.02365 | -2.24353 | -1.44225 | - | - |
| OS01G0880800 | Similar to Acyl-[acyl-carrier-protein] desaturase, chloroplast precursor (EC 1.14.19.2) (Stearoyl-ACP desaturase). (Os01t0880800-01) | -3.25761 | -2.21861 | - | 1.98664 | - |
| OS01G0922700 | Conserved hypothetical protein. (Os01t0922700-01) | -2.47917 | -2.14593 | -1.33582 | - | - |
| OS12G0194900 | Similar to Amino acid carrier (Fragment). (Os12t0194900-01) | -2.45024 | -2.10941 | -1.49812 | - | - |
| OS03G0281900 | ATP-binding cassette (ABC) transporter, Hypodermal suberization of roots, Salt stress tolerance (Os03t0281900-01) | -1.88593 | -2.05789 | - | - | - |
| OS08G0530100 | Nucleotide-diphospho-sugar transferase domain containing protein. (Os08t0530100-01) | -1.5634 | -1.95271 | - | - | - |
| OS12G0189300 | Pyruvate/Phosphoenolpyruvate kinase, catalytic core domain containing protein. (Os12t0189300-01);Non-protein coding transcript. (Os | -1.74844 | -1.93505 | -0.873105 | - | - |
| OS06G0294600 | Cytochrome P450 family protein. (Os06t0294600-02) | -1.93307 | -1.91845 | -1.43845 | - | - |
| OS06G0561000 | Myo-inositol oxygenase, Drought stress tolerance (Os06t0561000-01) | -1.61086 | -1.87632 | - | - | - |
| OS05G0400500 | Similar to plasma membrane associated protein. (Os05t0400500-01) | -1.81727 | -1.80436 | -1.41163 | - | - |
| OS02G0189200 | Lipase, GDSL domain containing protein. (Os02t0189200-01) | -1.66486 | -1.80067 | -1.39245 | - | - |
| OS06G0228200 | Aquaporin NIP III subfamily protein, Aquaporin, Silicon influx transporter, Water transport (Os06t0228200-01);Similar to NOD26-like me | -1.97578 | -1.76679 | -0.543913 | 1.32942 | - |
| OS05G0541750 | Hypothetical gene. (Os05t0541750-01) | -1.96009 | -1.76629 | -1.46485 | - | - |
| OS04G0464100 | Heavy metal transport/detoxification protein domain containing protein. (Os04t0464100-01) | -1.82406 | -1.76159 | -1.24676 | - | - |
| OS05G0462000 | Similar to plant-specific domain TIGR01589 family protein. (Os05t0462000-00) | -1.94493 | -1.74219 | -0.960341 | - | - |
| OS05G0518300 | Lipase, GDSL domain containing protein. (Os05t0518300-01) | -1.82775 | -1.72925 | -1.43552 | - | - |
| OS03G0221500 | Glycoside hydrolase, family 17 protein. (Os03t0221500-01);Glycoside hydrolase, family 17 protein. (Os03t0221500-02);Glycoside hydrola | -1.8596 | -1.68396 | - | 1.10381 | - |
| OS12G0601800 | Similar to BZIP transcription factor family protein, expressed. (Os12t0601800-01) | -1.55911 | -1.68331 | -1.12782 | - | - |
| OS02G0768300 | Protein of unknown function DUF6, transmembrane domain containing protein. (Os02t0768300-01) | -3.48506 | -1.67958 | - | - | - |
| OS02G0629800 | Similar to Defensin precursor. (Os02t0629800-01) | -1.66745 | -1.67597 | - | - | - |
| OS01G0644600 | Glutelin family protein. (Os01t0644600-00) | -1.95474 | -1.67493 | -1.19665 | - | - |
| OS06G0274800 | Similar to Peroxidase 11 precursor (EC 1.11.1.7) (Atperox P11) (ATP23a/ATP23b). (Os06t0274800-01) | -1.55842 | -1.65004 | -0.969284 | - | - |
| OS12G0451300 | Hypothetical conserved gene. (Os12t0451300-00) | -1.91622 | -1.64856 | - | - | - |
| OS12G0569200 | Conserved hypothetical protein. (Os12t0569200-01) | -1.60729 | -1.64825 | -1.35506 | - | - |
| OS11G0155900 | Histone H3. (Os11t0155900-00) | -1.79906 | -1.63571 | - | 1.24511 | - |
| OS08G0420600 | Similar to Permease 1. (Os08t0420600-01) | -2.46146 | -1.62938 | 1.23424 | - | - |
| OS10G0518000 | Non-protein coding transcript. (Os10t0518000-00) | -1.83595 | -1.62627 | - | - | - |
| OS03G0756300 | X8 domain containing protein. (Os03t0756300-01) | -2.02837 | -1.61674 | - | - | - |
| OS01G0247900 | AWPM-19-like family protein. (Os01t0247900-01) | -1.5362 | -1.60929 | -1.41738 | - | - |
| OS10G0164001 | Non-protein coding transcript. (Os10t0164001-00) | -2.34473 | -1.60809 | - | 1.16978 | - |
| OS09G0567500 | Similar to Fatty acyl coA reductase. (Os09t0567500-01);Similar to Fatty acyl coA reductase. (Os09t0567500-02) | -2.08468 | -1.60409 | -1.12032 | - | - |
| OS03G0574600 | Conserved hypothetical protein. (Os03t0574600-01);Conserved hypothetical protein. (Os03t0574600-02) | -1.84407 | -1.60308 | - | - | - |
| OS08G0478000 | Similar to mucin-2. (Os08t0478000-01);Conserved hypothetical protein. (Os08t0478000-02) | -1.94486 | -1.57855 | -1.13893 | 0.594667 | - |
| OS02G0797400 | MCM family protein. (Os02t0797400-01) | -1.57715 | -1.55045 | - | - | - |
| OS07G0617000 | Similar to Ethylene response factor 2. (Os07t0617000-01);Similar to Ethylene response factor 2. (Os07t0617000-02) | -1.54768 | -1.54874 | -0.781239 | 0.899457 | - |
| OS10G0396300 | Similar to MOB1. (Os10t0396300-01) | -1.96324 | -1.5399 | -1.32697 | - | - |
| OS06G0666500 | Zinc finger, RING/FYVE/PHD-type domain containing protein. (Os06t0666500-00) | -2.34242 | -1.5263 | -1.34087 | - | - |
| OS07G0257200 | Manganese and Cadmium transporter, Mn and Cd uptake (Os07t0257200-01) | -2.72228 | -1.52477 | -1.17227 | - | - |
| OS01G0893400 | Zinc finger, TAZ-type domain containing protein. (Os01t0893400-01);Zinc finger, TAZ-type domain containing protein. (Os01t0893400-02 | -1.57056 | -1.51616 | -1.04716 | - | - |
| OS01G0838350 | Conserved hypothetical protein. (Os01t0838350-01) | -1.98042 | -1.51439 | - | - | - |

| | | | | | | |
|---|---|---|---|---|---|---|
| OS02G0483500 | Transferase family protein. (Os02t0483500-01) | -1.74102 | -1.51096 | - | - | - |
| OS01G0529800 | Similar to Very-long-chain fatty acid condensing enzyme CUT1. (Os01t0529800-01) | -1.81073 | -1.50806 | -1.2713 | - | - |
| OS11G0649600 | Uncharacterised protein family UPF0497, trans-membrane plant subgroup domain containing protein. (Os11t0649600-01) | -2.00524 | -1.50501 | -1.16813 | - | - |
| OS01G0807900 | Similar to Dihydropyrimidinase (Dihydropyrimidine amidohydrolase) (EC 3.5.2.2). (Os01t0807900-01);D-hydantoinase domain containing | 1.53259 | 1.55258 | - | -1.13606 | -1.25017 |
| OS01G0715400 | Hypothetical conserved gene. (Os01t0715400-01);Conserved hypothetical protein. (Os01t0715400-02);Glycoside hydrolase, subgroup, c | 1.95147 | 1.5551 | 1.03879 | - | - |
| OS02G0194700 | Similar to Lipoxygenase 2.3, chloroplast precursor (EC 1.13.11.12) (LOX2:Hv:3). (Os02t0194700-01);Similar to Lipoxygenase. (Os02t0194 | 1.99897 | 1.57291 | 1.03866 | - | - |
| OS03G0244200 | Similar to thaumatin-like protein 1. (Os03t0244200-01);Similar to Thaumatin-like protein. (Os03t0244200-02) | 1.85872 | 1.5773 | 1.38124 | -1.00673 | -1.03954 |
| OS04G0167800 | Similar to Chalcone reductase homologue (Fragment). (Os04t0167800-01) | 1.63124 | 1.58518 | - | - | -1.22724 |
| OS05G0566800 | Cold acclimation protein COR413-TM1. (Os05t0566800-01) | 1.88544 | 1.58701 | - | - | -1.18194 |
| OS08G0442900 | Hypothetical protein. (Os08t0442900-01) | 2.40702909 | 1.60475929 | 1.31564738 | - | - |
| OS02G0580900 | TGF-beta receptor, type I/II extracellular region family protein. (Os02t0580900-01);Similar to peptide transporter PTR2. (Os02t0580900- | 2.10652 | 1.65468 | 1.40035 | - | - |
| OS06G0264500 | TGF-beta receptor, type I/II extracellular region family protein. (Os06t0264500-01) | 1.79022 | 1.66502 | - | - | -1.1191 |
| OS06G0698785 | Similar to Choline monooxygenase. (Os06t0698785-01) | 2.30546 | 1.67266 | - | -1.29311 | -1.72271 |
| OS01G0667200 | Mitochondrial sulfur dioxygenase, Abiotic stress response (Os01t0667200-01);Similar to Glyoxalase II. (Os01t0667200-02) | 1.61479 | 1.67633 | 1.06824 | -0.684898 | -0.806744 |
| OS03G0154000 | Aromatic-ring hydroxylase family protein. (Os03t0154000-01) | 1.54118 | 1.67722 | 0.933871 | - | - |
| OS01G0105900 | Carbohydrate/purine kinase domain containing protein. (Os01t0105900-01) | 1.71079 | 1.681 | 1.17034 | - | - |
| OS05G0557200 | Armadillo-type fold domain containing protein. (Os05t0557200-01);Armadillo-type fold domain containing protein. (Os05t0557200-02) | 1.64121 | 1.68834 | 0.851835 | - | -0.975552 |
| OS09G0555500 | Similar to Chloroplast phytoene synthase 3. (Os09t0555500-01) | 1.58839 | 1.72173 | 0.871318 | - | - |
| OS09G0419100 | Hypothetical protein. (Os09t0419100-01);Non-protein coding transcript. (Os09t0419100-02);Hypothetical gene. (Os09t0419100-03);Nor | 2.41431 | 1.72348 | - | - | -0.602205 |
| OS06G0129100 | Short-chain dehydrogenase/reductase SDR family protein. (Os06t0129100-01) | 2.94707 | 1.74352 | 1.43801 | - | - |
| OS08G0408500 | Pathogenesis-related transcriptional factor and ERF domain containing protein. (Os08t0408500-01) | 1.57005 | 1.75972 | - | - | - |
| OS05G0550300 | Similar to Lipid transfer protein (Fragment). (Os05t0550300-01) | 1.77151 | 1.76626 | 1.46773 | -1.63631 | -2.15177 |
| OS08G0405700 | Similar to Copper chaperone homolog CCH. (Os08t0405700-01) | 1.80686 | 1.79534 | -1.44255 | - | - |
| OS07G0190800 | Similar to Thioredoxin h. (Os07t0190800-01) | 2.3552 | 1.79702 | 1.37374 | - | - |
| OS03G0807900 | Chaperonin-like RbcX family protein. (Os03t0807900-01);Chaperonin-like RbcX family protein. (Os03t0807900-02) | 1.57668 | 1.80618 | - | - | - |
| OS08G0560000 | Similar to gibberellin 20 oxidase 2. (Os08t0560000-01) | 1.96186 | 1.80773 | - | - | - |
| OS06G0728700 | Homeodomain-like containing protein. (Os06t0728700-01) | 2.80037 | 1.80833 | 1.41237 | - | - |
| OS03G0817200 | Amino acid transporter, transmembrane domain containing protein. (Os03t0817200-01) | 2.34447 | 1.84059 | 0.986716 | - | - |
| OS09G0572400 | Similar to Abcf2-prov protein. (Os09t0572400-01) | 1.71791 | 1.84827 | 1.29611 | - | - |
| OS04G0630300 | NAD(P)-binding domain containing protein. (Os04t0630300-01) | 1.83709 | 1.85376 | 0.866219 | - | -0.945785 |
| OS04G0526800 | GRAM domain containing protein. (Os04t0526800-01) | 2.18504 | 1.91352 | - | - | -1.03992 |
| OS08G0425800 | Similar to DRP2 protein (Fragment). (Os08t0425800-01);Hypothetical conserved gene. (Os08t0425800-02) | 1.55389 | 1.9196 | - | - | - |
| OS12G0197600 | Hypothetical gene. (Os12t0197600-01) | 1.79284 | 1.92621 | - | - | - |
| OS04G0443200 | Protein of unknown function DUF538 family protein. (Os04t0443200-01) | 2.40851 | 1.95069 | - | - | - |
| OS07G0638100 | Six-bladed beta-propeller, TolB-like domain containing protein. (Os07t0638100-01) | 1.91231 | 2.02411 | - | - | - |
| OS05G0267800 | Cellular retinaldehyde-binding/triple function, C-terminal domain containing protein. (Os05t0267800-01) | 1.58774 | 2.03341 | 1.06678 | - | - |
| OS10G0422600 | Protein of unknown function DUF581 family protein. (Os10t0422600-01) | 1.62515 | 2.04314 | - | - | - |
| OS05G0171200 | Embryo-specific 3 family protein. (Os05t0171200-01) | 2.14479 | 2.06025 | 1.14869 | - | - |
| OS01G0944900 | Similar to Glucan endo-1,3-beta-D-glucosidase. (Os01t0944900-01) | 2.96158 | 2.07456 | 1.288 | - | - |
| OS06G0710300 | Uncharacterised conserved protein UCP022348 domain containing protein. (Os06t0710300-01) | 1.64866 | 2.11153 | - | - | - |
| OS05G0195200 | CCCH-tandem zinc finger protein, Photomorphogenesis and ABA responses, Abiotic stress tolerance (Os05t0195200-01) | 3.76295 | 2.12154 | - | - | - |
| OS09G0367700 | Similar to GST6 protein (EC 2.5.1.18). (Os09t0367700-01) | 1.86413 | 2.14068 | 1.24595 | - | -2.45145 |
| OS02G0766700 | bZIP transcription factor, Regulation of ABA signaling and biosynthesis, Drought resistance (Os02t0766700-01) | 1.75318 | 2.15479 | - | - | - |
| OS05G0244700 | Aminotransferase, class IV family protein. (Os05t0244700-01);Aminotransferase, class IV family protein. (Os05t0244700-02) | 3.25318 | 2.16811 | - | - | - |
| OS01G0583100 | Similar to Protein phosphatase 2C. (Os01t0583100-01) | 2.15293 | 2.20283 | 1.42884 | - | - |
| OS06G0253100 | Heat shock protein Hsp20 domain containing protein. (Os06t0253100-01) | 2.24569 | 2.24354 | - | - | - |
| OS05G0206100 | Conserved hypothetical protein. (Os05t0206100-00) | 2.14807181 | 2.26333193 | 1.09083988 | - | - |

| | | | | | | |
|---|---|---|---|---|---|---|
| OS09G0441000 | Peptidase S8, subtilisin-related domain containing protein. (Os09t0441000-00) | 1.78866 | 2.30029 | - | - | - |
| OS01G0863300 | Similar to MCB2 protein. (Os01t0863300-01) | 2.31011 | 2.33434 | - | -1.14234 | -1.96421 |
| OS02G0770800 | Similar to Nitrate reductase [NAD(P)H] (EC 1.7.1.2). (Os02t0770800-01) | 2.3244 | 2.36517 | - | -0.99182 | -1.55906 |
| OS02G0162600 | Conserved hypothetical protein. (Os02t0162600-01) | 2.04541 | 2.42825 | 1.46205 | - | - |
| OS03G0200200 | Conserved hypothetical protein. (Os03t0200200-01) | 3.87756 | 2.44038 | 1.49123 | -1.44076 | -1.9624 |
| OS03G0724700 | Similar to cDNA clone:002-104-C06, full insert sequence. (Os03t0724700-01);Similar to cDNA clone:002-144-A12, full insert sequence. (C | 2.40733 | 2.49823 | 1.4482 | - | - |
| OS03G0663400 | Similar to Thaumatin-like protein. (Os03t0663400-02) | 3.02689 | 2.55698 | - | - | -1.65184 |
| OS05G0477900 | Similar to nonspecific lipid-transfer protein. (Os05t0477900-01) | 1.98418 | 2.58963 | - | - | - |
| OS12G0258700 | Cupredoxin domain containing protein. (Os12t0258700-01) | 4.0818 | 2.94202 | - | - | - |
| OS10G0463800 | Domain of unknown function, DUF1338, containing green-plant-unique protein, Regulation of starch synthesis and amyloplast developm | 2.81222 | 3.30293 | - | - | - |
| OS07G0188700 | Similar to EXO. (Os07t0188700-01);Similar to oxidoreductase. (Os07t0188700-02);Hypothetical conserved gene. (Os07t0188700-03) | 2.99923 | 3.48077 | - | - | - |
| OS12G0569500 | Thaumatin, pathogenesis-related family protein. (Os12t0569500-01) | 2.60169 | 3.54066 | 0.669619 | - | - |
| OS07G0209100 | Similar to Seed imbibition protein (Fragment). (Os07t0209100-01) | 3.10463 | 3.7293 | 1.39924 | - | - |
| OS01G0256500 | Similar to ZnI. (Os01t0256500-02) | 2.69842 | 3.93591 | - | - | -2.03054 |
| OS09G0455300 | Similar to INDEHISCENT protein. (Os09t0455300-01) | 3.28316 | 4.08938 | 1.36869 | - | - |
| OS10G0465700 | Similar to Beta-amylase PCT-BMYI (EC 3.2.1.2). (Os10t0465700-01) | 2.97267 | 4.42394 | 1.46334 | - | - |
| OS03G0826900 | Conserved hypothetical protein. (Os03t0826900-01) | 5.49997341 | 4.84929375 | - | - | -2.47107 |
| OS02G0537000 | Pectinesterase inhibitor domain containing protein. (Os02t0537000-01) | 3.08069 | 4.93281 | 1.48288 | - | - |
| OS04G0415800 | Similar to OSIGBa0092M08.2 protein. (Os04t0415800-00) | 7.45125233 | 7.69315935 | - | - | - |

| 3. Stress-responsive in SLR1-OX | | | | | |
|---|---|---|---|---|---|
| Identifier | Description | FC.SmSs | FC.SSmSSs | FC.NmNs | FC.NmSm |
| OS02G0726000 | FAS1 domain domain containing protein. (Os02t0726000-01) | -2.98251 | - | - | - |
| OS06G0695900 | Zinc finger, RING-type domain containing protein. (Os06t0695900-01) | -2.95454 | - | - | - |
| OS08G0239300 | Dienelactone hydrolase domain containing protein. (Os08t0239300-01) | -2.83415 | - | - | - |
| OS12G0191500 | Similar to Peroxidase 43 precursor (EC 1.11.1.7) (Atperox P43). (Os12t0191500-01) | -2.66447 | -1.06856 | -1.46788 | - |
| OS11G0687100 | von Willebrand factor, type A domain containing protein. (Os11t0687100-01) | -2.53163 | -1.35515 | 0.641604 | - |
| OS01G0669100 | S-domain receptor-like kinase, OsPR1b-interacting factor, Resistance protein, Grain yield, Disease resistance (Os01t0669100-01);Simila | -2.50721 | - | -1.27078 | - |
| OS11G0109700 | Protein of unknown function Cys-rich family protein. (Os11t0109700-01);Protein of unknown function Cys-rich family protein. (Os11t01 | -2.4863 | - | - | - |
| OS01G0831250 | Hypothetical gene. (Os01t0831250-00) | -2.48198 | - | - | - |
| OS04G0687800 | Protein of unknown function DUF6, transmembrane domain containing protein. (Os04t0687800-01) | -2.3297 | - | - | - |
| OS08G0136800 | Protein of unknown function DUF26 domain containing protein. (Os08t0136800-01) | -2.19609 | - | - | - |
| OS01G0751600 | Class III lipase, Regulation of tillering, plant height, and spikelet fertility (Os01t0751600-01) | -2.16203 | - | - | - |
| OS07G0530400 | Ribosomal protein S14, conserved site domain containing protein. (Os07t0530400-01) | -2.13849 | - | - | - |
| OS04G0357000 | Conserved hypothetical protein. (Os04t0357000-00) | -2.13733 | - | - | - |
| OS11G0152025 | Conserved hypothetical protein. (Os11t0152025-00) | -2.12086 | - | - | - |
| OS03G0199500 | Conserved hypothetical protein. (Os03t0199500-01) | -2.11664 | - | - | - |
| OS03G0733600 | SSXT family protein. (Os03t0733600-01) | -2.10814 | - | - | - |
| OS01G0133200 | Conserved hypothetical protein. (Os01t0133200-01) | -2.04376 | - | - | - |
| OS01G0827300 | Putative laccase precursor, Abiotic stress response (Os01t0827300-01) | -2.00174 | -1.35768 | -1.2399 | - |
| OS12G0125000 | Myb factor. (Os12t0125000-01) | -1.9944 | -1.27304 | -0.751127 | - |
| OS08G0162700 | Aldehyde dehydrogenase, conserved site domain containing protein. (Os08t0162700-01) | -1.98471 | - | -1.24187 | - |
| OS09G0433650 | Tobacco mosaic virus coat protein family protein. (Os09t0433650-01) | -1.95953 | - | - | - |
| OS04G0574200 | FAS1 domain domain containing protein. (Os04t0574200-01) | -1.93925 | - | - | - |
| OS01G0942200 | Cyclin-like F-box domain containing protein. (Os01t0942200-01) | -1.89293 | - | - | - |
| OS03G0305800 | Galactosyl transferase family protein. (Os03t0305800-02) | -1.87409 | - | - | - |
| OS04G0410600 | Similar to Purple acid phosphatase. (Os04t0410600-01) | -1.83918 | - | - | - |
| OS07G0619400 | EF-Hand type domain containing protein. (Os07t0619400-01) | -1.8315 | - | - | - |
| OS07G0484800 | Similar to Adenine phosphoribosyltransferase (EC 2.4.2.7)-like protein. (Os07t0484800-01) | -1.82236 | -1.45073 | - | - |
| OS04G0116200 | Protein of unknown function DUF827, plant family protein. (Os04t0116200-01) | -1.80936 | -1.06272 | - | - |
| OS11G0559600 | Conserved hypothetical protein. (Os11t0559600-01) | -1.79004 | - | - | - |
| OS11G0661600 | Similar to Peroxidase (EC 1.11.1.7). (Os11t0661600-01) | -1.78256 | -1.20808 | -1.3832 | - |
| OS02G0666700 | Permease for cytosine/purines, uracil, thiamine, allantoin family protein. (Os02t0666700-01) | -1.78236 | - | -1.06714 | - |
| OS02G0666600 | Harpin-induced 1 domain containing protein. (Os02t0666600-01) | -1.75914 | - | -0.968684 | - |
| OS01G0892800 | Similar to Ankyrin-kinase. (Os01t0892800-01);Integrin-linked protein kinase domain containing protein. (Os01t0892800-02) | -1.7316 | - | - | - |
| OS03G0151300 | Similar to JmjC domain containing protein, expressed. (Os03t0151300-01) | -1.72067 | - | - | - |
| OS11G0169800 | Similar to Long-chain-fatty-acid--CoA ligase 4 (EC 6.2.1.3) (Long-chain acyl-CoA synthetase 4) (LACS 4). (Os11t0169800-01) | -1.68099 | - | - | - |
| OS03G0135700 | Transcriptional activator Rb homolog (Fragment). (Os03t0135700-01);Transcriptional activator Rb homolog (Fragment). (Os03t013570( | -1.68033 | - | - | - |
| OS01G0638600 | UDP-glucuronosyl/UDP-glucosyltransferase family protein. (Os01t0638600-00) | -1.68012 | - | -1.09269 | - |
| OS01G0896300 | Hypothetical conserved gene. (Os01t0896300-00) | -1.67752 | - | - | - |
| OS03G0296500 | Similar to Glycosyl hydrolases family 17 protein, expressed. (Os03t0296500-00) | -1.67409 | - | - | - |
| OS11G0431100 | Similar to Serine carboxypeptidase family protein. (Os11t0431100-00) | -1.6704869 | - | -0.9843633 | - |
| OS05G0448300 | Similar to glycerol-3-phosphate acyltransferase 8. (Os05t0448300-01) | -1.66002 | -1.06992 | -1.15649 | - |
| OS07G0105900 | NB-ARC domain containing protein. (Os07t0105900-01) | -1.65845 | -1.13051 | -0.895418 | - |
| OS05G0435800 | Similar to Subtilisin-like protease. (Os05t0435800-01) | -1.65692 | - | - | 0.651401 |

| | | | | | |
|---|---|---|---|---|---|
| OS01G0300900 | Galactose oxidase/kelch, beta-propeller domain containing protein. (Os01t0300900-01);Hypothetical conserved gene. (Os01t0300900- | -1.65141 | - | - | - |
| OS04G0664400 | Auxin response factor 1. (Os04t0664400-01);Similar to Auxin response factor 11. (Os04t0664400-02) | -1.62495 | - | - | - |
| OS04G0381400 | Similar to OSIGBa0092E09.2 protein. (Os04t0381400-00) | -1.60478 | - | -0.984116 | - |
| OS01G0832100 | Esterase, SGNH hydrolase-type domain containing protein. (Os01t0832100-01) | -1.60064 | -1.47085 | - | - |
| OS06G0229400 | Conserved hypothetical protein. (Os06t0229400-00) | -1.59722 | - | - | - |
| OS07G0176400 | Hypothetical protein. (Os07t0176400-00) | -1.59427 | - | - | - |
| OS06G0115700 | Conserved hypothetical protein. (Os06t0115700-01);Hypothetical conserved gene. (Os06t0115700-02) | -1.59144 | -1.2418 | - | - |
| OS08G0244500 | Similar to hydrolase, hydrolyzing O-glycosyl compounds. (Os08t0244500-00) | -1.5658 | - | -1.19531 | - |
| OS01G0331100 | Similar to Alpha-L-fucosidase 2. (Os01t0331100-01) | -1.54715 | -1.49087 | - | - |
| OS01G0183600 | Cytochrome P450 family protein. (Os01t0183600-01) | -1.53756 | - | - | - |
| OS03G0119100 | Similar to Phospholipase D beta 2. (Os03t0119100-01) | -1.53633 | - | -1.29825 | - |
| OS03G0149300 | Protein of unknown function DUF6, transmembrane domain containing protein. (Os03t0149300-01) | -1.53592 | -1.2228 | - | - |
| OS02G0189300 | Esterase, SGNH hydrolase-type domain containing protein. (Os02t0189300-01) | -1.50964 | - | - | - |
| OS02G0753700 | VQ domain containing protein. (Os02t0753700-01) | -1.50925 | - | - | - |
| OS07G0101000 | Cupredoxin domain containing protein. (Os07t0101000-00) | -1.50596 | - | -1.36777 | - |
| OS10G0163370 | Lecithin:cholesterol acyltransferase family protein. (Os10t0163370-01) | -1.50399 | -1.48346 | - | - |
| OS01G0259300 | Non-protein coding transcript. (Os01t0259300-01) | -1.50144 | -1.31826 | - | - |
| OS11G0123033 | Amidase family protein. (Os11t0123033-01) | 1.51233 | - | - | - |
| OS01G0731100 | Similar to Pathogen-related protein. (Os01t0731100-01) | 1.51266 | - | - | - |
| OS03G0287100 | Similar to Phosphatidylinositol transfer protein, expressed. (Os03t0287100-01) | 1.51321 | 1.47451 | 0.854047 | - |
| OS02G0126500 | Pentatricopeptide repeat domain containing protein. (Os02t0126500-01) | 1.51476 | - | - | - |
| OS09G0513800 | Similar to H0425E08.1 protein. (Os09t0513800-01) | 1.51797 | - | - | - |
| OS06G0140200 | Leucine-rich repeat, plant specific containing protein. (Os06t0140200-01) | 1.52944 | - | - | -1.84692 |
| OS06G0563900 | Similar to Diacylglycerol acyltransferase. (Os06t0563900-01) | 1.53124 | - | -1.20387 | -2.2614 |
| OS09G0453700 | Peptidase S9A, prolyl oligopeptidase family protein. (Os09t0453700-01) | 1.532 | - | - | - |
| OS09G0316000 | Similar to cupin, RmlC-type. (Os09t0316000-01) | 1.53231 | - | 0.699889 | -1.45131 |
| OS06G0525200 | Similar to avr9/Cf-9 rapidly elicited protein 194. (Os06t0525200-00) | 1.53307 | - | - | - |
| OS01G0278200 | Similar to Ran-binding protein 17. (Os01t0278200-01) | 1.53373 | - | - | - |
| OS11G0541600 | Protein of unknown function DUF247, plant domain containing protein. (Os11t0541600-00) | 1.53469 | - | - | -1.47982 |
| OS10G0502500 | Cytochrome b5 domain containing protein. (Os10t0502500-04) | 1.53562 | - | -1.08592 | -0.824409 |
| OS07G0685700 | Transcription factor, Response to ethylene stimulus, Wound signaling (Os07t0685700-01) | 1.53731 | 0.939897 | - | - |
| OS04G0206700 | UDP-glucuronosyl/UDP-glucosyltransferase family protein. (Os04t0206700-01);UDP-glucuronosyl/UDP-glucosyltransferase family prote | 1.54175 | - | - | -1.43234 |
| OS05G0170950 | Similar to Plastid phosphoenolpyruvate/phosphate translocator. (Os05t0170950-00) | 1.54342 | - | - | - |
| OS01G0508000 | Similar to Beta-glucosidase. (Os01t0508000-01);Similar to Beta-mannosidase 2. (Os01t0508000-02) | 1.54697 | 1.3431 | 1.2435 | - |
| OS06G0127500 | Hypothetical conserved gene. (Os06t0127500-01) | 1.54767 | - | - | - |
| OS07G0522500 | Similar to PDR6 ABC transporter. (Os07t0522500-01) | 1.5519 | - | -0.577144 | -2.71308 |
| OS07G0618400 | Similar to Leucine-rich repeat transmembrane protein kinase 1 (Fragment). (Os07t0618400-01);Similar to Leucine-rich repeat transmer | 1.55973 | 1.32756 | 1.21299 | - |
| OS03G0133800 | Similar to predicted protein. (Os03t0133800-01) | 1.56023 | - | - | - |
| OS02G0615500 | Protein kinase, core domain containing protein. (Os02t0615500-01);Hypothetical conserved gene. (Os02t0615500-02) | 1.56224 | - | - | -1.44781 |
| OS02G0701700 | Conserved hypothetical protein. (Os02t0701700-01) | 1.57259 | - | - | - |
| OS08G0337100 | Conserved hypothetical protein. (Os08t0337100-01) | 1.57593 | - | - | -1.53554 |
| OS06G0724400 | Zinc finger, Zim17-type family protein. (Os06t0724400-01) | 1.57755 | 1.08097 | - | - |
| OS02G0244100 | RING-type E3 ubiquitin ligase, Regulation of grain width and weight (Os02t0244100-01) | 1.57888 | - | - | - |
| OS04G0586200 | Similar to H0307D04.13 protein. (Os04t0586200-01) | 1.591 | 1.26856 | - | - |
| OS05G0312000 | Hypothetical conserved gene. (Os05t0312000-01) | 1.5917 | - | 1.23572 | - |
| OS09G0426100 | Similar to cDNA, clone: J075189B12, full insert sequence. (Os09t0426100-01);Similar to cDNA, clone: J075189B12, full insert sequence. | 1.60151 | - | - | - |

| | | | | | |
|---|---|---|---|---|---|
| OS07G0240400 | Similar to MEE60 (maternal effect embryo arrest 60). (Os07t0240400-01) | 1.6027 | - | - | - |
| OS01G0770700 | Similar to Copper transporter 1. (Os01t0770700-00) | 1.61431 | - | -0.966774 | -0.801307 |
| OS03G0313800 | Hypothetical conserved gene. (Os03t0313800-00) | 1.61712 | - | -1.32424 | - |
| OS08G0497900 | Conserved hypothetical protein. (Os08t0497900-01) | 1.61848 | - | - | - |
| OS12G0635500 | Protein of unknown function DUF266, plant domain containing protein. (Os12t0635500-01) | 1.61925 | - | -1.24348 | -2.05047 |
| OS03G0342900 | Dormancyauxin associated family protein. (Os03t0342900-02);Dormancyauxin associated family protein. (Os03t0342900-04) | 1.62494 | 0.830549 | - | - |
| OS04G0475600 | 2OG-Fe(II) oxygenase domain containing protein. (Os04t0475600-01) | 1.63876 | 1.39011 | - | - |
| OS03G0659300 | Glyoxalase/bleomycin resistance protein/dioxygenase domain containing protein. (Os03t0659300-01) | 1.64138 | - | -1.16714 | - |
| OS03G0306900 | Haem oxygenase-like, multi-helical domain containing protein. (Os03t0306900-01);Similar to Seed maturation protein PM36. (Os03t03 | 1.65857 | 1.43872 | - | - |
| OS06G0235300 | ATPase, F1/V1/A1 complex, alpha/beta subunit, nucleotide-binding domain containing protein. (Os06t0235300-01) | 1.66417 | - | - | - |
| OS06G0255900 | Conserved hypothetical protein. (Os06t0255900-01) | 1.66903 | - | - | - |
| OS07G0563600 | FAR1 domain containing protein. (Os07t0563600-01) | 1.66993 | - | - | -1.74246 |
| OS09G0378700 | Similar to ubiquitin-protein ligase. (Os09t0378700-00) | 1.67058204 | 0.86117725 | - | - |
| OS03G0298300 | Uncharacterised protein family UPF0497, trans-membrane plant domain containing protein. (Os03t0298300-01);Uncharacterised prote | 1.67181 | 1.25127 | - | -0.853574 |
| OS08G0503700 | Sodium/sulphate symporter family protein. (Os08t0503700-01);Sodium/sulphate symporter family protein. (Os08t0503700-02) | 1.674 | 0.834338 | -0.653494 | -0.718588 |
| OS07G0223766 | Hypothetical conserved gene. (Os07t0223766-01) | 1.67747 | - | - | - |
| OS11G0303800 | Hypothetical protein. (Os11t0303800-01) | 1.67764 | - | 1.42918 | - |
| OS02G0570400 | Similar to Ent-kaurene synthase 1A. (Os02t0570400-01) | 1.67986 | - | -0.84182 | -3.03213 |
| OS06G0264650 | Hypothetical conserved gene. (Os06t0264650-00) | 1.69154 | - | - | - |
| OS02G0781600 | Hypothetical conserved gene. (Os02t0781600-00) | 1.71852 | - | - | - |
| OS08G0224000 | NB-ARC domain containing protein. (Os08t0224000-01) | 1.72301 | - | - | -1.68338 |
| OS04G0466600 | Agmatine deiminase domain containing protein. (Os04t0466600-01) | 1.72919 | 1.2072 | 1.44044 | - |
| OS07G0535800 | Similar to SRK15 protein (Fragment). (Os07t0535800-01) | 1.75073 | - | - | - |
| OS03G0397300 | Metallophosphoesterase domain containing protein. (Os03t0397300-01) | 1.75324 | - | - | - |
| OS03G0146400 | Pheophorbide a oxygenase, Leaf senescence, Wound responses (Os03t0146400-01);Similar to Lethal leaf-spot 1 (Fragment). (Os03t014 | 1.75349 | 1.28921 | - | - |
| OS04G0470150 | Hypothetical conserved gene. (Os04t0470150-00) | 1.75599 | - | - | - |
| OS06G0694200 | Esterase, SGNH hydrolase-type domain containing protein. (Os06t0694200-01) | 1.75801 | 1.10876 | - | - |
| OS04G0504700 | Conserved hypothetical protein. (Os04t0504700-01) | 1.75813 | - | - | - |
| OS11G0186800 | Similar to inorganic phosphate cotransporter. (Os11t0186800-01) | 1.78303 | - | - | - |
| OS06G0701600 | Monovalent cation transporter, Na+ and K+ transport (Os06t0701600-01) | 1.78462 | - | - | - |
| OS06G0147000 | Conserved hypothetical protein. (Os06t0147000-01) | 1.79098 | 1.37536 | 0.980388 | - |
| OS12G0638300 | Similar to Peptide transporter. (Os12t0638300-01) | 1.7923 | 1.11479 | - | -1.15061 |
| OS08G0499900 | Conserved hypothetical protein. (Os08t0499900-01) | 1.79409 | - | - | - |
| OS05G0568100 | Similar to Iron sulfur cluster assembly protein 1, mitochondrial precursor (Iron sulfur cluster scaffold protein 1). (Os05t0568100-01) | 1.80642 | 1.07776 | - | - |
| OS08G0290700 | Winged helix repressor DNA-binding domain containing protein. (Os08t0290700-01) | 1.80851 | - | - | -1.59345 |
| OS10G0374600 | Hypothetical conserved gene. (Os10t0374600-00) | 1.81159 | - | - | - |
| OS04G0683700 | 4-coumarate-Co-A ligase (4CL) like protein, Adenosine monophosphate binding protein, Regulation of rice blast resistance, floret devel | 1.81827 | 1.28326 | - | - |
| OS01G0869200 | Magnesium transporter, Mg-mediated aluminum tolerance (Os01t0869200-01) | 1.82269 | - | -1.20327 | -1.61982 |
| OS04G0635400 | Conserved hypothetical protein. (Os04t0635400-00) | 1.8243 | 1.12688 | 1.22457 | - |
| OS04G0628000 | Protein of unknown function DUF794, plant family protein. (Os04t0628000-01) | 1.82536 | - | - | - |
| OS06G0713800 | Alpha-amylase isozyme 2A precursor (EC 3.2.1.1) (1,4-alpha-D-glucan glucanohydrolase). (Os06t0713800-01) | 1.83516 | - | - | - |
| OS03G0184550 | Similar to Dihydroflavonol-4-reductase. (Os03t0184550-01) | 1.83719 | - | - | - |
| OS04G0529500 | Similar to C-terminal domain phosphatase-like 1. (Os04t0529500-01);Similar to OSIGBa0155K17.3 protein. (Os04t0529500-02);Similar t | 1.84235 | - | - | - |
| OS04G0320700 | Similar to Glucosyltransferase (Fragment). (Os04t0320700-01);UDP-glucuronosyl/UDP-glucosyltransferase domain containing protein. ( | 1.84847 | 1.03416 | - | - |
| OS02G0735700 | Similar to ankyrin repeat family protein. (Os02t0735700-01) | 1.86438 | - | - | -1.29087 |
| OS01G0948200 | Similar to GRAS family transcription factor containing protein. (Os01t0948200-00) | 1.86829 | - | -1.23856 | - |

| | | | | | |
|---|---|---|---|---|---|
| OS03G0118450 | Non-protein coding transcript. (Os03t0118450-00) | 1.88196 | 1.3857 | - | - |
| OS05G0239900 | Non-protein coding transcript. (Os05t0239900-01) | 1.90754 | - | - | - |
| OS04G0537100 | Similar to Auxin-induced protein X15. (Os04t0537100-01) | 1.91394 | - | -1.47048 | - |
| OS02G0210600 | Hypothetical protein. (Os02t0210600-01) | 1.92218 | - | - | - |
| OS05G0169200 | WD40 repeat-like domain containing protein. (Os05t0169200-01) | 1.93226 | - | - | - |
| OS12G0284850 | Hypothetical gene. (Os12t0284850-01) | 1.99911 | - | - | -2.2846 |
| OS01G0844300 | Similar to Peptidylprolyl isomerase. (Os01t0844300-01) | 2.02827 | - | - | - |
| OS03G0269900 | Protein of unknown function DUF604 family protein. (Os03t0269900-01) | 2.04926 | 1.30654 | -0.968211 | -1.5685 |
| OS01G0692100 | Glutathione S-transferase, C-terminal-like domain containing protein. (Os01t0692100-01) | 2.05884 | - | - | - |
| OS02G0684400 | Similar to OXS3 (OXIDATIVE STRESS 3). (Os02t0684400-01) | 2.08746 | - | - | - |
| OS08G0191433 | Starch synthase, Starch biosynthesis (Os08t0191433-01) | 2.12637 | - | - | - |
| OS01G0314800 | Late embryogenesis abundant protein 3 family protein. (Os01t0314800-01) | 2.13673 | 1.49884 | -1.11407 | -1.4599 |
| OS04G0419700 | Similar to H0525E10.7 protein. (Os04t0419700-00) | 2.14039 | - | 0.886736 | - |
| OS03G0168200 | Similar to F16A14.21. (Os03t0168200-01) | 2.15114 | 1.22063 | - | -1.34831 |
| OS01G0848200 | Similar to Delta 1-pyrroline-5-carboxylate synthetase (P5CS) [Includes: Glutamate 5-kinase (EC 2.7.2.11) (Gamma-glutamyl kinase) (GK) | 2.15453 | - | 1.26052 | - |
| OS10G0548100 | Similar to DM280 protein. (Os10t0548100-01) | 2.15807522 | - | - | - |
| OS06G0133400 | Conserved hypothetical protein. (Os06t0133400-01) | 2.18247 | - | - | - |
| OS01G0225600 | Similar to Dehydrin. (Os01t0225600-00) | 2.18493 | - | - | - |
| OS04G0677300 | Harpin-induced 1 domain containing protein. (Os04t0677300-01) | 2.18501 | - | - | -2.2962 |
| OS03G0141200 | Similar to Beta-amylase PCT-BMYI (EC 3.2.1.2). (Os03t0141200-01) | 2.18586 | - | - | - |
| OS03G0767900 | Uncharacterised protein family UPF0497, trans-membrane plant domain containing protein. (Os03t0767900-01) | 2.18797 | - | - | -1.45862 |
| OS01G0200300 | Similar to Homeobox-leucine zipper protein HOX29. (Os01t0200300-01) | 2.18838 | - | - | - |
| OS06G0166400 | Similar to TINY-like protein (AP2 domain containing protein RAP2.10) (Fragment). (Os06t0166400-01) | 2.27687 | - | - | - |
| OS05G0355450 | Hypothetical protein. (Os05t0355450-01) | 2.31661 | - | - | - |
| OS02G0686300 | Similar to H0307D04.3 protein. (Os02t0686300-01) | 2.32035 | - | -1.17586 | -2.82294 |
| OS03G0696300 | Similar to Nuclear transcription factor Y subunit A-1. (Os03t0696300-01);Similar to nuclear transcription factor Y subunit A-1. (Os03t06 | 2.3276 | 1.10436 | - | - |
| OS11G0644800 | Similar to Tyrosine/nicotianamine aminotransferases family protein, expressed. (Os11t0644800-00) | 2.34572 | 1.28591 | - | -1.34721 |
| OS04G0116800 | Similar to 3-ketoacyl-CoA synthase. (Os04t0116800-01) | 2.34842 | - | - | - |
| OS08G0240200 | Conserved hypothetical protein. (Os08t0240200-01) | 2.39021 | - | - | - |
| OS07G0218200 | Similar to terpene synthase 7. (Os07t0218200-00) | 2.41155 | - | - | - |
| OS05G0217700 | BURP domain containing protein. (Os05t0217700-01) | 2.43103 | 1.24685 | -1.30553 | -0.969808 |
| OS05G0374500 | Tetratricopeptide-like helical domain containing protein. (Os05t0374500-01) | 2.6347 | - | 1.40517 | - |
| OS06G0127100 | Dehydration-responsive element-binding protein 1C. (Os06t0127100-01) | 2.7058983 | - | - | -0.6782072 |
| OS05G0149200 | PWWP domain containing protein. (Os05t0149200-01) | 2.73117 | - | - | -3.00213 |
| OS10G0173000 | Conserved hypothetical protein. (Os10t0173000-01) | 2.75266 | 0.973211 | -1.14913 | -1.47836 |
| OS02G0291000 | Calcineurin B (Fragment). (Os02t0291000-00) | 2.79498 | - | - | - |
| OS12G0257600 | Similar to Laccase-25. (Os12t0257600-00) | 2.87947 | - | - | - |
| OS07G0249800 | Similar to IAA-amino acid hydrolase ILR1-like 8. (Os07t0249800-01) | 2.9722 | - | - | - |
| OS05G0466000 | Non-protein coding transcript. (Os05t0466000-01) | 3.03437 | - | - | - |
| OS07G0499800 | Ubiquitin-protein ligase E3 MDM2 domain containing protein. (Os07t0499800-01) | 3.12432 | - | - | - |
| OS04G0357700 | Hypothetical conserved gene. (Os04t0357700-00) | 3.36481 | - | - | - |
| OS01G0627800 | Similar to Cytochrome P450 monooxygenase CYP72A5 (Fragment). (Os01t0627800-01) | 3.63373 | - | - | - |
| OS05G0572700 | Similar to protein phosphatase 2C ABI1. (Os05t0572700-01);Similar to Protein phosphatase 2C. (Os05t0572700-02) | 3.66801 | - | - | - |
| OS08G0332600 | Disease resistance protein domain containing protein. (Os08t0332600-00) | 3.81352 | - | - | - |
| OS01G0692200 | Conserved hypothetical protein. (Os01t0692200-01) | 3.97616 | - | - | - |
| OS03G0799900 | Similar to sec12-like protein 1. (Os03t0799900-01) | 4.82895 | - | - | - |

| 4. Stress-resp in SUMO1SLR1-OX | | | | | |
|---|---|---|---|---|---|
| Identifier | Description | FC.SSmSSs | FC.SmSs | FC.NmNs | FC.NmSSm |
| OS08G0373000 | Hypothetical conserved gene. (Os08t0373000-00) | -3.74858 | - | - | - |
| OS04G0468600 | Hypothetical conserved gene. (Os04t0468600-01);Heavy metal transport/detoxification protein domain containing protein | -3.15701 | - | - | -1.24465 |
| OS11G0528400 | Similar to aminopeptidase. (Os11t0528400-01) | -3.08958 | - | - | - |
| OS02G0699433 | Hypothetical conserved gene. (Os02t0699433-00) | -2.97757 | - | - | - |
| OS02G0640100 | Hypothetical conserved gene. (Os02t0640100-00) | -2.87845 | - | - | - |
| OS03G0106400 | Similar to Branched-chain-amino-acid aminotransferase 5, chloroplast precursor (EC 2.6.1.42) (Atbcat-5). (Os03t0106400-0 | -2.51721 | - | - | - |
| OS02G0590200 | Lecithin:cholesterol/phospholipid:diacylglycerol acyltransferase domain containing protein. (Os02t0590200-01) | -2.50127 | - | - | - |
| OS01G0856000 | Similar to CDC6 protein. (Os01t0856000-01);Similar to predicted protein. (Os01t0856000-02) | -2.44386 | - | - | - |
| OS08G0508700 | Similar to EIL3. (Os08t0508700-01) | -2.41902 | - | - | - |
| OS03G0386800 | Similar to EIL3. (Os08t0508700-01) | -2.36459 | -1.44788 | -1.42313 | - |
| OS04G0223901 | Dimethylaniline monooxygenase, N-oxide-forming domain containing protein. (Os04t0223901-00) | -2.36377 | - | - | - |
| OS03G0146800 | Non-protein coding transcript. (Os03t0146800-01) | -2.36303 | -1.21896 | -1.22238 | 1.1619 |
| OS01G0711800 | Nucleosome assembly protein (NAP) family protein. (Os01t0711800-00) | -2.2633 | -1.33216 | -1.09439 | - |
| OS02G0445600 | Similar to Auxin-induced SAUR-like protein (Fragment). (Os02t0445600-01) | -2.23597 | -1.35757 | - | - |
| OS10G0403800 | Similar to DNA binding protein. (Os10t0403800-00) | -2.22672 | - | - | - |
| OS09G0132200 | Esterase, SGNH hydrolase-type domain containing protein. (Os09t0132200-01);Similar to anther-specific proline-rich prote | -2.21481 | - | - | - |
| OS01G0312800 | Similar to CEL4=CELLULASE 4 (Fragment). (Os01t0312800-01) | -2.16709 | - | - | - |
| OS02G0589900 | Hypothetical conserved gene. (Os02t0589900-00) | -2.14327 | - | - | - |
| OS05G0201700 | Hypothetical conserved gene. (Os05t0201700-00) | -2.1016702 | - | - | - |
| OS05G0217000 | Protein of unknown function DUF1070 family protein. (Os05t0217000-01) | -2.09434 | -1.46966 | -0.958217 | - |
| OS08G0206950 | Similar to Pectinesterase. (Os08t0206950-00) | -2.07097 | - | - | - |
| OS09G0466400 | Zinc finger homeodomain (ZF-HD) class homeobox transcription factor, Rice morphogenesis, Modulation of leaf rolling (Os | -2.05348 | - | -1.2325 | - |
| OS12G0189900 | Las1-like family protein. (Os12t0189900-01) | -2.04964 | - | -0.929916 | - |
| OS06G0158900 | Similar to Multidrug-resistance associated protein 3. (Os06t0158900-00) | -2.04941 | - | - | - |
| OS02G0230300 | En/Spm-like transposon protein (Protodermal factor 1). (Os02t0230300-01) | -2.04543 | -1.18173 | - | - |
| OS05G0573200 | Similar to Isocitrate dehydrogenase (Fragment). (Os05t0573200-00) | -2.04437 | -1.41877 | - | - |
| OS07G0668300 | Lipase, GDSL domain containing protein. (Os07t0668300-01) | -2.03416 | -1.46753 | -1.03154 | - |
| OS10G0438600 | Similar to Family II lipase EXL3. (Os10t0438600-01) | -2.02388 | - | - | - |
| OS07G0197100 | Similar to Hexokinase-4, chloroplastic. (Os07t0197100-01) | -1.97565 | - | - | - |
| OS05G0579600 | Homeodomain-like containing protein. (Os05t0579600-01) | -1.97525 | -1.45011 | -1.05656 | - |
| OS07G0148600 | Sulfotransferase family protein. (Os07t0148600-00) | -1.95007 | -1.06982 | -1.16604 | - |
| OS01G0748000 | Similar to Dynamin family protein. (Os01t0748000-00) | -1.93774 | - | - | - |
| OS11G0213600 | Peptidase S10, serine carboxypeptidase family protein. (Os11t0213600-01) | -1.93543 | -0.845983 | -0.691811 | - |
| OS05G0409400 | Similar to Microtubule-associated protein MAP65-1a. (Os05t0409400-01) | -1.90719 | - | 0.784181 | - |
| OS04G0496000 | Glucose/ribitol dehydrogenase family protein. (Os04t0496000-01) | -1.8955 | -1.23853 | - | - |
| OS02G0153900 | Protein kinase, core domain containing protein. (Os02t0153900-01) | -1.88631 | - | - | - |
| OS07G0523100 | Similar to 60S ribosomal protein L44. (Os07t0523100-00) | -1.8785096 | - | - | 1.83967747 |
| OS11G0521000 | Lipase, GDSL domain containing protein. (Os11t0521000-01) | -1.87464 | -1.33881 | -1.18597 | - |
| OS02G0699500 | Conserved hypothetical protein. (Os02t0699500-01) | -1.83232 | -1.34776 | - | - |
| OS06G0713300 | Conserved hypothetical protein. (Os06t0713300-01) | -1.82091 | - | - | - |
| OS11G0128500 | Similar to Myb factor. (Os11t0128500-01) | -1.81974 | -1.1519 | - | 1.06611 |
| OS11G0109600 | Hypothetical conserved gene. (Os11t0109600-00) | -1.81768 | - | - | - |
| OS10G0479500 | Similar to carboxy-lyase. (Os10t0479500-01) | -1.81045 | -1.2438 | -0.901199 | - |
| OS06G0173000 | Armadillo-type fold domain containing protein. (Os06t0173000-00) | -1.80339 | - | - | - |
| OS05G0102000 | SAM dependent carboxyl methyltransferase family protein. (Os05t0102000-01) | -1.80256 | -1.47622 | - | - |
| OS10G0160100 | Glycoside hydrolase, family 17 protein. (Os10t0160100-01);Glycoside hydrolase, family 17 protein. (Os10t0160100-02) | -1.79934 | -1.46864 | -1.02035 | - |

| | | | | | |
|---|---|---|---|---|---|
| OS02G0230250 | Non-protein coding transcript. (Os02t0230250-00) | -1.79663 | -1.02276 | - | - |
| OS04G0644400 | Similar to Proline-rich-like protein. (Os04t0644400-01) | -1.77006 | -1.48651 | -1.3183 | - |
| OS03G0338400 | Basic helix-loop-helix dimerisation region bHLH domain containing protein. (Os03t0338400-01) | -1.7597 | -1.48305 | - | - |
| OS01G0170500 | Protein of unknown function DUF239, plant domain containing protein. (Os01t0170500-01) | -1.7574 | - | -1.31548 | - |
| OS04G0382400 | Similar to OSIGBa0092E09.7 protein. (Os04t0382400-01) | -1.74482 | -1.44178 | - | - |
| OS04G0419600 | Histone H3. (Os04t0419600-01) | -1.74268 | -1.05883 | - | 0.840063 |
| OS05G0443800 | Similar to Plastid division protein ftsZ1 precursor. (Os05t0443800-01) | -1.74119 | -0.978745 | - | - |
| OS07G0406800 | DNA primase, large subunit, eukaryotic domain containing protein. (Os07t0406800-01);Similar to predicted protein. (Os07t | -1.73819 | - | - | - |
| OS01G0854800 | Similar to Cytochrome P450 86A1 (EC 1.14.-.-) (CYPLXXXVI) (P450-dependent fatty acid omega-hydroxylase). (Os01t085480 | -1.73714 | -1.06168 | -0.88315 | - |
| OS01G0709000 | Similar to Transcription factor MYB1. (Os01t0709000-01) | -1.72639 | - | - | - |
| OS05G0245300 | Uncharacterised protein family UPF0497, trans-membrane plant subgroup domain containing protein. (Os05t0245300-01) | -1.72288 | -1.05246 | - | 0.915611 |
| OS01G0728100 | Lipase, GDSL domain containing protein. (Os01t0728100-01);Similar to anther-specific proline-rich protein APG. (Os01t072 | -1.71959 | -1.38416 | -0.833062 | - |
| OS05G0355900 | Conserved hypothetical protein. (Os05t0355900-01) | -1.71933 | - | - | - |
| OS02G0529700 | Similar to Acidic ribosomal protein P2a-4 (Fragment). (Os02t0529700-01) | -1.7118 | -1.21865 | - | - |
| OS02G0833100 | Similar to antiporter/ drug transporter. (Os02t0833100-01) | -1.7023 | - | -0.708567 | - |
| OS11G0135400 | 60S acidic ribosomal protein P0. (Os11t0135400-00) | -1.69422 | - | - | - |
| OS04G0689900 | Similar to Pathogenesis-related protein 5 precursor (PR-5). (Os04t0689900-01) | -1.68149 | - | -1.1308 | - |
| OS10G0109900 | Major facilitator superfamily, general substrate transporter domain containing protein. (Os10t0109900-00) | -1.6751 | -1.21079 | - | - |
| OS04G0554800 | Similar to RCc3 protein. (Os04t0554800-01) | -1.66725 | -1.28477 | -0.922935 | - |
| OS11G0302800 | Protein of unknown function DUF724 family protein. (Os11t0302800-01) | -1.66188 | -1.3874 | -1.12672 | - |
| OS03G0853600 | Conserved hypothetical protein. (Os03t0853600-01) | -1.66082 | - | - | - |
| OS09G0560000 | Armadillo-like helical domain containing protein. (Os09t0560000-01) | -1.64498 | - | - | - |
| OS11G0522900 | Peptidase S10, serine carboxypeptidase family protein. (Os11t0522900-01) | -1.63857 | -1.28623 | - | - |
| OS02G0574600 | Similar to H0523F07.2 protein. (Os02t0574600-01) | -1.63847 | -1.13288 | - | - |
| OS01G0544450 | Similar to DNA replication licensing factor mcm4. (Os01t0544450-01) | -1.62765 | - | - | - |
| OS02G0699466 | Conserved hypothetical protein. (Os02t0699466-00) | -1.62655 | -1.41621 | - | - |
| OS10G0505500 | Similar to Nonspecific lipid-transfer protein 2G (LTP2G) (Lipid transfer protein 2 isoform 1) (LTP2-1) (7 kDa lipid transfer pro | -1.61748 | -1.21993 | - | 0.654005 |
| OS02G0555700 | Similar to Chaperone protein dnaJ 10 (AtJ10) (AtDjC10). (Os02t0555700-01) | -1.61736 | - | 1.22843 | - |
| OS09G0474000 | bZIP transcription factor, bZIP-1 domain containing protein. (Os09t0474000-01) | -1.60766 | - | - | - |
| OS11G0641300 | Conserved hypothetical protein. (Os11t0641300-00) | -1.60329 | - | - | - |
| OS06G0257600 | Esterase, SGNH hydrolase-type domain containing protein. (Os06t0257600-01) | -1.60232 | -1.43628 | -0.922024 | - |
| OS05G0130600 | Conserved hypothetical protein. (Os05t0130600-01) | -1.60224 | -1.05735 | -0.889009 | - |
| OS09G0427125 | Hypothetical conserved gene. (Os09t0427125-00) | -1.5904 | -1.26005 | - | - |
| OS12G0139400 | A-type response regulator, Cytokinin signaling (Os12t0139400-01) | -1.58684 | -1.07952 | -1.27804 | - |
| OS04G0682500 | Peptidase T2, asparaginase 2 domain containing protein. (Os04t0682500-01) | -1.58511 | - | - | - |
| OS06G0217300 | Similar to Transcription factor MADS55. (Os06t0217300-01) | -1.58481 | - | - | - |
| OS07G0119400 | Similar to Pectinesterase like protein. (Os07t0119400-01) | -1.58092 | -1.06827 | - | 1.05003 |
| OS01G0699900 | Similar to Plasma membrane associated protein-like. (Os01t0699900-01) | -1.58021 | -1.49286 | -0.864318 | - |
| OS01G0685900 | Similar to 65kD microtubule associated protein. (Os01t0685900-01);Similar to 65kD microtubule associated protein. (Os01t | -1.5742 | - | 0.714537 | - |
| OS06G0169900 | Similar to GOS9 protein. (Os06t0169900-00) | -1.56916 | - | -1.36997 | - |
| OS10G0104900 | Chromomethylase, Plant development, Silencing of transposable elements (TEs) (Os10t0104900-01) | -1.55945 | - | - | - |
| OS03G0644400 | Amino acid permease. (Os03t0644400-01);Similar to Amino acid permease. (Os03t0644400-02) | -1.55872 | -1.28679 | -1.35534 | - |
| OS03G0850100 | NLI interacting factor domain containing protein. (Os03t0850100-01) | -1.55003 | - | - | - |
| OS10G0159600 | Heat shock protein Hsp20 domain containing protein. (Os10t0159600-01) | -1.54429 | -1.11894 | - | - |
| OS02G0653400 | Similar to OSIGBa0101C23.8 protein. (Os02t0653400-01) | -1.53356 | -1.2546 | - | - |
| OS08G0116800 | Exoribonuclease domain containing protein. (Os08t0116800-01) | -1.53024 | - | - | - |
| OS06G0217700 | Conserved hypothetical protein. (Os06t0217700-01) | -1.52955 | -1.31338 | -1.34029 | - |
| OS03G0249400 | Similar to 40S ribosomal protein S20 (S22) (Fragment). (Os03t0249400-01) | -1.52669 | -1.04121 | -0.910097 | 0.641374 |

| | | | | | |
|---|---|---|---|---|---|
| OS12G0590500 | Similar to Kinesin motor domain containing protein, expressed. (Os12t0590500-00) | -1.52483 | - | - | - |
| OS08G0538700 | Retinoblastoma-related protein. (Os08t0538700-01);Retinoblastoma-related protein. (Os08t0538700-02) | -1.52322 | - | 0.960184 | - |
| OS11G0133900 | Conserved hypothetical protein. (Os11t0133900-00) | -1.51676 | -1.13866 | - | - |
| OS01G0191800 | Similar to Aurora kinase. (Os01t0191800-01);Similar to Aurora kinase. (Os01t0191800-02) | -1.51643 | - | - | - |
| OS03G0171700 | Basic helix-loop-helix dimerisation region bHLH domain containing protein. (Os03t0171700-01) | -1.51097 | - | - | - |
| OS09G0539800 | Similar to Acyl carrier protein III, chloroplast precursor (ACP III). (Os09t0539800-01) | -1.50033 | -1.09601 | - | - |
| OS01G0876400 | Sad1/UNC-like, C-terminal domain containing protein. (Os01t0876400-01);Hypothetical conserved gene. (Os01t0876400-0 | 1.50009 | - | 0.717691 | - |
| OS10G0552400 | Zinc finger, RING/FYVE/PHD-type domain containing protein. (Os10t0552400-01) | 1.50595 | 1.26009 | 1.32809 | -1.48651 |
| OS05G0424300 | Cytochrome P450 family protein. (Os05t0424300-01) | 1.50749 | - | 0.872604 | - |
| OS03G0338200 | Conserved hypothetical protein. (Os03t0338200-01) | 1.50824 | - | - | - |
| OS01G0504100 | Protein of unknown function DUF250 domain containing protein. (Os01t0504100-01) | 1.50858 | 1.16865 | - | - |
| OS05G0541800 | Similar to Ipomoelin. (Os05t0541800-01) | 1.51057 | - | - | - |
| OS03G0122300 | Similar to Flavanone 3-hydroxylase-like protein. (Os03t0122300-01);Similar to Oxidoreductase, 2OG-Fe oxygenase family p | 1.51356 | 1.21539 | 1.17294 | - |
| OS03G0266300 | Class I low-molecular-weight heat shock protein 17.9. (Os03t0266300-01);Class I low-molecular-weight heat shock protein | 1.51676 | 1.30949 | 1.26328 | - |
| OS06G0651300 | Zinc finger, U1-type domain containing protein. (Os06t0651300-01) | 1.51756 | - | - | - |
| OS10G0392400 | Tify domain containing protein. (Os10t0392400-01) | 1.52376 | - | 1.39017 | - |
| OS12G0169600 | Similar to Fructose-bisphosphate aldolase. (Os12t0169600-01) | 1.52598 | - | - | - |
| OS01G0860601 | Similar to Ferredoxin, root R-B1. (Os01t0860601-01) | 1.52954 | 1.15392 | 1.44059 | - |
| OS05G0449600 | Similar to glycosyl hydrolase family 3 protein. (Os05t0449600-01);Hypothetical conserved gene. (Os05t0449600-02) | 1.52962 | 1.21191 | - | -1.11145 |
| OS03G0717900 | Similar to ABRH7 (Fragment). (Os03t0717900-01);Similar to mRNA, clone: RTFL01-13-K15. (Os03t0717900-02) | 1.53782 | 1.42686 | 0.950275 | - |
| OS08G0386800 | Adenine nucleotide translocator 1 domain containing protein. (Os08t0386800-01);Adenine nucleotide translocator 1 doma | 1.5448 | - | - | - |
| OS03G0591700 | Conserved hypothetical protein. (Os03t0591700-01) | 1.54828 | - | - | - |
| OS12G0234000 | Hypothetical protein. (Os12t0234000-01) | 1.55499 | - | - | - |
| OS01G0123700 | Zinc finger, RING-type domain containing protein. (Os01t0123700-00) | 1.55652261 | - | - | - |
| OS06G0324400 | Late embryogenesis abundant (LEA) group 1 family protein. (Os06t0324400-01) | 1.56251353 | -0.1317099 | 0.08177592 | - |
| OS05G0163700 | Similar to Acyl-coenzyme A oxidase 4, peroxisomal (EC 1.3.3.6) (AOX 4) (Short- chain acyl-CoA oxidase) (SAOX) (AtCX4) (G6p | 1.56635 | 1.29343 | 1.30951 | - |
| OS11G0515700 | Non-protein coding transcript. (Os11t0515700-00) | 1.57143 | - | 1.33312 | - |
| OS12G0112000 | Similar to Peroxidase precursor (EC 1.11.1.7) (Fragment). (Os12t0112000-01) | 1.58175 | 1.1331 | 0.967162 | - |
| OS01G0837800 | Metal tolerance protein, Manganese transporter, Mn-cation diffusion facilitator (CDF ), Mn and other heavy metal toleranc | 1.58852 | 1.18872 | - | -0.860016 |
| OS08G0202400 | Disease resistance protein domain containing protein. (Os08t0202400-01) | 1.59313 | - | - | - |
| OS05G0508400 | Mannose-binding lectin domain containing protein. (Os05t0508400-01) | 1.59586 | 1.11761 | 0.718813 | - |
| OS03G0794200 | Similar to N-acetyltransferase. (Os03t0794200-00) | 1.59602 | - | - | - |
| OS05G0522600 | Leucine-rich repeat, plant specific containing protein. (Os05t0522600-01);Leucine-rich repeat domain containing protein. ( | 1.6004 | - | - | -1.38768 |
| OS09G0376300 | Conserved hypothetical protein. (Os09t0376300-01) | 1.60414 | - | - | - |
| OS06G0586000 | Conserved hypothetical protein. (Os06t0586000-01) | 1.61516 | 1.30848 | 0.78487 | -2.20091 |
| OS07G0241800 | UDP-glucuronosyl/UDP-glucosyltransferase family protein. (Os07t0241800-01) | 1.6176 | 1.34336 | - | - |
| OS01G0799100 | NB-ARC domain containing protein. (Os01t0799100-01) | 1.62697 | - | - | - |
| OS02G0565600 | Similar to Homeodomain leucine zipper protein (Fragment). (Os02t0565600-01) | 1.62856 | - | - | - |
| OS10G0528100 | Similar to Glutathione S-transferase GST 42 (EC 2.5.1.18) (Fragment). (Os10t0528100-01) | 1.62938 | - | 0.855519 | - |
| OS01G0601625 | Leucine rich repeat, N-terminal domain containing protein. (Os01t0601625-00) | 1.64138 | - | - | -2.09333 |
| OS04G0128900 | Flavin monooxygenase-like enzyme, Auxin biosynthesis (Os04t0128900-01) | 1.64288 | - | 1.47971 | - |
| OS07G0520900 | Similar to galactolipase/ phospholipase. (Os07t0520900-01);Similar to predicted protein. (Os07t0520900-02) | 1.64717 | - | 0.964909 | - |
| OS05G0584600 | Similar to ATP binding / nucleoside-triphosphatase/ nucleotide binding. (Os05t0584600-01);ATPase, AAA-type, core domai | 1.65808 | - | -1.01655 | - |
| OS05G0132400 | Agenet domain containing protein. (Os05t0132400-01) | 1.66484 | - | - | - |
| OS01G0788400 | Similar to Pectinesterase (EC 3.1.1.11) (Fragment). (Os01t0788400-01) | 1.68174 | - | - | - |
| OS01G0647850 | Hypothetical conserved gene. (Os01t0647850-01) | 1.68516 | - | - | - |
| OS10G0564800 | Calcineurin B-like protein, Calcium sensor protein, Reguration of potassium uptake by CBL1-CIPK23 complex (Os10t056480 | 1.69449 | - | - | - |
| OS03G0145500 | DENN domain containing protein. (Os03t0145500-01) | 1.70512 | 1.13415 | - | -1.09502 |

| | | | | | |
|---|---|---|---|---|---|
| OS10G0477200 | Pentatricopeptide repeat domain containing protein. (Os10t0477200-01) | 1.70525 | - | 1.40493 | - |
| OS11G0181800 | Similar to Short-chain dehydrogenase Tic32. (Os11t0181800-01) | 1.7068 | - | - | - |
| OS07G0273900 | Disease resistance protein domain containing protein. (Os07t0273900-01) | 1.70851 | - | - | -1.79023 |
| OS03G0684400 | Mg2+ transporter protein, CorA-like domain containing protein. (Os03t0684400-01) | 1.71053 | 1.37436 | - | - |
| OS02G0756800 | Phosphate-induced protein 1 conserved region family protein. (Os02t0756800-01) | 1.7109 | 1.44096 | 0.917145 | - |
| OS01G0160800 | Similar to Protein synthesis inhibitor II (EC 3.2.2.22) (Ribosome-inactivating protein II) (rRNA N-glycosidase). (Os01t016080 | 1.71216 | 1.46247 | 1.46519 | - |
| OS10G0527800 | Similar to Tau class GST protein 3. (Os10t0527800-01) | 1.71367 | - | - | - |
| OS08G0110400 | Protein of unknown function DUF266, plant family protein. (Os08t0110400-01) | 1.71812 | - | - | - |
| OS01G0846500 | Hypothetical conserved gene. (Os01t0846500-01);PAP/25A core domain containing protein. (Os01t0846500-02) | 1.72698 | - | - | - |
| OS05G0460000 | Similar to 70 kDa heat shock cognate protein 1. (Os05t0460000-01) | 1.73671 | - | 1.31267 | - |
| OS06G0326700 | Diacylglycerol acyltransferase family protein. (Os06t0326700-01) | 1.7475 | - | - | - |
| OS02G0129700 | Hypothetical protein. (Os02t0129700-01) | 1.7519 | - | - | -1.23024 |
| OS10G0517500 | Cys/Met metabolism, pyridoxal phosphate-dependent enzyme domain containing protein. (Os10t0517500-02) | 1.75452 | - | - | - |
| OS10G0542900 | Similar to chitinase. (Os10t0542900-01) | 1.76404 | 0.928762 | 1.29672 | -2.76851 |
| OS01G0757200 | GA 2-oxidase3, GA metabolism (Os01t0757200-01) | 1.76972 | 1.24826 | 0.799602 | - |
| OS01G0850100 | Similar to Phosphatidic acid phosphatase-like protein. (Os01t0850100-01) | 1.779 | - | - | - |
| OS05G0421600 | No apical meristem (NAM) protein domain containing protein. (Os05t0421600-01) | 1.78261 | 1.33829 | - | - |
| OS01G0817800 | WD40 repeat-like domain containing protein. (Os01t0817800-01);Similar to predicted protein. (Os01t0817800-02) | 1.78714 | - | 1.15067 | - |
| OS01G0695200 | Protein of unknown function DUF266, plant family protein. (Os01t0695200-01) | 1.80244 | 1.34885 | -0.780444 | -1.73098 |
| OS09G0426000 | Protein of unknown function DUF6, transmembrane domain containing protein. (Os09t0426000-01) | 1.80712 | - | 1.31952 | - |
| OS10G0205802 | Conserved hypothetical protein. (Os10t0205802-01) | 1.8171 | 1.32674 | - | - |
| OS04G0461700 | Non-protein coding transcript. (Os04t0461700-01);Non-protein coding transcript. (Os04t0461700-02) | 1.81874 | - | 1.10479 | - |
| OS10G0501500 | Protein of unknown function DUF607 family protein. (Os10t0501500-01) | 1.83933 | - | - | - |
| OS07G0591800 | Conserved hypothetical protein. (Os07t0591800-01) | 1.84183 | - | - | -1.36388 |
| OS07G0686300 | Zinc finger, RING/FYVE/PHD-type domain containing protein. (Os07t0686300-01) | 1.8467 | 1.39767 | - | - |
| OS01G0226500 | Conserved hypothetical protein. (Os01t0226500-01);Similar to predicted protein. (Os01t0226500-02) | 1.85126 | - | - | -1.77802 |
| OS06G0295500 | Hypothetical gene. (Os06t0295500-01);Hypothetical protein. (Os06t0295500-02);Non-protein coding transcript. (Os06t029 | 1.85385 | - | - | - |
| OS08G0407600 | Protein of unknown function DUF581 family protein. (Os08t0407600-01) | 1.85856 | 1.17693 | - | - |
| OS08G0198900 | Conserved hypothetical protein. (Os08t0198900-01) | 1.86525 | - | - | -2.0602 |
| OS07G0442900 | Uncharacterised protein family UPF0497, trans-membrane plant subgroup domain containing protein. (Os07t0442900-01) | 1.87955 | - | - | - |
| OS11G0537300 | Similar to DedA. (Os11t0537300-01) | 1.92663 | - | - | -1.65774 |
| OS12G0442900 | DNA-binding SAP domain containing protein. (Os12t0442900-01) | 1.93669 | - | - | - |
| OS02G0181300 | WRKY transcription factor, Defense response (Os02t0181300-01) | 1.94026 | - | - | - |
| OS09G0513100 | Similar to Phospholipase A1. (Os09t0513100-01);Similar to phospholipase A1. (Os09t0513100-02) | 1.94295 | - | - | - |
| OS09G0452900 | Glycosyl transferase, family 31 protein. (Os09t0452900-01) | 1.94807 | 1.4245 | 1.2328 | - |
| OS05G0515200 | Cytochrome P450 family protein. (Os05t0515200-01) | 1.95256 | 1.45008 | - | -1.21756 |
| OS10G0491000 | Plant Basic Secretory Protein family protein. (Os10t0491000-01) | 1.97129 | 1.35118 | 1.15443 | -1.11731 |
| OS06G0674800 | Hypothetical conserved gene. (Os06t0674800-01) | 1.97759 | - | - | - |
| OS07G0489000 | Plant lipid transfer protein/Par allergen family protein. (Os07t0489000-01) | 1.97864228 | - | - | -3.2403052 |
| OS03G0371000 | Similar to Cytochrome P450 family protein, expressed. (Os03t0371000-01) | 1.98437 | - | - | - |
| OS08G0385000 | Similar to seed maturation protein PM41. (Os08t0385000-01) | 1.98913537 | - | 0.69213256 | - |
| OS08G0184600 | Conserved hypothetical protein. (Os08t0184600-01);Conserved hypothetical protein. (Os08t0184600-02);Conserved hypot | 2.01216 | - | - | - |
| OS10G0393800 | Lipase, GDSL domain containing protein. (Os10t0393800-01) | 2.02058 | - | - | - |
| OS01G0883800 | Similar to GA C20oxidase2. (Os01t0883800-01);GA 20-oxidase2, GA metabolism (Os01t0883800-02) | 2.02637 | 1.29828 | - | - |
| OS04G0515400 | NTP pyrophosphohydrolase MazG, putative catalytic core domain containing protein. (Os04t0515400-00) | 2.02721 | - | - | - |
| OS05G0461600 | Similar to SIK1 protein (Nucleolar protein NOP56). (Os05t0461600-01) | 2.04094 | - | - | - |
| OS02G0484200 | Transferase family protein. (Os02t0484200-01) | 2.04721 | - | - | - |
| OS03G0773300 | Protein kinase, core domain containing protein. (Os03t0773300-01);Similar to wound and phytochrome signaling involved | 2.0919 | - | - | -1.95667 |

| | | | | | |
|---|---|---|---|---|---|
| OS04G0409900 | Plant neutral invertase family protein. (Os04t0409900-01) | 2.0975 | - | - | - |
| OS03G0654700 | Protein of unknown function DUF1637 family protein. (Os03t0654700-01) | 2.10097428 | - | - | - |
| OS01G0805400 | UDP-glucuronosyl/UDP-glucosyltransferase family protein. (Os01t0805400-01) | 2.1165 | - | - | - |
| OS10G0535800 | Protein of unknown function Cys-rich domain containing protein. (Os10t0535800-01) | 2.11766 | 1.28237 | - | -2.17533 |
| OS04G0301700 | Glycosyl transferase, family 14 protein. (Os04t0301700-01) | 2.17646 | - | 1.20936 | - |
| OS07G0682000 | Heavy metal transport/detoxification protein domain containing protein. (Os07t0682000-01) | 2.18107 | - | - | - |
| OS12G0281600 | NB-ARC domain containing protein. (Os12t0281600-01);Hypothetical conserved gene. (Os12t0281600-02) | 2.19775 | - | - | - |
| OS09G0255400 | Similar to Indole-3-glycerol phosphate synthase, chloroplast precursor (EC 4.1.1.48) (IGPS). (Os09t0255400-01);Similar to ir | 2.22695 | - | - | -3.14574 |
| OS02G0216600 | Similar to Protein ROOT HAIR DEFECTIVE 3. (Os02t0216600-01) | 2.227 | - | - | - |
| OS06G0323100 | Similar to H1005F08.18 protein. (Os06t0323100-01) | 2.2705 | - | 0.889409 | -2.75428 |
| OS02G0770100 | WD40 repeat domain containing protein. (Os02t0770100-01) | 2.27666 | - | - | -2.21351 |
| OS07G0663900 | Similar to Oxidoreductase, short chain dehydrogenase/reductase family protein, expressed. (Os07t0663900-00) | 2.29101115 | - | - | -1.1481076 |
| OS09G0491740 | Auxin efflux carrier domain containing protein. (Os09t0491740-01) | 2.29867 | - | - | -1.79841 |
| OS09G0135100 | Hypothetical conserved gene. (Os09t0135100-00) | 2.30006 | - | - | - |
| OS05G0324700 | Conserved hypothetical protein. (Os05t0324700-01) | 2.32117 | - | - | - |
| OS07G0670300 | C2 calcium-dependent membrane targeting domain containing protein. (Os07t0670300-01) | 2.37234 | 1.10549 | - | - |
| OS12G0555200 | Similar to Probenazole-inducible protein PBZ1. (Os12t0555200-01) | 2.38069 | - | - | -4.50082 |
| OS08G0310001 | Non-protein coding transcript. (Os08t0310001-01) | 2.3998 | - | - | - |
| OS12G0595800 | Similar to cDNA clone:J023098M23, full insert sequence. (Os12t0595800-01);Similar to Protein kinase domain containing p | 2.41341 | - | - | - |
| OS06G0697600 | ATPase, AAA-type, core domain containing protein. (Os06t0697600-01) | 2.42751 | - | - | - |
| OS03G0800000 | Similar to Nitrate and chloride transporter. (Os03t0800000-01);Similar to nitrate and chloride transporter. (Os03t0800000- | 2.42803 | 1.05637 | - | -1.26106 |
| OS12G0555500 | Probenazole-inducible protein PBZ1. (Os12t0555500-01) | 2.44189 | 1.16947 | - | -5.03856 |
| OS02G0609500 | Hypothetical protein. (Os02t0609500-01) | 2.50223 | - | - | - |
| OS10G0389200 | Red chlorophyll catabolite reductase, Leaf senescence, Wound responses (Os10t0389200-01) | 2.51691 | - | 1.27891 | - |
| OS07G0521100 | Leucine-rich repeat, typical subtype containing protein. (Os07t0521100-01) | 2.52619 | - | - | -3.14274 |
| OS07G0271000 | Similar to GDP dissociation inhibitor protein OsGDI1. (Os07t0271000-01) | 2.62346 | - | - | - |
| OS09G0498000 | Hypothetical gene. (Os09t0498000-01) | 2.63361 | 0.903562 | - | -2.28434 |
| OS01G0959200 | Similar to Ci21A protein. (Os01t0959200-01) | 2.64401 | - | - | -2.50483 |
| OS03G0790500 | Alpha/beta hydrolase fold-3 domain containing protein. (Os03t0790500-01) | 2.64876 | - | - | - |
| OS05G0182600 | Similar to SSRP1 protein. (Os05t0182600-01) | 2.70023 | - | - | -2.97544 |
| OS04G0629300 | Similar to H0303G06.18 protein. (Os04t0629300-01);DEAD-like helicase, N-terminal domain containing protein. (Os04t062[illegible] | 2.79876 | 1.29313 | 0.837086 | - |
| OS11G0641500 | *Cupredoxin domain containing protein. (Os11t0641500-01)* | 2.83335 | - | -1.21786 | -4.38129 |
| OS06G0110200 | Late embryogenesis abundant (LEA) group 1 family protein. (Os06t0110200-01) | 2.88059 | - | - | - |
| OS02G0759400 | Zinc finger, RING/FYVE/PHD-type domain containing protein. (Os02t0759400-01) | 2.9076 | - | - | - |
| OS07G0656400 | Protein of unknown function DUF288 domain containing protein. (Os07t0656400-01) | 2.9266 | - | -1.11079 | -2.58221 |
| OS03G0834466 | Similar to MADS-box transcription factor 21. (Os03t0834466-00) | 3.16078307 | - | - | - |
| OS02G0676800 | Similar to Dehydration responsive element binding protein 1E (DREB1E protein). (Os02t0676800-01) | 3.17059 | - | - | - |
| OS03G0795400 | Hypothetical conserved gene. (Os03t0795400-00) | 3.18329 | - | - | -2.38033 |
| OS04G0538400 | Similar to Nodulin 21 (N-21). (Os04t0538400-01) | 3.20447 | - | - | - |
| OS03G0432100 | Similar to Pyruvate, phosphate dikinase 2. (Os03t0432100-01) | 3.44585 | - | - | - |
| OS11G0514500 | Sorghum bicolor leucine-rich repeat-containing extracellular glycoprotein precursor. (Os11t0514500-01) | 3.45449 | 1.09952 | - | -3.2422 |
| OS11G0619000 | Hypothetical protein. (Os11t0619000-01) | 3.51362 | - | - | - |
| OS03G0661600 | Similar to Alpha-amylase/trypsin inhibitor (Antifungal protein). (Os03t0661600-01) | 3.67793 | - | 0.89187 | - |
| OS10G0409400 | beta subunit of polygalacturonase 1, Abiotic stress response (Os10t0409400-01) | 3.76247 | - | - | - |
| OS07G0154100 | Similar to Viviparous-14. (Os07t0154100-01) | 4.56582 | - | - | - |
| OS11G0181200 | Hypothetical protein. (Os11t0181200-01) | 5.58544632 | - | - | - |

| 5. Stress-responsive in WT | | | | |
|---|---|---|---|---|
| Identifier | Description | FC.NmNs | FC.NmSm | FC.NmSSm |
| OS02G0787400 | Non-protein coding transcript. (Os02t0787400-00) | -5.3071553 | - | - |
| OS10G0508700 | Pectinesterase inhibitor domain containing protein. (Os10t0508700-01) | -4.85304 | -1.17874 | -0.940467 |
| OS12G0199800 | Similar to Cyt-P450 monooxygenase. (Os12t0199800-01) | -4.1946 | - | - |
| OS03G0431600 | Hypothetical conserved gene. (Os03t0431600-00) | -4.14665 | - | - |
| OS02G0162200 | Similar to Early salt stress and cold acclimation-induced protein 2-3. (Os02t0162200-00) | -3.9151 | - | - |
| OS04G0415600 | Similar to H0622F05.9 protein. (Os04t0415600-00) | -3.80573 | - | - |
| OS03G0773000 | Protein of unknown function DUF1005 family protein. (Os03t0773000-01) | -3.63242 | -1.72485 | -1.35082 |
| OS03G0323900 | Bifunctional inhibitor/plant lipid transfer protein/seed storage domain containing protein. (Os03t0323900-01) | -3.5962 | - | - |
| OS01G0608101 | Conserved hypothetical protein. (Os01t0608101-00) | -3.51229 | - | - |
| OS03G0782400 | Conserved hypothetical protein. (Os03t0782400-00) | -3.45909 | - | - |
| OS03G0397600 | Glycoside hydrolase, family 17 protein. (Os03t0397600-00) | -3.44869 | - | - |
| OS01G0873200 | Similar to Amidophosphoribosyltransferase, chloroplast precursor (EC 2.4.2.14) (Glutamine phosphoribosylpyrophosphate amidotra | -3.42079 | - | - |
| OS04G0495300 | Similar to OSIGBa0159F11.7 protein. (Os04t0495300-01) | -3.41145 | - | - |
| OS05G0567600 | Similar to SANT/MYB protein. (Os05t0567600-00) | -3.39765 | - | - |
| OS01G0313300 | Similar to EREBP-3 protein (Fragment). (Os01t0313300-01) | -3.3893 | - | - |
| OS05G0541400 | Helix-loop-helix DNA-binding domain containing protein. (Os05t0541400-00) | -3.34181 | - | - |
| OS07G0241050 | Hypothetical protein. (Os07t0241050-00) | -3.32448 | - | - |
| OS05G0568300 | Similar to 50S ribosomal protein L12, chloroplast precursor (CL12). (Os05t0568300-00) | -3.31485 | - | - |
| OS08G0549600 | bZIP transcription factor, bZIP-1 domain containing protein. (Os08t0549600-00) | -3.30132 | -1.3697 | #N/A |
| OS03G0860600 | Similar to 2-oxoglutarate-dependent oxygenase. (Os03t0860600-01) | -3.27809 | -1.3868 | #N/A |
| OS01G0237500 | NmrA-like family protein. (Os01t0237500-01) | -3.25881 | #N/A | -4.3038319 |
| OS03G0233000 | Protein of unknown function DUF607 family protein. (Os03t0233000-01) | -3.23715 | - | - |
| OS05G0588250 | Hypothetical protein. (Os05t0588250-00) | -3.20202 | - | - |
| OS03G0431100 | Conserved hypothetical protein. (Os03t0431100-01) | -3.18043 | -0.842096 | #N/A |
| OS09G0119600 | UDP-glucuronosyl/UDP-glucosyltransferase family protein. (Os09t0119600-01) | -3.14604 | - | - |
| OS02G0265200 | WRKY transcription factor 39. (Os02t0265200-01);Similar to WRKY39v2 - superfamily of TFs having WRKY and zinc finger domains. ( | -3.14508 | - | - |
| OS09G0273600 | Hypothetical gene. (Os09t0273600-00) | -3.13046 | - | - |
| OS07G0162700 | Alpha/beta hydrolase fold-3 domain containing protein. (Os07t0162700-01) | -3.1156 | - | - |
| OS05G0215300 | UDP-glucuronosyl/UDP-glucosyltransferase family protein. (Os05t0215300-01) | -3.11204 | -0.936757 | #N/A |
| OS08G0390600 | Conserved hypothetical protein. (Os08t0390600-01) | -3.10735 | - | - |
| OS01G0916100 | Similar to loricrin. (Os01t0916100-01) | -3.09034 | - | - |
| OS02G0308800 | FAS1 domain domain containing protein. (Os02t0308800-01);Conserved hypothetical protein. (Os02t0308800-02) | -3.07992 | -1.23411 | #N/A |
| OS01G0210300 | Conserved hypothetical protein. (Os01t0210300-01) | -3.06824 | - | - |
| OS09G0498500 | NAD-binding site containing protein. (Os09t0498500-00) | -3.06426 | - | - |
| OS01G0785900 | Zinc finger, C2H2-type domain containing protein. (Os01t0785900-01);Zinc finger, C2H2-type domain containing protein. (Os01t078 | -3.03065 | - | - |
| OS07G0105000 | Cupredoxin domain containing protein. (Os07t0105000-00) | -2.96666 | - | - |
| OS12G0455100 | Hypothetical protein. (Os12t0455100-01) | -2.95707 | - | - |
| OS03G0128000 | FAS1 domain domain containing protein. (Os03t0128000-01);Similar to Fasciclin-like protein FLA2. (Os03t0128000-02) | -2.9509 | - | - |
| OS07G0673600 | Conserved hypothetical protein. (Os07t0673600-00) | -2.94948 | - | - |

| | | | | |
|---|---|---|---|---|
| OS12G0578600 | Zinc finger, C2H2-type domain containing protein. (Os12t0578600-01) | -2.93376 | - | - |
| OS06G0253600 | Similar to secretory protein-like. (Os06t0253600-00) | -2.91732 | - | - |
| OS01G0198500 | Conserved hypothetical protein. (Os01t0198500-01) | -2.9069 | - | - |
| OS01G0296700 | Glycoside hydrolase, family 3, N-terminal domain containing protein. (Os01t0296700-01) | -2.88258 | - | - |
| OS03G0399800 | Mannose-binding lectin domain containing protein. (Os03t0399800-01) | -2.85394 | - | - |
| OS03G0675300 | Hypothetical conserved gene. (Os03t0675300-01) | -2.85271 | - | - |
| OS05G0565701 | Conserved hypothetical protein. (Os05t0565701-00) | -2.8514 | - | - |
| OS03G0391400 | Similar to Phospholipase D nu-2 (Fragment). (Os03t0391400-01) | -2.83972 | - | - |
| OS09G0482660 | Similar to Subtilisin-type protease. (Os09t0482660-01) | -2.82494 | - | - |
| OS10G0546900 | Similar to Zinc finger, C3HC4 type family protein, expressed. (Os10t0546900-00) | -2.81559 | - | - |
| OS03G0196600 | Similar to satase isoform II. (Os03t0196600-01) | -2.79784 | - | - |
| OS02G0489500 | Conserved hypothetical protein. (Os02t0489500-01) | -2.79436 | -1.66737 | #N/A |
| OS01G0963400 | Thioredoxin family protein. (Os01t0963400-01) | -2.788 | - | - |
| OS02G0269600 | Similar to Subtilase. (Os02t0269600-00) | -2.78471 | -3.09277 | #N/A |
| OS04G0568500 | Conserved hypothetical protein. (Os04t0568500-00) | -2.77955 | - | - |
| OS10G0536700 | Similar to Peroxidase 1. (Os10t0536700-01) | -2.75688 | - | - |
| OS06G0594400 | Kelch-type beta propeller domain containing protein. (Os06t0594400-02) | -2.74482 | - | - |
| OS09G0517900 | UDP-glucuronosyl/UDP-glucosyltransferase family protein. (Os09t0517900-01) | -2.74366 | - | - |
| OS07G0673000 | Ureohydrolase domain containing protein. (Os07t0673000-01) | -2.74183 | -1.34045 | -1.09579 |
| OS08G0392100 | 2OG-Fe(II) oxygenase domain containing protein. (Os08t0392100-00) | -2.74077 | - | - |
| OS09G0371625 | Hypothetical protein. (Os09t0371625-00) | -2.73491 | - | - |
| OS09G0306650 | Hypothetical conserved gene. (Os09t0306650-01) | -2.70582 | -1.18129 | #N/A |
| OS06G0711900 | Bifunctional inhibitor/plant lipid transfer protein/seed storage domain containing protein. (Os06t0711900-01) | -2.70472 | #N/A | -0.856185 |
| OS10G0501700 | Similar to Protein Rf1, mitochondrial. (Os10t0501700-00) | -2.697 | 0.5222025 | 0.53212791 |
| OS07G0520600 | Conserved hypothetical protein. (Os07t0520600-01);Conserved hypothetical protein. (Os07t0520600-02) | -2.69493 | - | - |
| OS06G0155400 | Conserved hypothetical protein. (Os06t0155400-00) | -2.69189 | - | - |
| OS08G0556300 | Glycolipid transfer protein domain domain containing protein. (Os08t0556300-01) | -2.67961 | - | - |
| OS01G0276900 | Conserved hypothetical protein. (Os01t0276900-01) | -2.67341 | -1.01071 | #N/A |
| OS06G0157900 | Similar to Non-specific lipid-transfer protein. (Os06t0157900-00) | -2.65341 | - | - |
| OS07G0543100 | Similar to Beta-amylase (EC 3.2.1.2). (Os07t0543100-00) | -2.64647 | -1.75405 | #N/A |
| OS03G0843400 | Similar to 30S ribosomal protein S6, chloroplast precursor (Fragment). (Os03t0843400-01) | -2.64435 | -0.655478 | #N/A |
| OS03G0377100 | Similar to Expansin (Expansin2). (Os03t0377100-01) | -2.63301 | - | - |
| OS04G0423100 | Monooxygenase, FAD-binding domain containing protein. (Os04t0423100-00) | -2.62957 | - | - |
| OS08G0490600 | FAS1 domain domain containing protein. (Os08t0490600-01) | -2.62516 | - | - |
| OS12G0204500 | Protein of unknown function DUF579, plant family protein. (Os12t0204500-01) | -2.62079 | -0.757155 | #N/A |
| OS09G0517200 | Hypothetical conserved gene. (Os09t0517200-00) | -2.61636 | #N/A | 0.29728218 |
| OS03G0738900 | NPH3 domain containing protein. (Os03t0738900-00) | -2.60071 | - | - |
| OS03G0159600 | Similar to Rab28 protein. (Os03t0159600-01) | -2.59251 | - | - |
| OS06G0288300 | C-glucosyltransferase, Flavone-C-glycoside synthesis (Os06t0288300-01) | -2.5829 | - | - |
| OS03G0729700 | Ribosomal RNA small subunit methyltransferase E domain containing protein. (Os03t0729700-01) | -2.58005 | - | - |
| OS03G0182800 | Similar to Ethylene responsive element binding factor3 (OsERF3). (Os03t0182800-01) | -2.5712 | - | - |
| OS12G0541000 | Lumazine-binding protein family protein. (Os12t0541000-01) | -2.56971 | -0.948518 | #N/A |

| | | | | |
|---|---|---|---|---|
| OS04G0674100 | Tetratricopeptide-like helical domain containing protein. (Os04t0674100-01);Similar to H0403D02.12 protein. (Os04t0674100-02) | -2.54681 | - | - |
| OS08G0524400 | Protein of unknown function DUF568, DOMON-like domain containing protein. (Os08t0524400-00) | -2.54501 | -0.984106 | #N/A |
| OS01G0849100 | Small GTPase Rac/ROP guanine nucleotide exchange factor, Signal transduction (Os01t0849100-01) | -2.53774 | - | - |
| OS03G0297600 | Streptomyces cyclase/dehydrase family protein. (Os03t0297600-01) | -2.5281 | - | - |
| OS08G0530300 | Exo70 exocyst complex subunit family protein. (Os08t0530300-01) | -2.52394 | -1.44009 | -1.00062 |
| OS09G0274100 | Similar to H0402C08.3 protein. (Os09t0274100-01) | -2.52097 | - | - |
| OS06G0236300 | Conserved hypothetical protein. (Os06t0236300-01) | -2.5172 | -1.42027 | #N/A |
| OS04G0495700 | Similar to OSIGBa0159F11.10 protein. (Os04t0495700-01) | -2.51365 | - | - |
| OS01G0342500 | Hypothetical conserved gene. (Os01t0342500-00) | -2.51231 | -1.37969 | #N/A |
| OS03G0758900 | Similar to predicted protein. (Os03t0758900-01) | -2.51181 | -0.813904 | -0.758007 |
| OS04G0523500 | Hypothetical conserved gene. (Os04t0523500-00) | -2.51155 | - | - |
| OS04G0568600 | Similar to 6-phospho-3-hexuloisomerase. (Os04t0568600-00) | -2.50922 | - | - |
| OS01G0794800 | Similar to Subtilase. (Os01t0794800-01);Similar to Subtilase. (Os01t0794800-02) | -2.50751 | -2.8148 | -2.74152 |
| OS05G0179950 | Similar to cDNA, clone: J100088D24, full insert sequence. (Os05t0179950-00) | -2.50489 | - | - |
| OS01G0237750 | Hypothetical conserved gene. (Os01t0237750-00) | -2.50305 | - | - |
| OS05G0337400 | Similar to heavy-metal-associated domain-containing protein. (Os05t0337400-01) | -2.49378 | - | - |
| OS07G0243150 | Hypothetical conserved gene. (Os07t0243150-01) | -2.48108 | -0.773889 | #N/A |
| OS01G0284700 | Similar to Peptidyl-prolyl cis-trans isomerase. (Os01t0284700-01) | -2.46758 | -0.836929 | -0.668992 |
| OS04G0662600 | Similar to Naringenin,2-oxoglutarate 3-dioxygenase. (Os04t0662600-00) | -2.46501 | - | - |
| OS04G0208900 | Hypothetical conserved gene. (Os04t0208900-00) | -2.46023 | - | - |
| OS08G0102000 | Serotonin N-acetyltransferase, Melatonin synthesis (Os08t0102000-01) | -2.45597 | -1.22989 | #N/A |
| OS08G0117100 | Nucleotide-binding, alpha-beta plait domain containing protein. (Os08t0117100-01) | -2.4531 | - | - |
| OS06G0214300 | Alpha/beta hydrolase fold-3 domain containing protein. (Os06t0214300-01) | -2.44601 | - | - |
| OS03G0761000 | SWIB/MDM2 domain containing protein. (Os03t0761000-01) | -2.43717 | - | - |
| OS03G0719000 | MAP65/ASE1 family protein. (Os03t0719000-01) | -2.41874 | - | - |
| OS09G0371650 | Non-protein coding transcript. (Os09t0371650-00) | -2.4183 | - | - |
| OS03G0826200 | Conserved hypothetical protein. (Os03t0826200-01) | -2.41828 | - | - |
| OS05G0419000 | Alpha/beta hydrolase family protein. (Os05t0419000-01) | -2.4173 | -0.775528 | #N/A |
| OS09G0438100 | Conserved hypothetical protein. (Os09t0438100-01) | -2.4026 | - | - |
| OS03G0205800 | Similar to Acetyltransferase, GNAT family protein, expressed. (Os03t0205800-00) | -2.40012 | - | - |
| OS01G0201100 | Similar to xylosyltransferase oxt. (Os01t0201100-01) | -2.39773 | - | - |
| OS02G0773500 | Protein of unknown function DUF3464 domain containing protein. (Os02t0773500-01) | -2.38236 | -1.31551 | #N/A |
| OS03G0581400 | Similar to anther-specific proline-rich protein APG. (Os03t0581400-00) | -2.37785 | - | - |
| OS02G0302200 | Similar to aminotransferase family protein. (Os02t0302200-01) | -2.36557 | -1.0837 | -0.921307 |
| OS02G0809800 | Phosphate (Pi) transporter, Root-to-shoot Pi transfer (Os02t0809800-01) | -2.36518 | - | - |
| OS08G0562600 | C2 calcium-dependent membrane targeting domain containing protein. (Os08t0562600-03) | -2.36447 | -1.36723 | -0.994625 |
| OS12G0209000 | Similar to sporulation protein-related. (Os12t0209000-00) | -2.3614 | -1.12787 | -1.01175 |
| OS12G0600701 | Conserved hypothetical protein. (Os12t0600701-00) | -2.34617 | -0.780583 | -0.728428 |
| OS07G0144500 | Similar to Patatin-like phospholipase family protein, expressed. (Os07t0144500-00) | -2.34431 | - | - |
| OS07G0240600 | UDP-glucuronosyl/UDP-glucosyltransferase family protein. (Os07t0240600-01) | -2.3379 | - | - |
| OS03G0229500 | Similar to MATE efflux family protein, expressed. (Os03t0229500-00) | -2.32931 | -1.08579 | #N/A |
| OS10G0181600 | Methyltransferase type 11 domain containing protein. (Os10t0181600-01);Similar to CMV 1a interacting protein 1. (Os10t0181600- | -2.32453 | -0.768382 | #N/A |

| | | | | |
|---|---|---|---|---|
| OS05G0582000 | Plant Basic Secretory Protein domain containing protein. (Os05t0582000-01) | -2.31726 | - | - |
| OS04G0447600 | Similar to NADPH-dependent codeinone reductase (EC 1.1.1.247). (Os04t0447600-01) | -2.31581 | - | - |
| OS08G0235800 | Similar to WRKY transcription factor 25. (Os08t0235800-00) | -2.30863 | - | - |
| OS02G0241200 | Myb-like DNA-binding domain, SHAQKYF class domain containing protein. (Os02t0241200-00) | -2.30681 | - | - |
| OS01G0678600 | Ribosomal protein S20 family protein. (Os01t0678600-01);Similar to ribosomal protein rpS20. (Os01t0678600-02);Similar to riboson | -2.30659 | -0.633511 | #N/A |
| OS07G0631200 | Zinc finger, RING/FYVE/PHD-type domain containing protein. (Os07t0631200-01) | -2.30361 | -1.64056 | #N/A |
| OS02G0594700 | Similar to Non-phototropic hypocotyl 3. (Os02t0594700-01) | -2.2861 | - | - |
| OS11G0463700 | Penicillin-binding protein, transpeptidase fold domain containing protein. (Os11t0463700-01) | -2.28479 | - | - |
| OS02G0569000 | Cytochrome P450 family protein. (Os02t0569000-01) | -2.28402 | #N/A | -2.49274 |
| OS07G0589000 | Lateral organ boundaries, LOB domain containing protein. (Os07t0589000-01) | -2.28226 | -0.904384 | #N/A |
| OS08G0401800 | Similar to Phospholipase D. (Os08t0401800-00) | -2.27541 | - | - |
| OS06G0712200 | NUDIX domain containing protein. (Os06t0712200-01) | -2.2712 | - | - |
| OS03G0239000 | Concanavalin A-like lectin/glucanase, subgroup domain containing protein. (Os03t0239000-01) | -2.27106 | - | - |
| OS01G0732300 | Protein of unknown function DUF623, plant domain containing protein. (Os01t0732300-01) | -2.27087 | - | - |
| OS08G0545700 | Similar to TraB protein-related. (Os08t0545700-01) | -2.25418 | -0.924907 | -0.679067 |
| OS06G0484500 | Conserved hypothetical protein. (Os06t0484500-01) | -2.25009 | - | - |
| OS08G0513051 | Hypothetical protein. (Os08t0513051-00) | -2.24948 | - | - |
| OS06G0306300 | Haem peroxidase domain containing protein. (Os06t0306300-01) | -2.24521 | - | - |
| OS03G0800500 | Putative small multi-drug export family protein. (Os03t0800500-01);Putative small multi-drug export family protein. (Os03t080050( | -2.24344 | -1.11026 | #N/A |
| OS08G0103100 | Hypothetical protein. (Os08t0103100-01) | -2.23128 | - | - |
| OS11G0474800 | Similar to Isoform 2 of Stemar-13-ene synthase. (Os11t0474800-01) | -2.2251 | #N/A | -0.7842694 |
| OS03G0692000 | Glycosyl transferase, family 14 protein. (Os03t0692000-00) | -2.22251 | - | - |
| OS07G0614700 | SPX, N-terminal domain containing protein. (Os07t0614700-01) | -2.22 | - | - |
| OS01G0967200 | Similar to Rac GTPase activating protein 1. (Os01t0967200-01) | -2.21865 | - | - |
| OS01G0814100 | Similar to Bindin (Fragment). (Os01t0814100-01) | -2.2172 | - | - |
| OS01G0281600 | Cupredoxin domain containing protein. (Os01t0281600-01) | -2.21391 | -0.982714 | #N/A |
| OS05G0439400 | Similar to Arm repeat containing protein. (Os05t0439400-01) | -2.21117 | - | - |
| OS08G0438600 | Exostosin-like family protein. (Os08t0438600-01);Exostosin-like family protein. (Os08t0438600-02) | -2.20899 | - | - |
| OS07G0162900 | Lipase, GDXG, active site domain containing protein. (Os07t0162900-01) | -2.20872 | - | - |
| OS02G0635800 | Transcription factor, TCP domain containing protein. (Os02t0635800-01) | -2.20839 | - | - |
| OS07G0162400 | Alpha/beta hydrolase fold-3 domain containing protein. (Os07t0162400-01) | -2.20651 | -1.8305 | -1.18788 |
| OS08G0269600 | EGF-like region domain containing protein. (Os08t0269600-01) | -2.20423 | - | - |
| OS09G0516750 | Hypothetical protein. (Os09t0516750-00) | -2.20199 | - | - |
| OS09G0503100 | Similar to Quinone-oxidoreductase QR1 (Fragment). (Os09t0503100-01) | -2.19248 | - | - |
| OS06G0640800 | Similar to Cytochrome P450 CYP71Y10. (Os06t0640800-00) | -2.19216 | - | - |
| OS08G0375400 | Plant disease resistance response protein family protein. (Os08t0375400-01) | -2.1804 | -1.58319 | -0.842383 |
| OS02G0200300 | Similar to Beta-1,3-glucanase-like protein. (Os02t0200300-01);Similar to Beta-1,3-glucanase. (Os02t0200300-02) | -2.17774 | - | - |
| OS01G0179600 | UDP-glucuronosyl/UDP-glucosyltransferase family protein. (Os01t0179600-01) | -2.17692 | - | - |
| OS08G0547800 | Alpha/beta hydrolase fold-3 domain containing protein. (Os08t0547800-01) | -2.16742 | - | - |
| OS03G0115000 | Cupredoxin domain containing protein. (Os03t0115000-01) | -2.1673 | - | - |
| OS02G0775300 | Conserved hypothetical protein. (Os02t0775300-01) | -2.16122 | -1.0364 | #N/A |
| OS08G0339200 | Uncharacterized conserved protein UCP016210 domain containing protein. (Os08t0339200-01) | -2.15801 | - | - |

| | | | | |
|---|---|---|---|---|
| OS08G0485000 | Similar to PHI-1. (Os08t0485000-01) | -2.14788 | - | - |
| OS08G0340900 | Pentatricopeptide repeat domain containing protein. (Os08t0340900-01) | -2.14768 | -2.47285 | -1.47888 |
| OS12G0137100 | EF-Hand type domain containing protein. (Os12t0137100-01) | -2.14087 | - | - |
| OS05G0500500 | Heat shock protein Hsp20 domain containing protein. (Os05t0500500-01) | -2.14028 | - | - |
| OS05G0391600 | Glycosyltransferase AER61, uncharacterized domain containing protein. (Os05t0391600-01) | -2.13437 | - | - |
| OS02G0507100 | Conserved hypothetical protein. (Os02t0507100-01) | -2.12685 | - | - |
| OS11G0173900 | Similar to Leucine Rich Repeat family protein, expressed. (Os11t0173900-00) | -2.12453 | #N/A | -2.40773 |
| OS02G0246900 | Conserved hypothetical protein. (Os02t0246900-01) | -2.12409 | -0.924848 | #N/A |
| OS09G0242800 | Zinc finger, RING/FYVE/PHD-type domain containing protein. (Os09t0242800-01) | -2.11947 | - | - |
| OS03G0769700 | Uncharacterised conserved protein UCP031279 domain containing protein. (Os03t0769700-01) | -2.1169 | - | - |
| OS06G0219900 | Similar to Phi-1 protein. (Os06t0219900-01) | -2.11414 | - | - |
| OS04G0483500 | Beta-ketoacyl-CoA reductase, Cuticular wax biosynthesis, Fatty acid elongation (Os04t0483500-01) | -2.1141 | -0.596536 | #N/A |
| OS04G0305700 | Similar to OSIGBa0096F13.5 protein. (Os04t0305700-01);Similar to OSIGBa0096F13.5 protein. (Os04t0305700-02) | -2.11336 | - | - |
| OS07G0633500 | Hypothetical conserved gene. (Os07t0633500-00) | -2.11033 | -1.48158 | -1.3997 |
| OS03G0219900 | Similar to 50S ribosomal protein L15, chloroplast precursor (CL15) (Fragment). (Os03t0219900-01) | -2.1086 | - | - |
| OS04G0463000 | Serine/threonine protein kinase domain containing protein. (Os04t0463000-01) | -2.10525 | - | - |
| OS05G0232700 | Conserved hypothetical protein. (Os05t0232700-01) | -2.1031 | -1.11138 | #N/A |
| OS04G0116600 | Glucose/ribitol dehydrogenase family protein. (Os04t0116600-01) | -2.0974 | -0.719342 | -0.798051 |
| OS11G0703100 | Thaumatin, pathogenesis-related family protein. (Os11t0703100-01) | -2.09655 | - | - |
| OS07G0491900 | Similar to CTF2A (Fragment). (Os07t0491900-00) | -2.0937 | - | - |
| OS01G0749200 | Chloroplast ribosome L13 protein, Chloroplast development under low temperature conditions (Os01t0749200-01) | -2.09314 | -0.495623 | #N/A |
| OS09G0363800 | Similar to metallo-beta-lactamase family protein. (Os09t0363800-00) | -2.0926 | -0.78973 | #N/A |
| OS09G0412300 | Similar to Calmodulin-like protein. (Os09t0412300-01) | -2.08972 | -2.11682 | -1.48449 |
| OS12G0169000 | Similar to Amidase family protein, expressed. (Os12t0169000-00) | -2.08809 | - | - |
| OS10G0213100 | Similar to Gamma-aminobutyrate transaminase subunit isozyme 3 (EC 2.6.1.19). (Os10t0213100-01) | -2.08517 | -0.770429 | #N/A |
| OS04G0569300 | Similar to Membrane protein. (Os04t0569300-01);Similar to Membrane protein. (Os04t0569300-02) | -2.0829 | - | - |
| OS03G0177300 | Zinc finger, RING/FYVE/PHD-type domain containing protein. (Os03t0177300-01) | -2.08034 | -1.58541 | -1.40848 |
| OS10G0554900 | Protein of unknown function DUF566 family protein. (Os10t0554900-01);Protein of unknown function DUF566 domain containing p | -2.07845 | -0.827183 | #N/A |
| OS03G0410100 | Similar to cDNA clone:J033024M21, full insert sequence. (Os03t0410100-01) | -2.07718 | -2.51992 | #N/A |
| OS03G0839800 | Protein of unknown function DUF688 domain containing protein. (Os03t0839800-01) | -2.07356 | - | - |
| OS01G0873100 | Similar to Amidophosphoribosyltransferase, chloroplast precursor (EC 2.4.2.14) (Glutamine phosphoribosylpyrophosphate amidotra | -2.07316 | - | - |
| OS04G0578400 | Similar to Beta-ring hydroxylase (Fragment). (Os04t0578400-01) | -2.0694 | - | - |
| OS11G0249800 | Non-protein coding transcript. (Os11t0249800-00) | -2.06753 | - | - |
| OS02G0652000 | PREG-like protein. (Os02t0652000-01) | -2.06515 | - | - |
| OS01G0688600 | Protein of unknown function DUF1639 family protein. (Os01t0688600-01);Protein of unknown function DUF1639 family protein. (O | -2.06448 | -1.38685 | #N/A |
| OS07G0628600 | Origin recognition complex subunit 6 family protein. (Os07t0628600-00) | -2.0626 | - | - |
| OS02G0203300 | Similar to UDP-glycosyltransferase UGT75E3. (Os02t0203300-00) | -2.06222 | - | - |
| OS08G0446800 | Similar to GDU1. (Os08t0446800-01) | -2.05769 | - | - |
| OS07G0252800 | Conserved hypothetical protein. (Os07t0252800-01) | -2.05549 | - | - |
| OS11G0595200 | PAP fibrillin family protein. (Os11t0595200-01) | -2.05361 | -0.798134 | #N/A |
| OS01G0964900 | Similar to Mitochondrial carrier protein-like. (Os01t0964900-00) | -2.05188 | - | - |
| OS12G0630600 | Carbohydrate/puine kinase, PfkB, conserved site containing protein. (Os12t0630600-01) | -2.0462 | -0.607591 | #N/A |

| | | | | |
|---|---|---|---|---|
| OS06G0641866 | Hypothetical protein. (Os06t0641866-00) | -2.04606 | - | - |
| OS01G0197700 | Similar to Cytokinin dehydrogenase 2. (Os01t0197700-01) | -2.04254 | -2.90949 | -1.75741 |
| OS01G0863500 | Control of leaf rolling and bending (Os01t0863500-01) | -2.04049 | - | - |
| OS01G0949300 | EF-Hand type domain containing protein. (Os01t0949300-01) | -2.0348 | - | - |
| OS04G0585700 | Protein of unknown function DUF581 family protein. (Os04t0585700-01) | -2.02592 | -1.56654 | #N/A |
| OS03G0581100 | Hypothetical conserved gene. (Os03t0581100-01) | -2.02385 | - | - |
| OS07G0502900 | UDP-glucuronosyl/UDP-glucosyltransferase family protein. (Os07t0502900-00) | -2.02137 | - | - |
| OS06G0693200 | Protein kinase, core domain containing protein. (Os06t0693200-01);Hypothetical conserved gene. (Os06t0693200-02) | -2.0113 | - | - |
| OS08G0400200 | Similar to SET domain protein. (Os08t0400200-00) | -2.011 | -1.93455 | #N/A |
| OS01G0221900 | Similar to peptidase C45, acyl-coenzyme A/6-aminopenicillanic acid acyl-transferase. (Os01t0221900-01) | -2.00844 | - | - |
| OS01G0675700 | Similar to Auxin-responsive protein IAA14 (Indoleacetic acid-induced protein 14) (SOLITARY-ROOT protein). (Os01t0675700-01);Sim | -2.00729 | -0.914167 | #N/A |
| OS06G0239500 | TGF-beta receptor, type I/II extracellular region family protein. (Os06t0239500-01) | -2.00692 | - | - |
| OS08G0504500 | Mog1/PsbP/DUF1795, alpha/beta/alpha sandwich domain containing protein. (Os08t0504500-01) | -2.00663 | - | - |
| OS03G0292200 | Adenine nucleotide translocator 1 domain containing protein. (Os03t0292200-01) | -2.00535 | -1.18263 | #N/A |
| OS04G0691400 | Similar to POT family protein. (Os04t0691400-00) | -2.00398 | - | - |
| OS06G0129800 | Hypothetical conserved gene. (Os06t0129800-00) | -2.0036 | - | - |
| OS03G0284500 | HAD-superfamily subfamily IIA hydrolase, CECR5 protein. (Os03t0284500-01);Similar to cat eye syndrome critical region protein 5. ( | -2.00249 | - | - |
| OS03G0856700 | Gibberellin 20 oxidase 1 (EC 1.14.11.-) (Gibberellin C-20 oxidase 1) (GA 20-oxidase 1) (Os20ox). (Os03t0856700-01);GA 20-oxidase1, | -2.00196 | - | - |
| OS04G0623800 | Similar to Aminomethyltransferase, mitochondrial precursor (EC 2.1.2.10) (Glycine cleavage system T protein) (GCVT). (Os04t06238 | -2.00135 | - | - |
| OS01G0640700 | Similar to vegetative cell wall protein gp1. (Os01t0640700-00) | -1.9982 | #N/A | #N/A |
| OS11G0558400 | Leucine-rich repeat, N-terminal domain containing protein. (Os11t0558400-00) | -1.99661 | #N/A | #N/A |
| OS09G0116900 | Conserved hypothetical protein. (Os09t0116900-01) | -1.99512 | #N/A | #N/A |
| OS04G0401700 | Potassium transporter, Potassium mediated growth, Salt tolerance (Os04t0401700-01);Similar to Isoform 2 of Potassium transporte | -1.99016 | -1.84784 | #N/A |
| OS10G0352000 | Conserved hypothetical protein. (Os10t0352000-01) | -1.98889 | #N/A | #N/A |
| OS02G0170000 | Conserved hypothetical protein. (Os02t0170000-01);Pentatricopeptide repeat domain containing protein. (Os02t0170000-02) | -1.9883 | -1.36274 | -0.713721 |
| OS03G0363400 | Uncharacterised protein family UPF0153 domain containing protein. (Os03t0363400-01) | -1.98783 | -0.996852 | -0.921394 |
| OS03G0161400 | Similar to IQ calmodulin-binding motif family protein, expressed. (Os03t0161400-01) | -1.98537 | #N/A | #N/A |
| OS02G0625300 | Similar to PEPC kinase. (Os02t0625300-00) | -1.98359 | -0.703938 | #N/A |
| OS07G0613500 | Similar to Protein kinase APK1B, chloroplast precursor (EC 2.7.1.-). (Os07t0613500-01) | -1.98147 | -1.216 | #N/A |
| OS07G0142100 | Conserved hypothetical protein. (Os07t0142100-00) | -1.98056 | #N/A | 1.47313 |
| OS03G0780900 | Protein of unknown function DUF858, methyltransferase-like family protein. (Os03t0780900-01) | -1.97153 | #N/A | -1.03325 |
| OS05G0215033 | Hypothetical gene. (Os05t0215033-01) | -1.96975 | -1.38497 | #N/A |
| OS02G0243300 | UDP-glucuronosyl/UDP-glucosyltransferase family protein. (Os02t0243300-01) | -1.96446 | #N/A | #N/A |
| OS02G0557800 | A-type response regulator, Cytokinin signaling (Os02t0557800-01);A-type response regulator, Cytokinin signaling (Os02t0557800-02 | -1.95885 | #N/A | #N/A |
| OS02G0771700 | Glycoside hydrolase, family 17 protein. (Os02t0771700-01) | -1.95847 | #N/A | #N/A |
| OS12G0541700 | Similar to Rapid alkalinization factor 2. (Os12t0541700-01) | -1.95659 | #N/A | #N/A |
| OS02G0703300 | Protein of unknown function DUF1218 family protein. (Os02t0703300-01);Protein of unknown function DUF1218 family protein. (Os | -1.94439 | #N/A | #N/A |
| OS02G0596300 | Cytochrome P450 domain containing protein. (Os02t0596300-01) | -1.94377 | #N/A | #N/A |
| OS06G0691200 | Similar to Thaumatin-like protein precursor. (Os06t0691200-01) | -1.93863 | #N/A | #N/A |
| OS02G0660100 | Hypothetical conserved gene. (Os02t0660100-00) | -1.93832 | #N/A | #N/A |
| OS09G0454100 | Peptidase S54, rhomboid domain containing protein. (Os09t0454100-01) | -1.93727 | #N/A | #N/A |
| OS01G0891000 | Glycoside hydrolase, family 20 protein. (Os01t0891000-01) | -1.93697 | #N/A | #N/A |

| | | | | |
|---|---|---|---|---|
| OS01G0221700 | Similar to GDT1-like protein 1, chloroplastic. (Os01t0221700-00) | -1.93276 | #N/A | #N/A |
| OS08G0452500 | Auxin responsive SAUR protein family protein. (Os08t0452500-01);Auxin responsive SAUR protein family protein. (Os08t0452500-0[illegible] | -1.9301 | #N/A | #N/A |
| OS11G0702400 | Zinc finger, C2H2-type domain containing protein. (Os11t0702400-01) | -1.92901 | -1.97594 | -3.96894 |
| OS05G0338900 | Oligopeptide transporter domain containing protein. (Os05t0338900-00) | -1.92577 | #N/A | #N/A |
| OS07G0558300 | Inositol monophosphatase family protein. (Os07t0558300-01) | -1.92104 | -0.565395 | -0.590643 |
| OS05G0475700 | Nodulin-like domain containing protein. (Os05t0475700-01);Similar to nodulin-like protein. (Os05t0475700-02) | -1.9208 | #N/A | #N/A |
| OS01G0819700 | Hypothetical conserved gene. (Os01t0819700-01) | -1.9193 | -1.17685 | #N/A |
| OS08G0433600 | Protein of unknown function DUF962 family protein. (Os08t0433600-01) | -1.91568 | -0.668555 | #N/A |
| OS12G0547100 | Conserved hypothetical protein. (Os12t0547100-01);Conserved hypothetical protein. (Os12t0547100-02) | -1.91212 | #N/A | #N/A |
| OS11G0598500 | NB-ARC domain containing protein. (Os11t0598500-00) | -1.91209 | #N/A | #N/A |
| OS04G0477000 | BTB/POZ fold domain containing protein. (Os04t0477000-01) | -1.91159 | #N/A | #N/A |
| OS04G0185100 | DNA replication factor CDT1-like domain containing protein. (Os04t0185100-01) | -1.90648 | #N/A | #N/A |
| OS01G0668100 | Similar to Arabinogalactan-like protein. (Os01t0668100-01) | -1.90374 | #N/A | #N/A |
| OS01G0952800 | Achaete-scute transcription factor related domain containing protein. (Os01t0952800-01);Conserved hypothetical protein. (Os01t0[illegible] | -1.9034 | -1.46362 | #N/A |
| OS01G0175700 | Boron (B) transporter (Os01t0175700-01) | -1.90044 | -1.78697 | -1.67997 |
| OS10G0502600 | Cytochrome b5 domain containing protein. (Os10t0502600-01);Similar to Membrane steroid-binding protein 1 (Fragment). (Os10t0[illegible] | -1.89859 | #N/A | #N/A |
| OS02G0140300 | Molybdopterin biosynthesis MoaE family protein. (Os02t0140300-01) | -1.89826 | #N/A | #N/A |
| OS08G0235550 | Conserved hypothetical protein. (Os08t0235550-01) | -1.8977 | #N/A | #N/A |
| OS08G0463100 | Conserved hypothetical protein. (Os08t0463100-01) | -1.89408 | #N/A | #N/A |
| OS02G0567200 | Protein phosphatase 2C domain containing protein. (Os02t0567200-01) | -1.89237 | #N/A | #N/A |
| OS04G0136700 | Cystathionine beta-synthase, core domain containing protein. (Os04t0136700-01) | -1.89203 | -0.886066 | #N/A |
| OS02G0773600 | Hypothetical conserved gene. (Os02t0773600-00) | -1.88476 | #N/A | #N/A |
| OS03G0708500 | Capsid/spike protein, ssDNA virus domain containing protein. (Os03t0708500-00) | -1.88305 | -0.820401 | #N/A |
| OS09G0504000 | Similar to Nucleotide sugar epimerase-like protein (UDP-D-glucuronate 4- epimerase) (EC 5.1.3.6). (Os09t0504000-01) | -1.88025 | #N/A | #N/A |
| OS05G0147500 | Similar to DEGP2 (DEGP PROTEASE 2); serine-type peptidase/ trypsin. (Os05t0147500-01);Serine endopeptidase DegP2 domain con[illegible] | -1.87369 | -0.620543 | -0.61502 |
| OS11G0643100 | Transferase family protein. (Os11t0643100-01) | -1.87188 | #N/A | #N/A |
| OS04G0644433 | Hypothetical genes. (Os04t0644433-00) | -1.86951 | #N/A | #N/A |
| OS04G0412800 | Thioredoxin fold domain containing protein. (Os04t0412800-01) | -1.86915 | #N/A | #N/A |
| OS10G0575900 | Conserved hypothetical protein. (Os10t0575900-00) | -1.86683 | #N/A | #N/A |
| OS01G0266100 | Similar to RING finger family protein. (Os01t0266100-01) | -1.86609 | #N/A | #N/A |
| OS01G0974400 | Zinc finger, RING/FYVE/PHD-type domain containing protein. (Os01t0974400-01) | -1.86538 | #N/A | #N/A |
| OS06G0153600 | Conserved hypothetical protein. (Os06t0153600-01);Conserved hypothetical protein. (Os06t0153600-02) | -1.86432 | #N/A | #N/A |
| OS03G0249100 | Conserved hypothetical protein. (Os03t0249100-01) | -1.86396 | #N/A | #N/A |
| OS02G0174100 | Similar to Isoform 2 of Squamosa promoter-binding-like protein 4. (Os02t0174100-01);SBP domain containing protein. (Os02t01741[illegible] | -1.85828 | #N/A | #N/A |
| OS11G0425600 | Four F5 protein family protein. (Os11t0425600-01) | -1.85413 | -0.803032 | #N/A |
| OS01G0349600 | Hypothetical protein. (Os01t0349600-01) | -1.84937 | #N/A | #N/A |
| OS12G0508266 | Histone deacetylase superfamily protein. (Os12t0508266-01) | -1.84923 | #N/A | #N/A |
| OS10G0531400 | Glutathione S-transferase GST 30 (EC 2.5.1.18). (Os10t0531400-01) | -1.84618 | -1.46025 | -1.10839 |
| OS04G0454200 | Similar to Monosaccharide transporter 1. (Os04t0454200-01) | -1.84398 | #N/A | #N/A |
| OS01G0915900 | Conserved hypothetical protein. (Os01t0915900-01) | -1.84381 | #N/A | #N/A |
| OS04G0658800 | Similar to OSIGBa0132E09-OSIGBa0108L24.9 protein. (Os04t0658800-01) | -1.84242 | #N/A | #N/A |
| OS12G0215950 | Similar to Leucine Rich Repeat family protein, expressed. (Os12t0215950-01) | -1.84211 | -3.20577 | -4.03763 |

| | | | | |
|---|---|---|---|---|
| OS03G0705800 | Protein of unknown function DUF3049 domain containing protein. (Os03t0705800-00) | -1.8374 | #N/A | #N/A |
| OS11G0215100 | Plant disease resistance response protein family protein. (Os11t0215100-01) | -1.83699 | #N/A | #N/A |
| OS03G0135600 | Similar to Ankyrin repeat protein. (Os03t0135600-01) | -1.83471 | -0.933318 | -0.755165 |
| OS07G0542600 | Similar to receptor-like protein kinase RK20-1. (Os07t0542600-01) | -1.83276 | #N/A | #N/A |
| OS01G0881300 | Non-protein coding transcript. (Os01t0881300-01);MtN3 and saliva related transmembrane protein family protein. (Os01t0881300- | -1.83061 | -0.738141 | #N/A |
| OS02G0717100 | Esterase/lipase/thioesterase domain containing protein. (Os02t0717100-01) | -1.82644 | #N/A | #N/A |
| OS01G0839200 | DUF966 family protein, Abiotic stress response (Os01t0839200-01) | -1.82498 | -1.32432 | #N/A |
| OS09G0460400 | Alpha/beta hydrolase fold-3 domain containing protein. (Os09t0460400-01) | -1.82384 | #N/A | #N/A |
| OS10G0140300 | Esterase, SGNH hydrolase-type domain containing protein. (Os10t0140300-01) | -1.82327 | #N/A | #N/A |
| OS10G0433700 | Hypothetical protein. (Os10t0433700-01) | -1.82006 | #N/A | #N/A |
| OS01G0756900 | Hypothetical protein. (Os01t0756900-01) | -1.81492 | #N/A | #N/A |
| OS08G0550600 | Hypothetical conserved gene. (Os08t0550600-01) | -1.81262 | #N/A | #N/A |
| OS03G0116900 | Armadillo-like helical domain containing protein. (Os03t0116900-01);Armadillo-like helical domain containing protein. (Os03t01169 | -1.81125 | #N/A | #N/A |
| OS01G0931200 | Armadillo-type fold domain containing protein. (Os01t0931200-01) | -1.81079 | #N/A | #N/A |
| OS05G0563600 | FAS1 domain domain containing protein. (Os05t0563600-01) | -1.80778 | #N/A | #N/A |
| OS02G0542500 | Similar to NF-180. (Os02t0542500-01) | -1.80515 | #N/A | #N/A |
| OS08G0490100 | Similar to PBF protein. (Os08t0490100-01) | -1.80464 | #N/A | #N/A |
| OS10G0562000 | Similar to glycoprotein. (Os10t0562000-01) | -1.79325 | #N/A | #N/A |
| OS11G0153300 | Conserved hypothetical protein. (Os11t0153300-01);Conserved hypothetical protein. (Os11t0153300-02) | -1.79251 | #N/A | #N/A |
| OS01G0205600 | Crotonase, core domain containing protein. (Os01t0205600-01) | -1.79177 | #N/A | #N/A |
| OS03G0388100 | ATPase, P-type, K/Mg/Cd/Cu/Zn/Na/Ca/Na/H-transporter domain containing protein. (Os03t0388100-01);ATPase, P-type, K/Mg/Cd | -1.7906 | #N/A | #N/A |
| OS06G0664200 | Inositol monophosphatase/Fructose-1,6-bisphosphatase domain containing protein. (Os06t0664200-01);Hypothetical gene. (Os06t( | -1.79033 | #N/A | #N/A |
| OS04G0480200 | Transporter, high affinity nitrate, Nar2 domain containing protein. (Os04t0480200-01) | -1.78968 | -1.56087 | #N/A |
| OS01G0152700 | Similar to Histone H2B (CaH2B). (Os01t0152700-01) | -1.78758 | #N/A | #N/A |
| OS09G0278700 | Hypothetical conserved gene. (Os09t0278700-00) | -1.78757 | #N/A | #N/A |
| OS10G0165400 | Hypothetical gene. (Os10t0165400-01) | -1.78727 | #N/A | #N/A |
| OS07G0669200 | Similar to GTP1/OBG family protein. (Os07t0669200-00) | -1.78613 | -1.34974 | -0.952073 |
| OS12G0613250 | Similar to BTB/POZ; Superoxide dismutase, copper/zinc binding; NPH3. (Os12t0613250-00) | -1.7805 | #N/A | #N/A |
| OS10G0577900 | Similar to Glycerol-3-phosphate acyltransferase (Fragment). (Os10t0577900-01) | -1.7792 | -0.913521 | -0.725819 |
| OS12G0273900 | Leucine-rich repeat domain containing protein. (Os12t0273900-00) | -1.77883 | #N/A | #N/A |
| OS05G0560100 | Similar to Pollen-specific kinase partner protein. (Os05t0560100-01) | -1.77866 | #N/A | #N/A |
| OS03G0769100 | Similar to 9S ribosomal protein. (Os03t0769100-01);Non-protein coding transcript. (Os03t0769100-02) | -1.77606 | #N/A | #N/A |
| OS04G0179200 | Similar to Stem secoisolariciresinol dehydrogenase (Fragment). (Os04t0179200-01) | -1.7745 | -2.77597 | #N/A |
| OS03G0807100 | Protein of unknown function DUF239 domain containing protein. (Os03t0807100-01) | -1.77245 | #N/A | #N/A |
| OS08G0506500 | Similar to tRNA/rRNA methyltransferase (SpoU) family protein. (Os08t0506500-00) | -1.77187 | #N/A | #N/A |
| OS02G0177700 | Uncharacterised protein family UPF0497, trans-membrane plant subgroup domain containing protein. (Os02t0177700-01) | -1.77047 | #N/A | #N/A |
| OS08G0335600 | Protein of unknown function DUF568, DOMON-like domain containing protein. (Os08t0335600-01) | -1.76216 | #N/A | #N/A |
| OS06G0301500 | Hypothetical protein. (Os06t0301500-01) | -1.76086 | #N/A | #N/A |
| OS06G0146300 | Conserved hypothetical protein. (Os06t0146300-01) | -1.7606 | #N/A | #N/A |
| OS03G0797800 | AUX/IAA protein family protein. (Os03t0797800-01) | -1.76009 | #N/A | #N/A |
| OS12G0155100 | Protein of unknown function DUF3049 domain containing protein. (Os12t0155100-01) | -1.76005 | #N/A | #N/A |
| OS01G0140100 | Peptidase A1 domain containing protein. (Os01t0140100-01) | -1.75999 | -0.766706 | #N/A |

| | | | | |
|---|---|---|---|---|
| OS01G0651800 | Lipase, class 3 family protein. (Os01t0651800-01) | -1.75839 | #N/A | #N/A |
| OS04G0481300 | Methyltransferase type 11 domain containing protein. (Os04t0481300-01) | -1.75709 | #N/A | #N/A |
| OS01G0629750 | Conserved hypothetical protein. (Os01t0629750-00) | -1.75451 | #N/A | #N/A |
| OS01G0279400 | Major facilitator superfamily antiporter. (Os01t0279400-01) | -1.75328 | -0.945692 | -0.706929 |
| OS01G0585200 | Similar to cDNA clone:J023088C01, full insert sequence. (Os01t0585200-01) | -1.75144 | -2.4032 | #N/A |
| OS01G0796500 | Similar to predicted protein. (Os01t0796500-01) | -1.75087 | #N/A | #N/A |
| OS01G0836600 | ABC transporter-like domain containing protein. (Os01t0836600-01) | -1.7482 | #N/A | #N/A |
| OS01G0271400 | Peptidase S14, ClpP family protein. (Os01t0271400-00) | -1.74486 | -0.723456 | #N/A |
| OS11G0447300 | GTP-binding protein, HSR1-related domain containing protein. (Os11t0447300-01) | -1.74444 | #N/A | -1.49772 |
| OS03G0750700 | Six-bladed beta-propeller, TolB-like domain containing protein. (Os03t0750700-01) | -1.74388 | #N/A | #N/A |
| OS08G0310100 | Ribosomal protein L15 family protein. (Os08t0310100-01) | -1.74348 | -0.887111 | #N/A |
| OS08G0526100 | NAD(P)-binding domain containing protein. (Os08t0526100-01) | -1.74287 | #N/A | #N/A |
| OS07G0673550 | Photosystem II protein PsbX family protein. (Os07t0673550-00) | -1.74223 | #N/A | #N/A |
| OS02G0754300 | Ribosomal protein L29 family protein. (Os02t0754300-01) | -1.74113 | #N/A | #N/A |
| OS06G0141166 | Similar to hydrolase, NUDIX family protein. (Os06t0141166-00) | -1.73803 | -1.59114 | -1.20635 |
| OS10G0573700 | Similar to Mitochondrial carnitine/acylcarnitine carrier-like protein (A BOUT DE SOUFFLE) (Carnitine/acylcarnitine translocase-like p | -1.73197 | #N/A | #N/A |
| OS07G0117000 | NB-ARC domain containing protein. (Os07t0117000-01) | -1.72996 | -1.55393 | -1.78369 |
| OS10G0519700 | Mitochondrial import inner membrane translocase, subunit Tim17/22 family protein. (Os10t0519700-01) | -1.72868 | #N/A | #N/A |
| OS09G0284300 | Conserved hypothetical protein. (Os09t0284300-01);Conserved hypothetical protein. (Os09t0284300-02) | -1.72846 | #N/A | #N/A |
| OS05G0406000 | Conserved hypothetical protein. (Os05t0406000-01) | -1.7268 | -1.19943 | -0.64504 |
| OS03G0850000 | Glucose/ribitol dehydrogenase family protein. (Os03t0850000-01) | -1.72641 | #N/A | #N/A |
| OS11G0241700 | Protein of unknown function DUF538 family protein. (Os11t0241700-01) | -1.72535 | #N/A | #N/A |
| OS08G0440900 | Similar to Omega-6 fatty acid desaturase, chloroplast precursor (EC 1.14.19.-). (Os08t0440900-01) | -1.72511 | #N/A | #N/A |
| OS01G0971800 | Similar to Two-component response regulator ARR11 (Receiver-like protein 3). (Os01t0971800-00) | -1.72389 | -1.19361 | #N/A |
| OS03G0346500 | Conserved hypothetical protein. (Os03t0346500-01) | -1.72042 | #N/A | #N/A |
| OS08G0544800 | PCF2. (Os08t0544800-01) | -1.71973 | #N/A | #N/A |
| OS03G0301200 | COBRA-like protein 7 precursor. (Os03t0301200-01);Glycosyl-phosphatidyl inositol-anchored, plant domain containing protein. (Os0 | -1.7185 | -1.43033 | -1.8932 |
| OS01G0796700 | Zinc finger, RING/FYVE/PHD-type domain containing protein. (Os01t0796700-01) | -1.7176 | #N/A | #N/A |
| OS02G0219800 | Tetraspanin domain containing protein. (Os02t0219800-01) | -1.71748 | #N/A | #N/A |
| OS05G0499600 | UDP-glucuronosyl/UDP-glucosyltransferase family protein. (Os05t0499600-01) | -1.71687 | #N/A | #N/A |
| OS08G0548400 | Similar to chaperone protein dnaJ 11. (Os08t0548400-00) | -1.71614 | #N/A | #N/A |
| OS11G0242600 | Protein of unknown function DUF579, plant family protein. (Os11t0242600-00) | -1.71366 | #N/A | #N/A |
| OS03G0781000 | Conserved hypothetical protein. (Os03t0781000-01) | -1.71221 | #N/A | #N/A |
| OS02G0257300 | Rhodanese-like domain containing protein. (Os02t0257300-00) | -1.70817 | #N/A | #N/A |
| OS09G0488000 | Acyl-CoA N-acyltransferase domain containing protein. (Os09t0488000-01) | -1.70792 | #N/A | #N/A |
| OS07G0188233 | Conserved hypothetical protein. (Os07t0188233-01) | -1.70765 | #N/A | #N/A |
| OS01G0111100 | Cyclophilin-like domain containing protein. (Os01t0111100-01) | -1.705 | #N/A | #N/A |
| OS01G0958100 | Similar to chloroplast SRP receptor cpFtsY precursor. (Os01t0958100-01);Similar to chloroplast srp54 receptor1. (Os01t0958100-02) | -1.70342 | #N/A | #N/A |
| OS01G0934100 | C2 calcium-dependent membrane targeting domain containing protein. (Os01t0934100-01) | -1.70278 | -1.38984 | #N/A |
| OS06G0600301 | Similar to GTP-binding nuclear protein Ran1B (Fragment). (Os06t0600301-00) | -1.70272 | #N/A | #N/A |
| OS12G0514000 | Similar to Sorbitol transporter. (Os12t0514000-01) | -1.70244 | #N/A | #N/A |
| OS01G0776300 | Protein of unknown function DUF26 domain containing protein. (Os01t0776300-01) | -1.70243 | #N/A | #N/A |

| | | | | |
|---|---|---|---|---|
| OS01G0291500 | Transferase family protein. (Os01t0291500-01);Transferase family protein. (Os01t0291500-02) | -1.70052 | #N/A | #N/A |
| OS11G0432600 | Similar to Beta-keto acyl reductase (Fragment). (Os11t0432600-01) | -1.69504 | #N/A | #N/A |
| OS11G0163800 | Protein of unknown function DUF793 family protein. (Os11t0163800-01) | -1.69156 | #N/A | #N/A |
| OS11G0528500 | Similar to Rubredoxin 1 (Rd-1). (Os11t0528500-01) | -1.69101 | #N/A | #N/A |
| OS12G0575000 | Protein of unknown function DUF1118 family protein. (Os12t0575000-01) | -1.68773 | -1.04476 | #N/A |
| OS07G0625800 | Plant lipid transfer protein/Par allergen family protein. (Os07t0625800-01);Plant lipid transfer protein/Par allergen family protein. ( | -1.68538 | #N/A | #N/A |
| OS01G0338150 | Hypothetical gene. (Os01t0338150-01) | -1.6849 | #N/A | #N/A |
| OS07G0606800 | Alpha/beta hydrolase fold-3 domain containing protein. (Os07t0606800-01) | -1.68324 | #N/A | #N/A |
| OS02G0440000 | Similar to Diaminopimelate decarboxylase. (Os02t0440000-01);Similar to cDNA clone:J013098O07, full insert sequence. (Os02t044C | -1.68144 | -0.699542 | #N/A |
| OS07G0544800 | Similar to Oxygen-evolving enhancer protein 3-2, chloroplast precursor (OEE3) (16 kDa subunit of oxygen evolving system of photos | -1.68116 | #N/A | #N/A |
| OS07G0601600 | Similar to VEP1 (VEIN PATTERNING 1); binding / catalytic. (Os07t0601600-00) | -1.68095 | #N/A | #N/A |
| OS04G0572800 | Similar to OSIGBa0147H17.4 protein. (Os04t0572800-00) | -1.68086 | #N/A | #N/A |
| OS03G0763000 | Similar to Casein kinase II alpha subunit. (Os03t0763000-01) | -1.68066 | #N/A | #N/A |
| OS10G0509200 | Similar to plastid-lipid associated protein PAP / fibrillin family protein. (Os10t0509200-01) | -1.67817 | #N/A | #N/A |
| OS06G0693000 | Protein kinase, core domain containing protein. (Os06t0693000-01) | -1.67662 | #N/A | #N/A |
| OS04G0689300 | PLC-like phosphodiesterase, TIM beta/alpha-barrel domain domain containing protein. (Os04t0689300-01) | -1.676 | #N/A | #N/A |
| OS07G0681600 | DNA helicase, ATP-dependent, RecQ type domain containing protein. (Os07t0681600-01) | -1.67204 | #N/A | -2.34365 |
| OS12G0616000 | Similar to Kinesin-like protein KIFC3. (Os12t0616000-01) | -1.67146 | #N/A | #N/A |
| OS09G0286600 | Pathogenesis-related transcriptional factor and ERF domain containing protein. (Os09t0286600-01);Pathogenesis-related transcript | -1.67088 | #N/A | #N/A |
| OS12G0543600 | Similar to sarcosine oxidase. (Os12t0543600-00) | -1.6693 | #N/A | #N/A |
| OS05G0182201 | Similar to bifunctional protein tilS/hprT. (Os05t0182201-01) | -1.66885 | #N/A | #N/A |
| OS10G0144700 | Cytochrome P450 domain containing protein. (Os10t0144700-00) | -1.66775 | #N/A | #N/A |
| OS01G0149400 | Hypothetical protein. (Os01t0149400-00) | -1.66695 | #N/A | #N/A |
| OS02G0106800 | TMPIT-like family protein. (Os02t0106800-01) | -1.66499 | #N/A | #N/A |
| OS01G0676200 | Conserved hypothetical protein. (Os01t0676200-01);Conserved hypothetical protein. (Os01t0676200-02) | -1.66124 | -0.976017 | #N/A |
| OS06G0284500 | Zinc finger, Dof-type family protein. (Os06t0284500-00) | -1.65884 | -1.07367 | #N/A |
| OS01G0726001 | Hypothetical gene. (Os01t0726001-00) | -1.65873 | #N/A | #N/A |
| OS02G0281150 | Similar to Leucine Rich Repeat family protein. (Os02t0281150-01) | -1.65795 | #N/A | #N/A |
| OS10G0486100 | Cytochrome P450-like protein (CYP86B1). (Os10t0486100-00) | -1.65589 | #N/A | #N/A |
| OS07G0591100 | Protein of unknown function DUF620 family protein. (Os07t0591100-01) | -1.65525 | #N/A | #N/A |
| OS07G0160100 | Similar to cDNA clone:J023038F18, full insert sequence. (Os07t0160100-01);YABBY family transcription factor, Stamen and carpel d | -1.65345 | #N/A | -1.1666 |
| OS05G0568800 | Protein of unknown function DUF1645 family protein. (Os05t0568800-01) | -1.65302 | #N/A | #N/A |
| OS11G0291475 | Hypothetical protein. (Os11t0291475-00) | -1.65299 | #N/A | 0.828085 |
| OS06G0308200 | Similar to 60S ribosomal protein L15. (Os06t0308200-01) | -1.65285 | #N/A | #N/A |
| OS11G0585700 | Similar to 50S ribosomal protein L4. (Os11t0585700-01) | -1.65239 | #N/A | #N/A |
| OS11G0505300 | Sulfotransferase, Resistance to rice stripe virus (Os11t0505300-01) | -1.65212 | #N/A | #N/A |
| OS10G0130500 | Hypothetical protein. (Os10t0130500-01);Hypothetical protein. (Os10t0130500-02) | -1.64856 | -1.24258 | -0.851958 |
| OS05G0501200 | Hypothetical conserved gene. (Os05t0501200-00) | -1.64511 | #N/A | #N/A |
| OS05G0495900 | Similar to Beta-1,3-glucanase precursor (Fragment). (Os05t0495900-01) | -1.64456 | -0.659245 | #N/A |
| OS12G0507400 | Conserved hypothetical protein. (Os12t0507400-00) | -1.64422 | #N/A | #N/A |
| OS08G0194350 | F-box domain, Skp2-like domain containing protein. (Os08t0194350-00) | -1.64346 | #N/A | #N/A |
| OS05G0490500 | Conserved hypothetical protein. (Os05t0490500-01) | -1.64317 | #N/A | #N/A |

| | | | | |
|---|---|---|---|---|
| OS05G0112800 | Protein of unknown function DUF26 domain containing protein. (Os05t0112800-01) | -1.64287 | #N/A | #N/A |
| OS07G0656700 | Uncharacterised conserved protein UCP022348 domain containing protein. (Os07t0656700-01) | -1.64185 | -0.913853 | -0.764983 |
| OS02G0662000 | RCc3 protein. (Os02t0662000-01) | -1.64129 | #N/A | #N/A |
| OS01G0559500 | Similar to predicted protein. (Os01t0559500-01) | -1.64073 | -0.858081 | #N/A |
| OS05G0514400 | Hypothetical conserved gene. (Os05t0514400-00) | -1.63958 | -1.23464 | #N/A |
| OS06G0581000 | Similar to Nitrate transporter NTL1. (Os06t0581000-01) | -1.63945 | #N/A | #N/A |
| OS12G0633000 | Xylulose kinase. (Os12t0633000-01) | -1.63904 | #N/A | #N/A |
| OS01G0225400 | Pyruvate/Phosphoenolpyruvate kinase, catalytic core domain containing protein. (Os01t0225400-01) | -1.63871 | #N/A | #N/A |
| OS05G0429500 | Similar to dienelactone hydrolase family protein. (Os05t0429500-01);Dienelactone hydrolase domain containing protein. (Os05t042 | -1.63836 | #N/A | #N/A |
| OS11G0629400 | Similar to Nitrilase associated protein-like. (Os11t0629400-01) | -1.63669 | #N/A | #N/A |
| OS04G0676100 | Similar to Thioredoxin X, chloroplast precursor. (Os04t0676100-01) | -1.63406 | #N/A | #N/A |
| OS09G0416900 | Armadillo-like helical domain containing protein. (Os09t0416900-00) | -1.63368 | #N/A | #N/A |
| OS12G0491800 | Similar to Ent-kaurene synthase 1A. (Os12t0491800-01) | -1.63355 | #N/A | #N/A |
| OS07G0605400 | Similar to EGG APPARATUS-1 protein (ZmEA1). (Os07t0605400-01);Similar to EGG APPARATUS-1 protein (ZmEA1). (Os07t0605400-0 | -1.63179 | -1.57511 | -1.17663 |
| OS04G0110400 | Similar to H0201G08.9 protein. (Os04t0110400-01) | -1.63155 | #N/A | #N/A |
| OS04G0445700 | Similar to OSIGBa0140O07.8 protein. (Os04t0445700-01);Similar to 3-oxoacyl-[acyl-carrier-protein] synthase I, chloroplast precurso | -1.62921 | #N/A | #N/A |
| OS03G0388000 | Similar to Splicing factor SC35. (Os03t0388000-01);Hypothetical conserved gene. (Os03t0388000-02) | -1.62809 | #N/A | #N/A |
| OS07G0467200 | Similar to Clone ZZD536 mRNA sequence. (Os07t0467200-01) | -1.62738 | #N/A | #N/A |
| OS06G0700300 | Conserved hypothetical protein. (Os06t0700300-01) | -1.62609 | #N/A | #N/A |
| OS01G0194300 | Ankyrin-repeat protein, Herbivore-induced defense response, Blast disease resistance (Os01t0194300-01) | -1.62547 | #N/A | #N/A |
| OS07G0175500 | Similar to lipid binding protein. (Os07t0175500-00) | -1.62404 | #N/A | #N/A |
| OS08G0230000 | Conserved hypothetical protein. (Os08t0230000-01) | -1.61992 | #N/A | #N/A |
| OS04G0400600 | Similar to OSIGBa0113L04.4 protein. (Os04t0400600-00) | -1.61138 | #N/A | #N/A |
| OS03G0710700 | Similar to predicted protein. (Os03t0710700-00) | -1.61035 | #N/A | #N/A |
| OS07G0575800 | Similar to hydrolase, alpha/beta fold family protein. (Os07t0575800-01) | -1.60844 | -1.21447 | -1.19698 |
| OS01G0827700 | Lipase, GDSL domain containing protein. (Os01t0827700-00) | -1.60669 | #N/A | #N/A |
| OS07G0250501 | Conserved hypothetical protein. (Os07t0250501-00) | -1.60545 | #N/A | #N/A |
| OS03G0254000 | NB-ARC domain containing protein. (Os03t0254000-01) | -1.60438 | #N/A | #N/A |
| OS09G0480600 | Conserved hypothetical protein. (Os09t0480600-01) | -1.60401 | -0.979285 | -0.978426 |
| OS10G0463200 | Esterase, SGNH hydrolase-type domain containing protein. (Os10t0463200-01) | -1.60342 | #N/A | #N/A |
| OS05G0500000 | UDP-glucuronosyl/UDP-glucosyltransferase family protein. (Os05t0500000-00) | -1.60283 | #N/A | #N/A |
| OS05G0571400 | Protein of unknown function DUF3593 domain containing protein. (Os05t0571400-01);Hypothetical conserved gene. (Os05t057140 | -1.60278 | -0.707567 | #N/A |
| OS03G0315800 | Similar to 30S ribosomal protein S1, chloroplast precursor (CS1). (Os03t0315800-01);Similar to 30S ribosomal protein S1 (Fragment) | -1.60089 | #N/A | #N/A |
| OS10G0112500 | Similar to POT family protein, expressed. (Os10t0112500-00) | -1.59937 | #N/A | #N/A |
| OS01G0758500 | Similar to predicted protein. (Os01t0758500-01);Conserved hypothetical protein. (Os01t0758500-02) | -1.59658 | #N/A | #N/A |
| OS05G0478000 | Zinc finger, RING/FYVE/PHD-type domain containing protein. (Os05t0478000-01) | -1.59652 | #N/A | #N/A |
| OS04G0462300 | Conserved hypothetical protein. (Os04t0462300-01) | -1.59594 | #N/A | #N/A |
| OS02G0562600 | Similar to Seven trans-membrane protein (Fragment). (Os02t0562600-01) | -1.59528 | #N/A | -1.82641 |
| OS03G0736600 | Similar to ATP synthase. (Os03t0736600-01) | -1.59452 | -0.677925 | #N/A |
| OS05G0143600 | Similar to Jasmonate-induced protein. (Os05t0143600-01) | -1.59352 | #N/A | #N/A |
| OS02G0792000 | Hypothetical protein. (Os02t0792000-01) | -1.59346 | #N/A | #N/A |
| OS07G0174400 | Similar to Non-specific lipid-transfer protein. (Os07t0174400-00) | -1.59309 | #N/A | #N/A |

| | | | | |
|---|---|---|---|---|
| OS07G0417200 | Similar to Delta 12 oleic acid desaturase FAD2. (Os07t0417200-01) | -1.58926 | -4.11333 | -3.363928 |
| OS09G0299400 | Similar to TPK1. (Os09t0299400-00) | -1.58877 | #N/A | #N/A |
| OS06G0108900 | Similar to H0613A10.3 protein. (Os06t0108900-01);Homeodomain-related domain containing protein. (Os06t0108900-02) | -1.58722 | #N/A | #N/A |
| OS06G0291800 | Conserved hypothetical protein. (Os06t0291800-00) | -1.58627 | #N/A | #N/A |
| OS10G0155400 | Similar to Amidase family protein, expressed. (Os10t0155400-00) | -1.58583 | #N/A | #N/A |
| OS04G0628200 | Similar to H0303G06.9 protein. (Os04t0628200-01) | -1.58476 | #N/A | #N/A |
| OS08G0511400 | Conserved hypothetical protein. (Os08t0511400-01) | -1.58442 | #N/A | #N/A |
| OS06G0141200 | Similar to RNA-binding protein EWS. (Os06t0141200-01) | -1.58208 | #N/A | #N/A |
| OS03G0290300 | Similar to W-3 fatty acid desaturase (Fragment). (Os03t0290300-01) | -1.58207 | #N/A | #N/A |
| OS06G0505700 | Similar to AGP16. (Os06t0505700-01) | -1.57796 | #N/A | #N/A |
| OS11G0649500 | Uncharacterised protein family UPF0497, trans-membrane plant domain containing protein. (Os11t0649500-00) | -1.57013 | #N/A | #N/A |
| OS09G0555150 | Sulfotransferase family protein. (Os09t0555150-01) | -1.56905 | #N/A | #N/A |
| OS03G0279600 | Conserved hypothetical protein. (Os03t0279600-01);Conserved hypothetical protein. (Os03t0279600-02) | -1.56886 | -0.859947 | #N/A |
| OS11G0148600 | Conserved hypothetical protein. (Os11t0148600-01) | -1.56822 | #N/A | #N/A |
| OS02G0718000 | Conserved hypothetical protein. (Os02t0718000-01) | -1.56722 | #N/A | #N/A |
| OS10G0116400 | Hypothetical conserved gene. (Os10t0116400-01) | -1.56678 | #N/A | #N/A |
| OS07G0119000 | Similar to MAP3K gamma protein kinase (Fragment). (Os07t0119000-01) | -1.56633 | #N/A | #N/A |
| OS06G0140700 | Similar to Homeodomain leucine zipper protein (Fragment). (Os06t0140700-01) | -1.56501 | #N/A | #N/A |
| OS04G0298600 | Uncharacterised protein family FPL domain containing protein. (Os04t0298600-00) | -1.56465 | -1.29196 | -0.797827 |
| OS02G0106600 | Zinc finger, RING/FYVE/PHD-type domain containing protein. (Os02t0106600-01) | -1.56406 | #N/A | #N/A |
| OS03G0300000 | Similar to Xyloglucan 6-xylosyltransferase (EC 2.4.2.39) (AtXT1). (Os03t0300000-01);Similar to glycosyltransferase 5. (Os03t0300000 | -1.56342 | -1.01478 | #N/A |
| OS07G0489300 | UDP-glucuronosyl/UDP-glucosyltransferase family protein. (Os07t0489300-01) | -1.56337 | #N/A | #N/A |
| OS01G0938400 | Conserved hypothetical protein. (Os01t0938400-01) | -1.56208 | #N/A | #N/A |
| OS01G0722700 | Similar to Hexokinase. (Os01t0722700-01) | -1.55656 | #N/A | #N/A |
| OS03G0807000 | Conserved hypothetical protein. (Os03t0807000-01) | -1.55628 | #N/A | #N/A |
| OS10G0413101 | Similar to H0815C01.6 protein. (Os10t0413101-01) | -1.55622 | #N/A | #N/A |
| OS05G0110100 | Heat shock protein DnaJ, cysteine-rich domain domain containing protein. (Os05t0110100-01);Heat shock protein DnaJ, cysteine-ric | -1.55286 | -0.993482 | #N/A |
| OS07G0614600 | Transmembrane receptor, eukaryota domain containing protein. (Os07t0614600-01) | -1.55201 | -2.48989 | -2.29112 |
| OS01G0892500 | Similar to carboxylic ester hydrolase. (Os01t0892500-01) | -1.55186 | #N/A | -1.92985 |
| OS07G0663700 | Similar to Oxidoreductase, short chain dehydrogenase/reductase family protein, expressed. (Os07t0663700-01) | -1.55181 | -2.53788 | -4.91791 |
| OS07G0498400 | Leucine-rich repeat, typical subtype domain containing protein. (Os07t0498400-00) | -1.54657 | #N/A | #N/A |
| OS02G0571100 | Terpenoid synthase domain containing protein. (Os02t0571100-01) | -1.54607 | #N/A | #N/A |
| OS01G0332200 | GA 2-oxidase2, GA metabolism (Os01t0332200-01) | -1.54585 | #N/A | #N/A |
| OS01G0795000 | Subtilase. (Os01t0795000-01) | -1.5445 | -2.53613 | -4.03328 |
| OS04G0556550 | Conserved hypothetical protein. (Os04t0556550-00) | -1.54423 | #N/A | #N/A |
| OS08G0513000 | Similar to Serine/threonine protein phosphatase. (Os08t0513000-01) | -1.54204 | #N/A | #N/A |
| OS12G0578000 | Conserved hypothetical protein. (Os12t0578000-01) | -1.54197 | -0.972193 | #N/A |
| OS03G0776150 | Hypothetical protein. (Os03t0776150-00) | -1.54049 | #N/A | #N/A |
| OS09G0540600 | Similar to WD-40 repeat protein MSI1. (Os09t0540600-01) | -1.53963 | #N/A | #N/A |
| OS02G0793500 | Cyclin-like F-box domain containing protein. (Os02t0793500-01) | -1.53869 | #N/A | #N/A |
| OS05G0322900 | WRKY transcription factor, Benzothiadiazole (BTH)-inducible blast resistance (Os05t0322900-01);Similar to WRKY transcription facto | -1.53846 | -1.66278 | -1.30225 |
| OS06G0196600 | Beta-ketoacyl-[acyl carrier protein] synthase I, Root development (Os06t0196600-01) | -1.5351 | #N/A | #N/A |

| | | | | |
|---|---|---|---|---|
| OS10G0195600 | Transferase family protein. (Os10t0195600-01) | -1.53482 | #N/A | #N/A |
| OS10G0478100 | Heat shock protein DnaJ, N-terminal domain containing protein. (Os10t0478100-01);Heat shock protein DnaJ, N-terminal domain co | -1.53466 | #N/A | #N/A |
| OS03G0232500 | GTP-binding protein, HSR1-related domain containing protein. (Os03t0232500-01) | -1.53223 | #N/A | #N/A |
| OS11G0488500 | 2OG-Fe(II) oxygenase domain containing protein. (Os11t0488500-01);Similar to oxidoreductase, 2OG-Fe(II) oxygenase family protei | -1.5256 | #N/A | #N/A |
| OS07G0542400 | Similar to Receptor protein kinase. (Os07t0542400-01);Similar to receptor-like protein kinase RK20-1. (Os07t0542400-02) | -1.52473 | #N/A | #N/A |
| OS01G0743400 | Similar to Tryptophanyl-tRNA synthetase (Fragment). (Os01t0743400-01);Similar to predicted protein. (Os01t0743400-02) | -1.52171 | #N/A | #N/A |
| OS10G0522000 | Similar to Methyltransferase family protein, expressed. (Os10t0522000-01) | -1.51999 | #N/A | #N/A |
| OS08G0140300 | Aromatic L-amino acid decarboxylase (AADC), Senescence-induced serotonin biosynthesis (Os08t0140300-01) | -1.51986 | -1.79313 | -4.09093 |
| OS12G0555400 | Similar to Raffinose synthase protein. (Os12t0555400-01) | -1.51929 | #N/A | #N/A |
| OS08G0521800 | Protein of unknown function DUF1997 domain containing protein. (Os08t0521800-01);Protein of unknown function DUF1997 doma | -1.51778 | -0.647554 | #N/A |
| OS08G0557100 | Similar to Nucleic acid-binding protein precursor. (Os08t0557100-01);28 kDa ribonucleoprotein, chloroplast precursor (28RNP). (Os | -1.51714 | #N/A | #N/A |
| OS11G0566800 | Similar to Bibenzyl synthase (EC 2.3.1.-). (Os11t0566800-01) | -1.51644 | #N/A | -1.64164 |
| OS07G0664000 | Similar to Oxidoreductase, short chain dehydrogenase/reductase family protein, expressed. (Os07t0664000-00) | -1.51364 | #N/A | #N/A |
| OS08G0557400 | Protein tyrosine phosphatase-like protein. (Os08t0557400-01) | -1.51337 | #N/A | #N/A |
| OS01G0805300 | Mog1/PsbP, alpha/beta/alpha sandwich domain containing protein. (Os01t0805300-01);Mog1/PsbP, alpha/beta/alpha sandwich do | -1.51323 | #N/A | #N/A |
| OS10G0573400 | Similar to Cyclase/dehydrase family protein. (Os10t0573400-01) | -1.50807 | #N/A | #N/A |
| OS02G0302700 | Similar to Nicotianamine aminotransferase A. (Os02t0302700-01) | -1.50542 | -0.952016 | #N/A |
| OS04G0111200 | Similar to ATP sulfurylase (Fragment). (Os04t0111200-01) | -1.50487 | #N/A | #N/A |
| OS07G0643400 | Alpha/beta hydrolase fold-3 domain containing protein. (Os07t0643400-01) | -1.50337 | -0.642578 | #N/A |
| OS01G0882500 | Similar to NADH dehydrogenase I subunit N. (Os01t0882500-00) | -1.50157 | #N/A | #N/A |
| OS04G0689000 | Similar to Peroxidase (EC 1.11.1.7). (Os04t0689000-01) | 1.50009 | #N/A | #N/A |
| OS11G0296500 | Zinc finger, CCHC retroviral-type domain containing protein. (Os11t0296500-01) | 1.5021 | #N/A | 1.53363 |
| OS11G0665600 | Helix-turn-helix, Fis-type domain containing protein. (Os11t0665600-00) | 1.50234 | #N/A | #N/A |
| OS07G0418600 | Proline-rich glycoprotein, ABA-dependent inhibition of root growth (Os07t0418600-01) | 1.50361 | #N/A | #N/A |
| OS12G0132800 | Major facilitator superfamily antiporter. (Os12t0132800-01) | 1.50418 | #N/A | #N/A |
| OS03G0750300 | Conserved hypothetical protein. (Os03t0750300-00) | 1.50426 | #N/A | #N/A |
| OS04G0377600 | Similar to OSIGBa0135C09.4 protein. (Os04t0377600-01) | 1.5061 | #N/A | #N/A |
| OS03G0401100 | Similar to Protein kinase domain containing protein, expressed. (Os03t0401100-01) | 1.50656 | #N/A | #N/A |
| OS10G0544200 | Basic helix-loop-helix dimerisation region bHLH domain containing protein. (Os10t0544200-01) | 1.5107 | #N/A | #N/A |
| OS08G0501900 | Non-protein coding transcript. (Os08t0501900-01) | 1.51206 | #N/A | #N/A |
| OS11G0244300 | Similar to ATPase, AAA family protein. (Os11t0244300-01) | 1.51295 | #N/A | #N/A |
| OS08G0556200 | Similar to Dihydroneopterin aldolase. (Os08t0556200-01) | 1.51341 | #N/A | #N/A |
| OS06G0647700 | Hypothetical protein. (Os06t0647700-01) | 1.51348 | #N/A | #N/A |
| OS03G0277600 | Similar to (clone wusl1032) mRNA sequence. (Os03t0277600-01) | 1.51447 | #N/A | #N/A |
| OS09G0481100 | Similar to Phosphatidylinositol transfer-like protein IV. (Os09t0481100-01) | 1.51547 | #N/A | #N/A |
| OS10G0546400 | Conserved hypothetical protein. (Os10t0546400-01) | 1.51575 | #N/A | #N/A |
| OS10G0114300 | Similar to NADPH-dependent codeinone reductase (EC 1.1.1.247). (Os10t0114300-01) | 1.5167 | #N/A | #N/A |
| OS07G0123500 | BTB/POZ fold domain containing protein. (Os07t0123500-01) | 1.517 | #N/A | #N/A |
| OS04G0571300 | Cyclin-like F-box domain containing protein. (Os04t0571300-01);Similar to OSIGBa0127A14.1 protein. (Os04t0571300-02);Cyclin-like | 1.51775 | #N/A | #N/A |
| OS03G0324900 | Zinc finger, RING/FYVE/PHD-type domain containing protein. (Os03t0324900-01) | 1.51785 | #N/A | #N/A |
| OS02G0584900 | Conserved hypothetical protein. (Os02t0584900-01) | 1.51883 | #N/A | #N/A |
| OS05G0375400 | Beta-glucanase precursor. (Os05t0375400-01) | 1.51927 | #N/A | 0.988123 |

| | | | | |
|---|---|---|---|---|
| OS02G0504100 | Conserved hypothetical protein. (Os02t0504100-01) | 1.51983 | #N/A | #N/A |
| OS03G0793600 | Similar to type I inositol-1,4,5-trisphosphate 5-phosphatase CVP2. (Os03t0793600-00) | 1.52032 | #N/A | #N/A |
| OS05G0509500 | Similar to Signal recognition particle 54 kDa protein 2 (SRP54). (Os05t0509500-01) | 1.52224 | #N/A | #N/A |
| OS10G0358600 | Similar to OSIGBa0136O08-OSIGBa0153H12.4 protein. (Os10t0358600-01) | 1.52249 | #N/A | #N/A |
| OS03G0148800 | Zinc finger, C2H2-type domain containing protein. (Os03t0148800-01) | 1.52296 | #N/A | #N/A |
| OS11G0601700 | Helix-loop-helix DNA-binding domain containing protein. (Os11t0601700-01) | 1.52317 | #N/A | #N/A |
| OS04G0509300 | Rnase III family protein, Shoot apical meristem (SAM) formation and maintenance (Os04t0509300-01);Rnase III family protein, Tran | 1.52436 | #N/A | #N/A |
| OS08G0485800 | Barwin-related endoglucanase domain containing protein. (Os08t0485800-01) | 1.52503 | #N/A | #N/A |
| OS02G0213100 | Conserved hypothetical protein. (Os02t0213100-01) | 1.52609 | #N/A | 1.45253 |
| OS01G0299100 | Conserved hypothetical protein. (Os01t0299100-00) | 1.52800113 | -0.6684209 | -0.5042046 |
| OS01G0951400 | Similar to Uridine 5'-monophosphate synthase (UMP synthase). (Os01t0951400-01);Similar to UMP synthase. (Os01t0951400-02) | 1.53092 | #N/A | #N/A |
| OS06G0581500 | Protein kinase, core domain containing protein. (Os06t0581500-01) | 1.53473 | -1.78138 | #N/A |
| OS01G0954500 | Protein of unknown function DUF702 family protein. (Os01t0954500-01) | 1.53779 | #N/A | #N/A |
| OS04G0674500 | Hypothetical protein. (Os04t0674500-01) | 1.54347 | #N/A | #N/A |
| OS05G0380200 | Conserved hypothetical protein. (Os05t0380200-01) | 1.54475 | #N/A | #N/A |
| OS02G0134300 | Protein of unknown function DUF701, zinc-binding putative family protein. (Os02t0134300-01) | 1.54607 | #N/A | #N/A |
| OS07G0679300 | Similar to Alpha-galactosidase precursor (EC 3.2.1.22) (Melibiase) (Alpha-D- galactoside galactohydrolase). (Os07t0679300-01) | 1.54787 | #N/A | #N/A |
| OS01G0920100 | Conserved hypothetical protein. (Os01t0920100-01) | 1.54838 | #N/A | #N/A |
| OS02G0742800 | Conserved hypothetical protein. (Os02t0742800-01) | 1.54841 | #N/A | #N/A |
| OS10G0505700 | Similar to Nonspecific lipid-transfer protein 2 (nsLTP2) (7 kDa lipid transfer protein). (Os10t0505700-01) | 1.54843 | #N/A | #N/A |
| OS03G0265600 | Similar to Transformer-2-like protein. (Os03t0265600-01) | 1.54921 | #N/A | #N/A |
| OS09G0491100 | Similar to Beta-primeverosidase (EC 3.2.1.149). (Os09t0491100-01);Similar to Beta-glucosidase isozyme 2. (Os09t0491100-02);Simil | 1.55046 | #N/A | #N/A |
| OS01G0706600 | Conserved hypothetical protein. (Os01t0706600-00) | 1.5505 | #N/A | #N/A |
| OS02G0804000 | Non-protein coding transcript. (Os02t0804000-01);Non-protein coding transcript. (Os02t0804000-02) | 1.55279 | #N/A | 2.10536 |
| OS12G0572500 | Region of unknown function XH domain containing protein. (Os12t0572500-01) | 1.55623 | #N/A | #N/A |
| OS08G0190700 | Conserved hypothetical protein. (Os08t0190700-00) | 1.55901 | #N/A | #N/A |
| OS08G0412100 | NB-ARC domain containing protein. (Os08t0412100-01) | 1.56002 | #N/A | #N/A |
| OS07G0592600 | Indole-3-acetic acid (IAA)-amido synthetase, Auxin signaling and defense signaling in a pathogen-nonspecific manner (Os07t059260 | 1.56295 | #N/A | #N/A |
| OS11G0120300 | Protein of unknown function DUF567 family protein. (Os11t0120300-01) | 1.56384 | #N/A | #N/A |
| OS02G0558600 | Conserved hypothetical protein. (Os02t0558600-01) | 1.56455 | #N/A | #N/A |
| OS10G0145200 | Cyclin-like F-box domain containing protein. (Os10t0145200-01) | 1.56522 | #N/A | #N/A |
| OS10G0544600 | Zinc finger, RING/FYVE/PHD-type domain containing protein. (Os10t0544600-01);Zinc finger, RING/FYVE/PHD-type domain containi | 1.57192 | #N/A | #N/A |
| OS02G0671100 | Cyclin-like F-box domain containing protein. (Os02t0671100-01) | 1.57341 | #N/A | #N/A |
| OS11G0652100 | NB-ARC domain containing protein. (Os11t0652100-01) | 1.57351 | #N/A | #N/A |
| OS03G0590700 | NADH dehydrogenase [ubiquinone] 1 alpha subcomplex assembly factor 3 domain containing protein. (Os03t0590700-01) | 1.57453 | #N/A | #N/A |
| OS07G0130200 | Similar to Resistance protein candidate (Fragment). (Os07t0130200-01) | 1.57488 | #N/A | #N/A |
| OS03G0225200 | Cyclin, A/B/D/E domain containing protein. (Os03t0225200-01);Similar to cyclin, N-terminal domain containing protein. (Os03t0225 | 1.57546 | #N/A | #N/A |
| OS03G0229000 | Conserved hypothetical protein. (Os03t0229000-01) | 1.58086 | #N/A | #N/A |
| OS06G0354700 | Alpha/beta hydrolase-fold family protein, Chlorophyll degradation during senescence (Os06t0354700-01);Similar to Hydrolase, alph | 1.58122 | #N/A | #N/A |
| OS04G0394766 | Non-protein coding transcript. (Os04t0394766-00) | 1.58494 | #N/A | #N/A |
| OS06G0673500 | Similar to polyubiquitin containing 7 ubiquitin monomers. (Os06t0673500-01) | 1.58575 | #N/A | #N/A |
| OS11G0289700 | Cytochrome P450 family protein. (Os11t0289700-01) | 1.58696 | #N/A | #N/A |

| | | | | |
|---|---|---|---|---|
| OS07G0601100 | Similar to NADPH HC toxin reductase (Fragment). (Os07t0601100-01) | 1.58824 | #N/A | #N/A |
| OS12G0147800 | Similar to Phytosulfokines 5 precursor (Secretory protein SH27A). (Os12t0147800-01) | 1.5893 | #N/A | #N/A |
| OS01G0609300 | Similar to Pleiotropic drug resistance protein 3. (Os01t0609300-01);Hypothetical conserved gene. (Os01t0609300-02) | 1.58971 | #N/A | #N/A |
| OS06G0672801 | Conserved hypothetical protein. (Os06t0672801-01) | 1.59006 | #N/A | #N/A |
| OS04G0540500 | Hypothetical protein. (Os04t0540500-01) | 1.59063 | #N/A | #N/A |
| OS12G0131200 | Similar to zinc finger (DHHC type) family protein. (Os12t0131200-00) | 1.59407 | #N/A | #N/A |
| OS01G0811300 | Similar to SET domain protein SDG111. (Os01t0811300-01) | 1.59423 | #N/A | #N/A |
| OS01G0129800 | Conserved hypothetical protein. (Os01t0129800-01) | 1.5960287 | #N/A | #N/A |
| OS10G0135500 | Cyclin-like F-box domain containing protein. (Os10t0135500-01) | 1.59747 | #N/A | #N/A |
| OS02G0533800 | Similar to ATPase inhibitor. (Os02t0533800-01) | 1.59778 | #N/A | #N/A |
| OS02G0582300 | Pentatricopeptide repeat domain containing protein. (Os02t0582300-01) | 1.59818 | #N/A | #N/A |
| OS08G0120301 | Hypothetical gene. (Os08t0120301-00) | 1.60222 | #N/A | #N/A |
| OS06G0585900 | Similar to cDNA clone:002-174-F06, full insert sequence. (Os06t0585900-01);Hypothetical protein. (Os06t0585900-02) | 1.60624 | #N/A | #N/A |
| OS11G0633800 | Cyclin-like F-box domain containing protein. (Os11t0633800-01) | 1.60688 | #N/A | #N/A |
| OS05G0529700 | Similar to electron transporter/ heat shock protein binding protein. (Os05t0529700-01);Similar to electron transporter/ heat shock | 1.60775 | #N/A | #N/A |
| OS10G0139600 | F-box domain, Skp2-like domain containing protein. (Os10t0139600-01) | 1.60938 | #N/A | #N/A |
| OS05G0531100 | Protein of unknown function DUF584 family protein. (Os05t0531100-01) | 1.60987 | #N/A | #N/A |
| OS12G0270900 | Sulfotransferase family protein. (Os12t0270900-01);Similar to cDNA clone:J023121F12, full insert sequence. (Os12t0270900-02) | 1.61021 | #N/A | #N/A |
| OS08G0524100 | Similar to Type II inositol-1,4,5-trisphosphatase 5-phosphatase 12 (EC 3.1.3.36) (At5PTase12) (FRAGILE FIBER3 protein). (Os08t052410 | 1.61251 | #N/A | #N/A |
| OS01G0581900 | Conserved hypothetical protein. (Os01t0581900-01) | 1.61324 | #N/A | #N/A |
| OS04G0555600 | Similar to WD-40 repeat family protein / beige-related. (Os04t0555600-01) | 1.61563 | #N/A | #N/A |
| OS08G0128200 | Protein of unknown function DUF3537 domain containing protein. (Os08t0128200-01) | 1.6195 | #N/A | #N/A |
| OS06G0701700 | Ion transporter , Na+/K+ symport (Os06t0701700-01);Similar to Cation transporter HKT1. (Os06t0701700-02) | 1.6224 | #N/A | #N/A |
| OS08G0420700 | Conserved hypothetical protein. (Os08t0420700-01) | 1.62322 | #N/A | #N/A |
| OS09G0464000 | Similar to Carbonate dehydratase-like protein. (Os09t0464000-01);Similar to Carbonate dehydratase-like protein. (Os09t0464000-0 | 1.62411 | #N/A | #N/A |
| OS03G0334200 | Protein of unknown function DUF3133 domain containing protein. (Os03t0334200-01);Protein of unknown function DUF3133 doma | 1.62493 | #N/A | #N/A |
| OS10G0131800 | Similar to NBS-LRR class disease resistance protein. (Os10t0131800-01) | 1.62563 | #N/A | #N/A |
| OS02G0153200 | Protein kinase, core domain containing protein. (Os02t0153200-01) | 1.62709 | #N/A | 1.19811 |
| OS08G0439000 | Similar to pyrophosphate--fructose 6-phosphate 1-phosphotransferase. (Os08t0439000-01) | 1.62982 | #N/A | #N/A |
| OS11G0661101 | Conserved hypothetical protein. (Os11t0661101-01) | 1.63223 | #N/A | #N/A |
| OS01G0247500 | Protein kinase, core domain containing protein. (Os01t0247500-01) | 1.63291 | #N/A | #N/A |
| OS01G0823600 | Conserved hypothetical protein. (Os01t0823600-01) | 1.63416 | #N/A | #N/A |
| OS03G0321700 | Similar to WRKY transcription factor 55. (Os03t0321700-01) | 1.63485 | #N/A | #N/A |
| OS09G0241100 | Similar to nucleotide binding. (Os09t0241100-01) | 1.63666 | #N/A | #N/A |
| OS06G0691800 | Protein kinase, core domain containing protein. (Os06t0691800-01) | 1.63828 | #N/A | #N/A |
| OS12G0503000 | Similar to Allantoin permease. (Os12t0503000-01);Similar to Ureide permease 4. (Os12t0503000-03);Similar to Ureide permease 4. | 1.63874 | #N/A | #N/A |
| OS12G0477700 | Conserved hypothetical protein. (Os12t0477700-01) | 1.64146 | #N/A | #N/A |
| OS01G0392800 | Similar to DET1-like protein. (Os01t0392800-01) | 1.64322 | #N/A | 1.59081 |
| OS01G0115800 | Similar to Receptor-like kinase. (Os01t0115800-01) | 1.64684 | #N/A | #N/A |
| OS07G0180000 | Hypothetical protein. (Os07t0180000-01) | 1.64707 | #N/A | #N/A |
| OS08G0208700 | Zinc finger, BED-type predicted domain containing protein. (Os08t0208700-01) | 1.64842 | #N/A | #N/A |
| OS02G0153300 | Hypothetical protein. (Os02t0153300-01);Hypothetical protein. (Os02t0153300-02) | 1.65177 | #N/A | #N/A |

| | | | | |
|---|---|---|---|---|
| OS02G0104150 | Non-protein coding transcript. (Os02t0104150-00) | 1.65303 | #N/A | #N/A |
| OS02G0802500 | Similar to H(+)-translocating (Pyrophosphate-ENERGIZED) inorganic pyrophosphatase beta-1 polypeptide (EC 3.6.1.1) (Fragment). (O | 1.65889 | #N/A | #N/A |
| OS03G0153500 | Similar to FAD binding domain containing protein, expressed. (Os03t0153500-00) | 1.65915 | #N/A | #N/A |
| OS08G0411500 | Conserved hypothetical protein. (Os08t0411500-01) | 1.6605 | #N/A | #N/A |
| OS07G0272400 | Zinc finger, RING/FYVE/PHD-type domain containing protein. (Os07t0272400-01) | 1.66305 | #N/A | #N/A |
| OS03G0577100 | Nuf2 family protein. (Os03t0577100-01) | 1.66794 | #N/A | #N/A |
| OS01G0555200 | Asp/Glu racemase family protein. (Os01t0555200-01) | 1.66916 | #N/A | #N/A |
| OS02G0583700 | Conserved hypothetical protein. (Os02t0583700-00) | 1.66948944 | #N/A | #N/A |
| OS05G0594200 | Similar to Cation/proton exchanger 1a. (Os05t0594200-01) | 1.67013 | #N/A | #N/A |
| OS01G0178100 | Region of unknown function DUF1767 domain containing protein. (Os01t0178100-01) | 1.67044 | #N/A | 1.15327 |
| OS06G0138000 | Similar to IMB1. (Os06t0138000-01) | 1.67535 | #N/A | #N/A |
| OS06G0716100 | Similar to Chaperone protein dnaJ. (Os06t0716100-01) | 1.67723 | #N/A | #N/A |
| OS07G0601900 | Similar to NADPH HC toxin reductase (Fragment). (Os07t0601900-01) | 1.67838 | -1.70105 | -2.25909 |
| OS02G0731500 | Similar to cDNA, clone: J065098J18, full insert sequence. (Os02t0731500-01);AWPM-19-like family protein. (Os02t0731500-02) | 1.68095 | #N/A | #N/A |
| OS01G0162500 | Leucine-rich repeat-containing N-terminal, type 2 domain containing protein. (Os01t0162500-01) | 1.68165 | #N/A | #N/A |
| OS07G0538400 | Similar to Receptor-like protein kinase 4. (Os07t0538400-01) | 1.68201 | #N/A | #N/A |
| OS12G0602200 | Conserved hypothetical protein. (Os12t0602200-01) | 1.68452 | #N/A | #N/A |
| OS04G0629600 | Conserved hypothetical protein. (Os04t0629600-01);Similar to Transposase of Tn10 [Oryza sativa (japonica cultivar-group)]. (Os04t0 | 1.68559 | #N/A | #N/A |
| OS10G0416800 | Similar to Chitinase 2 (EC 3.2.1.14) (Tulip bulb chitinase-2) (TBC-2). (Os10t0416800-03) | 1.68667 | #N/A | #N/A |
| OS12G0527700 | Serine/threonine protein kinase-related domain containing protein. (Os12t0527700-01) | 1.6869 | #N/A | #N/A |
| OS11G0618850 | Hypothetical conserved gene. (Os11t0618850-00) | 1.69027 | #N/A | #N/A |
| OS01G0793800 | Conserved hypothetical protein. (Os01t0793800-01) | 1.69134 | #N/A | #N/A |
| OS03G0599000 | NB-ARC domain containing protein. (Os03t0599000-01) | 1.69994 | #N/A | #N/A |
| OS03G0596600 | Zinc finger, CCHC-type domain containing protein. (Os03t0596600-01) | 1.70243 | #N/A | #N/A |
| OS01G0797900 | Hypothetical protein. (Os01t0797900-01);Hypothetical protein. (Os01t0797900-02) | 1.70437 | #N/A | #N/A |
| OS08G0278100 | Osteocrin domain containing protein. (Os08t0278100-01) | 1.71171 | #N/A | #N/A |
| OS09G0356200 | Malectin-like carbohydrate-binding domain domain containing protein. (Os09t0356200-01) | 1.71192 | #N/A | #N/A |
| OS02G0284500 | FAR1 DNA binding domain domain containing protein. (Os02t0284500-01) | 1.71441 | #N/A | #N/A |
| OS05G0456900 | Conserved hypothetical protein. (Os05t0456900-01) | 1.71498 | #N/A | #N/A |
| OS01G0826000 | Heavy metal transport/detoxification protein domain containing protein. (Os01t0826000-00) | 1.715 | #N/A | #N/A |
| OS02G0740400 | Similar to GSDL-motif lipase. (Os02t0740400-01);Lipase, GDSL domain containing protein. (Os02t0740400-02) | 1.71624 | #N/A | #N/A |
| OS04G0644675 | Similar to H0413E07.9 protein. (Os04t0644675-00) | 1.71883 | #N/A | #N/A |
| OS10G0517050 | Hypothetical conserved gene. (Os10t0517050-01) | 1.72383 | #N/A | #N/A |
| OS11G0648400 | Protein of unknown function DUF3615 domain containing protein. (Os11t0648400-01) | 1.72483 | #N/A | #N/A |
| OS05G0209000 | Similar to Cytidine deaminase. (Os05t0209000-01) | 1.72513 | #N/A | #N/A |
| OS03G0348900 | Similar to CHY zinc finger family protein, expressed. (Os03t0348900-01);Ring finger E3 ubiquitin ligase, Abiotic stress tolerance (Os0 | 1.72527 | #N/A | #N/A |
| OS01G0838400 | Conserved hypothetical protein. (Os01t0838400-01) | 1.72534 | #N/A | #N/A |
| OS07G0604400 | Similar to COBRA-like protein 6. (Os07t0604400-00) | 1.72861 | #N/A | #N/A |
| OS02G0141300 | Arabinokinase-Like Protein, Pollen development (Os02t0141300-01) | 1.72933 | #N/A | #N/A |
| OS11G0212900 | Serine/threonine protein kinase-related domain containing protein. (Os11t0212900-01) | 1.73207 | #N/A | #N/A |
| OS02G0104100 | Similar to MADS-box transcription factor TaAGL18. (Os02t0104100-00) | 1.73225 | #N/A | #N/A |
| OS02G0153700 | Serine/threonine protein kinase domain containing protein. (Os02t0153700-01) | 1.73351 | #N/A | #N/A |

| | | | | |
|---|---|---|---|---|
| OS03G0728900 | Helix-loop-helix DNA-binding domain containing protein. (Os03t0728900-00) | 1.73789 | #N/A | #N/A |
| OS12G0606600 | DNA-binding HORMA domain containing protein. (Os12t0606600-01) | 1.74453 | #N/A | #N/A |
| OS01G0132000 | Similar to Wound-induced protease inhibitor (WIP1). (Os01t0132000-01) | 1.75045 | #N/A | #N/A |
| OS02G0751100 | Peptidase A1 domain containing protein. (Os02t0751100-01);Hypothetical conserved gene. (Os02t0751100-02) | 1.75274 | #N/A | #N/A |
| OS01G0976300 | Heavy metal transport/detoxification protein domain containing protein. (Os01t0976300-01) | 1.75524 | #N/A | #N/A |
| OS05G0172100 | F-box associated type 1 domain containing protein. (Os05t0172100-01) | 1.75826 | #N/A | #N/A |
| OS03G0288900 | Hypothetical protein. (Os03t0288900-01) | 1.7595 | #N/A | #N/A |
| OS11G0513700 | Protein kinase-like domain containing protein. (Os11t0513700-01) | 1.75958 | #N/A | #N/A |
| OS03G0231950 | Similar to Helix-loop-helix DNA-binding domain containing protein, expressed. (Os03t0231950-01) | 1.75991 | #N/A | #N/A |
| OS04G0649200 | Similar to H0212B02.1 protein. (Os04t0649200-01) | 1.76007 | #N/A | #N/A |
| OS01G0114100 | Similar to Protein kinase RLK17. (Os01t0114100-01) | 1.76153 | #N/A | #N/A |
| OS06G0649600 | Non-protein coding transcript. (Os06t0649600-01) | 1.7775 | #N/A | #N/A |
| OS01G0661500 | Mov34/MPN/PAD-1 family protein. (Os01t0661500-01) | 1.77941 | #N/A | #N/A |
| OS01G0283000 | Conserved hypothetical protein. (Os01t0283000-01);Conserved hypothetical protein. (Os01t0283000-02) | 1.78067 | #N/A | #N/A |
| OS03G0235000 | Similar to Peroxidase. (Os03t0235000-01) | 1.78513 | #N/A | #N/A |
| OS12G0610600 | Similar to NAM / CUC2-like protein. (Os12t0610600-01) | 1.7856 | #N/A | #N/A |
| OS12G0604700 | Similar to LSTK-1-like kinase. (Os12t0604700-01);Similar to cDNA clone:J023075D08, full insert sequence. (Os12t0604700-02) | 1.78684 | #N/A | #N/A |
| OS12G0148700 | Adipose-regulatory protein, Seipin domain containing protein. (Os12t0148700-01) | 1.79054 | #N/A | #N/A |
| OS11G0113700 | Serine/threonine protein kinase, Abiotic stresses (Os11t0113700-01);Serine/threonine protein kinase, Microbe-assosiated molecula | 1.79098 | #N/A | #N/A |
| OS01G0927400 | Molecular chaperone, heat shock protein, Hsp40, DnaJ domain containing protein. (Os01t0927400-01) | 1.79126 | #N/A | #N/A |
| OS01G0668400 | Similar to predicted protein. (Os01t0668400-00) | 1.79387 | #N/A | #N/A |
| OS05G0369300 | Exo70 exocyst complex subunit domain containing protein. (Os05t0369300-01) | 1.79723 | #N/A | #N/A |
| OS03G0834050 | Similar to predicted protein. (Os03t0834050-01) | 1.80111 | #N/A | #N/A |
| OS03G0689300 | Plasma membrane H+ ATPase (EC 3.6.3.6) (H-ATPase). (Os03t0689300-01);Similar to H-ATPase. (Os03t0689300-02) | 1.8033 | #N/A | #N/A |
| OS01G0147300 | Similar to membrane protein. (Os01t0147300-01) | 1.80438 | #N/A | #N/A |
| OS06G0295900 | Hypothetical conserved gene. (Os06t0295900-01) | 1.80531 | #N/A | 1.36065 |
| OS02G0282900 | Similar to 68 kDa protein HP68. (Os02t0282900-01) | 1.80701 | #N/A | #N/A |
| OS11G0132600 | Similar to Arabinoxylan arabinofuranohydrolase isoenzyme AXAH-II. (Os11t0132600-01);Similar to Arabinoxylan arabinofuranohydr | 1.80748 | #N/A | #N/A |
| OS02G0554900 | Similar to Protein disulfide isomerase (Fragment). (Os02t0554900-01);Similar to Protein disulfide isomerase-like 1-3. (Os02t055490 | 1.80953 | #N/A | #N/A |
| OS12G0441300 | O-methyltransferase, family 2 protein. (Os12t0441300-00) | 1.81071044 | -0.6444153 | -0.3638575 |
| OS07G0285200 | Cyclin-like F-box domain containing protein. (Os07t0285200-01) | 1.81926 | #N/A | #N/A |
| OS04G0685500 | Exo70 exocyst complex subunit domain containing protein. (Os04t0685500-01);Hypothetical conserved gene. (Os04t0685500-02) | 1.82131 | #N/A | #N/A |
| OS03G0125100 | Beta-carotene hydroxylase, Drought and oxidative stress resistance (Os03t0125100-01) | 1.82301 | #N/A | #N/A |
| OS04G0310400 | Similar to H0211A12.15 protein. (Os04t0310400-00) | 1.8245 | #N/A | #N/A |
| OS05G0247800 | Glycoside hydrolase, family 18 protein. (Os05t0247800-01) | 1.82637154 | #N/A | #N/A |
| OS09G0268100 | Similar to lectin-like receptor kinase 7. (Os09t0268100-00) | 1.82911 | #N/A | #N/A |
| OS03G0277500 | Similar to Glyoxalase family protein, expressed. (Os03t0277500-01) | 1.83275232 | #N/A | #N/A |
| OS12G0105500 | Similar to Phosphatidate cytidylyltransferase family protein, expressed. (Os12t0105500-01) | 1.83572 | #N/A | #N/A |
| OS05G0580000 | Similar to ADP-glucose pyrophosphorylase (EC 2.7.7.27) (Fragment). (Os05t0580000-01) | 1.84156 | #N/A | #N/A |
| OS10G0432200 | Conserved hypothetical protein. (Os10t0432200-01) | 1.84608 | #N/A | #N/A |
| OS10G0159300 | Conserved hypothetical protein. (Os10t0159300-01);Conserved hypothetical protein. (Os10t0159300-02) | 1.8465 | #N/A | 2.38329 |
| OS09G0454900 | Serine/threonine protein kinase-related domain containing protein. (Os09t0454900-01) | 1.84794 | #N/A | #N/A |

| | | | | |
|---|---|---|---|---|
| OS08G0369600 | Similar to Cysteine-rich polycomb-like protein. (Os08t0369600-01) | 1.84913 | #N/A | #N/A |
| OS06G0122800 | Hypothetical conserved gene. (Os06t0122800-01);Hypothetical conserved gene. (Os06t0122800-02) | 1.84927 | #N/A | #N/A |
| OS01G0391100 | Hypothetical gene. (Os01t0391100-01);Hypothetical protein. (Os01t0391100-02);Hypothetical protein. (Os01t0391100-03) | 1.85032 | #N/A | #N/A |
| OS02G0136150 | Hypothetical gene. (Os02t0136150-00) | 1.85262214 | #N/A | #N/A |
| OS12G0410050 | Hypothetical conserved gene. (Os12t0410050-01) | 1.85291 | #N/A | #N/A |
| OS01G0626100 | Armadillo-like helical domain containing protein. (Os01t0626100-01) | 1.85416 | #N/A | #N/A |
| OS11G0207400 | NB-ARC domain containing protein. (Os11t0207400-01) | 1.85463 | #N/A | #N/A |
| OS11G0592100 | Similar to Barwin. (Os11t0592100-01) | 1.85793 | #N/A | #N/A |
| OS01G0725400 | Uncharacterised protein family UPF0497, trans-membrane plant subgroup domain containing protein. (Os01t0725400-01) | 1.85887 | #N/A | #N/A |
| OS07G0484700 | Myb transcription factor domain containing protein. (Os07t0484700-01) | 1.85897 | #N/A | #N/A |
| OS05G0485300 | TRAM, LAG1 and CLN8 homology domain containing protein. (Os05t0485300-01) | 1.86562 | #N/A | #N/A |
| OS01G0504500 | Similar to Transparent testa 12 protein. (Os01t0504500-01);Multi antimicrobial extrusion protein MatE family protein. (Os01t05045 | 1.86908 | #N/A | #N/A |
| OS06G0683300 | Similar to Beta-glucosidase. (Os06t0683300-01) | 1.87 | #N/A | #N/A |
| OS08G0197400 | F-box domain, cyclin-like domain containing protein. (Os08t0197400-01) | 1.87318 | #N/A | #N/A |
| OS12G0425600 | Protein of unknown function DUF246, plant family protein. (Os12t0425600-01) | 1.87335 | #N/A | #N/A |
| OS04G0453500 | Conserved hypothetical protein. (Os04t0453500-01) | 1.87409 | #N/A | #N/A |
| OS11G0118400 | Hypothetical protein. (Os11t0118400-01) | 1.8754 | #N/A | 1.69889 |
| OS12G0571100 | Metallothionein-like protein, ROS (reactive oxygen species) scavenger, Antioxidant (Os12t0571100-01) | 1.87605 | #N/A | #N/A |
| OS02G0617100 | Conserved hypothetical protein. (Os02t0617100-01) | 1.8761 | #N/A | #N/A |
| OS06G0105700 | Protein of unknown function DUF6, transmembrane domain containing protein. (Os06t0105700-01) | 1.87967 | #N/A | #N/A |
| OS01G0609900 | Similar to Pleiotropic drug resistance protein 4. (Os01t0609900-01);Similar to PDR-like ABC transporter (PDR4 ABC transporter). (Os | 1.87992 | #N/A | #N/A |
| OS03G0277300 | Heat shock protein 70. (Os03t0277300-01) | 1.88279 | #N/A | #N/A |
| OS01G0347200 | Conserved hypothetical protein. (Os01t0347200-01) | 1.88719 | #N/A | #N/A |
| OS11G0182100 | Protein of unknown function DUF688 domain containing protein. (Os11t0182100-01) | 1.88943 | #N/A | #N/A |
| OS01G0915000 | Protein of unknown function DUF506, plant family protein. (Os01t0915000-01) | 1.88983 | #N/A | #N/A |
| OS05G0527000 | UDP-glucuronosyl/UDP-glucosyltransferase family protein. (Os05t0527000-01) | 1.89304 | #N/A | #N/A |
| OS01G0177100 | Similar to STYLOSA protein. (Os01t0177100-01);Similar to STYLOSA protein. (Os01t0177100-02) | 1.89449 | #N/A | #N/A |
| OS05G0320300 | Similar to Homeobox protein. (Os05t0320300-01) | 1.89632 | #N/A | 1.30985 |
| OS09G0101800 | Quinoprotein amine dehydrogenase, beta chain-like domain containing protein. (Os09t0101800-01) | 1.89704 | #N/A | #N/A |
| OS04G0611000 | Conserved hypothetical protein. (Os04t0611000-01);Non-protein coding transcript. (Os04t0611000-02) | 1.89836 | #N/A | #N/A |
| OS11G0117600 | Similar to WRKY transcription factor 50 (Fragment). (Os11t0117600-00) | 1.9012 | #N/A | #N/A |
| OS12G0187500 | Similar to C2 domain containing protein. (Os12t0187500-00) | 1.902 | #N/A | #N/A |
| OS04G0493600 | Similar to Lectin-C precursor (PL-C). (Os04t0493600-01) | 1.9099 | #N/A | #N/A |
| OS06G0564700 | Similar to Cysteine synthase (EC 4.2.99.8). (Os06t0564700-01) | 1.91073 | #N/A | #N/A |
| OS03G0622500 | Conserved hypothetical protein. (Os03t0622500-01);Hypothetical gene. (Os03t0622500-02) | 1.91454 | #N/A | #N/A |
| OS01G0162200 | Leucine-rich repeat domain containing protein. (Os01t0162200-00) | 1.91477 | #N/A | #N/A |
| OS08G0542300 | Similar to 60S ribosomal protein L7. (Os08t0542300-01) | 1.91778 | #N/A | #N/A |
| OS02G0574500 | General substrate transporter domain containing protein. (Os02t0574500-01);General substrate transporter domain containing pro | 1.91809 | #N/A | #N/A |
| OS10G0126700 | Similar to F-box domain containing protein, expressed. (Os10t0126700-01) | 1.9185 | #N/A | #N/A |
| OS04G0121100 | Similar to OSIGBa0115D20.4 protein. (Os04t0121100-01) | 1.91917 | #N/A | #N/A |
| OS06G0142200 | Early nodulin. (Os06t0142200-01);Similar to cDNA clone:J033147G13, full insert sequence. (Os06t0142200-02) | 1.92074 | #N/A | #N/A |
| OS04G0395800 | Tify domain containing protein. (Os04t0395800-01) | 1.92112 | #N/A | #N/A |

| | | | | |
|---|---|---|---|---|
| OS08G0143900 | Hypothetical protein. (Os08t0143900-01) | 1.92225 | #N/A | #N/A |
| OS01G0545900 | Pentatricopeptide repeat domain containing protein. (Os01t0545900-00) | 1.92327 | #N/A | #N/A |
| OS02G0528500 | Nucleic acid-binding, OB-fold domain containing protein. (Os02t0528500-01);Nucleic acid-binding, OB-fold-like domain containing p | 1.92379 | #N/A | 2.04303 |
| OS01G0720600 | Similar to Starch synthase IV. (Os01t0720600-01);Starch synthase, Starch biosynthesis (Os01t0720600-02);Similar to starch synthase | 1.92485 | #N/A | 0.920631 |
| OS06G0334100 | Hypothetical gene. (Os06t0334100-01);Conserved hypothetical protein. (Os06t0334100-02) | 1.92904 | #N/A | #N/A |
| OS06G0129900 | Similar to Cytochrome P450 CYPD. (Os06t0129900-01);Similar to cytochrome P450. (Os06t0129900-02) | 1.93011 | -0.741211 | #N/A |
| OS01G0543000 | Hypothetical gene. (Os01t0543000-02) | 1.93058 | #N/A | #N/A |
| OS12G0629700 | Similar to Thaumatin-like protein precursor. (Os12t0629700-01) | 1.94203 | #N/A | -2.5333 |
| OS07G0683200 | Similar to NAC domain transcription factor. (Os07t0683200-01) | 1.94261 | #N/A | #N/A |
| OS06G0653000 | Glossy1(GL1) homolog, Cuticular wax biosynthesis, Drought tolerance (Os06t0653000-01) | 1.94519 | #N/A | #N/A |
| OS08G0106000 | Similar to Bx2-like protein. (Os08t0106000-00) | 1.94627 | #N/A | #N/A |
| OS07G0526400 | Polyketide synthase, type III domain containing protein. (Os07t0526400-01) | 1.94734 | #N/A | #N/A |
| OS04G0107700 | Peptidase C1A, papain family protein. (Os04t0107700-01) | 1.94895 | #N/A | #N/A |
| OS07G0296000 | Conserved hypothetical protein. (Os07t0296000-01);Conserved hypothetical protein. (Os07t0296000-02) | 1.95232 | #N/A | 1.80117 |
| OS07G0694300 | Similar to Peroxidase. (Os07t0694300-01) | 1.95517 | #N/A | #N/A |
| OS03G0643300 | Ornithine delta-aminotransferase, Abiotic stress tolerance (Os03t0643300-02) | 1.95525 | #N/A | #N/A |
| OS09G0264400 | Cytochrome P450 family protein. (Os09t0264400-01) | 1.96195 | #N/A | #N/A |
| OS09G0417500 | Pentatricopeptide repeat domain containing protein. (Os09t0417500-01) | 1.96203 | #N/A | #N/A |
| OS10G0377400 | Similar to Ras-related protein Rab11D. (Os10t0377400-01);Similar to Ras-related protein Rab11D. (Os10t0377400-02);Non-protein c | 1.96547 | #N/A | #N/A |
| OS11G0551900 | Hypothetical conserved gene. (Os11t0551900-01) | 1.96644 | #N/A | #N/A |
| OS07G0677500 | Similar to Peroxidase precursor (EC 1.11.1.7). (Os07t0677500-00) | 1.96993 | #N/A | -1.0698877 |
| OS09G0482720 | Similar to lipid binding protein. (Os09t0482720-00) | 1.97454 | #N/A | #N/A |
| OS01G0349800 | Similar to Cytochrome P450. (Os01t0349800-01) | 1.97519 | #N/A | #N/A |
| OS12G0167700 | Similar to SAP domain containing protein, expressed. (Os12t0167700-01) | 1.97725 | #N/A | #N/A |
| OS09G0498700 | F-box domain, cyclin-like domain containing protein. (Os09t0498700-01) | 1.97954 | #N/A | #N/A |
| OS07G0181500 | Protein of unknown function DUF506, plant family protein. (Os07t0181500-01) | 1.98528 | #N/A | #N/A |
| OS02G0252600 | Aminotransferase, class IV family protein. (Os02t0252600-01) | 1.98537 | #N/A | #N/A |
| OS02G0154200 | Protein kinase, core domain containing protein. (Os02t0154200-01) | 1.98834 | #N/A | #N/A |
| OS08G0557800 | Similar to Fatty acyl coA reductase. (Os08t0557800-01) | 1.99021 | #N/A | #N/A |
| OS11G0687800 | NB-ARC domain containing protein. (Os11t0687800-01);NB-ARC domain containing protein. (Os11t0687800-02) | 1.99332 | #N/A | #N/A |
| OS04G0536300 | Transcription factor with zinc finger domain and helix-loop-helix domain (YABBY domain), Leaf development (Os04t0536300-01) | 2.00077 | - | - |
| OS07G0593400 | Similar to golgi transport 1 protein B. (Os07t0593400-00) | 2.00321 | - | - |
| OS12G0218100 | Similar to cDNA, clone: J065028F04, full insert sequence. (Os12t0218100-01) | 2.00541 | - | - |
| OS01G0692000 | Similar to Glutathione S-transferase GSTU6. (Os01t0692000-01) | 2.00656 | - | - |
| OS03G0720400 | Cyclin-like F-box domain containing protein. (Os03t0720400-01) | 2.00675 | - | - |
| OS03G0337200 | Hypothetical protein. (Os03t0337200-01);Hypothetical protein. (Os03t0337200-02);Non-protein coding transcript. (Os03t0337200-( | 2.00698 | - | - |
| OS02G0469300 | Similar to OSIGBa0148D14.7 protein. (Os02t0469300-01);ATP-binding region, ATPase-like domain containing protein. (Os02t046930 | 2.01262 | - | - |
| OS07G0647600 | Myb/SANT-like domain domain containing protein. (Os07t0647600-01) | 2.01378 | - | - |
| OS01G0359600 | Disease resistance protein domain containing protein. (Os01t0359600-00) | 2.02992 | #N/A | 1.37033 |
| OS10G0372800 | Conserved hypothetical protein. (Os10t0372800-01) | 2.03048 | - | - |
| OS04G0378400 | Non-protein coding transcript. (Os04t0378400-01) | 2.03582 | - | - |
| OS02G0167200 | Pentatricopeptide repeat domain containing protein. (Os02t0167200-01) | 2.03867 | #N/A | 2.50367 |

| | | | | |
|---|---|---|---|---|
| OS11G0116900 | Similar to WRKY DNA binding domain containing protein, expressed. (Os11t0116900-01) | 2.052 | - | - |
| OS08G0189200 | Germin-like protein 8-3, Disease resistance (Os08t0189200-01) | 2.05327 | -0.944552 | #N/A |
| OS03G0760000 | Cytochrome P450 family protein. (Os03t0760000-01) | 2.05674 | - | - |
| OS10G0556900 | Conserved hypothetical protein. (Os10t0556900-01) | 2.05713 | - | - |
| OS03G0754200 | Similar to Triosephosphate isomerase. (Os03t0754200-01);Similar to Triosephosphate isomerase. (Os03t0754200-02);Similar to Tric | 2.05829 | - | - |
| OS09G0262000 | Similar to Cinnamoyl CoA reductase. (Os09t0262000-00) | 2.0696 | -1.20191 | -1.27271 |
| OS04G0127300 | Peptidase S8 and S53, subtilisin, kexin, sedolisin domain containing protein. (Os04t0127300-01) | 2.0748 | - | - |
| OS02G0719600 | SAM dependent carboxyl methyltransferase family protein. (Os02t0719600-01) | 2.08126 | - | - |
| OS01G0953400 | NB-ARC domain containing protein. (Os01t0953400-01) | 2.08317 | #N/A | 1.75173 |
| OS01G0748100 | Conserved hypothetical protein. (Os01t0748100-01) | 2.09243 | - | - |
| OS04G0675800 | Similar to H0103C06.10 protein. (Os04t0675800-00) | 2.09347 | - | - |
| OS01G0136100 | 16.9 kDa class I heat shock protein 1. (Os01t0136100-01) | 2.09471 | - | - |
| OS02G0755900 | UDP-glucuronosyl/UDP-glucosyltransferase domain containing protein. (Os02t0755900-01) | 2.09591 | - | - |
| OS01G0328900 | Glycosyl transferase, family 31 domain containing protein. (Os01t0328900-01) | 2.10501 | - | - |
| OS11G0164900 | Conserved hypothetical protein. (Os11t0164900-00) | 2.1083 | - | - |
| OS02G0685600 | Similar to Protein phosphatase 2C-like. (Os02t0685600-01) | 2.10978 | - | - |
| OS01G0101800 | Conserved hypothetical protein. (Os01t0101800-01) | 2.11386 | - | - |
| OS01G0628900 | Cytochrome P450 family protein. (Os01t0628900-01) | 2.11396 | - | - |
| OS12G0552400 | Rossmann-like alpha/beta/alpha sandwich fold domain containing protein. (Os12t0552400-01) | 2.12201 | - | - |
| OS06G0695400 | Haem peroxidase family protein. (Os06t0695400-01) | 2.12893343 | - | - |
| OS12G0633600 | Protein of unknown function DUF221 domain containing protein. (Os12t0633600-01) | 2.1291 | - | - |
| OS11G0659200 | Protein of unknown function DUF581 family protein. (Os11t0659200-00) | 2.13951861 | - | - |
| OS08G0528700 | Conserved hypothetical protein. (Os08t0528700-01) | 2.14277 | - | - |
| OS04G0524400 | Similar to OSIGBa0153E02-OSIGBa0093I20.13 protein. (Os04t0524400-01) | 2.146 | - | - |
| OS06G0163701 | Non-protein coding transcript. (Os06t0163701-01) | 2.15254 | #N/A | 1.74495 |
| OS01G0720801 | Hypothetical gene. (Os01t0720801-01) | 2.15518 | - | - |
| OS01G0516700 | Similar to glycosyltransferase family 28 C-terminal domain containing protein. (Os01t0516700-01) | 2.15573 | - | - |
| OS08G0382800 | Hypothetical conserved gene. (Os08t0382800-01) | 2.16625 | - | - |
| OS01G0506200 | Tetratricopeptide-like helical domain containing protein. (Os01t0506200-01) | 2.16705 | - | - |
| OS04G0466000 | Similar to OSIGBa0115M15.3 protein. (Os04t0466000-01) | 2.16709 | - | - |
| OS07G0223700 | Similar to predicted protein. (Os07t0223700-00) | 2.16941 | - | - |
| OS11G0103100 | Hypothetical conserved gene. (Os11t0103100-00) | 2.17596 | - | - |
| OS03G0733200 | Non-protein coding transcript. (Os03t0733200-00) | 2.18318 | - | - |
| OS05G0150600 | Hypothetical conserved gene. (Os05t0150600-01);DNA helicase, ATP-dependent, RecQ type domain containing protein. (Os05t0150 | 2.18487 | - | - |
| OS04G0340300 | Terpenoid synthase domain containing protein. (Os04t0340300-01) | 2.18881 | -1.09718 | -1.35019 |
| OS03G0103400 | GRAS transcription factor domain containing protein. (Os03t0103400-01) | 2.1895 | - | - |
| OS08G0444440 | Hypothetical conserved gene. (Os08t0444440-01) | 2.19072 | - | - |
| OS09G0545300 | SAUR family protein, Negative regulator of auxin synthesis and transport (Os09t0545300-01) | 2.20204 | - | - |
| OS05G0211100 | Similar to Cytochrome P450-like protein. (Os05t0211100-01);Obtusifoliol 14a-demethylase, Phytosterol and Brassinosteroid biosynt | 2.20627 | - | - |
| OS04G0201200 | Hypothetical conserved gene. (Os04t0201200-01) | 2.21245 | - | - |
| OS04G0420801 | Hypothetical protein. (Os04t0420801-00) | 2.21384 | - | - |
| OS02G0782900 | Conserved hypothetical protein. (Os02t0782900-01) | 2.21553 | - | - |

| | | | | |
|---|---|---|---|---|
| OS01G0777300 | DNA repair protein (XPGC)/yeast Rad family protein. (Os01t0777300-00) | 2.2334 | - | - |
| OS01G0133100 | Conserved hypothetical protein. (Os01t0133100-01);Hypothetical conserved gene. (Os01t0133100-02) | 2.23481 | - | - |
| OS01G0307686 | Similar to embryonic abundant protein-like. (Os01t0307686-00) | 2.23652 | - | - |
| OS01G0764800 | Indole-3-acetic acid (IAA)-amido synthetase, Disease resistance, Abiotic stress tolerance (Os01t0764800-01) | 2.23801 | - | - |
| OS05G0417100 | Chloroplast-targeted Deg protease protein, Chloroplast development and maintenance of PSII function under high temperatures (O | 2.24056 | #N/A | 1.6977 |
| OS03G0187100 | Hypothetical conserved gene. (Os03t0187100-00) | 2.25002 | #N/A | 1.2239 |
| OS04G0614300 | Hypothetical conserved gene. (Os04t0614300-01) | 2.25119 | - | - |
| OS10G0167250 | Similar to Cytochrome P450 CYP76H18. (Os10t0167250-00) | 2.25171 | - | - |
| OS01G0884400 | Armadillo domain containing protein. (Os01t0884400-01) | 2.25273 | #N/A | 1.65555 |
| OS06G0602500 | Serine/threonine protein kinase-related domain containing protein. (Os06t0602500-01) | 2.26173 | - | - |
| OS05G0325200 | Cyclin-related 2 domain containing protein. (Os05t0325200-01) | 2.27415 | - | - |
| OS06G0585982 | Protein kinase, catalytic domain domain containing protein. (Os06t0585982-00) | 2.27521 | - | - |
| OS03G0107000 | Similar to ASK20. (Os03t0107000-01) | 2.27648 | #N/A | 1.98944 |
| OS04G0583200 | Conserved hypothetical protein. (Os04t0583200-01) | 2.27726 | - | - |
| OS04G0201000 | Similar to OSIGBa0113K06.10 protein. (Os04t0201000-00) | 2.27954 | - | - |
| OS11G0209600 | Cyclin-like F-box domain containing protein. (Os11t0209600-01);Similar to F-box family-1. (Os11t0209600-02) | 2.27983 | - | - |
| OS07G0537900 | Similar to SRK3 gene. (Os07t0537900-01);Hypothetical conserved gene. (Os07t0537900-02) | 2.28565 | - | - |
| OS03G0624300 | Conserved hypothetical protein. (Os03t0624300-01) | 2.28743 | - | - |
| OS09G0110300 | Putative cyclase family protein. (Os09t0110300-01);Putative cyclase family protein. (Os09t0110300-02) | 2.2904 | - | - |
| OS01G0107700 | Similar to LIMONENE cyclase like protein. (Os01t0107700-01) | 2.29053 | #N/A | 1.58867 |
| OS03G0240400 | Conserved hypothetical protein. (Os03t0240400-01) | 2.30128 | #N/A | 1.9279 |
| OS03G0108200 | Similar to Transposon protein. (Os03t0108200-00) | 2.30675 | - | - |
| OS02G0740500 | Conserved hypothetical protein. (Os02t0740500-01) | 2.30879 | - | - |
| OS01G0917500 | Leucine-rich repeat receptor-like kinase, Specification of anther cell identity, Control of early sporogenic development, Initiation of | 2.31755 | - | - |
| OS03G0816600 | Similar to pentatricopeptide repeat-containing protein. (Os03t0816600-01) | 2.32254 | - | - |
| OS01G0533900 | Similar to Multidrug resistance protein 1 homolog. (Os01t0533900-01);Similar to MDR-like ABC transporter. (Os01t0533900-02) | 2.32706 | - | - |
| OS03G0614901 | Conserved hypothetical protein. (Os03t0614901-01) | 2.32731 | - | - |
| OS11G0679900 | Conserved hypothetical protein. (Os11t0679900-01) | 2.32974 | #N/A | 2.2266 |
| OS01G0153700 | Conserved hypothetical protein. (Os01t0153700-01) | 2.33155 | - | - |
| OS06G0286500 | Similar to NBS-LRR disease resistance protein homologue. (Os06t0286500-01) | 2.3415 | - | - |
| OS06G0216300 | Similar to 12-oxophytodienoic acid reductase. (Os06t0216300-01) | 2.34162 | - | - |
| OS07G0418700 | Proline-rich glycoprotein, ABA-dependent inhibition of root growth (Os07t0418700-01) | 2.36677 | - | - |
| OS02G0191300 | Similar to Amino acid transporter-like protein. (Os02t0191300-01) | 2.36805 | - | - |
| OS12G0471100 | Similar to ATPase 2. (Os12t0471100-01) | 2.36898 | - | - |
| OS02G0218700 | Allene oxide synthase (CYP74A3), Fatty acid 9-/13-hydroperoxide lyase (CYP74C), Biosynthesis of jasmonic acid (JA), Plant defense ( | 2.37099004 | - | - |
| OS11G0282700 | Homeodomain-like containing protein. (Os11t0282700-01) | 2.3755 | - | - |
| OS02G0190300 | P-glycoprotein homologue. (Os02t0190300-01) | 2.38078 | - | - |
| OS01G0859100 | Similar to WIP1 protein (Fragment). (Os01t0859100-01) | 2.39834 | - | - |
| OS01G0795400 | Similar to Subtilase. (Os01t0795400-01) | 2.40577 | - | - |
| OS12G0580600 | Conserved hypothetical protein. (Os12t0580600-01) | 2.40954 | - | - |
| OS02G0481700 | Similar to lipase class 3 family protein. (Os02t0481700-01) | 2.41588 | - | - |
| OS09G0334600 | Similar to Cytochrome b5. (Os09t0334600-00) | 2.43418 | - | - |

| | | | | |
|---|---|---|---|---|
| OS11G0117400 | Similar to WRKY DNA binding domain containing protein, expressed. (Os11t0117400-00) | 2.43601 | - | - |
| OS06G0168700 | Similar to Prolin rich protein. (Os06t0168700-01) | 2.44025 | - | - |
| OS01G0673900 | Zinc finger, RING-type domain containing protein. (Os01t0673900-01) | 2.44628 | - | - |
| OS07G0539350 | Non-protein coding transcript. (Os07t0539350-00) | 2.44664 | - | - |
| OS02G0164000 | Peptidase C48, SUMO/Sentrin/Ubl1 domain containing protein. (Os02t0164000-00) | 2.4585 | - | - |
| OS04G0126900 | Similar to H0124E07.4 protein. (Os04t0126900-01) | 2.45993254 | - | - |
| OS10G0558700 | Similar to Oxidoreductase, 2OG-Fe oxygenase family protein, expressed. (Os10t0558700-01);2OG-Fe(II) oxygenase domain containi | 2.46195 | - | - |
| OS08G0461700 | F-box domain, cyclin-like domain containing protein. (Os08t0461700-01) | 2.47748 | - | - |
| OS07G0656200 | Similar to Beta-glucosidase 26. (Os07t0656200-01) | 2.48001 | - | - |
| OS09G0544000 | Transferase family protein. (Os09t0544000-01) | 2.49014 | - | - |
| OS11G0702100 | Similar to Class III chitinase homologue (OsChib3H-h) (Fragment). (Os11t0702100-01) | 2.51015 | - | - |
| OS08G0383250 | Hypothetical conserved gene. (Os08t0383250-01) | 2.52272 | - | - |
| OS05G0399400 | Chitinase 9. (Os05t0399400-00) | 2.53339 | - | - |
| OS07G0684800 | Similar to NAM / CUC2-like protein. (Os07t0684800-01);Similar to NAM / CUC2-like protein. (Os07t0684800-02) | 2.54136 | - | - |
| OS12G0161900 | Hypothetical conserved gene. (Os12t0161900-01) | 2.54983 | - | - |
| OS11G0260100 | SAM dependent carboxyl methyltransferase family protein. (Os11t0260100-00) | 2.5623 | - | - |
| OS09G0472700 | Similar to blight-associated protein p12. (Os09t0472700-00) | 2.57569 | - | - |
| OS11G0590400 | Similar to NB-ARC domain containing protein. (Os11t0590400-00) | 2.57713 | - | - |
| OS03G0734200 | Conserved hypothetical protein. (Os03t0734200-01) | 2.58674 | - | - |
| OS12G0637400 | Similar to Purple acid phosphatase (EC 3.1.3.2) (Fragment). (Os12t0637400-01) | 2.58704 | - | - |
| OS09G0532000 | Senescence-inducible chloroplast protein, Activation of the chlorophll-degrading pathway during leaf senescence (Os09t0532000-0 | 2.59811 | - | - |
| OS09G0275400 | Cytochrome P450 family protein. (Os09t0275400-01) | 2.60631667 | - | - |
| OS01G0706700 | Protein of unknown function DUF590 family protein. (Os01t0706700-01) | 2.6151 | #N/A | 2.26141 |
| OS12G0617100 | Similar to Translation initiation factor eIF-2B delta subunit (eIF-2B GDP-GTP exchange factor) (Guanine nucleotide exchange factor s | 2.61621 | - | - |
| OS03G0626500 | Similar to predicted protein. (Os03t0626500-00) | 2.6166 | - | - |
| OS09G0380450 | Conserved hypothetical protein. (Os09t0380450-01) | 2.62323 | - | - |
| OS11G0684000 | Similar to Transcription factor MYB21 (Myb-related protein 21) (AtMYB21) (Myb homolog 3) (AtMyb3). (Os11t0684000-01) | 2.63331 | - | - |
| OS08G0166400 | Zinc finger, PMZ-type domain containing protein. (Os08t0166400-01) | 2.6338 | - | - |
| OS03G0168300 | Protein of unknown function DUF1997 domain containing protein. (Os03t0168300-01) | 2.64491 | - | - |
| OS07G0593000 | Similar to Zinc-finger protein KNUCKLES. (Os07t0593000-01) | 2.64521 | - | - |
| OS08G0234400 | Hypothetical gene. (Os08t0234400-01) | 2.65269 | - | - |
| OS03G0693700 | Similar to Oxalate oxidase 1 (EC 1.2.3.4) (Germin). (Os03t0693700-01) | 2.66224652 | - | - |
| OS03G0198600 | Homeodomain-leucine zipper transcription factor, Regulation of panicle exsertion (Os03t0198600-01) | 2.66495 | #N/A | 1.90572 |
| OS03G0288000 | Similar to Metallothionein. (Os03t0288000-01);Similar to Metallothionein. (Os03t0288000-02) | 2.66685 | - | - |
| OS07G0132500 | Similar to Resistance protein candidate (Fragment). (Os07t0132500-01) | 2.66873 | - | - |
| OS02G0690500 | Similar to TA11 protein (Fragment). (Os02t0690500-01) | 2.68158 | - | - |
| OS04G0613000 | Zinc transporter, Preferential distribution of Zn to developing tissues (Os04t0613000-01) | 2.68761 | - | - |
| OS12G0548501 | Similar to CI2C. (Os12t0548501-01) | 2.70466395 | - | - |
| OS05G0500900 | Similar to Indole-3-acetic acid-amido synthetase GH3.5 (EC 6.3.2.-) (Auxin- responsive GH3-like protein 5) (AtGH3-5). (Os05t050090 | 2.73781 | - | - |
| OS01G0108500 | Protein of unknown function DUF3778 domain containing protein. (Os01t0108500-00) | 2.73986 | - | - |
| OS01G0670100 | Serine/threonine protein kinase-related domain containing protein. (Os01t0670100-01) | 2.76212 | - | - |
| OS01G0162300 | Leucine-rich repeat, plant specific containing protein. (Os01t0162300-01) | 2.77525 | - | - |

| | | | | |
|---|---|---|---|---|
| OS09G0471800 | Similar to WAK80 - OsWAK receptor-like protein kinase. (Os09t0471800-01) | 2.77566 | #N/A | 2.09613 |
| OS11G0208000 | Cyclin-like F-box domain containing protein. (Os11t0208000-01);Cyclin-like F-box domain containing protein. (Os11t0208000-02) | 2.77664 | - | - |
| OS02G0667500 | Similar to tetracycline transporter protein. (Os02t0667500-01);Hypothetical conserved gene. (Os02t0667500-02) | 2.78019 | - | - |
| OS01G0810401 | Non-protein coding transcript. (Os01t0810401-01) | 2.78684 | - | - |
| OS08G0518800 | Similar to Class III chitinase homologue (OsChib3H-h) (Fragment). (Os08t0518800-01) | 2.78977098 | - | - |
| OS02G0776900 | Transcription activator, Gibberellin (GA)-induced stem elongation (Os02t0776900-01);Similar to Growth-regulating factor 1. (Os02t( | 2.80663 | - | - |
| OS08G0433500 | Senescence-associated NAC transcription factor, Negative regulation of leaf senescence, Salt stress tolerance, Plant architecture (Os | 2.80969 | - | - |
| OS02G0208400 | Hypothetical gene. (Os02t0208400-01);Hypothetical gene. (Os02t0208400-02) | 2.82995 | - | - |
| OS08G0307000 | Conserved hypothetical protein. (Os08t0307000-01) | 2.83341 | - | - |
| OS12G0473900 | Similar to Protease inhibitor/seed storage/LTP family protein. (Os12t0473900-00) | 2.84657 | - | - |
| OS06G0352200 | Protein of unknown function DUF679 family protein. (Os06t0352200-01) | 2.86326 | - | - |
| OS02G0453800 | Conserved hypothetical protein. (Os02t0453800-01) | 2.88053 | - | - |
| OS06G0307900 | Hypothetical conserved gene. (Os06t0307900-01);Protein of unknown function DUF1618 domain containing protein. (Os06t030790 | 2.89245 | - | - |
| OS07G0535200 | Cyclin-like F-box domain containing protein. (Os07t0535200-01) | 2.91166 | - | - |
| OS04G0659300 | Receptor-like protein, Root development, Salt stress response, Regulation of iron acquisition (Os04t0659300-01) | 2.93877 | - | - |
| OS08G0470700 | Similar to carbonic anhydrase. (Os08t0470700-00) | 2.95099 | - | - |
| OS07G0116900 | Conserved hypothetical protein. (Os07t0116900-01) | 2.96528 | - | - |
| OS11G0492100 | Similar to NB-ARC domain containing protein, expressed. (Os11t0492100-01) | 2.97839 | - | - |
| OS07G0622700 | Alpha/beta hydrolase fold-1 domain containing protein. (Os07t0622700-01) | 2.99559 | - | - |
| OS04G0420900 | Similar to H0525E10.7 protein. (Os04t0420900-01);Similar to Receptor-like protein kinase. (Os04t0420900-02) | 3.00492 | - | - |
| OS02G0620800 | Conserved hypothetical protein. (Os02t0620800-01) | 3.06296 | - | - |
| OS02G0537400 | Similar to Heat shock protein. (Os02t0537400-01) | 3.06524 | - | - |
| OS02G0467300 | Conserved hypothetical protein. (Os02t0467300-01) | 3.07131 | - | - |
| OS05G0465000 | Conserved hypothetical protein. (Os05t0465000-01) | 3.09155 | - | - |
| OS03G0195100 | Putative aminotransferase, Disease resistance response against infection with rice blast fungus (Os03t0195100-01) | 3.09802 | - | - |
| OS04G0565200 | Similar to Cis-zeatin O-glucosyltransferase 1 (EC 2.4.1.215) (cisZOG1). (Os04t0565200-01) | 3.13262 | - | - |
| OS06G0207200 | Conserved hypothetical protein. (Os06t0207200-01) | 3.15113 | - | - |
| OS07G0290500 | Plant lipid transfer/seed storage/trypsin-alpha amylase inhibitor domain containing protein. (Os07t0290500-00) | 3.1630752 | - | - |
| OS11G0213800 | Similar to NBS-LRR disease resistance protein homologue (Fragment). (Os11t0213800-01) | 3.19665 | - | - |
| OS05G0455900 | Pentatricopeptide repeat domain containing protein. (Os05t0455900-01) | 3.23144 | #N/A | 2.85672 |
| OS03G0287400 | Similar to LOB domain protein 4. (Os03t0287400-01) | 3.23383 | - | - |
| OS08G0188000 | RNA-binding, CRM domain domain containing protein. (Os08t0188000-01) | 3.25084 | - | - |
| OS10G0515900 | Cytochrome P450 family protein. (Os10t0515900-01) | 3.29436 | - | - |
| OS07G0638400 | Similar to 1-Cys peroxiredoxin. (Os07t0638400-01) | 3.30809 | - | - |
| OS05G0122700 | Similar to Low-temperature induced protein lt101.2. (Os05t0122700-01) | 3.32558789 | - | - |
| OS11G0147150 | Hypothetical gene. (Os11t0147150-00) | 3.35907 | - | - |
| OS02G0742900 | Similar to Kinesin-like protein. (Os02t0742900-01) | 3.36136 | - | - |
| OS06G0244000 | Similar to anthranilic acid methyltransferase 3. (Os06t0244000-01) | 3.3619 | - | - |
| OS03G0361500 | Similar to Terpene synthase family, metal binding domain containing protein, expressed. (Os03t0361500-00) | 3.36343 | - | - |
| OS06G0125200 | Protein of unknown function DUF581 family protein. (Os06t0125200-01) | 3.36753 | - | - |
| OS09G0268000 | Similar to lectin-like receptor kinase 7. (Os09t0268000-00) | 3.41711 | - | - |
| OS03G0611100 | Non-protein coding transcript. (Os03t0611100-01) | 3.4989 | - | - |

| | | | | |
|---|---|---|---|---|
| OS01G0163000 | Leucine-rich repeat, N-terminal domain containing protein. (Os01t0163000-01) | 3.65299 | - | - |
| OS12G0559250 | Non-protein coding transcript. (Os12t0559250-00) | 3.6581 | - | - |
| OS12G0564100 | Similar to Myb-like DNA-binding domain containing protein, expressed. (Os12t0564100-01) | 3.70519 | - | - |
| OS01G0283700 | Similar to Cinnamoyl-CoA reductase (EC 1.2.1.44). (Os01t0283700-01) | 3.71645 | - | - |
| OS07G0130100 | Similar to Resistance protein candidate (Fragment). (Os07t0130100-01) | 3.76318 | - | - |
| OS10G0368902 | Pentatricopeptide repeat domain containing protein. (Os10t0368902-00) | 3.76358 | - | - |
| OS02G0783700 | Similar to Lysine ketoglutarate reductase/saccharopine dehydrogenase. (Os02t0783700-01) | 3.80267 | - | - |
| OS08G0248100 | Protein kinase, core domain containing protein. (Os08t0248100-01) | 3.81622 | - | - |
| OS11G0512600 | Similar to No apical meristem protein. (Os11t0512600-00) | 3.83189 | - | - |
| OS10G0504900 | Similar to Lipid transfer protein. (Os10t0504900-01) | 3.84952 | - | - |
| OS07G0117900 | NB-ARC domain containing protein. (Os07t0117900-00) | 3.97067 | - | - |
| OS12G0187800 | Conserved hypothetical protein. (Os12t0187800-00) | 3.9775 | - | - |
| OS03G0193300 | Conserved hypothetical protein. (Os03t0193300-01);Conserved hypothetical protein. (Os03t0193300-02) | 4.04184 | - | - |
| OS10G0118000 | Similar to 5-pentadecatrienyl resorcinol O-methyltransferase. (Os10t0118000-01) | 4.04909 | - | - |
| OS02G0733900 | Conserved hypothetical protein. (Os02t0733900-01) | 4.05174 | - | - |
| OS08G0207800 | Peptidase aspartic, catalytic domain containing protein. (Os08t0207800-01) | 4.07304 | - | - |
| OS09G0109600 | Conserved hypothetical protein. (Os09t0109600-01) | 4.09711 | - | - |
| OS09G0439800 | Control of root system architecture, Drought avoidance (Os09t0439800-01) | 4.26865 | - | - |
| OS03G0830500 | Similar to PGPS/D12. (Os03t0830500-01) | 4.27154 | -0.3871309 | #N/A |
| OS02G0586900 | Hypothetical conserved gene. (Os02t0586900-01);Similar to Glycine rich protein (Fragment). (Os02t0586900-02) | 4.35786 | - | - |
| OS03G0738600 | Lipoxygenase, Seed germination and longevity (Os03t0738600-01) | 4.47679 | - | - |
| OS03G0638400 | Similar to OSIGBa0153E02-OSIGBa0093I20.19 protein. (Os03t0638400-01) | 4.4784 | - | - |
| OS02G0555300 | No apical meristem (NAM) protein domain containing protein. (Os02t0555300-01) | 4.86947 | - | - |
| OS04G0108600 | Similar to Sesquiterpene synthase (Fragment). (Os04t0108600-00) | 5.10331719 | - | - |
| OS03G0305600 | Mitochondrial import inner membrane translocase, subunit Tim17/22 family protein. (Os03t0305600-01) | 5.50222 | - | - |
| OS07G0526600 | Alpha/beta hydrolase fold-3 domain containing protein. (Os07t0526600-01) | 5.75728 | - | - |
| OS08G0168000 | Similar to Sesquiterpene synthase. (Os08t0168000-01) | 6.18092 | #N/A | 0.59853773 |
| OS06G0242700 | SAM dependent carboxyl methyltransferase family protein. (Os06t0242700-01) | 6.20984 | - | - |

### 6. OXs compared to WT (mock)

| Identifier | Description | FC.NmSm | FC.NmSSm | FC.NmNs | FC.SmSs | FC.SSmSSs |
|---|---|---|---|---|---|---|
| OS07G0417200 | Similar to Delta 12 oleic acid desaturase FAD2. (Os07t0417200-01) | -4.11333 | -3.363928 | -1.58926 | - | - |
| OS12G0247700 | Similar to Jasmonate-induced protein. (Os12t0247700-01) | -4.10315 | -2.38519 | 3.98897 | 3.34788 | 4.74645 |
| OS05G0312600 | EF-Hand type domain containing protein. (Os05t0312600-01) | -3.54928 | -3.09261 | -3.10657 | 2.46671 | - |
| OS02G0570700 | Cytochrome P450 family protein. (Os02t0570700-01) | -3.48937 | -5.87605 | -1.67975 | 1.51277 | - |
| OS01G0796000 | Conserved hypothetical protein. (Os01t0796000-01) | -3.38126 | -3.32761 | -1.12522 | - | - |
| OS01G0382000 | Similar to Pathogenesis-related protein PRB1-2 precursor. (Os01t0382000-01) | -3.3488 | -4.91563 | -0.786623 | 1.36873 | 1.09739 |
| OS12G0215950 | Similar to Leucine Rich Repeat family protein, expressed. (Os12t0215950-01) | -3.20577 | -4.03763 | -1.84211 | - | - |
| OS12G0555500 | Probenazole-inducible protein PBZ1. (Os12t0555500-01) | -3.18905 | -5.03856 | - | 1.16947 | 2.44189 |
| OS05G0182600 | Similar to SSRP1 protein. (Os05t0182600-01) | -3.16674 | -2.97544 | - | - | 2.70023 |
| OS07G0129200 | Pathogenesis-related 1a protein, Regulation of abiotic stress responses (Os07t0129200-01) | -3.14396 | -6.4776 | -1.22483 | - | - |
| OS06G0587900 | Hypothetical conserved gene. (Os06t0587900-00) | -3.1136 | -2.37947 | -0.949622 | - | - |
| OS06G0569500 | Ent-kaurene oxidase, Diterpenoid phytoalexin biosynthesis (Os06t0569500-01) | -3.04974 | -4.77379 | -0.87372 | 1.44838 | - |
| OS07G0129300 | Similar to PR1a protein. (Os07t0129300-00) | -2.98887 | -2.96291 | -1.02941 | - | - |
| OS04G0104900 | Similar to 0-methyltransferase (EC 2.1.1.6) (Fragment). (Os04t0104900-01) | -2.94674 | -4.74839 | -0.942286 | - | - |
| OS01G0197700 | Similar to Cytokinin dehydrogenase 2. (Os01t0197700-01) | -2.90949 | -1.75741 | -2.04254 | - | - |
| OS12G0548401 | Similar to Proteinase inhibitor. (Os12t0548401-01) | -2.89271 | -1.98868 | 3.82512 | 3.59409 | 3.43253 |
| OS12G0628600 | Similar to Thaumatin-like pathogenesis-related protein 3 precursor. (Os12t0628600-01) | -2.88413 | -2.36398 | - | 1.17938 | 1.31315 |
| OS08G0167800 | Similar to Sesquiterpene synthase. (Os08t0167800-01) | -2.83597 | -2.32753 | 2.60229 | - | 2.48306 |
| OS02G0569900 | Cytochrome P450 family protein. (Os02t0569900-01) | -2.82853 | -3.24589 | -1.16642 | - | - |
| OS02G0686300 | Similar to H0307D04.3 protein. (Os02t0686300-01) | -2.82294 | -1.56719 | -1.17586 | 2.32035 | - |
| OS01G0794800 | Similar to Subtilase. (Os01t0794800-01);Similar to Subtilase. (Os01t0794800-02) | -2.8148 | -2.74152 | -2.50751 | - | - |
| OS12G0218300 | Conserved hypothetical protein. (Os12t0218300-01) | -2.7963 | -2.40839 | - | - | - |
| OS10G0537800 | Peptidase aspartic, catalytic domain containing protein. (Os10t0537800-01) | -2.78682 | -3.44302 | -1.39709 | - | - |
| OS07G0522500 | Similar to PDR6 ABC transporter. (Os07t0522500-01) | -2.71308 | -2.55305 | -0.577144 | 1.5519 | - |
| OS07G0656400 | Protein of unknown function DUF288 domain containing protein. (Os07t0656400-01) | -2.59734 | -2.58221 | -1.11079 | - | 2.9266 |
| OS07G0663700 | Similar to Oxidoreductase, short chain dehydrogenase/reductase family protein, expressed. (Os07t0663700-01) | -2.53788 | -4.91791 | -1.55181 | - | - |
| OS01G0795000 | Subtilase. (Os01t0795000-01) | -2.53613 | -4.03328 | -1.5445 | - | - |
| OS07G0614600 | Transmembrane receptor, eukaryota domain containing protein. (Os07t0614600-01) | -2.48989 | -2.29112 | -1.55201 | - | - |
| OS12G0629300 | Similar to Thaumatin-like protein. (Os12t0629300-00) | -2.48588 | -2.64623 | - | - | - |
| OS08G0340900 | Pentatricopeptide repeat domain containing protein. (Os08t0340900-01) | -2.47285 | -1.47888 | -2.14768 | - | - |
| OS03G0838400 | Ammonium transporter. (Os03t0838400-01);Similar to Ammonium transporter. (Os03t0838400-02) | -2.471 | -2.64931 | - | - | - |
| OS11G0514400 | Similar to BRASSINOSTEROID INSENSITIVE 1-associated receptor kinase 1. (Os11t0514400-00) | -2.44602 | -4.68931 | -0.958482 | - | - |
| OS04G0122200 | Similar to OSIGBa0137A06.2 protein. (Os04t0122200-00) | -2.42479 | -4.02222 | -0.877836 | - | - |
| OS01G0940700 | Similar to Glucan endo-1,3-beta-glucosidase GII precursor (EC 3.2.1.39) ((1->3)- beta-glucan endohydrolase GII) ((1->3)-beta-gluc | -2.3976 | -2.16351 | - | 0.870965 | 0.828667 |
| OS12G0221400 | Similar to cDNA clone:001-112-B09, full insert sequence. (Os12t0221400-01) | -2.35869 | -2.99783 | - | - | - |
| OS12G0170800 | Similar to Citrate binding protein precursor. (Os12t0170800-01) | -2.30423 | -2.22993 | - | - | - |
| OS11G0537300 | Similar to DedA. (Os11t0537300-01) | -2.30148 | -1.65774 | - | - | 1.92663 |
| OS04G0677300 | Harpin-induced 1 domain containing protein. (Os04t0677300-01) | -2.2962 | -3.79267 | - | 2.18501 | - |
| OS06G0493100 | Conserved hypothetical protein. (Os06t0493100-01) | -2.26626 | -2.15578 | - | - | - |
| OS06G0563900 | Similar to Diacylglycerol acyltransferase. (Os06t0563900-01) | -2.2614 | -1.48545 | -1.20387 | 1.53124 | - |
| OS12G0458100 | Transferase family protein. (Os12t0458100-01) | -2.2217 | -1.6181 | 3.90497 | 1.81144 | 3.62784 |
| OS06G0591200 | Conserved hypothetical protein. (Os06t0591200-01) | -2.22147 | -2.23148 | - | - | - |
| OS03G0103200 | Similar to Physical impedance induced protein. (Os03t0103200-01) | -2.21187 | -1.895 | - | - | 0.807515 |

| | | | | | | |
|---|---|---|---|---|---|---|
| OS01G0695800 | Similar to MDR-like ABC transporter. (Os01t0695800-01) | -2.18477 | -2.02224 | - | - | - |
| OS11G0641500 | Cupredoxin domain containing protein. (Os11t0641500-01) | -2.18341 | -4.38129 | -1.21786 | - | 2.83335 |
| OS04G0449500 | Similar to H0818E04.12 protein. (Os04t0449500-01) | -2.1677 | -1.99332 | - | - | - |
| OS12G0197600 | Hypothetical gene. (Os12t0197600-01) | -2.15177 | -1.63631 | - | 1.92621 | 1.79284 |
| OS10G0542900 | Similar to chitinase. (Os10t0542900-01) | -2.14407 | -2.76851 | 1.29672 | 0.928762 | 1.76404 |
| OS12G0555000 | Similar to Probenazole-inducible protein PBZ1. (Os12t0555000-01) | -2.12319 | -4.72791 | - | - | 1.49445 |
| OS09G0412300 | Similar to Calmodulin-like protein. (Os09t0412300-01) | -2.11682 | -1.48449 | -2.08972 | - | - |
| OS01G0601700 | Leucine-rich repeat domain containing protein. (Os01t0601700-01) | -2.11584 | -1.95781 | - | - | - |
| OS04G0556400 | Similar to UDP-glycosyltransferase UGT93B9. (Os04t0556400-01) | -2.10113 | -2.9835 | -0.932065 | - | - |
| OS08G0331800 | Conserved hypothetical protein. (Os08t0331800-01) | -2.09338 | -2.97227 | -1.29641 | - | - |
| OS12G0635500 | Protein of unknown function DUF266, plant domain containing protein. (Os12t0635500-01) | -2.05047 | -1.54133 | -1.24348 | 1.61925 | - |
| OS01G0714600 | Similar to cDNA clone:J023088C01, full insert sequence. (Os01t0714600-01) | -2.04591 | -1.70802 | - | - | - |
| OS08G0256700 | Conserved hypothetical protein. (Os08t0256700-01) | -2.04065 | -2.24167 | -1.42436 | - | - |
| OS08G0197000 | Cyclin-like F-box domain containing protein. (Os08t0197000-01) | -1.99387 | -3.51665 | -1.06283 | - | - |
| OS01G0660200 | Acidic class III chitinase OsChib3a precursor (Chitinase) (EC 3.2.1.14). (Os01t0660200-01) | -1.98831 | -1.79195 | - | - | - |
| OS07G0539900 | Similar to Beta-1,3-glucanase-like protein. (Os07t0539900-01) | -1.98554 | -1.87227 | - | - | - |
| OS11G0702400 | Zinc finger, C2H2-type domain containing protein. (Os11t0702400-01) | -1.97594 | -3.96894 | -1.92901 | - | - |
| OS01G0695200 | Protein of unknown function DUF266, plant family protein. (Os01t0695200-01) | -1.9728 | -1.73098 | -0.780444 | 1.34885 | 1.80244 |
| OS12G0569500 | Thaumatin, pathogenesis-related family protein. (Os12t0569500-01) | -1.96421 | -1.14234 | 0.669619 | 3.54066 | 2.60169 |
| OS04G0468600 | Hypothetical conserved gene. (Os04t0468600-01);Heavy metal transport/detoxification protein domain containing protein. (Os( | -1.96277 | -1.24465 | - | - | -3.15701 |
| OS10G0463800 | Domain of unknown function, DUF1338, containing green-plant-unique protein, Regulation of starch synthesis and amyloplast d | -1.9624 | -1.44076 | - | 3.30293 | 2.81222 |
| OS03G0580200 | Lipase, GDSL domain containing protein. (Os03t0580200-01) | -1.92745 | -1.12771 | -0.877818 | 1.18819 | - |
| OS02G0321000 | Pentatricopeptide repeat domain containing protein. (Os02t0321000-01) | -1.92214 | -1.79926 | -1.2743 | - | - |
| OS12G0217800 | Hypothetical gene. (Os12t0217800-01) | -1.87038 | -2.21497 | - | - | - |
| OS09G0272600 | Conserved hypothetical protein. (Os09t0272600-01) | -1.85172 | -1.54768 | 2.8115 | 3.27245 | 3.23559 |
| OS06G0140200 | Leucine-rich repeat, plant specific containing protein. (Os06t0140200-01) | -1.84692 | -2.06209 | - | 1.52944 | - |
| OS10G0569400 | RIR1a protein precursor. (Os10t0569400-01) | -1.83196 | -1.64854 | - | - | - |
| OS07G0162400 | Alpha/beta hydrolase fold-3 domain containing protein. (Os07t0162400-01) | -1.8305 | -1.18788 | -2.20651 | 1.07134 | - |
| OS03G0132900 | Similar to Chitinase 11. (Os03t0132900-01) | -1.82481 | -1.29056 | 0.592808 | - | - |
| OS06G0579100 | Conserved hypothetical protein. (Os06t0579100-01) | -1.79503 | -2.12852 | - | 1.07275 | - |
| OS08G0140300 | Aromatic L-amino acid decarboxylase (AADC), Senescence-induced serotonin biosynthesis (Os08t0140300-01) | -1.79313 | -4.09093 | -1.51986 | - | - |
| OS01G0175700 | Boron (B) transporter (Os01t0175700-01) | -1.78697 | -1.67997 | -1.90044 | - | - |
| OS06G0591400 | Conserved hypothetical protein. (Os06t0591400-01) | -1.73534 | -1.87305 | - | - | - |
| OS07G0190000 | Similar to 1-deoxy-D-xylulose 5-phosphate synthase 2 precursor. (Os07t0190000-01) | -1.73482 | -2.75246 | -1.42572 | - | - |
| OS03G0773000 | Protein of unknown function DUF1005 family protein. (Os03t0773000-01) | -1.72485 | -1.35082 | -3.63242 | 1.19021 | - |
| OS04G0167800 | Similar to Chalcone reductase homologue (Fragment). (Os04t0167800-01) | -1.72271 | -1.29311 | - | 1.58518 | 1.63124 |
| OS06G0582600 | Similar to Cysteine proteinase. (Os06t0582600-01) | -1.7208 | -1.22887 | 0.903223 | 1.0386 | - |
| OS01G0940800 | Similar to Beta-1,3-glucanase precursor. (Os01t0940800-01) | -1.71364 | -3.07042 | 0.726988 | - | - |
| OS07G0601900 | Similar to NADPH HC toxin reductase (Fragment). (Os07t0601900-01) | -1.70105 | -2.25909 | 1.67838 | - | - |
| OS12G0555200 | Similar to Probenazole-inducible protein PBZ1. (Os12t0555200-01) | -1.69959 | -4.50082 | - | - | 2.38069 |
| OS01G0124000 | Similar to Bowman Birk trypsin inhibitor. (Os01t0124000-01) | -1.68054 | -0.954225 | 4.2157 | 3.70393 | 4.27422 |
| OS07G0283050 | Concanavalin A-like lectin/glucanase, subgroup domain containing protein. (Os07t0283050-00) | -1.6747 | -1.21225 | -0.947544 | - | - |
| OS05G0322900 | WRKY transcription factor, Benzothiadiazole (BTH)-inducible blast resistance (Os05t0322900-01);Similar to WRKY transcription f | -1.66278 | -1.30225 | -1.53846 | - | -1.09877 |
| OS08G0371200 | Carotenoid oxygenase domain containing protein. (Os08t0371200-01) | -1.65839 | -1.37755 | -0.580422 | - | - |
| OS10G0552400 | Zinc finger, RING/FYVE/PHD-type domain containing protein. (Os10t0552400-01) | -1.64808 | -1.48651 | 1.32809 | 1.26009 | 1.50595 |

| | | | | | | |
|---|---|---|---|---|---|---|
| OS01G0623500 | ATPase, AAA-type, core domain containing protein. (Os01t0623500-01) | -1.64778 | -1.07732 | - | - | - |
| OS02G0111600 | Serine/threonine protein kinase-related domain containing protein. (Os02t0111600-01) | -1.64412 | -1.18323 | -0.789693 | - | - |
| OS12G0508400 | Conserved hypothetical protein. (Os12t0508400-00) | -1.63847 | -3.56594 | - | - | - |
| OS06G0586000 | Conserved hypothetical protein. (Os06t0586000-01) | -1.60714 | -2.20091 | 0.78487 | 1.30848 | 1.61516 |
| OS08G0335500 | Hypothetical conserved gene. (Os08t0335500-01) | -1.60435 | -1.61404 | 0.93029 | - | - |
| OS06G0141166 | Similar to hydrolase, NUDIX family protein. (Os06t0141166-00) | -1.59114 | -1.20635 | -1.73803 | 0.979754 | - |
| OS03G0177300 | Zinc finger, RING/FYVE/PHD-type domain containing protein. (Os03t0177300-01) | -1.58541 | -1.40848 | -2.08034 | 1.17078 | - |
| OS08G0375400 | Plant disease resistance response protein family protein. (Os08t0375400-01) | -1.58319 | -0.842383 | -2.1804 | 1.33815 | - |
| OS04G0182200 | 2OG-Fe(II) oxygenase domain containing protein. (Os04t0182200-01) | -1.57587 | -0.868122 | - | - | - |
| OS07G0605400 | Similar to EGG APPARATUS-1 protein (ZmEA1). (Os07t0605400-01);Similar to EGG APPARATUS-1 protein (ZmEA1). (Os07t06054( | -1.57511 | -1.17663 | -1.63179 | - | - |
| OS01G0584900 | WRKY transcription factor 28-like (WRKY5) (WRKY transcription factor 77). (Os01t0584900-01) | -1.57079 | -1.936 | - | - | - |
| OS10G0536450 | Hypothetical protein. (Os10t0536450-00) | -1.56883 | -1.43517 | - | - | - |
| OS03G0269900 | Protein of unknown function DUF604 family protein. (Os03t0269900-01) | -1.5685 | -1.07795 | -0.968211 | 2.04926 | 1.30654 |
| OS01G0256500 | Similar to ZnI. (Os01t0256500-02) | -1.55906 | -0.99182 | - | 3.93591 | 2.69842 |
| OS07G0117000 | NB-ARC domain containing protein. (Os07t0117000-01) | -1.55393 | -1.78369 | -1.72996 | - | - |
| OS05G0583500 | Phox-associated domain domain containing protein. (Os05t0583500-01);Hypothetical conserved gene. (Os05t0583500-02) | -1.53196 | -1.47583 | - | 1.09812 | - |
| OS09G0448200 | Similar to High-affinity potassium transporter. (Os09t0448200-01);Similar to High-affinity potassium transporter. (Os09t044820( | -1.53075 | -1.4922 | 0.526047 | 0.968802 | 1.01197 |
| OS04G0149400 | Conserved hypothetical protein. (Os04t0149400-01) | -1.52575 | -1.54904 | - | - | - |
| OS08G0261000 | NB-ARC domain containing protein. (Os08t0261000-01) | -1.5197 | -1.97165 | - | - | - |
| OS10G0359500 | No apical meristem (NAM) protein domain containing protein. (Os10t0359500-01) | -1.50973 | -1.84623 | -0.949986 | 1.18071 | 1.41515 |
| OS10G0491400 | Conserved hypothetical protein. (Os10t0491400-01) | -1.50521 | -1.40459 | - | - | - |
| OS05G0111000 | Similar to Gag polyprotein [Contains: Core protein p15 (Matrix protein); Core protein p24; Core protein p12]. (Os05t0111000-01 | -1.50133 | -1.23942 | - | - | - |
| OS04G0581100 | 2OG-Fe(II) oxygenase domain containing protein. (Os04t0581100-01) | -1.49873 | -2.11393 | -0.842091 | #N/A | #N/A |
| OS03G0174300 | Exostosin-like family protein. (Os03t0174300-01) | -1.49839 | -1.56525 | #N/A | #N/A | #N/A |
| OS07G0217600 | CytochromeP450 monooxygenase, Diterpenoid phytoalexin biosynthesis, Bacterial blight resistance (Os07t0217600-01) | -1.49545 | -1.48338 | -0.474139 | #N/A | #N/A |
| OS05G0217800 | BURP domain containing protein. (Os05t0217800-01);Similar to BURP domain-containing protein 2. (Os05t0217800-02) | -1.49136 | -0.919721 | -1.93499 | 1.94445 | #N/A |
| OS10G0137300 | Similar to BLN1-1. (Os10t0137300-00) | -1.49094 | -0.917507 | -0.830355 | #N/A | #N/A |
| OS07G0633500 | Hypothetical conserved gene. (Os07t0633500-00) | -1.48158 | -1.3997 | -2.11033 | #N/A | #N/A |
| OS11G0541600 | Protein of unknown function DUF247, plant domain containing protein. (Os11t0541600-00) | -1.47982 | -1.40079 | #N/A | 1.53469 | #N/A |
| OS10G0173000 | Conserved hypothetical protein. (Os10t0173000-01) | -1.47836 | -0.888413 | -1.14913 | 2.75266 | 0.973211 |
| OS09G0537700 | S-like ribonuclease, Salinity tolerance, Abiotic stress response, Regulation of photomorphogenesis (Os09t0537700-02) | -1.46442 | -0.873958 | 2.2403 | 2.06698 | 1.77198 |
| OS10G0531400 | Glutathione S-transferase GST 30 (EC 2.5.1.18). (Os10t0531400-01) | -1.46025 | -1.10839 | -1.84618 | #N/A | #N/A |
| OS01G0314800 | Late embryogenesis abundant protein 3 family protein. (Os01t0314800-01) | -1.4599 | -1.00771 | -1.11407 | 2.13673 | 1.49884 |
| OS04G0142400 | Conserved hypothetical protein. (Os04t0142400-01) | -1.45443 | -1.30629 | 0.834992 | 1.32408 | 1.28829 |
| OS09G0355400 | Protein kinase, core domain containing protein. (Os09t0355400-01);Similar to OsD305. (Os09t0355400-02) | -1.4514 | -1.01956 | #N/A | #N/A | -0.975336 |
| OS10G0497700 | Similar to Phytochelatin synthetase. (Os10t0497700-01);Similar to COBRA-like protein 4. (Os10t0497700-02) | -1.45015 | -1.90007 | -0.701281 | #N/A | #N/A |
| OS08G0530300 | Exo70 exocyst complex subunit family protein. (Os08t0530300-01) | -1.44009 | -1.00062 | -2.52394 | #N/A | #N/A |
| OS10G0535800 | Protein of unknown function Cys-rich domain containing protein. (Os10t0535800-01) | -1.43813 | -2.17533 | #N/A | 1.28237 | 2.11766 |
| OS08G0128300 | Similar to Actin-like protein 3 (Actin-related protein 3). (Os08t0128300-01) | -1.43454 | -1.20611 | -1.22143 | #N/A | #N/A |
| OS01G0795200 | Similar to Subtilase. (Os01t0795200-01) | -1.43239 | -1.78089 | #N/A | #N/A | #N/A |
| OS04G0206700 | UDP-glucuronosyl/UDP-glucosyltransferase family protein. (Os04t0206700-01);UDP-glucuronosyl/UDP-glucosyltransferase fami | -1.43234 | -1.1124 | #N/A | 1.54175 | #N/A |
| OS03G0301200 | COBRA-like protein 7 precursor. (Os03t0301200-01);Glycosyl-phosphatidyl inositol-anchored, plant domain containing protein. ( | -1.43033 | -1.8932 | -1.7185 | #N/A | #N/A |
| OS11G0514500 | Sorghum bicolor leucine-rich repeat-containing extracellular glycoprotein precursor. (Os11t0514500-01) | -1.42816 | -3.2422 | #N/A | 1.09952 | 3.45449 |
| OS04G0307500 | Similar to OSIGBa0113B06.2 protein. (Os04t0307500-01) | -1.42524 | -0.935529 | -0.598636 | #N/A | #N/A |
| OS03G0666100 | Similar to GN (GNOM); GTP:GDP antiporter/ protein homodimerization. (Os03t0666100-00) | -1.42428 | -1.39731 | #N/A | #N/A | 1.18285 |

| | | | | | | |
|---|---|---|---|---|---|---|
| OS01G0864500 | Harpin-induced 1 domain containing protein. (Os01t0864500-01) | -1.42314 | -1.11039 | #N/A | #N/A | #N/A |
| OS08G0384500 | ABC transporter-like domain containing protein. (Os08t0384500-01) | -1.41905 | -1.11285 | -1.18443 | #N/A | #N/A |
| OS10G0469700 | Leucine-rich repeat, typical subtype containing protein. (Os10t0469700-01) | -1.41859 | -1.10181 | 0.732778 | #N/A | #N/A |
| OS01G0615100 | Similar to Substilin /chymotrypsin-like inhibitor (Proteinase inhibitor). (Os01t0615100-01) | -1.41329 | -2.12569 | 2.32488 | 2.39173 | 3.72562 |
| OS02G0748300 | Similar to VMP3 protein. (Os02t0748300-01);Similar to Kelch motif family protein. (Os02t0748300-02) | -1.41286 | -1.12521 | #N/A | 1.11929 | 1.36644 |
| OS06G0323100 | Similar to H1005F08.18 protein. (Os06t0323100-01) | -1.39586 | -2.75428 | 0.889409 | #N/A | 2.2705 |
| OS03G0225900 | Allene oxide synthase (CYP74A2), Biosynthesis of jasmonic acid (JA) (Os03t0225900-01);Allene oxide synthase. (Os03t0225900-0 | -1.38912 | -1.56245 | -0.975954 | #N/A | 1.42565 |
| OS10G0536400 | Similar to Oxidoreductase, 2OG-Fe oxygenase family protein, expressed. (Os10t0536400-01) | -1.38677 | -1.40543 | #N/A | #N/A | #N/A |
| OS07G0414900 | Similar to Transposable element Ac. (Os07t0414900-01);Similar to Transposase (Fragment). (Os07t0414900-02) | -1.37982 | -1.62777 | #N/A | #N/A | #N/A |
| OS02G0581300 | TRAM, LAG1 and CLN8 homology domain containing protein. (Os02t0581300-01) | -1.3769 | -0.943967 | -1.10588 | 0.861282 | #N/A |
| OS03G0773300 | Protein kinase, core domain containing protein. (Os03t0773300-01);Similar to wound and phytochrome signaling involved recep | -1.37499 | -1.95667 | #N/A | #N/A | 2.0919 |
| OS08G0562600 | C2 calcium-dependent membrane targeting domain containing protein. (Os08t0562600-03) | -1.36723 | -0.994625 | -2.36447 | #N/A | #N/A |
| OS02G0170000 | Conserved hypothetical protein. (Os02t0170000-01);Pentatricopeptide repeat domain containing protein. (Os02t0170000-02) | -1.36274 | -0.713721 | -1.9883 | 1.25615 | #N/A |
| OS02G0684300 | Nucleoporin protein Ndc1-Nup domain containing protein. (Os02t0684300-01) | -1.36148 | -1.1206 | -1.0558 | #N/A | #N/A |
| OS07G0669200 | Similar to GTP1/OBG family protein. (Os07t0669200-00) | -1.34974 | -0.952073 | -1.78613 | 1.0823 | #N/A |
| OS01G0894000 | Metallophosphoesterase domain containing protein. (Os01t0894000-01);Similar to predicted protein. (Os01t0894000-02) | -1.34307 | -1.25839 | #N/A | #N/A | #N/A |
| OS07G0673000 | Ureohydrolase domain containing protein. (Os07t0673000-01) | -1.34045 | -1.09579 | -2.74183 | 1.35125 | #N/A |
| OS01G0551100 | Ribonuclease III domain containing protein. (Os01t0551100-01) | -1.33678 | -1.12324 | -0.720488 | 1.02391 | #N/A |
| OS08G0127100 | Amino acid transporter, transmembrane family protein. (Os08t0127100-01);Amino acid transporter, transmembrane family pro | -1.33474 | -1.19686 | #N/A | #N/A | #N/A |
| OS09G0417600 | WRKY transcription factor, Transcriptional repressor, Pathogen defense (Os09t0417600-01);WRKY transcription factor 76. (Os09 | -1.32912 | -1.47949 | #N/A | #N/A | #N/A |
| OS04G0423400 | Similar to OSIGBa0076I14.3 protein. (Os04t0423400-01);Similar to OSIGBa0076I14.3 protein. (Os04t0423400-02) | -1.32033 | -0.553876 | -0.950882 | 1.49656 | #N/A |
| OS08G0508800 | Lipoxygenase, chloroplast precursor (EC 1.13.11.12). (Os08t0508800-01) | -1.30968 | -1.6826 | #N/A | #N/A | #N/A |
| OS10G0416500 | Similar to Chitinase 1 precursor (EC 3.2.1.14) (Tulip bulb chitinase-1) (TBC-1). (Os10t0416500-04) | -1.29475 | -2.07662 | #N/A | #N/A | #N/A |
| OS04G0298600 | Uncharacterised protein family FPL domain containing protein. (Os04t0298600-00) | -1.29196 | -0.797827 | -1.56465 | 1.11968 | #N/A |
| OS06G0319600 | Poly(A) polymerase, central domain domain containing protein. (Os06t0319600-01) | -1.2786 | -1.08987 | #N/A | #N/A | #N/A |
| OS10G0450900 | Similar to Glycine-rich cell wall structural protein 2 precursor. (Os10t0450900-02) | -1.27442 | -0.901957 | 1.33275 | 1.14854 | 1.30784 |
| OS11G0184900 | Similar to OsNAC5 protein [imported]-rice. (Os11t0184900-01);NAC-type transcription factor, Regulation of stress-inducible gen | -1.26126 | -0.934547 | -1.2575 | 1.22857 | #N/A |
| OS04G0401000 | Proline-rich protein, Blast resistance (Os04t0401000-01);Similar to Pi21 protein. (Os04t0401000-02) | -1.25764 | -0.915853 | -1.45011 | 1.36688 | #N/A |
| OS12G0136900 | Similar to Calcium-transporting ATPase 4, plasma membrane-type (EC 3.6.3.8) (Ca(2+)-ATPase isoform 4). (Os12t0136900-00) | -1.25716 | -1.61977 | #N/A | #N/A | #N/A |
| OS07G0295800 | Similar to peptidase. (Os07t0295800-01) | -1.25318 | -1.04614 | #N/A | #N/A | #N/A |
| OS01G0807900 | Similar to Dihydropyrimidinase (Dihydropyrimidine amidohydrolase) (EC 3.5.2.2). (Os01t0807900-01);D-hydantoinase domain co | -1.25017 | -1.13606 | #N/A | 1.55258 | 1.53259 |
| OS05G0443300 | Similar to predicted protein. (Os05t0443300-01) | -1.24871 | -0.889494 | -0.589663 | 1.04855 | #N/A |
| OS10G0130500 | Hypothetical protein. (Os10t0130500-01);Hypothetical protein. (Os10t0130500-02) | -1.24258 | -0.851958 | -1.64856 | 1.17144 | #N/A |
| OS07G0173400 | Conserved hypothetical protein. (Os07t0173400-01) | -1.23906 | -0.884565 | #N/A | 0.924889 | #N/A |
| OS07G0677100 | Peroxidase. (Os07t0677100-01) | -1.2195 | -1.72968 | 2.21888 | 1.10616 | 2.12963 |
| OS08G0503800 | Beta 1,2-xylosyltransferase, Response to abiotic stresses and phytohormones (Os08t0503800-01) | -1.2174 | -1.09576 | -1.0142 | #N/A | #N/A |
| OS07G0575800 | Similar to hydrolase, alpha/beta fold family protein. (Os07t0575800-01) | -1.21447 | -1.19698 | -1.60844 | 0.939939 | #N/A |
| OS10G0470700 | Similar to Peptide transporter. (Os10t0470700-01) | -1.2124 | -1.48957 | #N/A | #N/A | #N/A |
| OS01G0713200 | Similar to Beta-glucanase. (Os01t0713200-01) | -1.20919 | -1.01894 | #N/A | #N/A | #N/A |
| OS09G0407900 | Peptidase C19, ubiquitin carboxyl-terminal hydrolase 2 family protein. (Os09t0407900-01) | -1.20831 | -1.08383 | #N/A | #N/A | #N/A |
| OS04G0688500 | Peroxidase (EC 1.11.1.7). (Os04t0688500-01) | -1.20635 | -0.978508 | #N/A | #N/A | #N/A |
| OS01G0602400 | Similar to Cytochrome P450 monooxygenase CYP72A5 (Fragment). (Os01t0602400-01) | -1.20413 | -1.33412 | #N/A | 1.227 | 1.49005 |
| OS09G0262000 | Similar to Cinnamoyl CoA reductase. (Os09t0262000-00) | -1.20191 | -1.27271 | 2.0696 | #N/A | #N/A |
| OS03G0703900 | SNARE associated Golgi protein domain containing protein. (Os03t0703900-01) | -1.20045 | -1.01679 | -1.18616 | #N/A | #N/A |
| OS05G0406000 | Conserved hypothetical protein. (Os05t0406000-01) | -1.19943 | -0.64504 | -1.7268 | 0.875884 | #N/A |

| | | | | | | |
|---|---|---|---|---|---|---|
| OS01G0921400 | Conserved hypothetical protein. (Os01t0921400-01) | -1.19508 | -1.06968 | -1.05337 | #N/A | #N/A |
| OS08G0395700 | Conserved hypothetical protein. (Os08t0395700-01) | -1.19387 | -1.06201 | 3.47867 | 2.67344 | 2.98476 |
| OS02G0763900 | Similar to Galactoside 2-alpha-L-fucosyltransferase (EC 2.4.1.69) (Xyloglucan alpha-(1,2)-fucosyltransferase) (AtFUT1). (Os02t076 | -1.19201 | -1.0356 | -1.22805 | #N/A | #N/A |
| OS07G0692900 | Similar to Ubiquitin-activating enzyme E1. (Os07t0692900-01) | -1.18791 | -1.37457 | #N/A | #N/A | #N/A |
| OS04G0127200 | Hypothetical conserved gene. (Os04t0127200-01);Similar to Subtilase. (Os04t0127200-02) | -1.18529 | -1.53791 | -0.92271 | #N/A | #N/A |
| OS03G0351100 | Similar to CONSTANS-like protein CO9 (Fragment). (Os03t0351100-01);Similar to CONSTANS-like protein CO9 (Fragment). (Os03 | -1.18067 | -0.752286 | -1.22746 | 1.12455 | #N/A |
| OS10G0508700 | Pectinesterase inhibitor domain containing protein. (Os10t0508700-01) | -1.17874 | -0.940467 | -4.85304 | -0.648523 | -1.32535 |
| OS09G0550000 | Bromodomain containing protein. (Os09t0550000-01) | -1.17799 | -1.08312 | -0.861777 | #N/A | #N/A |
| OS10G0457700 | Similar to chromatin remodeling complex subunit. (Os10t0457700-01) | -1.17355 | -1.05797 | -1.03816 | #N/A | #N/A |
| OS11G0266800 | Tetratricopeptide-like helical domain containing protein. (Os11t0266800-01) | -1.14887 | -1.56046 | -0.785848 | #N/A | 1.35424 |
| OS12G0209000 | Similar to sporulation protein-related. (Os12t0209000-00) | -1.12787 | -1.01175 | -2.3614 | #N/A | #N/A |
| OS09G0536700 | Similar to predicted protein. (Os09t0536700-01);Nodulin-like domain containing protein. (Os09t0536700-02) | -1.12375 | -0.701328 | -1.27547 | #N/A | #N/A |
| OS08G0309300 | RER1A protein (AtRER1A). (Os08t0309300-01) | -1.12073 | -0.945429 | -1.23336 | 1.31834 | #N/A |
| OS06G0644500 | Ankyrin domain containing protein. (Os06t0644500-01);Ankyrin domain containing protein. (Os06t0644500-02) | -1.10157 | -0.859611 | #N/A | #N/A | #N/A |
| OS04G0340300 | Terpenoid synthase domain containing protein. (Os04t0340300-01) | -1.09718 | -1.35019 | 2.18881 | 1.19147 | #N/A |
| OS02G0536500 | Similar to H0622F05.5 protein. (Os02t0536500-01) | -1.08911 | -1.22128 | #N/A | #N/A | 1.46283 |
| OS03G0574900 | Similar to Potassium transporter 1 (OsHAK1). Splice isoform 2. (Os03t0574900-01) | -1.08593 | -0.893536 | 0.543332 | 0.932171 | #N/A |
| OS02G0302200 | Similar to aminotransferase family protein. (Os02t0302200-01) | -1.0837 | -0.921307 | -2.36557 | 1.40388 | #N/A |
| OS12G0547600 | Calmodulin binding protein-like domain containing protein. (Os12t0547600-01);Hypothetical conserved gene. (Os12t0547600-0 | -1.07897 | -2.01017 | #N/A | #N/A | #N/A |
| OS09G0103200 | Conserved hypothetical protein. (Os09t0103200-01) | -1.07372 | -0.887344 | -1.42903 | #N/A | #N/A |
| OS09G0498000 | Hypothetical gene. (Os09t0498000-01) | -1.0717 | -2.28434 | #N/A | 0.903562 | 2.63361 |
| OS12G0583300 | Peptidase aspartic, catalytic domain containing protein. (Os12t0583300-01) | -1.06072 | -0.821148 | -0.669531 | #N/A | 0.78528 |
| OS04G0121800 | Hypothetical conserved gene. (Os04t0121800-01) | -1.05852 | -1.68932 | 1.14202 | #N/A | #N/A |
| OS11G0474900 | Similar to Stemar-13-ene synthase. (Os11t0474900-00) | -1.0580047 | -1.0482501 | #N/A | #N/A | #N/A |
| OS08G0451000 | Sec34-like protein family protein. (Os08t0451000-01) | -1.05739 | -1.04276 | -0.774146 | #N/A | #N/A |
| OS12G0613700 | Transcriptional factor B3 family protein. (Os12t0613700-01);Similar to Auxin response factor 25. (Os12t0613700-02) | -1.05492 | -0.787575 | #N/A | #N/A | #N/A |
| OS12G0123500 | Similar to Apyrase precursor (EC 3.6.1.5) (ATP-diphosphatase) (Adenosine diphosphatase) (ADPase) (ATP-diphosphohydrolase). | -1.04558 | -1.01149 | 0.950442 | #N/A | #N/A |
| OS05G0427400 | Similar to Phenylalanine ammonia-lyase. (Os05t0427400-00) | -1.04557 | -1.32401 | -0.866015 | #N/A | #N/A |
| OS09G0525300 | Cyclin-like F-box domain containing protein. (Os09t0525300-01) | -1.04307 | -0.749105 | -1.46185 | 0.935387 | #N/A |
| OS03G0363900 | Zinc finger, DHHC-type domain containing protein. (Os03t0363900-01);Zinc finger, DHHC-type domain containing protein. (Os03 | -1.04276 | -0.686895 | #N/A | 0.701678 | #N/A |
| OS08G0408500 | Pathogenesis-related transcriptional factor and ERF domain containing protein. (Os08t0408500-01) | -1.03954 | -1.00673 | #N/A | 1.75972 | 1.57005 |
| OS09G0565500 | Conserved hypothetical protein. (Os09t0565500-01) | -1.03558 | -0.917278 | #N/A | #N/A | #N/A |
| OS03G0184100 | Conserved hypothetical protein. (Os03t0184100-01) | -1.0301 | -0.582019 | #N/A | #N/A | #N/A |
| OS03G0669000 | DEAD-box RNA helicase, Regulation of thermotolerant growth, rRNA homeostasis at high temperature (Os03t0669000-01);Simil | -1.02934 | -1.05239 | -1.22357 | #N/A | #N/A |
| OS01G0952700 | Metal-dependent hydrolase, composite domain containing protein. (Os01t0952700-01) | -1.02492 | -1.22453 | #N/A | #N/A | #N/A |
| OS02G0122200 | Similar to predicted protein. (Os02t0122200-00) | -1.01234 | -1.34175 | #N/A | #N/A | 1.35739 |
| OS01G0963000 | Similar to Peroxidase BP 1 precursor. (Os01t0963000-01);Similar to Peroxidase BP 1 precursor. (Os01t0963000-04) | -1.01131 | -1.1826 | -1.74324 | -1.19766 | -1.52164 |
| OS04G0634700 | Similar to Diacylglycerol kinase. (Os04t0634700-01) | -1.01023 | -1.7537 | -1.10046 | #N/A | #N/A |
| OS01G0190000 | Similar to oxidoreductase. (Os01t0190000-01) | -1.00811 | -0.671165 | -1.70326 | 1.98432 | 0.9168 |
| OS03G0431800 | Conserved hypothetical protein. (Os03t0431800-00) | -1.0036 | -1.03059 | #N/A | #N/A | #N/A |
| OS03G0363400 | Uncharacterised protein family UPF0153 domain containing protein. (Os03t0363400-01) | -0.996852 | -0.921394 | -1.98783 | #N/A | #N/A |
| OS06G0361500 | Similar to cDNA clone:J013000C15, full insert sequence. (Os06t0361500-02) | -0.995024 | -0.778367 | #N/A | #N/A | 0.952233 |
| OS08G0542000 | Similar to Methionine aminopeptidase. (Os08t0542000-01) | -0.994532 | -1.31741 | #N/A | #N/A | #N/A |
| OS01G0290800 | Hypothetical protein. (Os01t0290800-01) | -0.990258 | -0.98033 | #N/A | #N/A | #N/A |
| OS09G0351700 | Protein kinase, catalytic domain domain containing protein. (Os09t0351700-00) | -0.987151 | -1.03176 | -2.36211 | -1.82898 | -1.50426 |

| | | | | | | |
|---|---|---|---|---|---|---|
| OS09G0480600 | Conserved hypothetical protein. (Os09t0480600-01) | -0.979285 | -0.978426 | -1.60401 | 1.3081 | #N/A |
| OS03G0218400 | Similar to Hexose transporter. (Os03t0218400-01) | -0.974867 | -1.59291 | 0.694098 | #N/A | #N/A |
| OS03G0772600 | Similar to Lectin-like receptor kinase 7;2. (Os03t0772600-01) | -0.974516 | -1.33448 | #N/A | #N/A | #N/A |
| OS03G0131400 | Polynucleotide adenylyltransferase region domain containing protein. (Os03t0131400-01) | -0.958662 | -1.06186 | -1.37373 | #N/A | #N/A |
| OS06G0701300 | Similar to ABC1 family protein. (Os06t0701300-01);Beta-lactamase family protein. (Os06t0701300-02);Similar to ABC1 family pr | -0.954298 | -0.711944 | #N/A | #N/A | #N/A |
| OS10G0444700 | Phosphate transporter, Pi homeostasis (Os10t0444700-01);Similar to Phosphate transporter 6. (Os10t0444700-02) | -0.950571 | -0.860961 | 0.641191 | #N/A | 0.854823 |
| OS05G0156800 | Conserved hypothetical protein. (Os05t0156800-01) | -0.949337 | -1.00802 | #N/A | 1.18059 | 0.972953 |
| OS10G0160000 | Similar to Ubiquitin carboxyl-terminal hydrolase 12 (EC 3.1.2.15) (Ubiquitin thiolesterase 12) (Ubiquitin-specific processing prote | -0.948304 | -0.908146 | #N/A | 0.780188 | #N/A |
| OS01G0279400 | Major facilitator superfamily antiporter. (Os01t0279400-01) | -0.945692 | -0.706929 | -1.75328 | #N/A | -1.07695 |
| OS11G0633500 | Similar to NBS-LRR disease resistance protein homologue (Fragment). (Os11t0633500-01) | -0.945563 | -1.22921 | -0.878437 | #N/A | #N/A |
| OS07G0582400 | Similar to Proton myo-inositol cotransporter. (Os07t0582400-01) | -0.940488 | -0.896653 | #N/A | #N/A | #N/A |
| OS03G0135600 | Similar to Ankyrin repeat protein. (Os03t0135600-01) | -0.933318 | -0.755165 | -1.83471 | #N/A | #N/A |
| OS12G0435200 | Similar to Zeaxanthin cleavage oxygenase. (Os12t0435200-01) | -0.933316 | -1.005 | 0.620553 | 1.21195 | 0.976536 |
| OS02G0550000 | Conserved hypothetical protein. (Os02t0550000-01) | -0.929533 | -0.780293 | -0.61341 | #N/A | #N/A |
| OS05G0515200 | Cytochrome P450 family protein. (Os05t0515200-01) | -0.927849 | -1.21756 | #N/A | 1.45008 | 1.95256 |
| OS08G0545700 | Similar to TraB protein-related. (Os08t0545700-01) | -0.924907 | -0.679067 | -2.25418 | 0.979422 | #N/A |
| OS01G0871800 | TGF-beta receptor, type I/II extracellular region family protein. (Os01t0871800-01) | -0.919548 | -0.898101 | #N/A | #N/A | #N/A |
| OS03G0708600 | DEAD-like helicase, N-terminal domain containing protein. (Os03t0708600-01);DEAD-like helicase, N-terminal domain containin | -0.918301 | -0.915406 | -0.956719 | #N/A | #N/A |
| OS05G0311500 | Protein of unknown function DUF567 family protein. (Os05t0311500-01) | -0.917427 | -0.674233 | -1.26879 | 1.07172 | 0.916656 |
| OS07G0656700 | Uncharacterised conserved protein UCP022348 domain containing protein. (Os07t0656700-01) | -0.913853 | -0.764983 | -1.64185 | 1.46558 | #N/A |
| OS10G0577900 | Similar to Glycerol-3-phosphate acyltransferase (Fragment). (Os10t0577900-01) | -0.913521 | -0.725819 | -1.7792 | 0.817608 | #N/A |
| OS01G0918500 | Conserved hypothetical protein. (Os01t0918500-01) | -0.912727 | -1.19423 | -0.846761 | #N/A | #N/A |
| OS06G0215400 | Peptidase S9, prolyl oligopeptidase active site region domain containing protein. (Os06t0215400-01) | -0.91226 | -1.5053 | #N/A | #N/A | 1.32594 |
| OS09G0542100 | Peptidase A1 domain containing protein. (Os09t0542100-01) | -0.90822 | -0.703524 | -0.690676 | #N/A | #N/A |
| OS06G0618600 | Rgp1 domain containing protein. (Os06t0618600-01) | -0.904271 | -0.976237 | -0.866829 | #N/A | #N/A |
| OS02G0739900 | Similar to microtubule-associated protein TORTIFOLIA1. (Os02t0739900-01) | -0.902645 | -0.895609 | -0.783375 | #N/A | #N/A |
| OS03G0158200 | DEAD-like helicase, N-terminal domain containing protein. (Os03t0158200-01) | -0.90204 | -0.779837 | -0.978614 | 0.749296 | #N/A |
| OS07G0558000 | ABC-1 domain containing protein. (Os07t0558000-01) | -0.901599 | -0.728091 | -0.946909 | #N/A | #N/A |
| OS10G0415200 | Exocyst complex subunit Sec15-like family protein. (Os10t0415200-01);Exocyst complex subunit Sec15-like family protein. (Os10 | -0.890497 | -0.983497 | #N/A | #N/A | #N/A |
| OS02G0704600 | Conserved hypothetical protein. (Os02t0704600-01) | -0.888813 | -0.963563 | -0.75594 | 1.21782 | 0.892936 |
| OS09G0344800 | Protein of unknown function DUF81 family protein. (Os09t0344800-01);Protein of unknown function DUF81 family protein. (Os0 | -0.887596 | -0.653107 | -0.510574 | 1.02618 | #N/A |
| OS03G0144000 | Conserved hypothetical protein. (Os03t0144000-01) | -0.884374 | -0.941112 | -0.992699 | #N/A | #N/A |
| OS03G0582200 | SCAMP family protein. (Os03t0582200-01);SCAMP family protein. (Os03t0582200-02) | -0.883726 | -1.06338 | 0.532629 | #N/A | 1.2571 |
| OS08G0197500 | Cyclin-like F-box domain containing protein. (Os08t0197500-01) | -0.880109 | -0.813109 | #N/A | #N/A | #N/A |
| OS10G0491000 | Plant Basic Secretory Protein family protein. (Os10t0491000-01) | -0.878434 | -1.11731 | 1.15443 | 1.35118 | 1.97129 |
| OS02G0729700 | Similar to HAHB-7 (Fragment). (Os02t0729700-01);Similar to HAHB-7 (Fragment). (Os02t0729700-02) | -0.871785 | -0.695389 | -0.65887 | 1.10027 | 0.910908 |
| OS02G0611400 | Pentatricopeptide repeat domain containing protein. (Os02t0611400-01) | -0.864631 | -0.96133 | -0.711727 | #N/A | #N/A |
| OS02G0633100 | Hypothetical conserved gene. (Os02t0633100-01);B-block binding subunit of TFIIIC domain containing protein. (Os02t0633100-0 | -0.863908 | -1.15105 | #N/A | #N/A | #N/A |
| OS04G0111500 | Similar to OSIGBa0127D24.4 protein. (Os04t0111500-01);Similar to OSIGBa0127D24.4 protein. (Os04t0111500-02) | -0.859798 | -0.648703 | #N/A | #N/A | #N/A |
| OS11G0242100 | Beta-glucosidase, GBA2 type domain containing protein. (Os11t0242100-01);Beta-glucosidase, GBA2 type domain containing pr | -0.85078 | -1.10635 | #N/A | #N/A | 0.879338 |
| OS11G0607200 | Protein kinase, core domain containing protein. (Os11t0607200-01);Similar to Brassinosteroid insensitive1-associated receptor | -0.850474 | -0.679229 | #N/A | #N/A | #N/A |
| OS05G0159501 | Conserved hypothetical protein. (Os05t0159501-00) | -0.8490686 | -0.8009804 | #N/A | #N/A | #N/A |
| OS05G0440250 | Histone deacetylase superfamily protein. (Os05t0440250-01) | -0.84843 | -0.931772 | #N/A | #N/A | #N/A |
| OS06G0655100 | Similar to D-3-phosphoglycerate dehydrogenase. (Os06t0655100-00) | -0.844516 | -0.870327 | -1.33334 | #N/A | #N/A |
| OS08G0328300 | ENT domain containing protein. (Os08t0328300-01);ENT domain containing protein. (Os08t0328300-02) | -0.84298 | -0.782341 | -0.642813 | #N/A | #N/A |

| | | | | | | |
|---|---|---|---|---|---|---|
| OS08G0107400 | Similar to GDP-mannose transporter. (Os08t0107400-01);Similar to predicted protein. (Os08t0107400-02) | -0.838521 | -0.763583 | #N/A | #N/A | #N/A |
| OS01G0284700 | Similar to Peptidyl-prolyl cis-trans isomerase. (Os01t0284700-01) | -0.836929 | -0.668992 | -2.46758 | #N/A | #N/A |
| OS01G0687400 | Similar to Chitinase (EC 3.2.1.14). (Os01t0687400-01) | -0.835282 | -0.945742 | #N/A | #N/A | #N/A |
| OS09G0376900 | Similar to Potassium transporter 13 (AtPOT13) (AtKT5). (Os09t0376900-01);Similar to Potassium transporter 23. (Os09t0376900 | -0.830342 | -0.728316 | -0.763536 | 1.35586 | #N/A |
| OS11G0454200 | Dehydrin RAB 16B. (Os11t0454200-01) | -0.8282295 | 0.21964869 | 3.02464414 | 5.90849 | 4.87375 |
| OS05G0426200 | No apical meristem (NAM) protein domain containing protein. (Os05t0426200-02) | -0.825592 | -0.511144 | #N/A | #N/A | #N/A |
| OS08G0296600 | NB-ARC domain containing protein. (Os08t0296600-01) | -0.824844 | -1.02406 | #N/A | 0.875058 | #N/A |
| OS10G0502500 | Cytochrome b5 domain containing protein. (Os10t0502500-04) | -0.824409 | -0.800297 | -1.08592 | 1.53562 | #N/A |
| OS02G0725100 | Thioredoxin fold domain containing protein. (Os02t0725100-01) | -0.823329 | -0.724006 | -0.797334 | 0.900375 | #N/A |
| OS08G0301500 | Similar to Sucrose-phosphate synthase 2 (EC 2.4.1.14) (Fragment). (Os08t0301500-01) | -0.822132 | -0.679915 | -0.581387 | 0.863123 | #N/A |
| OS05G0230700 | Similar to Auxin-responsive protein IAA17. (Os05t0230700-01);Similar to Auxin-responsive protein IAA17. (Os05t0230700-02);Si | -0.821764 | -0.663825 | -0.745721 | 1.12838 | 0.84794 |
| OS05G0449600 | Similar to glycosyl hydrolase family 3 protein. (Os05t0449600-01);Hypothetical conserved gene. (Os05t0449600-02) | -0.819718 | -1.11145 | #N/A | 1.21191 | 1.52962 |
| OS02G0730700 | Peptidase A1 domain containing protein. (Os02t0730700-02) | -0.818324 | -0.67116 | #N/A | 0.63791 | 0.953625 |
| OS01G0965900 | Conserved hypothetical protein. (Os01t0965900-01) | -0.818026 | -0.850879 | #N/A | #N/A | 1.07226 |
| OS08G0500800 | WLM domain containing protein. (Os08t0500800-01) | -0.815868 | -0.732718 | #N/A | #N/A | #N/A |
| OS11G0302500 | Similar to cycloeucalenol cycloisomerase. (Os11t0302500-01) | -0.814057 | -0.914195 | -1.27607 | 1.04101 | #N/A |
| OS03G0758900 | Similar to predicted protein. (Os03t0758900-01) | -0.813904 | -0.758007 | -2.51181 | #N/A | #N/A |
| OS03G0670100 | Similar to ATP-binding protein of ABC transporter. (Os03t0670100-01) | -0.813557 | -0.701823 | #N/A | #N/A | #N/A |
| OS01G0338200 | Mov34/MPN/PAD-1 family protein. (Os01t0338200-01) | -0.809631 | -0.696791 | #N/A | #N/A | #N/A |
| OS05G0474900 | Protein of unknown function Cys-rich family protein. (Os05t0474900-01) | -0.808617 | -0.780167 | #N/A | #N/A | #N/A |
| OS01G0734800 | UDP-glucuronosyl/UDP-glucosyltransferase family protein. (Os01t0734800-01);UDP-glucuronosyl/UDP-glucosyltransferase famil | -0.807666 | -1.11795 | -0.545 | #N/A | #N/A |
| OS05G0557200 | Armadillo-type fold domain containing protein. (Os05t0557200-01);Armadillo-type fold domain containing protein. (Os05t0557 | -0.806744 | -0.684898 | 0.851835 | 1.68834 | 1.64121 |
| OS01G0769000 | Topoisomerase II-associated protein PAT1 domain containing protein. (Os01t0769000-01);Similar to predicted protein. (Os01t07 | -0.801624 | -0.594138 | 0.738171 | 1.16091 | 0.826832 |
| OS01G0663400 | Similar to Aspartic proteinase oryzasin 1 precursor (EC 3.4.23.-). (Os01t0663400-01);Similar to aspartic proteinase oryzasin-1. (O | -0.798204 | -0.818086 | #N/A | #N/A | #N/A |
| OS03G0138000 | Conserved hypothetical protein. (Os03t0138000-01) | -0.792868 | -1.03549 | #N/A | #N/A | #N/A |
| OS03G0756700 | Hypothetical conserved gene. (Os03t0756700-01) | -0.790585 | -1.06109 | #N/A | #N/A | #N/A |
| OS02G0649800 | Similar to Cytochrome B5 homolog (Fragment). (Os02t0649800-01) | -0.78661 | -0.870108 | #N/A | #N/A | #N/A |
| OS12G0600701 | Conserved hypothetical protein. (Os12t0600701-00) | -0.780583 | -0.728428 | -2.34617 | 1.13321 | #N/A |
| OS09G0520600 | Bile acid:sodium symporter family protein. (Os09t0520600-01) | -0.773051 | -0.84524 | #N/A | #N/A | #N/A |
| OS07G0172600 | Pentatricopeptide repeat domain containing protein. (Os07t0172600-00) | -0.767778 | -0.847989 | -0.893511 | #N/A | #N/A |
| OS09G0434500 | Similar to Ethylene response factor 2. (Os09t0434500-01) | -0.756307 | -0.596663 | -1.39512 | 0.724982 | #N/A |
| OS06G0726200 | Similar to Chitinase 1. (Os06t0726200-02) | -0.756139 | -1.08374 | 1.7477 | 1.08524 | 2.07279 |
| OS03G0779500 | Hypothetical protein. (Os03t0779500-01) | -0.7511252 | -0.9121815 | -0.0596152 | #N/A | #N/A |
| OS04G0616300 | Similar to H0525G02.10 protein. (Os04t0616300-01) | -0.74642 | -0.78805 | #N/A | #N/A | #N/A |
| OS03G0255400 | FAR1 domain containing protein. (Os03t0255400-01) | -0.742954 | -0.703715 | #N/A | #N/A | #N/A |
| OS05G0399800 | Leucine-rich repeat, N-terminal domain containing protein. (Os05t0399800-01) | -0.7393815 | -0.7300704 | #N/A | #N/A | #N/A |
| OS05G0318700 | Similar to Resistance protein candidate (Fragment). (Os05t0318700-01);Similar to Resistance protein candidate (Fragment). (Os0 | -0.732256 | -0.652101 | #N/A | #N/A | #N/A |
| OS09G0467200 | Similar to Glutathione S-transferase GST 23 (EC 2.5.1.18) (Fragment). (Os09t0467200-01) | -0.729178 | -0.58153 | 0.426713 | #N/A | #N/A |
| OS12G0477200 | Hypothetical conserved gene. (Os12t0477200-00) | -0.7229365 | 0.47629873 | #N/A | #N/A | #N/A |
| OS12G0268000 | Similar to Cytochrome P450 71A1 (EC 1.14.-.-) (CYPLXXIA1) (ARP-2). (Os12t0268000-01) | -0.720295 | -0.828197 | #N/A | #N/A | 0.860194 |
| OS04G0116600 | Glucose/ribitol dehydrogenase family protein. (Os04t0116600-01) | -0.719342 | -0.798051 | -2.0974 | #N/A | #N/A |
| OS04G0376300 | Similar to 3-oxoacyl-reductase. (Os04t0376300-01) | -0.718161 | -0.6807 | -1.40214 | 0.796928 | #N/A |
| OS02G0437800 | Similar to vacuolar protein-sorting protein 45. (Os02t0437800-01);Similar to Vacuolar protein-sorting protein 45 homolog (AtVP | -0.716634 | -0.836292 | #N/A | #N/A | #N/A |
| OS08G0559300 | O-linked N-acetylglucosamine transferase, Negative regulator of gibberellin (GA) signaling, Brassinosteroid (BR) synthesis (Os08t | -0.70877 | -0.663467 | #N/A | #N/A | #N/A |
| OS09G0459800 | Similar to ARP protein. (Os09t0459800-01);Similar to ARP protein. (Os09t0459800-02);Similar to NADPH oxidoreductase homolo | -0.706141 | -0.730608 | #N/A | #N/A | #N/A |

| | | | | | | |
|---|---|---|---|---|---|---|
| OS03G0254900 | Zinc finger, RING/FYVE/PHD-type domain containing protein. (Os03t0254900-01);Similar to zinc finger (C3HC4-type RING finger) | -0.705556 | -0.768133 | -0.889506 | #N/A | #N/A |
| OS02G0618200 | Signal transduction response regulator, receiver region domain containing protein. (Os02t0618200-01) | -0.694978 | -0.68013 | -0.685973 | #N/A | #N/A |
| OS08G0511900 | Similar to aminoacylase-1. (Os08t0511900-01) | -0.691644 | -0.772447 | #N/A | 0.867862 | #N/A |
| OS02G0137500 | Similar to histone acetyltransferase. (Os02t0137500-01) | -0.688708 | -0.671618 | #N/A | #N/A | #N/A |
| OS01G0894300 | Similar to Fructokinase 1. (Os01t0894300-01);Fructokinase (Fragment). (Os01t0894300-02) | -0.677874 | -0.787533 | -1.28628 | #N/A | #N/A |
| OS09G0441625 | Similar to flavonoid 3-monooxygenase. (Os09t0441625-00) | -0.6715357 | 0.20467912 | #N/A | #N/A | #N/A |
| OS01G0299100 | Conserved hypothetical protein. (Os01t0299100-00) | -0.6684209 | -0.5042046 | 1.52800113 | #N/A | #N/A |
| OS10G0537400 | Similar to metal ion binding protein. (Os10t0537400-00) | -0.661093 | 1.10877199 | 0.05368135 | #N/A | #N/A |
| OS06G0229200 | Glycosyl transferase, family 31 protein. (Os06t0229200-01) | -0.654307 | -0.728211 | #N/A | #N/A | #N/A |
| OS03G0262900 | Hypothetical conserved gene. (Os03t0262900-01);Activator of Rab5 GTPase, Intracellular transport of the proglutelin (Os03t026 | -0.649955 | -0.867092 | #N/A | #N/A | #N/A |
| OS12G0441300 | O-methyltransferase, family 2 protein. (Os12t0441300-00) | -0.6444153 | -0.3638575 | 1.81071044 | #N/A | #N/A |
| OS07G0187700 | WD40 protein, Regulation of the plasma membrane localization of phosphate transporters, Phosphate uptake and translocation | -0.631597 | -0.582284 | -0.754302 | #N/A | #N/A |
| OS11G0465200 | Similar to Bx2-like protein. (Os11t0465200-01);Similar to Cytochrome P450 family protein, expressed. (Os11t0465200-02) | -0.630538 | -0.885341 | #N/A | 0.899934 | #N/A |
| OS05G0147500 | Similar to DEGP2 (DEGP PROTEASE 2); serine-type peptidase/ trypsin. (Os05t0147500-01);Serine endopeptidase DegP2 domain | -0.620543 | -0.61502 | -1.87369 | #N/A | #N/A |
| OS10G0511400 | Peptidase S28 family protein. (Os10t0511400-01) | -0.603129 | -0.860709 | #N/A | #N/A | 0.783262 |
| OS01G0627500 | Cytochrome P450 family protein. (Os01t0627500-01) | -0.597636 | -0.78511 | -0.689518 | #N/A | #N/A |
| OS05G0241000 | Prefoldin domain containing protein. (Os05t0241000-01) | -0.593441 | -0.661667 | #N/A | #N/A | #N/A |
| OS11G0635500 | Cytochrome P450 family protein. (Os11t0635500-01) | -0.578887 | -0.575394 | #N/A | #N/A | -0.799842 |
| OS07G0558300 | Inositol monophosphatase family protein. (Os07t0558300-01) | -0.565395 | -0.590643 | -1.92104 | #N/A | #N/A |
| OS10G0421800 | Pentatricopeptide repeat domain containing protein. (Os10t0421800-01) | -0.551196 | -0.58065 | -0.905884 | #N/A | #N/A |
| OS01G0895600 | Similar to Calreticulin-3. (Os01t0895600-01) | -0.528156 | -0.500539 | #N/A | #N/A | #N/A |
| OS04G0407500 | Similar to H0321H01.2 protein. (Os04t0407500-00) | -0.4969566 | -0.3360233 | #N/A | #N/A | #N/A |
| OS08G0443800 | Tetraspanin domain containing protein. (Os08t0443800-01) | -0.4834676 | -0.473309 | -0.6256259 | -0.2721963 | #N/A |
| OS03G0658800 | Cytochrome P450 family protein. (Os03t0658800-01);Hypothetical conserved gene. (Os03t0658800-02);Cytochrome P450 family | -0.466751 | -0.582053 | #N/A | #N/A | #N/A |
| OS04G0127100 | Peptidase S8, subtilisin-related domain containing protein. (Os04t0127100-01) | -0.3268594 | -0.3176977 | #N/A | #N/A | #N/A |
| OS01G0126100 | Multicopper oxidase (Os01t0126100-01) | -0.3229853 | 0.12118785 | #N/A | #N/A | #N/A |
| OS11G0417800 | Non-protein coding transcript. (Os11t0417800-01) | -0.2621714 | -0.4018616 | 0.30961859 | #N/A | #N/A |
| OS07G0625400 | SKP1 component domain containing protein. (Os07t0625400-01) | -0.1994326 | -0.6116385 | #N/A | #N/A | #N/A |
| OS02G0769700 | Protein kinase, core domain containing protein. (Os02t0769700-01) | -0.0986755 | -0.5878862 | #N/A | #N/A | 0.55283132 |
| OS02G0643200 | Transcription factor, DNA-binding intermediate protein for SLR1, Modulation of gibberellin signaling pathway, Regulation of pla | 0.18119162 | -0.1367529 | 0.18733703 | #N/A | #N/A |
| OS04G0555700 | Similar to Actin-depolymerizing factor (ADF). (Os04t0555700-01) | 0.28879559 | -0.0592529 | #N/A | #N/A | #N/A |
| OS12G0184300 | Conserved hypothetical protein. (Os12t0184300-00) | 0.40321367 | 0.41348023 | #N/A | #N/A | #N/A |
| OS08G0189900 | Germin-like protein 8-10, Disease resistance (Os08t0189900-01) | 0.422387 | 0.4317616 | #N/A | #N/A | #N/A |
| OS05G0189300 | Vegetative storage protein/acid phosphatase domain containing protein. (Os05t0189300-01) | 0.448941 | 0.73474 | -0.891964 | -1.22439 | -1.41964 |
| OS08G0193900 | Cyclin-like F-box domain containing protein. (Os08t0193900-01) | 0.47830999 | 0.48848541 | #N/A | #N/A | #N/A |
| OS08G0269700 | Conserved hypothetical protein. (Os08t0269700-01) | 0.484704 | 0.698596 | -0.595735 | -0.786579 | -1.25541 |
| OS07G0674200 | Similar to 60S ribosomal protein L22-2. (Os07t0674200-01) | 0.489124 | 0.723858 | #N/A | -0.944056 | -1.10535 |
| OS02G0662100 | Similar to Tfm5 protein. (Os02t0662100-01) | 0.492094 | 0.663097 | -0.842415 | -1.21148 | -1.4763 |
| OS07G0562700 | Similar to Type III chlorophyll a/b-binding protein (Fragment). (Os07t0562700-01);Similar to Type III chlorophyll a/b-binding pro | 0.494644 | 0.683138 | #N/A | -0.691918 | -0.726806 |
| OS01G0662800 | Zinc finger, NHR/GATA-type domain containing protein. (Os01t0662800-00) | 0.50377614 | 0.30124815 | #N/A | #N/A | #N/A |
| OS01G0773700 | Similar to Photosystem II reaction center W protein (PSII 6.1 kDa protein) (Fragment). (Os01t0773700-02) | 0.50745 | 0.801365 | -0.593809 | #N/A | -0.789746 |
| OS01G0823300 | Similar to Ribosomal protein S26. (Os01t0823300-00) | 0.519685 | 0.514862 | #N/A | -0.860179 | -1.12385 |
| OS10G0501700 | Similar to Protein Rf1, mitochondrial. (Os10t0501700-00) | 0.5222025 | 0.53212791 | -2.697 | #N/A | #N/A |
| OS07G0517100 | Heat shock protein Hsp20 domain containing protein. (Os07t0517100-01);Heat shock protein Hsp20 domain containing protein. | 0.522943 | 0.898191 | -1.17811 | -1.37446 | -1.25026 |
| OS02G0622300 | Ribosomal protein L14 family protein. (Os02t0622300-00) | 0.527659 | 0.507994 | #N/A | -0.914984 | -1.07835 |

| | | | | | | |
|---|---|---|---|---|---|---|
| OS03G0592500 | Similar to Photosystem II type II chlorophyll a/b binding protein (Fragment). (Os03t0592500-01);Similar to Chloroplast chloroph | 0.532513 | 0.636869 | -0.73947 | -0.696443 | -0.836746 |
| OS06G0218600 | Cupredoxin domain containing protein. (Os06t0218600-01) | 0.535155 | 0.604812 | #N/A | #N/A | #N/A |
| OS04G0376000 | Similar to 60S ribosomal protein L35-1. (Os04t0376000-01) | 0.540512 | 0.620688 | #N/A | -0.933021 | -1.07593 |
| OS02G0192700 | Similar to Thioredoxin peroxidase. (Os02t0192700-02) | 0.548946 | 0.513254 | -1.35994 | -1.26062 | -0.994836 |
| OS01G0368900 | Similar to Glutaredoxin-C1. (Os01t0368900-01) | 0.5544 | 0.826205 | #N/A | #N/A | -0.912837 |
| OS07G0107300 | Plant disease resistance response protein family protein. (Os07t0107300-01) | 0.555435 | 0.672515 | -0.599147 | -0.709424 | -0.872025 |
| OS03G0781501 | Non-protein coding transcript. (Os03t0781501-01) | 0.556679 | 0.796727 | #N/A | -0.68435 | -1.00988 |
| OS01G0971400 | Similar to Cysteine proteinase. (Os01t0971400-01) | 0.559018 | 0.587433 | #N/A | #N/A | #N/A |
| OS05G0557000 | Similar to 60S ribosomal protein L37a. (Os05t0557000-01) | 0.559738 | 0.540748 | #N/A | -0.728939 | -0.925585 |
| OS05G0355500 | Similar to Ribosomal protein L29. (Os05t0355500-01) | 0.562618 | 0.698799 | -0.648071 | -0.721664 | -0.894347 |
| OS07G0148900 | Photosystem I protein-like protein. (Os07t0148900-01) | 0.563046 | 0.81963 | #N/A | #N/A | -0.69801 |
| OS03G0761500 | Similar to Subtilisin-like protease (Fragment). (Os03t0761500-01);Similar to Subtilisin-like protease (Fragment). (Os03t0761500- | 0.576153 | 0.613913 | #N/A | -0.899407 | -0.744143 |
| OS04G0598200 | Similar to 60S ribosomal protein L12. (Os04t0598200-00) | 0.576961 | 0.634028 | -0.717597 | -1.01611 | -0.921329 |
| OS12G0189400 | Similar to Photosystem I reaction centre subunit N, chloroplast precursor (PSI- N). (Os12t0189400-01) | 0.57803 | 0.571523 | -0.523258 | -0.607285 | #N/A |
| OS07G0139600 | Non-protein coding transcript. (Os07t0139600-01) | 0.583311 | 1.00916 | #N/A | #N/A | -0.890343 |
| OS09G0538400 | Similar to P-type R2R3 Myb protein (Fragment). (Os09t0538400-01) | 0.587941 | 0.716927 | #N/A | -0.765493 | -0.623427 |
| OS03G0823400 | Similar to Bowman-Birk type trypsin inhibitor (WTI). (Os03t0823400-01) | 0.592522 | 0.792957 | 1.04712 | #N/A | #N/A |
| OS03G0333400 | Similar to photosystem II 11 kD protein. (Os03t0333400-01) | 0.597486 | 0.87708 | -0.810503 | -0.652998 | -0.782845 |
| OS06G0181566 | Similar to 60S ribosomal protein L39. (Os06t0181566-01) | 0.603801 | 0.68054 | #N/A | #N/A | -0.866767 |
| OS02G0284600 | Similar to 60S ribosomal protein L27. (Os02t0284600-01) | 0.604226 | 0.619508 | -0.447259 | -0.906152 | -1.03724 |
| OS08G0129800 | Hypothetical conserved gene. (Os08t0129800-01);Hypothetical conserved gene. (Os08t0129800-02) | 0.611672 | 0.756797 | #N/A | #N/A | #N/A |
| OS04G0517000 | Similar to OSIGBa0145M07.3 protein. (Os04t0517000-01) | 0.611932 | 0.625338 | #N/A | -0.695249 | -0.851231 |
| OS05G0408900 | Similar to 1-deoxy-D-xylulose-5-phosphate synthase. (Os05t0408900-01);Similar to 1-deoxy-D-xylulose-5-phosphate synthase. ( | 0.612262 | 0.577002 | #N/A | #N/A | #N/A |
| OS03G0811800 | Ribosomal protein L36 family protein. (Os03t0811800-01) | 0.621353 | 0.801059 | #N/A | #N/A | #N/A |
| OS07G0110000 | Conserved hypothetical protein. (Os07t0110000-01);Hypothetical conserved gene. (Os07t0110000-02) | 0.624679 | 0.723475 | 0.494406 | #N/A | #N/A |
| OS02G0496900 | Mitochondrial import receptor, TOM9-2 subunit, plant domain containing protein. (Os02t0496900-01) | 0.628468 | 0.720024 | #N/A | -0.625369 | -0.983283 |
| OS04G0692600 | Staygreen protein domain containing protein. (Os04t0692600-01);Staygreen protein domain containing protein. (Os04t069260 | 0.632711 | 0.670645 | #N/A | #N/A | #N/A |
| OS03G0249400 | Similar to 40S ribosomal protein S20 (S22) (Fragment). (Os03t0249400-01) | 0.634139 | 0.641374 | -0.910097 | -1.04121 | -1.52669 |
| OS09G0132900 | Lipase, GDSL domain containing protein. (Os09t0132900-01);Similar to anther-specific proline-rich protein APG. (Os09t0132900 | 0.636208 | 0.817516 | #N/A | -0.76307 | -1.22117 |
| OS03G0335200 | Similar to WRKY1 (WRKY transcription factor 17). (Os03t0335200-01) | 0.63893913 | 0.64719177 | #N/A | -0.961724 | #N/A |
| OS03G0782200 | Hypothetical protein. (Os03t0782200-01) | 0.649096 | 0.623305 | -1.31612 | #N/A | #N/A |
| OS07G0142300 | Similar to Mucin-2. (Os07t0142300-01) | 0.651778 | 0.670183 | -1.98842 | -2.38192 | -2.07945 |
| OS08G0559200 | Similar to Ribosomal protein S25 (40S ribosomal 25S subunit). (Os08t0559200-01) | 0.651842 | 0.600456 | -0.708567 | -0.910178 | -0.899437 |
| OS01G0263300 | Similar to Peroxidase 72 precursor (EC 1.11.1.7) (Atperox P72) (PRXR8) (ATP6a). (Os01t0263300-01) | 0.652288 | 0.630251 | -0.466361 | -0.820195 | -1.26262 |
| OS11G0671000 | Similar to Dormancy-associated protein. (Os11t0671000-01) | 0.661 | 0.726715 | 1.45956 | #N/A | 0.653916 |
| OS10G0539500 | Similar to Histone H4. (Os10t0539500-01) | 0.665312 | 0.728453 | 0.536544 | #N/A | #N/A |
| OS01G0222600 | Conserved hypothetical protein. (Os01t0222600-01) | 0.665332 | 0.810567 | 0.753913 | #N/A | #N/A |
| OS05G0459900 | Similar to 60S ribosomal protein L36-1. (Os05t0459900-01);Similar to 60S ribosomal protein L36. (Os05t0459900-02) | 0.670066 | 0.770324 | #N/A | -0.91302 | -1.12078 |
| OS09G0325220 | Conserved hypothetical protein. (Os09t0325220-00) | 0.67276793 | 0.68259636 | #N/A | #N/A | #N/A |
| OS02G0731600 | Similar to Threonine endopeptidase. (Os02t0731600-01) | 0.67428 | 0.760973 | -0.750225 | -0.863981 | -1.21649 |
| OS02G0658800 | Beta-expansin. (Os02t0658800-01) | 0.675698 | 0.738119 | #N/A | #N/A | #N/A |
| OS07G0489100 | Similar to male sterility protein 2. (Os07t0489100-00) | 0.68032195 | 0.68978321 | #N/A | #N/A | #N/A |
| OS01G0823100 | Alpha-expansin OsEXPA2. (Os01t0823100-01) | 0.683242 | 0.676259 | #N/A | #N/A | -0.773792 |
| OS01G0215700 | Esterase, SGNH hydrolase-type domain containing protein. (Os01t0215700-01);Esterase, SGNH hydrolase-type domain containi | 0.686627 | 0.710071 | #N/A | #N/A | #N/A |
| OS06G0553200 | Similar to Meiosis 5. (Os06t0553200-01) | 0.687683 | 0.864282 | #N/A | -1.02926 | -1.27511 |

| | | | | | | |
|---|---|---|---|---|---|---|
| OS12G0174300 | Conserved hypothetical protein. (Os12t0174300-00) | 0.68826989 | 0.50269721 | -0.3465382 | #N/A | #N/A |
| OS11G0151300 | Similar to 60S ribosomal protein L26-1. (Os11t0151300-01) | 0.689564 | 0.578043 | -0.493946 | -0.992094 | -1.0358 |
| OS01G0914300 | Plant lipid transfer protein/seed storage/trypsin-alpha amylase inhibitor domain containing protein. (Os01t0914300-01) | 0.699541 | 0.861422 | #N/A | #N/A | -0.801524 |
| OS03G0819400 | Heavy metal transport/detoxification protein domain containing protein. (Os03t0819400-01) | 0.706591 | 0.64653 | #N/A | -0.569353 | -0.614627 |
| OS01G0938100 | Photosystem II protein Psb28, class 1 domain containing protein. (Os01t0938100-01) | 0.710852 | 0.67308 | -1.40905 | -1.07569 | -0.985426 |
| OS12G0530000 | Similar to Histone H2A. (Os12t0530000-01) | 0.714742 | 0.746856 | #N/A | -0.961681 | -0.989674 |
| OS11G0703600 | Conserved hypothetical protein. (Os11t0703600-01);Conserved hypothetical protein. (Os11t0703600-02);Hypothetical conserve | 0.716101 | 0.699773 | #N/A | -0.739286 | #N/A |
| OS11G0474100 | Conserved hypothetical protein. (Os11t0474100-01) | 0.718228 | 0.842096 | #N/A | #N/A | #N/A |
| OS11G0433900 | Similar to 60S ribosomal protein L38. (Os11t0433900-01) | 0.718361 | 0.591831 | -0.762627 | -1.12109 | -1.26516 |
| OS02G0815000 | Similar to 60S ribosomal protein L37 (G1.16). (Os02t0815000-00) | 0.732406 | 0.844213 | #N/A | -0.81101 | -1.07735 |
| OS04G0545600 | Similar to Uclacyanin 3-like protein. (Os04t0545600-01) | 0.73243 | 0.761197 | #N/A | -1.24912 | #N/A |
| OS04G0583600 | Similar to Histone H4. (Os04t0583600-01) | 0.735617 | 0.64292 | #N/A | -0.72332 | -0.924554 |
| OS02G0102700 | Similar to AGL157Cp. (Os02t0102700-01);Similar to AGL157Cp. (Os02t0102700-02) | 0.737198 | 0.768152 | #N/A | #N/A | -0.769693 |
| OS05G0375400 | Beta-glucanase precursor. (Os05t0375400-01) | 0.7401 | 0.988123 | 1.51927 | #N/A | #N/A |
| OS10G0505500 | Similar to Nonspecific lipid-transfer protein 2G (LTP2G) (Lipid transfer protein 2 isoform 1) (LTP2-1) (7 kDa lipid transfer protein | 0.741191 | 0.654005 | #N/A | -1.21993 | -1.61748 |
| OS03G0279200 | Similar to Histone H2A. (Os03t0279200-01) | 0.743443 | 0.644973 | #N/A | -0.896062 | -0.946367 |
| OS08G0490900 | Similar to Histone H2B.2. (Os08t0490900-00) | 0.747691 | 0.650448 | #N/A | -0.781975 | -1.05034 |
| OS03G0765900 | PetM of cytochrome b6/f complex subunit 7 domain containing protein. (Os03t0765900-01) | 0.747738 | 0.810235 | -1.20954 | -1.06992 | -0.985389 |
| OS01G0191200 | Similar to Acid phosphatase. (Os01t0191200-01);Similar to Acid phosphatase. (Os01t0191200-02) | 0.766207 | 0.812889 | #N/A | #N/A | #N/A |
| OS02G0264800 | Protein of unknown function DUF1070 family protein. (Os02t0264800-01) | 0.790016 | 0.869118 | #N/A | -0.950482 | -1.35076 |
| OS12G0576600 | Metallophosphoesterase domain containing protein. (Os12t0576600-01);Metallophosphoesterase domain containing protein. ( | 0.790021 | 0.819809 | 0.816691 | #N/A | #N/A |
| OS06G0701400 | Similar to 60S acidic ribosomal protein P3 (P1/P2-like) (P3A). (Os06t0701400-01) | 0.793004 | 0.599186 | -0.790873 | -0.933372 | -0.684718 |
| OS03G0162200 | Similar to Histone H2A. (Os03t0162200-01) | 0.796622 | 0.676322 | #N/A | -0.717768 | -1.22919 |
| OS04G0657900 | Similar to Fortune-1. (Os04t0657900-01) | 0.806431 | 1.09676 | #N/A | #N/A | #N/A |
| OS06G0159450 | Similar to Histone H3. (Os06t0159450-00) | 0.808603 | 0.6588 | 0.498683 | -0.704014 | -0.998244 |
| OS05G0462000 | Similar to plant-specific domain TIGR01589 family protein. (Os05t0462000-00) | 0.812326 | 1.10381 | -0.960341 | -1.74219 | -1.94493 |
| OS06G0571800 | Similar to GATA transcription factor 20. (Os06t0571800-01);Similar to GATA transcription factor 3 (AtGATA-3). (Os06t0571800-0 | 0.815547 | 0.8426 | #N/A | #N/A | #N/A |
| OS03G0108300 | Similar to xyloglucan endotransglucosylase/hydrolase protein 32. (Os03t0108300-01);Similar to Cellulase (EC 3.2.1.4). (Os03t010 | 0.816465 | 0.711388 | #N/A | -0.552949 | #N/A |
| OS05G0441400 | Protein of unknown function DUF623, plant domain containing protein. (Os05t0441400-01) | 0.816763 | 0.835482 | #N/A | -0.730938 | #N/A |
| OS02G0771600 | Similar to 1-aminocyclopropane-1-carboxylate oxidase (Fragment). (Os02t0771600-01);ACC oxidase, Ethylene biosynthesis (Os0 | 0.824911 | 0.652782 | #N/A | #N/A | #N/A |
| OS06G0132400 | Similar to Magnesium-protoporphyrin O-methyltransferase (EC 2.1.1.11) (Magnesium- protoporphyrin IX methyltransferase). (O | 0.84241 | 0.865962 | -0.987764 | -1.16873 | #N/A |
| OS03G0648600 | Similar to Zinc finger, C3HC4 type family protein, expressed. (Os03t0648600-00) | 0.84427667 | 0.8533526 | 0.70621754 | #N/A | #N/A |
| OS06G0320500 | Similar to Light-harvesting complex I (Fragment). (Os06t0320500-01) | 0.84684 | 0.7884 | #N/A | -0.725207 | #N/A |
| OS03G0119900 | Similar to Histone H4. (Os03t0119900-01) | 0.852965 | 0.808073 | #N/A | -0.666442 | -1.04493 |
| OS12G0538300 | Dor1-like protein family protein. (Os12t0538300-01) | 0.864001 | 1.02108 | 1.0503 | #N/A | #N/A |
| OS04G0683800 | Similar to Phosphoribosyltransferase (Fragment). (Os04t0683800-01) | 0.866613 | 1.26346 | 1.14951 | #N/A | #N/A |
| OS05G0466600 | Similar to Histone H4. (Os05t0466600-01) | 0.883751 | 0.778505 | #N/A | -0.791321 | -0.974979 |
| OS06G0116900 | Conserved hypothetical protein. (Os06t0116900-01) | 0.93409 | 1.03716 | #N/A | -1.06087 | -0.918667 |
| OS10G0518000 | Non-protein coding transcript. (Os10t0518000-00) | 0.942234 | 1.24511 | #N/A | -1.62627 | -1.83595 |
| OS08G0126700 | Nucleotide-binding, alpha-beta plait domain containing protein. (Os08t0126700-01);RNA recognition motif domain domain cont | 0.957672 | 0.943693 | #N/A | #N/A | #N/A |
| OS10G0557900 | Peptidase M10A and M12B, matrixin and adamalysin family protein. (Os10t0557900-00) | 0.962098 | 0.77715 | #N/A | -1.00547 | #N/A |
| OS02G0629800 | Similar to Defensin precursor. (Os02t0629800-01) | 0.97225 | 0.594667 | #N/A | -1.67597 | -1.66745 |
| OS07G0564200 | Conserved hypothetical protein. (Os07t0564200-01) | 0.977173 | 1.04291 | #N/A | #N/A | #N/A |
| OS08G0444500 | Conserved hypothetical protein. (Os08t0444500-00) | 1.00593 | 1.05002 | #N/A | #N/A | #N/A |
| OS01G0720600 | Similar to Starch synthase IV. (Os01t0720600-01);Starch synthase, Starch biosynthesis (Os01t0720600-02);Similar to starch synt | 1.01251 | 0.920631 | 1.92485 | #N/A | #N/A |

| | | | | | | |
|---|---|---|---|---|---|---|
| OS06G0561000 | Myo-inositol oxygenase, Drought stress tolerance (Os06t0561000-01) | 1.05523 | 0.899457 | #N/A | -1.87632 | -1.61086 |
| OS04G0419600 | Histone H3. (Os04t0419600-01) | 1.05621 | 0.840063 | #N/A | -1.05883 | -1.74268 |
| OS01G0359600 | Disease resistance protein domain containing protein. (Os01t0359600-00) | 1.07619 | 1.37033 | 2.02992 | #N/A | #N/A |
| OS07G0175300 | Conserved hypothetical protein. (Os07t0175300-01) | 1.09506 | 0.833067 | #N/A | -1.34001 | -1.34332 |
| OS05G0419800 | Lipase, GDSL domain containing protein. (Os05t0419800-01) | 1.12088 | 1.18814 | 0.767618 | #N/A | #N/A |
| OS01G0342800 | Similar to UPF0631 protein. (Os01t0342800-01) | 1.12914 | 1.13065 | #N/A | #N/A | #N/A |
| OS02G0190500 | Serine/threonine protein kinase-related domain containing protein. (Os02t0190500-01) | 1.14598 | 0.813398 | #N/A | -0.96597 | -1.20549 |
| OS11G0155900 | Histone H3. (Os11t0155900-00) | 1.16102 | 1.16978 | #N/A | -1.63571 | -1.79906 |
| OS03G0187100 | Hypothetical conserved gene. (Os03t0187100-00) | 1.19786 | 1.2239 | 2.25002 | #N/A | #N/A |
| OS01G0532300 | Conserved hypothetical protein. (Os01t0532300-01) | 1.26999 | 1.11253 | #N/A | -0.997955 | #N/A |
| OS05G0500600 | GRAS transcription factor domain containing protein. (Os05t0500600-01) | 1.28314 | 1.11824 | -2.76276 | -2.92463 | -2.17042 |
| OS12G0438000 | Similar to Histone H2A. (Os12t0438000-02) | 1.28714 | 1.17771 | #N/A | -1.35346 | -1.23108 |
| OS07G0592000 | Gibberellin regulated protein family protein. (Os07t0592000-01) | 1.29721 | 1.43902 | 1.26012 | #N/A | #N/A |
| OS01G0603500 | Similar to Cf2/Cf5-like disease resistance protein (Fragment). (Os01t0603500-00) | 1.32175 | 1.43265 | #N/A | #N/A | #N/A |
| OS04G0644100 | Sterile alpha motif homology domain containing protein. (Os04t0644100-01) | 1.37378 | 1.11832 | #N/A | -0.945525 | #N/A |
| OS11G0118400 | Hypothetical protein. (Os11t0118400-01) | 1.47229 | 1.69889 | 1.8754 | #N/A | #N/A |
| OS01G0854000 | Similar to AER. (Os01t0854000-01) | 1.51237 | 1.47752 | 1.06165 | - | - |
| OS05G0521600 | Pectin lyase fold/virulence factor domain containing protein. (Os05t0521600-01) | 1.53012 | 1.44654 | - | -1.07623 | - |
| OS05G0112200 | Similar to Zn finger protein (Fragment). (Os05t0112200-01) | 1.53433 | 1.63274 | - | - | - |
| OS10G0178500 | UDP-glucuronosyl/UDP-glucosyltransferase family protein. (Os10t0178500-01) | 1.61314 | 2.39407 | - | - | - |
| OS10G0159300 | Conserved hypothetical protein. (Os10t0159300-01);Conserved hypothetical protein. (Os10t0159300-02) | 1.65535 | 2.38329 | 1.8465 | - | - |
| OS01G0392800 | Similar to DET1-like protein. (Os01t0392800-01) | 1.67482 | 1.59081 | 1.64322 | - | - |
| OS05G0463100 | Hypothetical gene. (Os05t0463100-01) | 1.68232 | 1.98664 | -1.30485 | -2.46887 | -2.47781 |
| OS01G0953400 | NB-ARC domain containing protein. (Os01t0953400-01) | 1.70267 | 1.75173 | 2.08317 | - | - |
| OS05G0320300 | Similar to Homeobox protein. (Os05t0320300-01) | 1.71161 | 1.30985 | 1.89632 | - | - |
| OS03G0707600 | DELLA repressor protein, Gibberellin signaling (Os03t0707600-01) | 1.74177 | 1.9669 | - | - | - |
| OS08G0484100 | Similar to predicted protein. (Os08t0484100-01) | 1.74906 | 1.59625 | - | -1.27372 | - |
| OS06G0295900 | Hypothetical conserved gene. (Os06t0295900-01) | 1.81893 | 1.36065 | 1.80531 | - | - |
| OS09G0538500 | Similar to Male sterility MS5. (Os09t0538500-01) | 1.87199 | 1.49755 | - | - | - |
| OS11G0679900 | Conserved hypothetical protein. (Os11t0679900-01) | 1.95114 | 2.2266 | 2.32974 | - | - |
| OS10G0446400 | AT-rich interaction region domain containing protein. (Os10t0446400-01);SANT domain, DNA binding domain containing protei | 2.07222 | 2.07384 | 1.49351 | - | - |
| OS01G0706700 | Protein of unknown function DUF590 family protein. (Os01t0706700-01) | 2.10425 | 2.26141 | 2.6151 | - | - |
| OS02G0528500 | Nucleic acid-binding, OB-fold domain containing protein. (Os02t0528500-01);Nucleic acid-binding, OB-fold-like domain containi | 2.13592 | 2.04303 | 1.92379 | - | - |
| OS03G0107000 | Similar to ASK20. (Os03t0107000-01) | 2.18268 | 1.98944 | 2.27648 | - | - |
| OS08G0389601 | Non-protein coding transcript. (Os08t0389601-01) | 3.50315 | 4.79083 | - | - | - |

| 7. SLR1-OX compa. to WT (mock) | | | | | |
|---|---|---|---|---|---|
| Identifier | Description | Nm.Sm | FC.NmNs | FC.SmSs | FC.SSmSSs |
| OS07G0162450 | Protein of unknown function DUF3778 domain containing protein. (Os07t0162450-01) | -5.65788 | 0.671612 | - | - |
| OS07G0122000 | Protein of unknown function DUF1719, Oryza sativa family protein. (Os07t0122000-01) | -5.07211 | - | - | - |
| OS06G0726100 | Similar to Seed chitinase-c. (Os06t0726100-01) | -3.29707 | 1.53334 | - | 3.49369 |
| OS07G0456500 | Hypothetical conserved gene. (Os07t0456500-01) | -3.24462 | - | - | - |
| OS10G0538200 | Peptidase aspartic, catalytic domain containing protein. (Os10t0538200-01) | -3.23145 | - | - | - |
| OS12G0431100 | ATPase, AAA-type, core domain containing protein. (Os12t0431100-01) | -3.1639 | -1.22113 | - | - |
| OS02G0269600 | Similar to Subtilase. (Os02t0269600-00) | -3.09277 | -2.78471 | - | - |
| OS02G0570400 | Similar to Ent-kaurene synthase 1A. (Os02t0570400-01) | -3.03213 | -0.84182 | 1.67986 | - |
| OS05G0149200 | PWWP domain containing protein. (Os05t0149200-01) | -3.00213 | - | 2.73117 | - |
| OS07G0513600 | Hypothetical conserved gene. (Os07t0513600-00) | -2.92858 | - | - | - |
| OS01G0597600 | Amino acid transporter, transmembrane domain containing protein. (Os01t0597600-01) | -2.90166 | - | - | - |
| OS02G0569400 | Cytochrome P450 family protein. (Os02t0569400-01);Similar to Cyt-P450 monooxygenase. (Os02t0569400-02) | -2.89963 | -0.881338 | - | - |
| OS11G0608300 | Barley stem rust resistance protein. (Os11t0608300-01) | -2.78507 | - | - | - |
| OS04G0654500 | Conserved hypothetical protein. (Os04t0654500-01) | -2.7822 | - | - | - |
| OS04G0179200 | Similar to Stem secoisolariciresinol dehydrogenase (Fragment). (Os04t0179200-01) | -2.77597 | -1.7745 | - | - |
| OS01G0113700 | Similar to Receptor-like kinase. (Os01t0113700-00) | -2.76318 | - | - | - |
| OS03G0838800 | Zinc finger, C2H2 domain containing protein. (Os03t0838800-00) | -2.70818 | - | - | - |
| OS03G0197200 | Similar to Sorbitol transporter. (Os03t0197200-01) | -2.69751 | - | - | - |
| OS12G0562400 | Similar to Phospholipase C (Fragment). (Os12t0562400-00) | -2.69162 | - | - | - |
| OS04G0178300 | Similar to Isoform 3 of Syn-copalyl diphosphate synthase. (Os04t0178300-01);Similar to Syn-copalyl diphosphate synthase | -2.58839 | -1.12275 | - | - |
| OS03G0410100 | Similar to cDNA clone:J033024M21, full insert sequence. (Os03t0410100-01) | -2.51992 | -2.07718 | - | - |
| OS12G0258700 | Cupredoxin domain containing protein. (Os12t0258700-01) | -2.47107 | - | 2.94202 | 4.0818 |
| OS02G0770800 | Similar to Nitrate reductase [NAD(P)H] (EC 1.7.1.2). (Os02t0770800-01) | -2.45145 | - | 2.36517 | 2.3244 |
| OS01G0585200 | Similar to cDNA clone:J023088C01, full insert sequence. (Os01t0585200-01) | -2.4032 | -1.75144 | - | - |
| OS04G0468000 | Similar to OSIGBa0128P10.7 protein. (Os04t0468000-01) | -2.40149 | - | - | - |
| OS03G0661600 | Similar to Alpha-amylase/trypsin inhibitor (Antifungal protein). (Os03t0661600-01) | -2.37404 | 0.89187 | - | 3.67793 |
| OS01G0650800 | Transposon, En/Spm-like domain containing protein. (Os01t0650800-01) | -2.36803 | - | - | - |
| OS05G0409500 | Similar to MtN21 protein. (Os05t0409500-00) | -2.35227 | - | - | - |
| OS12G0556500 | Calmodulin binding protein-like family protein. (Os12t0556500-01) | -2.33061 | -1.47294 | - | - |
| OS12G0284850 | Hypothetical gene. (Os12t0284850-01) | -2.2846 | - | 1.99911 | - |
| OS10G0439924 | Cytochrome P450 family protein. (Os10t0439924-01) | -2.25662 | - | - | - |
| OS03G0315400 | Similar to Typical P-type R2R3 Myb protein (Fragment). (Os03t0315400-01) | -2.255 | -1.90328 | 3.21363 | - |
| OS03G0832500 | Nucleic acid-binding, OB-fold domain containing protein. (Os03t0832500-01) | -2.21588 | - | - | - |
| OS08G0509100 | Similar to Lipoxygenase, chloroplast precursor (EC 1.13.11.12). (Os08t0509100-01);Similar to Lipoxygenase. (Os08t050910 | -2.20174 | -1.44623 | - | - |
| OS01G0775400 | Similar to Cytokinin dehydrogenase 5 precursor (EC 1.5.99.12) (Cytokinin oxidase 5) (CKO5) (AtCKX5) (AtCKX6). (Os01t077 | -2.18756 | -0.931208 | - | - |
| OS12G0573900 | Conserved hypothetical protein. (Os12t0573900-01) | -2.15097 | - | - | - |
| OS07G0121800 | Protein of unknown function DUF1719, Oryza sativa domain containing protein. (Os07t0121800-01) | -2.08706 | - | - | - |
| OS01G0740650 | Similar to glutamate carboxypeptidase 2. (Os01t0740650-00) | -2.07177 | -1.52519 | 1.96164 | - |
| OS09G0455300 | Similar to INDEHISCENT protein. (Os09t0455300-01) | -2.03054 | 1.36869 | 4.08938 | 3.28316 |

| | | | | | |
|---|---|---|---|---|---|
| OS02G0163500 | Conserved hypothetical protein. (Os02t0163500-01) | -1.98558 | - | - | - |
| OS04G0571600 | Multi antimicrobial extrusion protein MatE family protein. (Os04t0571600-01) | -1.93756 | - | - | - |
| OS08G0400200 | Similar to SET domain protein. (Os08t0400200-00) | -1.93455 | -2.011 | - | - |
| OS01G0605100 | Similar to BCS1 protein-like protein. (Os01t0605100-01) | -1.88607 | - | - | - |
| OS02G0649000 | Non-protein coding transcript. (Os02t0649000-01) | -1.87544 | - | - | - |
| **OS03G0327800** | **NAC Family transcriptional activator, Abiotic stress response, Positive regulator of leaf senescence (Os03t0327800-01)** | **-1.86983** | **-** | **-** | **-** |
| OS02G0129600 | Glycoside hydrolase, subgroup, catalytic core domain containing protein. (Os02t0129600-01) | -1.85462 | - | - | - |
| OS04G0401700 | Potassium transporter, Potassium mediated growth, Salt tolerance (Os04t0401700-01);Similar to Isoform 2 of Potassium t | -1.84784 | -1.99016 | 1.27954 | - |
| OS04G0178400 | Similar to Cytochrome P450 CYP99A1 (EC 1.14.-.-) (Fragment). (Os04t0178400-01) | -1.8415022 | -1.31089 | - | - |
| OS02G0570500 | Similar to Cytochrome P450 family protein, expressed. (Os02t0570500-01) | -1.82238 | - | - | - |
| OS12G0443800 | Hypothetical conserved gene. (Os12t0443800-01);Similar to ATP binding / ATPase/ nucleoside-triphosphatase/ nucleotide | -1.79528 | -1.49658 | - | - |
| OS10G0490100 | Virulence factor, pectin lyase fold family protein. (Os10t0490100-01) | -1.79131 | -1.21924 | - | - |
| OS06G0581500 | Protein kinase, core domain containing protein. (Os06t0581500-01) | -1.78138 | 1.53473 | - | - |
| OS12G0248600 | Hypothetical protein. (Os12t0248600-01) | -1.77414 | - | - | - |
| OS01G0613800 | Peptidase C1A, papain family protein. (Os01t0613800-01) | -1.77031 | - | - | - |
| OS04G0556000 | Heavy metal P-type ATPase, Xylem loading of copper (Os04t0556000-01);Similar to heavy metal ATPase. (Os04t0556000-( | -1.76321 | -1.19296 | - | - |
| OS10G0570200 | Similar to RIR1b protein precursor. (Os10t0570200-01) | -1.75629 | - | - | - |
| OS07G0543100 | Similar to Beta-amylase (EC 3.2.1.2). (Os07t0543100-00) | -1.75405 | -2.64647 | - | - |
| OS09G0280500 | Similar to Transcription factor HBP-1b(C38) (Fragment). (Os09t0280500-01) | -1.74974 | - | - | - |
| OS07G0563600 | FAR1 domain containing protein. (Os07t0563600-01) | -1.74246 | - | 1.66993 | - |
| OS01G0930400 | Potassium transporter, High-affinity K acquisition, Root-to-shoot K transport, K-regulated salt tolerance (Os01t0930400-0 | -1.73483 | -0.744179 | - | - |
| OS03G0743500 | Calcium-binding EF-hand domain containing protein. (Os03t0743500-01);Similar to Calmodulin 1 (Fragment). (Os03t0743! | -1.70468 | - | - | - |
| OS01G0627916 | Hypothetical protein. (Os01t0627916-00) | -1.70006 | - | - | - |
| OS07G0492100 | DNA helicase, UvrD/REP type domain containing protein. (Os07t0492100-01) | -1.69623 | - | 1.35789 | - |
| OS08G0224000 | NB-ARC domain containing protein. (Os08t0224000-01) | -1.68338 | - | 1.72301 | - |
| OS09G0472900 | Similar to Blight-associated protein p12 precursor. (Os09t0472900-01) | -1.68006 | 1.19631 | 1.30538 | - |
| OS03G0231400 | Hypothetical protein. (Os03t0231400-01) | -1.67782 | -2.39278 | 1.56956 | - |
| OS02G0260200 | Cyclin-like F-box domain containing protein. (Os02t0260200-01) | -1.66927 | -1.26019 | - | - |
| OS04G0586500 | Malectin-like carbohydrate-binding domain domain containing protein. (Os04t0586500-01) | -1.66744 | -0.989874 | - | - |
| OS02G0489500 | Conserved hypothetical protein. (Os02t0489500-01) | -1.66737 | -2.79436 | 1.38686 | - |
| OS05G0181700 | Conserved hypothetical protein. (Os05t0181700-01) | -1.655 | - | - | - |
| OS01G0944900 | Similar to Glucan endo-1,3-beta-D-glucosidase. (Os01t0944900-01) | -1.65184 | 1.288 | 2.07456 | 2.96158 |
| OS06G0554700 | Similar to kinesin motor protein-related. (Os06t0554700-01) | -1.64313 | -1.29628 | - | - |
| OS07G0631200 | Zinc finger, RING/FYVE/PHD-type domain containing protein. (Os07t0631200-01) | -1.64056 | -2.30361 | 1.14264 | - |
| OS01G0869200 | Magnesium transporter, Mg-mediated aluminum tolerance (Os01t0869200-01) | -1.61982 | -1.20327 | 1.82269 | - |
| OS05G0275100 | Pentatricopeptide repeat domain containing protein. (Os05t0275100-00) | -1.61567 | - | - | - |
| OS01G0562600 | Protein of unknown function DUF247, plant domain containing protein. (Os01t0562600-01) | -1.59715 | - | - | - |
| OS05G0358700 | Similar to predicted protein. (Os05t0358700-01) | -1.59584 | - | - | - |
| OS08G0290700 | Winged helix repressor DNA-binding domain containing protein. (Os08t0290700-01) | -1.59345 | - | 1.80851 | - |
| OS04G0430800 | Similar to OSIGBa0160I14.3 protein. (Os04t0430800-01);Similar to OSIGBa0160I14.3 protein. (Os04t0430800-02) | -1.56855 | - | - | - |
| OS04G0585700 | Protein of unknown function DUF581 family protein. (Os04t0585700-01) | -1.56654 | -2.02592 | 1.25769 | - |

| | | | | | |
|---|---|---|---|---|---|
| OS04G0480200 | Transporter, high affinity nitrate, Nar2 domain containing protein. (Os04t0480200-01) | -1.56087 | -1.78968 | - | - |
| OS01G0830100 | Pyridine nucleotide-disulphide oxidoreductase, NAD-binding region domain containing protein. (Os01t0830100-01);Simila | -1.54621 | - | - | - |
| OS08G0337100 | Conserved hypothetical protein. (Os08t0337100-01) | -1.53554 | - | 1.57593 | - |
| OS06G0586900 | Similar to Elicitor-inducible LRR receptor-like protein EILP. (Os06t0586900-01);Similar to Elicitor-inducible LRR receptor-lik | -1.52007 | - | - | - |
| OS10G0170300 | Alkaline-phosphatase-like, core domain domain containing protein. (Os10t0170300-01) | -1.51645 | - | - | - |
| OS04G0303100 | Similar to H0215E01.10 protein. (Os04t0303100-00) | -1.50846 | - | - | - |
| OS03G0223900 | UDP-3-O-acyl N-acetylglucosamine deacetylase domain containing protein. (Os03t0223900-01) | 1.51459 | - | - | - |
| OS06G0234600 | Domain of unknown function DUF231, plant domain containing protein. (Os06t0234600-01) | 1.53718 | - | - | - |
| OS01G0390600 | Pentatricopeptide repeat domain containing protein. (Os01t0390600-01) | 1.56097 | 1.39202 | - | - |
| OS09G0498700 | F-box domain, cyclin-like domain containing protein. (Os09t0498700-01) | 1.58906 | 1.97954 | - | - |
| OS03G0750300 | Conserved hypothetical protein. (Os03t0750300-00) | 1.59664 | 1.50426 | -1.06163 | - |
| OS07G0601100 | Similar to NADPH HC toxin reductase (Fragment). (Os07t0601100-01) | 1.61477 | 1.58824 | - | - |
| OS06G0585982 | Protein kinase, catalytic domain domain containing protein. (Os06t0585982-00) | 1.64033 | 2.27521 | - | - |
| OS11G0175500 | Zinc finger, RING/FYVE/PHD-type domain containing protein. (Os11t0175500-01);Zinc finger, RING/FYVE/PHD-type domai | 1.69032 | - | - | - |
| OS11G0552600 | Similar to Kinesin-like protein. (Os11t0552600-01) | 1.69507 | - | - | - |
| OS08G0157000 | Similar to Mitogen-activated protein kinase 4. (Os08t0157000-00) | 1.70907 | - | - | - |
| OS01G0238700 | Oligopeptide transporter OPT superfamily protein. (Os01t0238700-01) | 1.72731 | - | - | - |
| OS11G0687800 | NB-ARC domain containing protein. (Os11t0687800-01);NB-ARC domain containing protein. (Os11t0687800-02) | 1.79142 | 1.99332 | - | - |
| OS10G0556900 | Conserved hypothetical protein. (Os10t0556900-01) | 1.79535 | 2.05713 | - | - |
| OS10G0561200 | Similar to CigA protein. (Os10t0561200-01) | 1.81741 | - | - | - |
| OS11G0551900 | Hypothetical conserved gene. (Os11t0551900-01) | 1.81918 | 1.96644 | - | - |
| OS09G0554101 | Hypothetical conserved gene. (Os09t0554101-00) | 1.83543097 | - | - | - |
| OS03G0376000 | emp24/gp25L/p24 family protein. (Os03t0376000-01) | 1.84055 | - | - | - |
| OS11G0130600 | Conserved hypothetical protein. (Os11t0130600-01) | 1.85045 | - | - | - |
| OS08G0166400 | Zinc finger, PMZ-type domain containing protein. (Os08t0166400-01) | 1.90868 | 2.6338 | - | - |
| OS01G0506200 | Tetratricopeptide-like helical domain containing protein. (Os01t0506200-01) | 1.92572 | 2.16705 | - | - |
| OS03G0199500 | Conserved hypothetical protein. (Os03t0199500-01) | 1.92948 | - | -2.11664 | - |
| OS12G0167700 | Similar to SAP domain containing protein, expressed. (Os12t0167700-01) | 1.94055 | 1.97725 | - | - |
| OS03G0364700 | Transcription elongation factor, TFIIS/CRSP70, N-terminal domain containing protein. (Os03t0364700-01) | 1.99541 | - | - | - |
| OS01G0351200 | Similar to Poly. (Os01t0351200-00) | 2.04018 | - | - | - |
| OS07G0275475 | Hypothetical conserved gene. (Os07t0275475-00) | 2.22093 | - | - | - |
| OS12G0615100 | Similar to Protein kinase domain containing protein, expressed. (Os12t0615100-01);Hypothetical conserved gene. (Os12t0 | 2.27794 | - | - | - |
| OS06G0543601 | Protein of unknown function DUF3778 domain containing protein. (Os06t0543601-01) | 2.39839 | - | - | - |
| OS02G0620800 | Conserved hypothetical protein. (Os02t0620800-01) | 2.41909 | 3.06296 | - | - |
| OS12G0578300 | Calmodulin-binding, plant family protein. (Os12t0578300-01) | 2.44952 | - | - | 0.1306434 |
| OS11G0209100 | Cyclin-like F-box domain containing protein. (Os11t0209100-01) | 2.47482 | - | - | - |
| OS07G0167300 | Conserved hypothetical protein. (Os07t0167300-01) | 2.52397 | - | - | - |
| OS04G0524400 | Similar to OSIGBa0153E02-OSIGBa0093I20.13 protein. (Os04t0524400-01) | 2.67148 | 2.146 | - | - |

| 8. SUMO1SLR1-OX c. to WT (mock) | | | | | |
|---|---|---|---|---|---|
| Identifier | Description | FC.NmSSm | FC.NmNs | FC.SmSs | FC.SSmSSs |
| OS12G0117700 | Similar to RNA binding protein. (Os12t0117700-01) | -5.78643 | 0.571282 | - | - |
| OS01G0237500 | NmrA-like family protein. (Os01t0237500-01) | -4.3038319 | -3.25881 | - | - |
| OS02G0571900 | Cytochrome P450 family protein. (Os02t0571900-01) | -3.7111975 | -1.00862 | - | 0.23107599 |
| OS01G0186600 | Conserved hypothetical protein. (Os01t0186600-01) | -3.37321 | - | - | - |
| OS09G0471500 | Conserved hypothetical protein. (Os09t0471500-01) | -3.30853 | - | - | - |
| OS07G0489000 | Plant lipid transfer protein/Par allergen family protein. (Os07t0489000-01) | -3.2403052 | - | - | 1.97864228 |
| OS09G0255400 | Similar to Indole-3-glycerol phosphate synthase, chloroplast precursor (EC 4.1.1.48) (IGPS). (Os09t0255400-01);Similar to indole-3-glyc | -3.14574 | - | - | 2.22695 |
| OS07G0521100 | Leucine-rich repeat, typical subtype containing protein. (Os07t0521100-01) | -3.14274 | - | - | 2.52619 |
| OS02G0614966 | Leucine-rich repeat domain containing protein. (Os02t0614966-00) | -2.82784 | -1.0555 | - | - |
| OS12G0629700 | Similar to Thaumatin-like protein precursor. (Os12t0629700-01) | -2.5333 | 1.94203 | - | - |
| OS01G0959200 | Similar to Ci21A protein. (Os01t0959200-01) | -2.50483 | - | - | 2.64401 |
| OS02G0569000 | Cytochrome P450 family protein. (Os02t0569000-01) | -2.49274 | -2.28402 | - | - |
| OS09G0314200 | Disease resistance protein domain containing protein. (Os09t0314200-00) | -2.43886 | - | - | - |
| OS07G0256200 | RNA recognition motif, RNP-1 domain containing protein. (Os07t0256200-01) | -2.43564 | - | - | - |
| OS08G0233400 | Ankyrin repeat domain containing protein. (Os08t0233400-01) | -2.40801 | - | - | - |
| OS11G0173900 | Similar to Leucine Rich Repeat family protein, expressed. (Os11t0173900-00) | -2.40773 | -2.12453 | - | - |
| OS03G0795400 | Hypothetical conserved gene. (Os03t0795400-00) | -2.38033 | - | - | 3.18329 |
| OS07G0681600 | DNA helicase, ATP-dependent, RecQ type domain containing protein. (Os07t0681600-01) | -2.34365 | -1.67204 | - | - |
| OS10G0553300 | Similar to Trehalose-6-phosphate phosphatase. (Os10t0553300-01) | -2.22699 | - | - | - |
| OS02G0770100 | WD40 repeat domain containing protein. (Os02t0770100-01) | -2.21351 | - | - | 2.27666 |
| OS01G0793300 | Di-copper centre-containing domain containing protein. (Os01t0793300-01) | -2.10591 | 3.01992 | - | 2.33952 |
| OS01G0601625 | Leucine rich repeat, N-terminal domain containing protein. (Os01t0601625-00) | -2.09333 | - | - | 1.64138 |
| OS07G0120800 | Protein of unknown function DUF1719, Oryza sativa family protein. (Os07t0120800-01) | -2.07408 | - | - | - |
| OS05G0564000 | Conserved hypothetical protein. (Os05t0564000-00) | -2.07367 | - | - | - |
| OS12G0512700 | ABC transporter-like domain containing protein. (Os12t0512700-01) | -2.07193 | -1.0453 | - | - |
| OS11G0690466 | Hypothetical protein. (Os11t0690466-00) | -2.06844 | 1.16976 | - | - |
| OS08G0198900 | Conserved hypothetical protein. (Os08t0198900-01) | -2.0602 | - | - | 1.86525 |
| OS05G0365700 | Glycoside hydrolase, family 1 protein. (Os05t0365700-01) | -2.05809 | - | - | - |
| OS01G0961200 | Cystathionine beta-synthase, core domain containing protein. (Os01t0961200-01) | -2.05757 | - | - | - |
| OS08G0360300 | Calmodulin binding protein-like domain containing protein. (Os08t0360300-00) | -1.95595 | - | - | - |
| OS01G0892500 | Similar to carboxylic ester hydrolase. (Os01t0892500-01) | -1.92985 | -1.55186 | - | - |
| OS04G0296301 | Conserved hypothetical protein. (Os04t0296301-00) | -1.8974883 | - | - | - |
| OS02G0572000 | Hypothetical protein. (Os02t0572000-01) | -1.8728248 | - | - | 0.78937871 |
| OS02G0562600 | Similar to Seven trans-membrane protein (Fragment). (Os02t0562600-01) | -1.82641 | -1.59528 | - | - |
| OS02G0156600 | Hypothetical conserved gene. (Os02t0156600-00) | -1.81545 | - | - | - |
| OS01G0890600 | Protein kinase, catalytic domain domain containing protein. (Os01t0890600-00) | -1.80444 | - | - | - |
| OS09G0491740 | Auxin efflux carrier domain containing protein. (Os09t0491740-01) | -1.79841 | - | - | 2.29867 |
| OS07G0273900 | Disease resistance protein domain containing protein. (Os07t0273900-01) | -1.79023 | - | - | 1.70851 |
| OS01G0226500 | Conserved hypothetical protein. (Os01t0226500-01);Similar to predicted protein. (Os01t0226500-02) | -1.77802 | - | - | 1.85126 |
| OS07G0283125 | Similar to lectin-like receptor kinase 7. (Os07t0283125-00) | -1.75574 | 0.862919 | - | - |
| OS11G0586001 | Putative protein phosphatase 2C 76. (Os11t0586001-00) | -1.75043 | -2.41758 | 2.32361 | - |

| | | | | | |
|---|---|---|---|---|---|
| OS03G0346800 | Similar to metal tolerance protein. (Os03t0346800-00) | -1.74312 | -1.21435 | - | - |
| OS04G0510700 | Similar to OSIGBa0157K09-H0214G12.12 protein. (Os04t0510700-01) | -1.73782 | - | - | - |
| OS08G0139700 | Similar to terpene synthase 6. (Os08t0139700-01) | -1.73635 | 0.592926 | - | - |
| OS04G0631800 | Similar to H0105C05.10 protein. (Os04t0631800-00) | -1.69726 | - | - | - |
| OS11G0490900 | Similar to WRKY transcription factor 72. (Os11t0490900-01) | -1.67428 | - | - | - |
| OS11G0566800 | Similar to Bibenzyl synthase (EC 2.3.1.-). (Os11t0566800-01) | -1.64164 | -1.51644 | - | - |
| OS08G0205100 | Disease resistance protein domain containing protein. (Os08t0205100-01) | -1.62425 | - | - | - |
| OS03G0668400 | Peptidase C12, ubiquitin carboxyl-terminal hydrolase 1 domain containing protein. (Os03t0668400-01) | -1.53237 | - | - | - |
| OS04G0370900 | Similar to H0607F01.5 protein. (Os04t0370900-01);Similar to cDNA, clone: J013067H13, full insert sequence. (Os04t0370900-02) | -1.52815 | - | - | - |
| OS01G0728300 | Similar to Cytochrome P450 monooxygenase CYP72A5 (Fragment). (Os01t0728300-01) | -1.50181 | - | - | - |
| OS07G0142100 | Conserved hypothetical protein. (Os07t0142100-00) | 1.47313 | -1.98056 | #N/A | -0.888735 |
| OS05G0275600 | Conserved hypothetical protein. (Os05t0275600-01);Conserved hypothetical protein. (Os05t0275600-02) | 1.52014 | 1.29551 | - | - |
| OS11G0296500 | Zinc finger, CCHC retroviral-type domain containing protein. (Os11t0296500-01) | 1.53363 | 1.5021 | - | - |
| OS01G0756200 | Similar to VirE2-interacting protein VIP1. (Os01t0756200-01) | 1.55823 | - | - | - |
| OS08G0536350 | Non-protein coding transcript. (Os08t0536350-01) | 1.56401 | - | - | - |
| OS12G0502700 | Heat shock protein DnaJ, N-terminal domain containing protein. (Os12t0502700-01) | 1.58237 | 1.31906 | - | - |
| OS01G0107700 | Similar to LIMONENE cyclase like protein. (Os01t0107700-01) | 1.58867 | 2.29053 | - | - |
| OS01G0884400 | Armadillo domain containing protein. (Os01t0884400-01) | 1.65555 | 2.25273 | - | - |
| OS03G0168400 | Hypothetical conserved gene. (Os03t0168400-01) | 1.67549 | 1.22318 | - | - |
| OS05G0417100 | Chloroplast-targeted Deg protease protein, Chloroplast development and maintenance of PSII function under high temperatures (Os05 | 1.6977 | 2.24056 | - | - |
| OS06G0163701 | Non-protein coding transcript. (Os06t0163701-01) | 1.74495 | 2.15254 | - | - |
| OS07G0527000 | Similar to nodulin-like protein. (Os07t0527000-00) | 1.74535 | - | - | -1.40436 |
| OS02G0772300 | Acyl-CoA N-acyltransferase domain containing protein. (Os02t0772300-01) | 1.76586 | - | - | - |
| OS10G0419400 | Similar to SIPL. (Os10t0419400-01) | 1.76967 | 1.4013 | - | - |
| OS07G0296000 | Conserved hypothetical protein. (Os07t0296000-01);Conserved hypothetical protein. (Os07t0296000-02) | 1.80117 | 1.95232 | - | - |
| OS07G0523100 | Similar to 60S ribosomal protein L44. (Os07t0523100-00) | 1.83967747 | - | - | -1.8785096 |
| OS03G0198600 | Homeodomain-leucine zipper transcription factor, Regulation of panicle exsertion (Os03t0198600-01) | 1.90572 | 2.66495 | - | - |
| OS03G0307900 | Conserved hypothetical protein. (Os03t0307900-01) | 1.9193 | - | - | - |
| OS03G0240400 | Conserved hypothetical protein. (Os03t0240400-01) | 1.9279 | 2.30128 | - | - |
| OS12G0516800 | Hypothetical conserved gene. (Os12t0516800-00) | 1.97025 | - | - | - |
| **OS09G0471800** | **Similar to WAK80 - OsWAK receptor-like protein kinase. (Os09t0471800-01)** | **2.09613** | **2.77566** | **1.24725** | - |
| OS02G0804000 | Non-protein coding transcript. (Os02t0804000-01);Non-protein coding transcript. (Os02t0804000-02) | 2.10536 | 1.55279 | - | - |
| **OS11G0454300** | **Similar to Water-stress inducible protein RAB21. (Os11t0454300-01)** | **2.17294** | **6.76968** | **7.65379** | **6.27587** |
| OS03G0795200 | Pentatricopeptide repeat domain containing protein. (Os03t0795200-01) | 2.26289 | 1.25639 | - | - |
| OS02G0167200 | Pentatricopeptide repeat domain containing protein. (Os02t0167200-01) | 2.50367 | 2.03867 | - | - |
| OS02G0630650 | Conserved hypothetical protein. (Os02t0630650-01) | 2.65767201 | - | - | - |
| OS05G0455900 | Pentatricopeptide repeat domain containing protein. (Os05t0455900-01) | 2.85672 | 3.23144 | - | - |
| OS04G0168300 | Conserved hypothetical protein. (Os04t0168300-01) | 3.43195 | - | - | - |

**Supplemental Table S2**. Expression changes of selected putative SLR1 TF interactors **(A)** and the respective downstream target genes **(B)**. Fold-change shown as $log_2$ FC. Comparison of condition A with condition B corresponding to respective selected analysis in Cufflinks (FPKM1 x FPKM2, for condition A and B, respectively). Nipp-Nipponbare; (m)-mock; (s)-stress

**A**

| Gene | Gene ID | Comparison | FPKM1 | FPKM2 | log2 FC | p value | q value |
|---|---|---|---|---|---|---|---|
| bHLH089 | Os03g0802900 | (no variation) | - | - | - | - | - |
| bHLH094 | Os07g0193800 | (no variation) | - | - | - | - | - |
| YABBY4 | Os02g0643200 | Nipp(m) x SLR1-OX(m) | 1 | 1,13382 | 0,18119162 | 5,00E-05 | 0,00322046 |
| | | Nipp(m) x SUMO1SLR1-OX(m) | 1 | 0,909564 | -0,1367529 | 5,00E-05 | 0,00413016 |
| OSH1 | Os03g0727000 | (no variation) | - | - | - | - | - |
| BZIP23 | Os02g0766700 | SUMO1SLR1-OX(m) x SUMO1SLR1-OX(s) | 5,65916 | 19,0771 | 1,75318 | 5,00E-05 | 0,00161363 |
| | | SLR1-OX(m) x SLR1-OX(s) | 5,15858 | 22,9713 | 2,15479 | 5,00E-05 | 0,00145967 |

**B**

| TF | Gene | Gene ID | Comparison | FPKM1 | FPKM2 | log2 FC | p value | q value |
|---|---|---|---|---|---|---|---|---|
| **bHLH089/094** | JAZ8 | Os09g0439200 | Nipp(m) x SLR1-OX(m) | 6,9285 | 3,14158 | -1,14105 | 0,00175 | 0,04149 |
| | PAL | Os05g0427400 | Nipp(m) x SLR1-OX(m) | 19,9094 | 9,64515 | -1,04557 | 0,00025 | 0,0109921 |
| | | | Nipp(m) x SUMOsLR1-OX(m) | 19,7858 | 7,90293 | -1,32401 | 5,00E-05 | 0,00413016 |
| | | | Nipp(m) x Nipp(s) | 21,8887 | 12,0094 | -0,866015 | 4,00E-04 | 0,00401929 |
| | JAMYB | Os11g0684000 | Nipp(m) x Nipp(s) | 4,66016 | 28,9139 | 2,63331 | 5,00E-05 | 0,0006964 |
| | | | SUMO1SLR1-OX(m) x SUMO1SLR1-OX(s) | 4,38065 | 10,2144 | 1,22139 | 0,00095 | 0,0174526 |
| **YABBY4/OSH1** | GA20ox2 | Os01g0883800 | SLR1-OX(m) x SLR1-OX(s) | 6,48556 | 15,9503 | 1,29828 | 0,00205 | 0,0289236 |
| | | | SUMO1SLR1-OX(m) x SUMO1SLR1-OX(s) | 4,93558 | 20,1065 | 2,02637 | 0,00035 | 0,00823851 |
| **BZIP23** | LEA | Os01g0705200 | Nipp(m) x Nipp(s) | 0,415116 | 9,4774 | 4,5129 | 0,0052 | 0,0303399 |
| | | | SLR1-OX(m) x SLR1-OX(s) | 0,294715 | 89,7753 | 8,25085 | 0,00415 | 0,0482317 |
| | BURP3 | Os01g0733500 | Nipp(m) x Nipp(s) | 174,1 | 266,382 | 0,613583 | 0,00015 | 0,0018036 |
| | | | SLR1-OX(m) x SLR1-OX(s) | 163,172 | 273,174 | 0,743428 | 5,00E-05 | 0,00145967 |
| | | | SUMO1SLR1-OX(m) x SUMO1SLR1-OX(s) | 205,282 | 361,207 | 0,81522 | 5,00E-05 | 0,00161363 |
| | Rab16C | Os11g0454000 | Nipp(m) x Nipp(s) | 3,70507 | 27,4644 | 2,88999 | 5,00E-05 | 0,0006964 |
| | | | SLR1-OX(m) x SLR1-OX(s) | 2,0665 | 29,641 | 3,84233 | 5,00E-05 | 0,00145967 |
| | | | SUMO1SLR1-OX(m) x SUMO1SLR1-OX(s) | 3,75271 | 52,5847 | 3,80864 | 5,00E-05 | 0,00161363 |

**Supplemental Table S3.** RNA-seq read count for each of the sequenced libraries and respective legend

| | Nm1 | Nm2 | Nm3 | Ns1 | Ns2 | Ns3 | SSm1 | SSm2 | SSm3 | SSs1 | SSs2 | SSs3 | Sm1 | Sm2 | Sm3 | Ss1 | Ss2 | Ss3 |
|---|---|---|---|---|---|---|---|---|---|---|---|---|---|---|---|---|---|---|
| **Mapped** | 24402466 | 21142008 | 26942628 | 19737389 | 24149789 | 27448789 | 22895169 | 24499364 | 26507478 | 18174287 | 20291054 | 32336235 | 28584410 | 28037380 | 22495137 | 32622076 | 28311844 | 24890888 |
| **Multi-align** | 3599340 | 3362109 | 4231368 | 2849750 | 3202344 | 3801961 | 3371838 | 3665242 | 3904528 | 2893930 | 4535501 | 4629288 | 4182229 | 3825617 | 3440968 | 5241837 | 4293603 | 3489718 |
| **Unique** | 20803126 | 17779899 | 22711260 | 16887639 | 20947445 | 23646828 | 19523331 | 20834122 | 22602950 | 15280357 | 15755553 | 27706947 | 24402181 | 24211763 | 19054169 | 27380239 | 24018241 | 21401170 |

Legend:

| | |
|---|---|
| **N** | Nipponbare (WT) |
| **SS** | SUMO1SLR1-OX |
| **S** | SLR1-OX |
| **m** | Condition: mock |
| **s** | Condition: salt |
| **1,2,3** | Replicates |

**Supplemental Table S4.** List of primers used for gene cloning, site directed mutagenesis (SDM) and gene expression quantification (RT-qPCR). Details for the use of specific primers are given in the text.

| Primers | Sequence (5' -> 3') |
|---|---|
| OsSLR1-GW-Fw | GGGGACAAGTTTGTACAAAAAAGCAGGCTTAATGAAGCGCGAGTACCAAGA |
| OsSLR1-GW-Rv | GGGGACCACTTTGTACAAGAAAGCTGGGTTAGCGTCCCAAAAACTTTGC |
| OsSLR1-CDS-Fw | ATGAAGCGCGAGTACCAAGA |
| OsSLR1-CDS-Rv | AGCGTCCCAAAAACTTTGC |
| pET-Fw-EcoRI | ATAGAATTCATGAAGCGCGAGTACCAAGA |
| pET-Rv-PspOMI | ATATGGGCCCAGCGTCCCAAAAACTTTGC |
| SLR1t-STOP-Fw | GGCAGCACGTCGTAATCCTCATCGTCG |
| SLR1t-STOP-Rv | CGACGATGAGGATTACGACGTGCTGCC |
| SLR1t-K2R-Fw | GTACTCGCGCCTCATGAATTCGGATCCGC |
| SLR1t-K2R-Rv | CGCGGATCCGAATTCATGAGGCGCGAGTAC |
| SLR1t-K60R-Fw | GCTCCAGCCTCTGCGCGACGTCGG |
| SLR1t-K60R-Rv | CCGACGTCGCGCAGAGGCTGGAGC |
| OL-SLR1+SUMO1-3'-Fw | CGGGGACGAGATCGACGCCATGCTCCACCAGACTGGAGGCAAGCGCGAGTACCAAGAAGC |
| OL-SUMO1+SLR1-5'-Rv | CGCCGCCGCTGCTCCCGCCGGCTTCTTGGTACTCGCGCTTGCCTCCAGTCTGGTGGAGCAT |
| OsSUMO1-GW-Fw | GGGGACAAGTTTGTACAAAAAAGCAGGCTTAATGTCGGCCGCCGGGGAGGAGGA |
| OsUBQ10-Fw | TGGTCAGTAATCAGCCAGTTTGG |
| OsUBQ10-Rv | GCACCACAAATACTTGACGAACAG |
| OsUBC2-Fw | TTGCATTCTCTATTCCTGAGCA |
| OsUBC2-Rv | CAGGCAAATCTCACCTGTCTT |
| OsSLR1-Fw | GATCGGGCTTACGGTTCTCG |
| OsSLR1-Rv | GCTAGGAGGACCAAGGAACG |
| OsP5CS-Fw | AAGATGGAAGATTGGCTTTGGGCAG |
| OsP5CS-Rv | TCATGCCTCCTCTACCTACACGAGA |
| OsRab21-Fw | CACACCACAGCAAGAGCTAAGTG |
| OsRab21-Rv | TGGTGCTCCATCCTGCTTAAG |
| OsSPY-Fw | CATTGGCTGACCCACCTGAT |
| OsSPY-Rv | CAGCTTCTGGGGAAGGACTG |
| OsSEC-Fw | TGCCATGTTTCGCGATGTTG |
| OsSEC-Rv | CGATGGCATGGAAAGGTTGC |
